# Meiosis-specific genes play roles in ploidy reduction in *Cryptococcus neoformans* titan cells

**DOI:** 10.1101/2025.09.16.676405

**Authors:** Zhuyun Bian, Kayla Wilhoit, Julian Liber, Anushka Peer, Ziyan Xu, Paul Magwene, Sheng Sun, Joseph Heitman

## Abstract

*Cryptococcus neoformans* is a fungal pathogen of humans that causes life-threatening meningoencephalitis. During infection, enlarged, polyploid titan cells are produced that promote persistence in the host, in part by resisting phagocytosis; under stress conditions, such as exposure to the antifungal drug fluconazole, titan cells can produce aneuploid or diploid, drug-resistant daughter cells. However, the mechanism underlying this ploidy reduction remains poorly understood. Interestingly, meiosis related genes have been shown to be activated during *Cryptococcus* infection, leading us to hypothesize that the depolyploidization of *C. neoformans* titan cells may occur through a process resembling the ploidy reduction during meiosis. In this study, we show that titan cells developed from diploid strains predominantly produce diploid daughter cells with haploid daughters observed infrequently. We further demonstrate that meiosis-specific genes, including *DMC1* and *SPO11*, are critical for stable inheritance of a diploid genome in the daughter cells. Specifically, deletion of these genes in a heterozygous diploid background resulted in: 1) titan cells with a significantly reduced capacity to produce daughter cells; 2) increased phenotypic variation among daughter cells produced by the titan cells, including traits that could be relevant to cell growth and viability; and 3) daughter cells produced by the titan cells exhibiting high levels of loss of heterozygosity (LOH) and aneuploidy, suggesting elevated genome instability. Taken together, these findings demonstrate the importance of meiosis-specific genes in the ploidy reduction process of titan cells derived from a heterozygous diploid background in an important human fungal pathogen.

**SIGNIFICANCE:** Polyploid titan cells are central to *Cryptococcus neoformans* persistence during infection, yet how they reduce ploidy to generate daughter cells was unclear. In most eukaryotes, the meiosis-related proteins Spo11 and Dmc1 are best known for initiating (double-strand break formation) and executing (strand exchange) meiotic recombination, respectively. Here, we show that in *C. neoformans*, Dmc1 and Spo11 are critical for polyploid titan cells to produce viable daughter cells with reduced ploidy specifically in a heterozygous diploid context, ensuring genome stability without introducing detectable canonical meiotic recombination. Loss of either gene compromises daughter cell production and causes genome instability (LOH, aneuploidy) and phenotypic heterogeneity. These findings reveal a “para-meiosis” safeguard that preserves genomic configurations during titan cell depolyploidization in *C. neoformans*.

## INTRODUCTION

*Cryptococcus neoformans* is a globally distributed pathogen that causes meningoencephalitis predominantly in immunocompromised hosts, most notably people living with HIV/AIDS or immunosuppressed for organ transplant or during cancer chemotherapy (1, 2). A recent study revealed that cryptococcal disease causes ∼112,000 deaths and accounts for ∼19% of AIDS-related mortality worldwide each year, underscoring its disproportionate impact on advanced HIV disease (3). During infection, *C. neoformans* undergoes marked morphogenetic transitions including the production of titan cells and seed cells (4–9). While seed cells are small cells (∼5 µm in diameter) that facilitate dissemination to extrapulmonary organs, titan cells are enlarged polyploid cells (>10 µm) with thickened cell wall and capsule that contribute to persistence and virulence of *Cryptococcus* in part through their size and altered surface properties that inhibit phagocytosis (10–14). More specifically, titan cells are a stress-associated morphotype induced by hostile and environmental cues including nutrient limitation, elevated CO_2_, serum, and low cell density (7, 15, 16). Moreover, the formation of titan cells is governed by stress-responsive pathways (e.g., cAMP/PKA, Rim101) (17–20). These conditions mirror the lung niche and position titanization within a broader Polyploid-and-Stress framework. In addition to their inhibitory activity against phagocytes, titan cells tend to be more resistant to stresses including oxidative and nitrosative (H_2_O_2_ and NaNO_2_) challenge. Moreover, polyploid titan cells can bud off smaller drug-resistant aneuploid, or diploid daughter cells when exposed to the antifungal drug fluconazole, whereas under nutrient-replete conditions they produce haploid daughter cells (21, 22). However, the mechanisms and regulation of this ploidy reduction process remain poorly understood. Because titan cells arise under hostile stress conditions and generate stress-tolerant progeny, defining how their polyploid genomes are reduced is directly relevant to how polyploidy buffers and resolves stress during infection.

Sexual reproduction in *C. neoformans* can proceed via different cycles including **a** × α heterothallic sexual reproduction, unisexual reproduction, and pseudosexual reproduction (23–27). All three modes of sexual reproduction involve regulated ploidy transitions and a terminal meiotic ploidy reduction. In both **a** × α sexual and unisexual reproduction, haploid cells undergo plasmogamy and form dikaryotic hyphae, and the nuclei then fuse (karyogamy) within the basidium to generate a transient diploid nucleus that undergoes meiosis (28). During meiosis, programmed meiotic recombination shuffles alleles between parental homologs. In *C. neoformans*, an obligatory ∼0.94 crossovers per chromosome for offspring from unisexual crosses and ∼1.27 crossovers per chromosome for offspring from the **a** x α sexual crosses were predicted (29). The *DMC1* and *SPO11* genes are both highly conserved from yeasts to humans and usually regarded as meiosis-specific genes because of the extensive functional and expression evidence across eukaryotes. Spo11 initiates the recombination process by generating DNA double-strand breaks (DSBs) (30). It is required exclusively during meiosis, as *spo11* deletion mutants in fungi, plants, and animals exhibit complete failure of meiotic recombination while maintaining normal mitotic growth (31). The recombinase Dmc1 promotes a homology search and strand invasion to repair these DSBs and produce meiotic recombination (32).

Deletion mutants of *dmc1* show normal vegetative proliferation but arrest during meiotic prophase with unpaired homologs (32). In *Cryptococcus*, both genes are transcriptionally induced during sexual development and are dispensable for mitotic growth, allowing their classification as meiosis-specific factors (33). Proper homolog pairing and crossover formation ensure accurate reductional segregation, thereby achieving ploidy reduction and producing haploid basidiospores with genetic variation (34, 35).

In addition to the facts that meiosis involves a ploidy reduction process and meiosis-specific genes have also been observed to be upregulated or activated during ploidy reduction processes across diverse systems, whether and how these factors functionally drive depolyploidization has remained unclear. In human cancer models, polyploid cells generated after genotoxic stress or mitotic catastrophe up-regulate canonical meiotic components (e.g., *DMC1*, *REC8*, *SYCP* proteins, *SPO11*) as they revert to smaller, proliferative progeny, linking a “pro-meiotic” transcriptional state to depolyploidization (36). In a Notch-induced Drosophila tumor model, tumor growth relies on cycles of polyploidization and reduction, and comparative RNA-seq analyses show that DNA-damage response genes with meiotic roles are up-regulated and required for the ploidy-reduction division (37). Analogously, the human fungal pathogen *Candida albicans* undergoes a parasexual cycle in which diploid cells fuse to form tetraploids and then return toward diploidy via concerted (largely random) chromosome loss, rather than meiosis. This depolyploidization and homologous recombination require meiosis-toolkit proteins such as Spo11 and Rec8 (38, 39).

In *C. neoformans*, meiotic genes were shown to be transiently expressed during infection, and deletion of meiotic genes increases the proportion of large cells during infection and delays the ploidy reduction of these cells (22). However, it is not yet clear whether the depolyploidization of *C. neoformans* titan cells is a meiotic or meiotic-like process. Determining whether depolyploidization is a process resembling meiosis could be helpful to reveal how this pathogen stabilizes its genome under stress, generates viable progeny, and adapts to hostile environments. In this study, we utilized a heterozygous diploid strain to carry out a detailed analysis of genome dynamics and transitions during the depolyploidization of *C. neoformans* titan cells. We show that the meiosis-specific genes *DMC1* and *SPO11* are required for efficient production of viable daughter cells from polyploid titan cells containing genome-wide heterozygosity, establishing a direct, genetics-based link between meiosis factors and titan-cell ploidy reduction. Loss of the meiosis-specific genes *DMC1* or *SPO11* led to the production of titan daughter cells with significantly increased phenotypic and genotypic heterogeneity, associated with rampant genomic instability during ploidy reduction, including LOH and aneuploidy that were often associated with reduced colony size and possibly inviability. Despite the observed dependence on meiosis-specific genes for production of viable daughter cells, we detected no canonical meiotic recombination in daughter cells derived from the titan cells formed by the wild-type heterozygous diploid strains. Thus, our findings indicate that titan cell depolyploidization co-opts meiotic proteins not to drive crossover formation, but instead to stabilize chromosome segregation, thereby safeguarding genome integrity during this ploidy reduction program in *C. neoformans*.

## RESULTS

### Meiosis-specific genes *DMC1* and *SPO11* are required for ploidy reduction when heterozygous diploid titan cells produce daughter cells

In this study, we induced *C. neoformans* cells to form titan cells and isolated titan cells through micro-dissection for analysis. Because titan cells were dissected based on size, usually defined as >10 µm, it was essential to confirm that our size-based isolation approach enriches for polyploid titan cells rather than enlarged haploid or diploid cells that might be arrested in G1 phase. To achieve this, we first grew the diploid strain CnLC6683 (40) under titan cell inducing conditions, then filtered the cell culture through a 10 μm filter to collect enlarged cells. Cell diameter measurements revealed a clear shift in size distribution toward large cells, with the proportion of large cells (>10 μm) increasing from ∼12% in the unfiltered cell culture to ∼65% post-filtration (Figure 1A). Because propidium-iodide staining followed by flow cytometry has been well established to assess the cell ploidy in *C. neoformans* (13, 14), we next assessed DNA content of both pre- and post-filtration cell cultures by this approach. We observed a clear enrichment of cells with elevated DNA content (>4C) in the filtered cultures (∼43%) than the pre-filtration cultures (∼11%) (Figure 1A). The filtration experiment was repeated, and independent scatter plots of different PI intensity are presented in Supplemental Figure S1. Together, these data confirm that size-based isolation effectively enriches for polyploid titan cells, therefore ensuring that subsequent analyses reflect *bona fide* titan cell biology.

**Figure 1.**
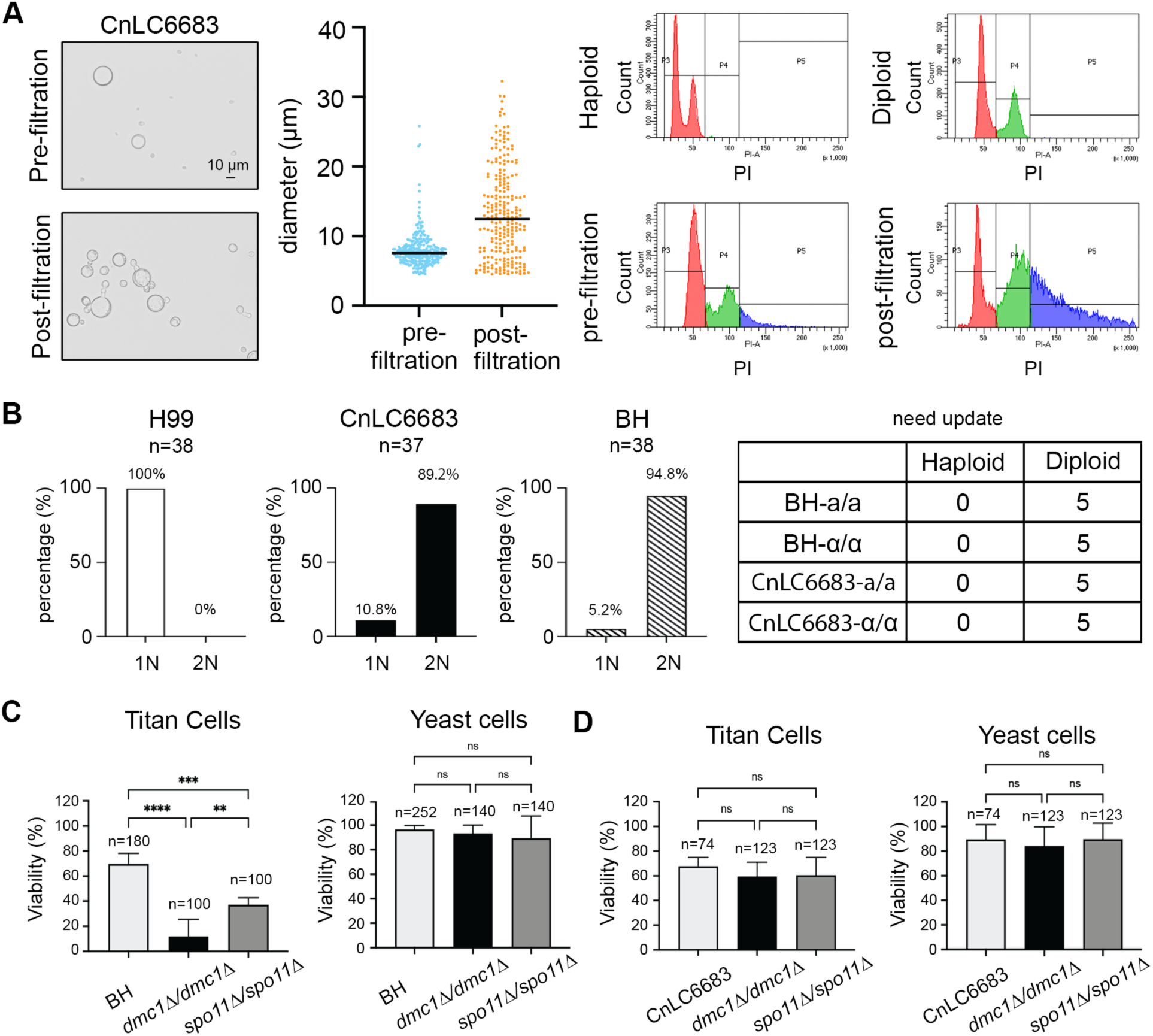
Investigating the role of *DMC1* and *SPO11* during ploidy reduction in different genetic backgrounds. **(A)** Pre- and post-filtration titan cell cultures of CnLC6683 were imaged by light microscopy and then used for cell diameter measurement. Cells collected from each culture were analyzed by flow cytometry for their ploidy. Post-filtration culture has increased proportion of larger polyploid cells. **(B)** Left panel: Ploidy of cells produced by the micro-dissected titan cells in the H99, CnLC6683, and BH genetic backgrounds were analyzed by flow cytometry. Percentages of haploid and diploid cells are presented. 1N: Haploid; 2N: Diploid. Right panel: Strains with different mating types were generated under BH and CnLC6683 background and used for titan cell induction and dissection. Ploidy of titan colony cells were analyzed by flow cytometry. **(C and D)** Percentage of dissected titan or yeast-like cells that can form colonies was assessed in the BH (C) and CnLC6683 (D) backgrounds.

The central question of this study was whether the mechanism by which polypoid titan cells produce haploid or aneuploid daughter cells resembled a meiotic process. Because meiotic recombination, a hallmark of meiosis, can only be tracked when homologous chromosomes carry distinguishable sequences, we constructed a heterozygous diploid strain BH by fusing the genetically distinct haploid strains Bt63**a** and H99α. While H99α belongs to the *C. neoformans* lineage VNI, Bt63**a** is a natural isolate that belongs to the lineage VNBI and exhibits ∼0.5% genetic divergence and one chromosomal translocation from H99α (23, 41). Previous studies on titan cells in *C. neoformans* mainly analyzed haploid strains and it has been reported that haploid-derived titan cells can produce haploid, aneuploid, or diploid daughter cells (7, 13, 21). To investigate these features in a diploid cell background, the BH strain was employed for titan cell induction. Once titan cells formed, we micro-dissected individual titan cells from the induction cell suspension and allowed them to grow on YPD agar media at 30°C for 48 hours. Colonies (titan colony cells) that formed on YPD agar media were then picked and analyzed by flow cytometry. Interestingly, for the CnLC6683 strain, a majority of the titan colony cells exhibited a diploid DNA content with only 10.8% being haploid cells (4 out of 38) (Figure 1B, left panel). Similarly, for the BH strain, only 5.2% titan colony cells exhibited a haploid DNA profile (2 out of 38) (Figure 1B, left panel). In contrast, when the haploid strain H99α was induced to form titan cells and then micro-dissected, all of the titan colony cells had a haploid DNA content (36 out of 36) (Figure 1B, left panel).

Because both the CnLC6683 and the BH strains are heterozygous at the *MAT* locus (**a**/α), to test whether the composition of *MAT* could influence the observed ploidy inheritance pattern, we generated *MAT*-homozygous (**a**/**a** and α/α) strains in both the BH and CnLC6683 backgrounds, and repeated the titan colony cell ploidy analysis. For the four tested strains, all of the micro-dissected titan cells (5 for each strain) formed colonies with cells that exhibited diploid DNA profiles (Figure 1B, right panel). Our data suggest that the ploidy levels of the cells produced by the titan cells correspond to those of the progenitor cells. We next investigated whether the meiosis-specific genes *DMC1* and *SPO11* are involved in the ploidy reduction processes when titan cells are forming colonies.

To this end, we first constructed a BH *dmc1*Δ/*dmc1*Δ homozygous deletion mutant by fusing two haploid *dmc1* deletion mutants (H99α *dmc1*Δ and Bt63**a** *dmc1*Δ). Similarly, we also generated a homozygous deletion mutant BH *spo11*Δ/*spo11*Δ. The two mutant strains, together with the BH strain, were analyzed for induction of titan cells, which were then micro-dissected onto YPD agar media, and allowed to form colonies. At least three biological replicates were performed for each genotype. We found that while approximately 70% of dissected titan cells formed by the BH wild-type strain gave rise to viable colonies (Figure 1C), the proportion of titan cells that formed viable colonies was significantly reduced in both the BH *dmc1*Δ/*dmc1*Δ (∼10%) and the BH *spo11*Δ/*spo11*Δ (∼40%) mutant strains (Figure 1C). We also micro-dissected normal sized yeast cells from the same cell cultures that were used for titan cell dissection, and all three genotypes displayed comparable colony-forming efficiencies of ∼90%, indicating that deletion of *DMC1* or *SPO11* does not compromise general cell viability under titan cell inducing conditions.

Because the BH strains are heterozygous diploids, we next assessed whether the requirement for Dmc1 and Spo11 in colony formation by titan cells produced by these strains is related to the heterozygosity present in their genomes. To this end, we conducted similar titan cell induction and dissection analyses with CnLC6683, a wild-type homozygous diploid strain, as well as *dmc1*Δ/*dmc1*Δ and *spo11*Δ/*spo11*Δ deletion mutants in the CnLC6683 background. In this case, we observed similarly high colony forming efficiency for all three genotypes in both titan cells (∼60%) and yeast-like cells (∼90%) (Figure 1D), suggesting the deletion of *DMC1* and *SPO11* does not have a significant effect on the titan cell colony forming ability in a homozygous diploid genetic background.

Taken together, these results demonstrate that the two meiosis-specific genes, *DMC1* and *SPO11*, are essential for the depolyploidization of titan cells produced by diploid cells when there is genome-wide heterozygosity, but they are dispensable if the progenitor strains are either haploid or homozygous diploid.

### Deletion of meiosis-specific genes leads to elevated phenotypic heterogeneity among cells produced by titan cells with genome-wide heterozygosity

Given the critical roles of *DMC1* and *SPO11* during the process of ploidy reduction and their canonical function in meiotic recombination that generates offspring diversity, we next investigated whether phenotypic variation is present among cells produced by the dissected titan cells. For this, we first performed carbon source utilization assays with Biolog YT microplates for three titan colony cells and their corresponding progenitor strains in each of the three genotypic backgrounds (BH wild-type, BH *dmc1*Δ/*dmc1*Δ, and BH *spo11*Δ/*spo11*Δ). This pilot screen not only revealed considerable variation in carbon source utilization among the cells produced from titan cells, but also allowed us to identify a subset of carbon sources that accounted for the most phenotypic variation among these cells, from which we chose D-ribose, L-arabinose, maltose, and sucrose for further single-carbon assays with an expanded set of titan colony cells (Supplemental Figure S2).

Interestingly, for each of the four carbon sources tested, cells produced by the BH *dmc1*Δ/*dmc1*Δ and BH *spo11*Δ/*spo11*Δ titan cells consistently showed significantly higher phenotypic variation, manifested as wider spread of the phenotypic values than those produced by their corresponding BH wild-type strains, indicating greater heterogeneity in carbon source utilization among cells produced by BH mutant titan cells (Figure 2A and Supplemental Figure S3A, Levene’s test). In contrast, when the same assays were performed for the cells produced by titan cells formed by CnLC6683 wild-type, CnLC6683 *dmc1*Δ/*dmc1*Δ, and CnLC6683 *spo11*Δ/*spo11*Δ strains, we observed significantly less difference in the levels of phenotypic variation among cells in the different genetic backgrounds compared to what was observed among the strains in the BH background (Figure 2B and Supplemental Figure S2B, Levene’s test).

**Figure 2.**
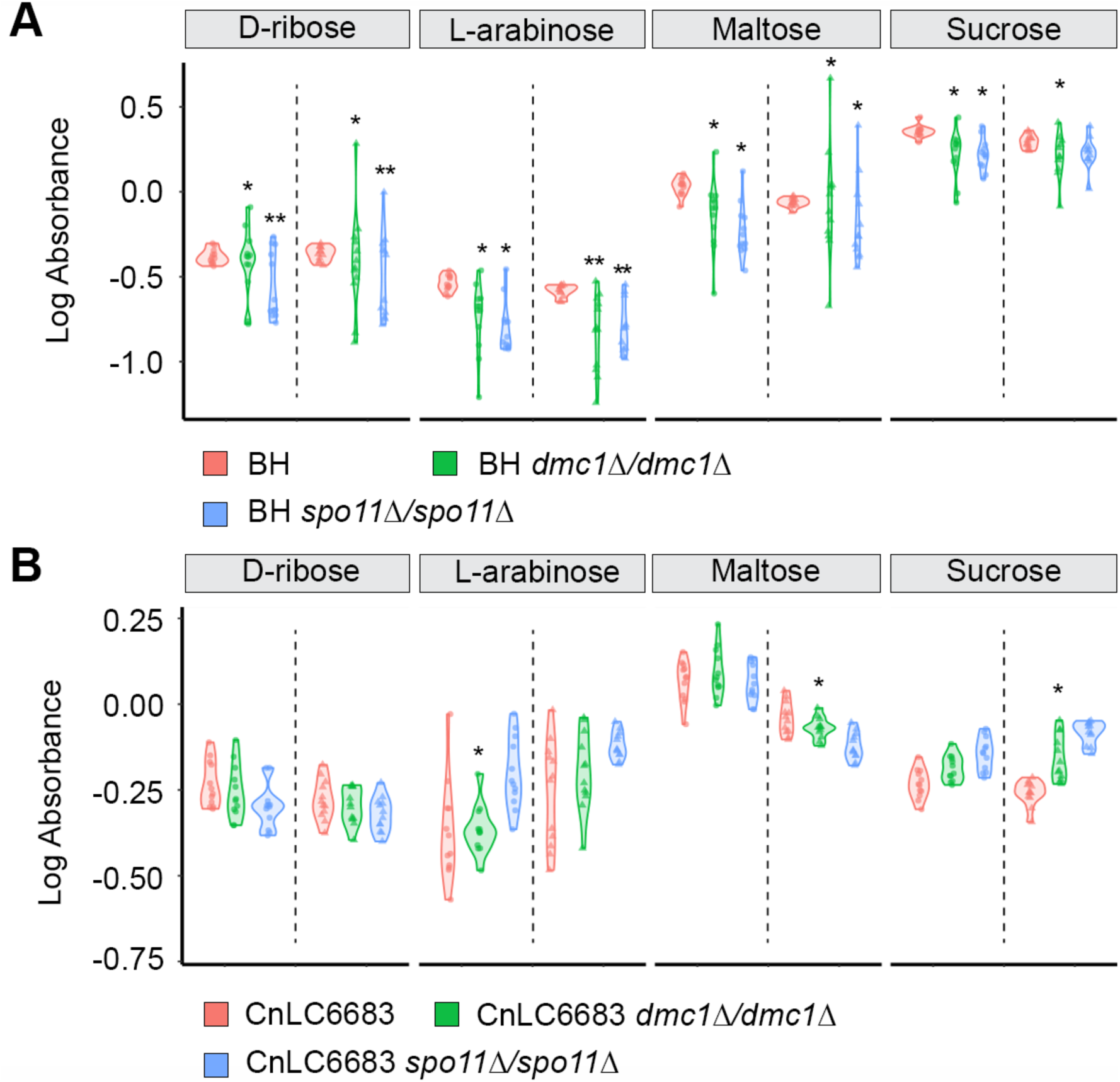
Titan colony cells derived from mutant BH strains have diverse carbon source utilization profiles. **(A)** 12 titan colony cells from each the BH, BH *dmc1*Δ/*dmc1*Δ and BH *spo11*Δ/*spo11*Δ mutant strains were used for carbon sources utilization assay. Cell growth in carbon sources including D-ribose, L-arabinose, maltose, and sucrose were calculated using optical density measured 6-dpi with normalization against their growth in glucose. The assay was repeated twice and data from both were plotted. The difference in variation among titan colony cells derived from different genotypes was assessed by Levene’s test (**, P <0.01; *, P <0.1). **(B)** The same assay was performed for titan colony cells derived from CnLC6683 wild type, CnLC6683 *dmc1*Δ/*dmc1*Δ, and CnLC6683 *spo11*Δ/*spo11*Δ strains. In contrast to the BH background, for majority tested group of cells, Levene’s test showed no significant difference in variation among titan colony cells derived from the three genotypes under CnLC6683 background.

Taken together, these results show that when heterozygosity is present in the genome, the absence of Dmc1 or Spo11 leads to titan cells producing daughter cells with significantly increased phenotypic variation.

### Deletion of the meiosis-specific genes leads to increased genome instability during depolyploidization of titan cells with genome-wide heterozygosity

Given the increased phenotypic variation in carbon-source utilization observed among cells produced by the BH *dmc1*Δ/*dmc1*Δ and BH *spo11*Δ/*spo11*Δ titan cells, we hypothesized that these phenotypic changes might be associated with underlying genomic changes that occurred during titan cell depolyploidization. To test this, we conducted whole-genome sequencing of colonies formed by titan cells derived from BH wild-type (29 colonies), BH *dmc1*Δ/*dmc1*Δ (12 colonies) and BH *spo11*Δ/*spo11*Δ (37 colonies) and mapped the reads to both the H99 and the Bt63 reference genomes. Our analyses revealed six different types of large scale genetic changes. Five of them (types 1-5) involve various types of loss of heterozygosity (LOH) through segmental or whole-chromosome deletion/duplication, as well as aneuploidy, and type 6 represents cases where non-diploid cells were produced from the titan cells (Figure 3A and Supplemental Figure S4). Notably, we found more cases with whole-chromosome alterations than those that only involved segmental chromosomal changes (Figure 3B). Full genome-wide plots for all sequenced titan colony cells are provided in Supplemental Figure S5 (BH wild-type), Supplemental Figure S6 (BH *dmc1*Δ/*dmc1*Δ), and Supplemental Figure S7 (BH *spo11*Δ/*spo11*Δ). A summary of mating type of all sequenced titan colony cells is presented in the Supplemental Table S2.

**Figure 3.**
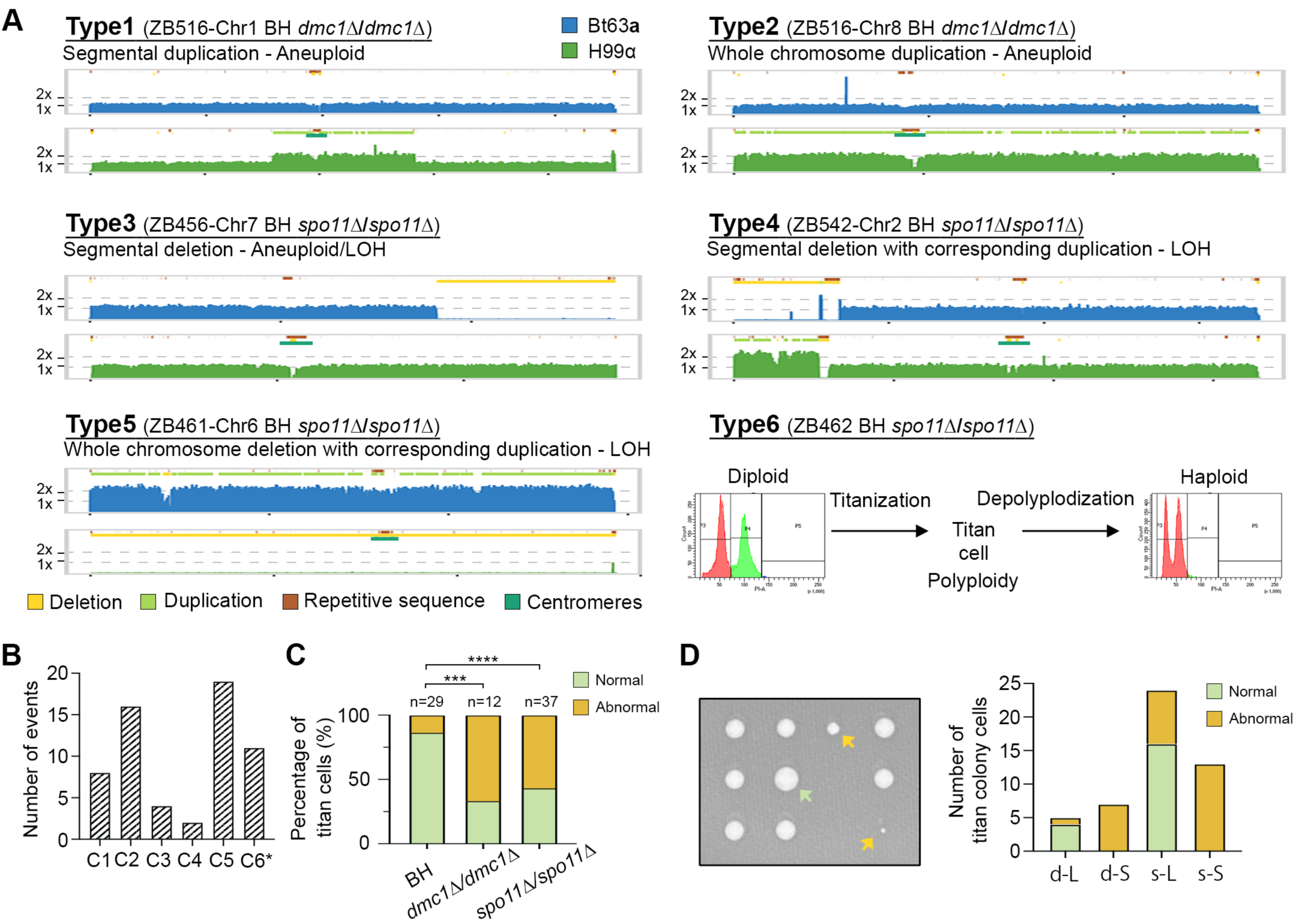
*DMC1* and *SPO11* are critical for chromosome stability during the ploidy reduction of titan cells with heterozygous genomes. (A) Representative LOH and aneuploid events identified in titan colony cells derived from BH *dmc1*Δ/*dmc1*Δ and BH *spo11*Δ/*spo11*Δ mutants, which were categorized into six types. For types 1 to 5, the information within the parentheses indicates the cell ID (i.e. ZB#), chromosome in which the event is identified (Chr#), and the genotype of the cell (BH *dmc1*Δ/*dmc1*Δ or BH *spo11*Δ/*spo11*Δ). (B) Number of events identified for each of the six types in titan colony cells derived from BH mutant strains. *: Number of cells with non-diploid DNA content based on flow cytometry analysis. (C) Percentages of viable titan cells produced daughter cell populations that have genomes that are normal (without any events belonging to the aforementioned six types) or abnormal (containing at least one event belonging to any of the six types). Statistical comparisons between different genetic backgrounds were based on Fisher’s Exact Test, and P values were adjusted for multiple comparisons using the Bonferroni method (****, *P* <0.0001; ***, *P* <0.001; ns, not significant). (D) Left: micro-dissected BH mutant titan cells can form both large and small colonies that are highlighted with green and yellow arrows, respectively. Right: similar to in (C), the bar chart shows the numbers of BH mutant cells with normal and abnormal genome contents. d-L: BH *dmc1*Δ/*dmc1*Δ large cells; d-S: BH *dmc1*Δ/*dmc1*Δ small cells; s-L: BH *spo11*Δ/*spo11*Δ large cells; s-S: BH *spo11*Δ/*spo11*Δ small cells.

When comparing among cells produced from different genetic backgrounds, a vast majority of those produced by the BH wild-type titan cells (78%) retained an unaltered genome, with only 22% containing genomic changes that belong to the six types characterized above (Figure 3C). In contrast, 57% of cells derived from the BH *spo11*Δ/*spo11*Δ titan cells and 67% of the cells derived from the BH *dmc1*Δ/*dmc1*Δ titan cells harbored various genomic alterations (Figure 3C), indicating that loss of *DMC1* or *SPO11* markedly increases the genomic instability and compromises the depolyploidization process in mutant titan cells. Interestingly, micro-dissected titan cells from both wild-type and mutant backgrounds formed colonies of varying sizes on YPD solid plates, and our analyses showed that every small-sized colony contained genomic alterations of various types (Figure 3D), suggesting that such chromosomal changes likely compromise cell growth, and that titan cells incapable of forming colonies may harbor even more extensive genomic disruptions that prevent daughter cell production.

Taken together, we indeed detected increased levels of genomic alteration in cells produced by the BH *dmc1*Δ/*dmc1*Δ and BH *spo11*Δ/*spo11*Δ mutant titan cells, which is consistent with the observed increase in their phenotypic variation.

### No canonical meiotic recombination or crossovers were observed in BH titan daughter cells

Our phenotypic and genotypic analyses demonstrated that the meiosis-specific genes *DMC1* and *SPO11* are critical for faithful depolyploidization process of heterozygous titan cells to produce viable daughter cells and form colonies. It is known that Spo11 and Dmc1 function prominently during meiosis by introducing and repairing double-strand breaks, respectively, contributing to the formation of crossovers and proper segregation of homologous chromosomes. We hypothesized that these two proteins might play similar roles during titan cell depolyploidization. To investigate this, we sought to analyze the cells produced from the BH wild-type titan cells to see if there were any signs of crossing over in their genomes.

Our hypothesis is that meiosis and crossovers occur when daughter cells are produced from titan cells; thus, the population of cells within the colony formed by a titan cell could be heterogenous in their genomic compositions, complicating the detection of meiotic crossovers. Instead, we first dissected titan cells formed by the BH wild-type strain onto YPD. Next, for each dissected titan cell, we separated and transferred every daughter cell that directly budded off the titan cell onto a different location (i.e. primary daughter cells), as well as cells that budded off these primary daughter cells (i.e. secondary daughter cells) within a 12-hour window (Figure 4A and Supplemental Figure S8). In total, we dissected 12 titan cells. Three of them did not produce any daughter cell, corresponding to a colony forming efficiency of 75%, which is comparable to our earlier observations of colony forming efficiency for the BH wild-type titan cell. Among the nine titan cells that produced daughter cells, we observed considerable variation in their ability to produce daughter cells: five titan cells produced at least four primary daughter cells, one titan cell produced three primary daughter cells, two titan cells produced two primary daughters, and one titan cell only produced a single primary daughter cell during the 12-hour window (Figure 4A and Supplemental Figure S8). In addition, primary daughter cells were observed to be produced simultaneously in four titan cells and were marked as Bd1 and Bd2 (Figure 4A and Supplemental Figure S8).

**Figure 4.**
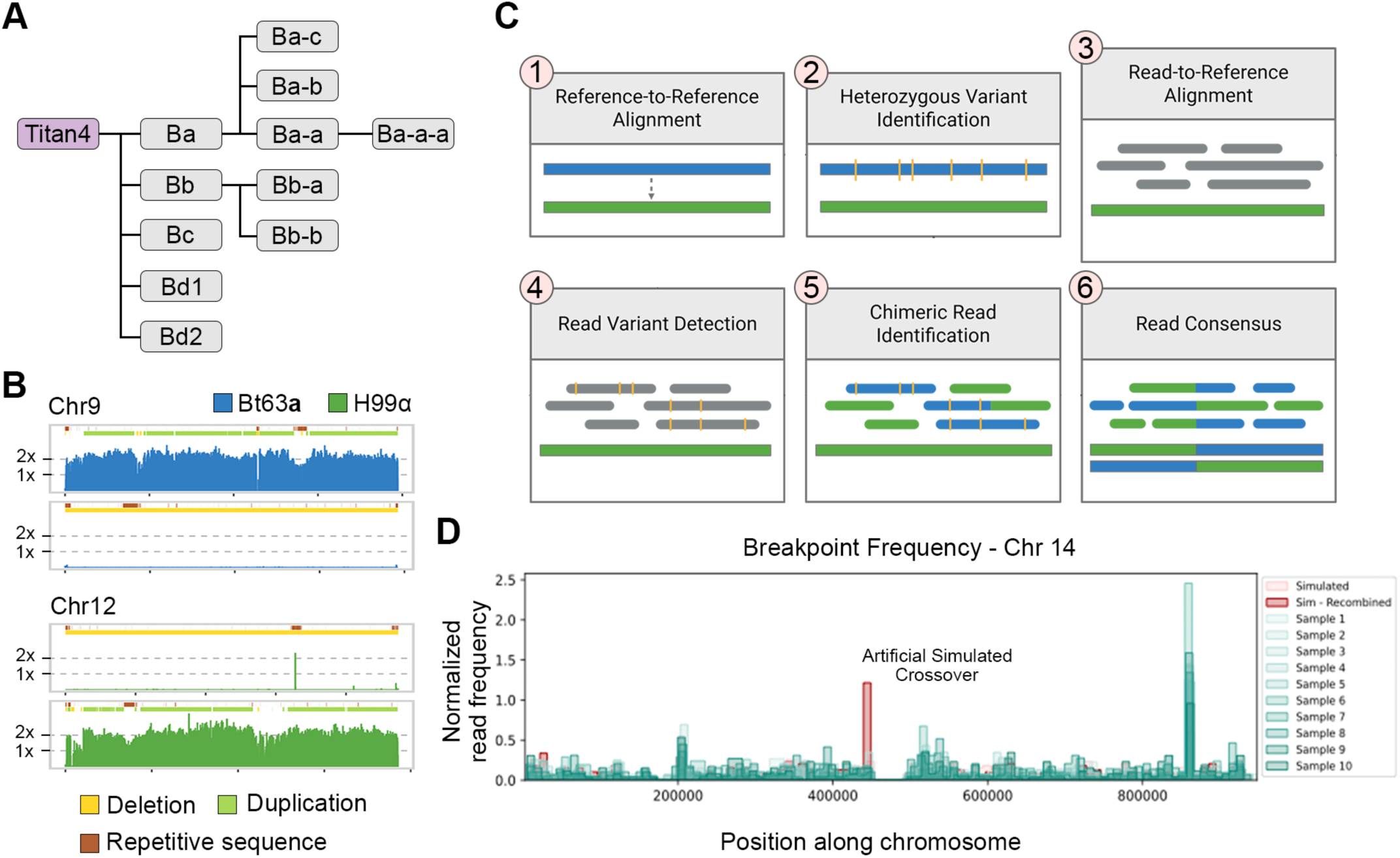
No canonical meiotic recombination was detected in BH titan daughter cells. **(A)** Titan daughter cells were used for recombination analysis and lineage dissection of BH titan cell 4 is shown as an example. **(B)** LOH events identified in all of the cells dissected in group 6 using whole-genome sequencing data. **(C)** Pipeline of recombination analysis in heterozygous diploid using nanopore sequencing results. **(D)** Output of the nanopore read analysis pipeline of 10 progeny samples in green compared to simulated samples with and without artificial crossover events in red. Counts of potentially chimeric reads containing breakpoints within 10kb windows were normalized to the genome-wide read depth for each sample.

We first subjected all of the dissected cells to Illumina whole genome sequencing. Except for the cells in group 6, all other cells exhibited a euploid 2N karyotype, carrying one H99-derived homolog and one Bt63-derived homolog for each chromosome. Loss of heterozygosity (LOH) on chromosome 9 (only the Bt63**a** allele) and on Chromosome 12 (only the H99α allele) was observed in all samples in group 6 (Figure 4B). Furthermore, the secondary daughter cell Ba-a harbors an additional copy of the H99α-derived chromosome 7, which likely arose during vegetative growth. Interestingly, all four samples in group 6 formed small colonies compared to the wild type BH (Supplemental Figure S8), suggesting that the identified LOH events may confer a growth disadvantage.

Because there is only ∼0.5% genetic divergence between the two parental genomes, Bt63 and H99, which constitute the diploid genome of the BH wild-type strain, the level of polymorphism may not be sufficient for detection of crossovers using the short reads generated from Illumina whole genome sequencing. To overcome this, we conducted Nanopore long-read sequencing for two primary daughter cells produced by titan cell No. 1, and one primary daughter cell from each of the other eight titan cells. We developed an analysis pipeline that is able to detect sequence breakpoints, which are junctions within the Nanopore reads where sequences on the two sides of the junction are of different origin (i.e. Bt63 and H99), indicative of crossovers (Figure 4C), and validated the pipeline with a simulated dataset that contains crossovers (Figure 4D and Supplemental Figure S9).

We next analyzed the Nanopore long-reads generated from the 10 primary daughter cells. To our surprise, we did not detect any statistically significant signal of crossover in any of the daughter cells. Specifically, the Nanopore reads all showed profiles that are distinct from the simulated dataset with crossovers, and resembled the simulated dataset without recombination. The plot generated for Chromosome 14 is shown as an example (Figure 4D). Further inspection of the reads underlying the minor breakpoint candidates confirmed that they are not crossovers and all of them are located within genomic regions with highly repetitive sequences or transposable elements (e.g. centromeric and telomeric regions). Thus, while Dmc1 and Spo11 are important for faithful depolyploidization of titan cells with genome-wide heterozygosity, our analyses of daughter cells produced by the titan cells showed that they have no canonical meiotic crossovers in their genomes, indicating that the two meiosis-specific genes might be exerting their influence on titan cell depolyploidization through novel mechanisms yet to be characterized.

## DISCUSSION

In this study, our data revealed a severe, heterozygosity-dependent defect in ploidy reduction of titan cells when the meiosis-specific genes *DMC1* or *SPO11* were deleted. The percentage of titan cells that produced viable daughter cells decreased significantly, and the surviving titan cell colonies frequently carried LOH or aneuploidy that was often linked to smaller colony size. The reason why these segregation errors are uniquely deleterious in a heterozygous genome could be that LOH converts heterozygous loci with divergent Bt63/H99 alleles to homozygosity, therefore unmasking recessive defects and/or disrupting favorable allele combinations of co-evolved genes, and aneuploidy distorts allelic dosage compared to the original euploid genome. In contrast, while Dmc1 and Spo11 may still participate in ploidy reduction of titan cells produced by homozygous diploid strains, the phenotypic consequences of their absence could be attenuated because: 1) without heterozygosity, genomes with the reciprocal crossovers and whole chromosome LOH observed in the BH backgrounds are both effectively dosage-balanced diploids resembling that of the wild-type progenitor strain; and 2) in cases of aneuploidy, gene expression imbalance would be less disruptive when both homologs are identical. Indeed, Dmc1 has been shown previously to be activated during infection of the host, when titan cell formation is induced (36). It should be noted that the H99 genome exhibits one reciprocal chromosomal translocation compared to Bt63, which can further sensitize the system in strains of the BH background as LOH or aneuploidy affecting these chromosomes can cause severe local copy number or structure imbalances, amplifying fitness costs and explaining the strong dependency on *DMC1* and *SPO11* to stabilize chromosome transmission during titan cell depolyploidization.

Because *DMC1* and *SPO11* are involved in the ploidy reduction process when titan cells produce daughter cells, and the canonical function of these two meiosis-specific genes is to generate (Spo11) and repair (Dmc1) DNA DSBs, which will result in crossover events, we hypothesized that cross-over events might exist in the titan daughter cells derived from the heterozygous diploid BH strain. Interestingly, we found no evidence of canonical crossovers between Bt63 and H99 homologs with apparent breakpoint enrichments all localized to telomeric, centromeric, or repetitive regions. Importantly, however, the absence of a Bt63/H99 crossover signal does not exclude recombination during the ploidy reduction process. Polyploid titan cells can harbor multiple copies of the same parental homolog, and recombination that occurs between genetically identical templates, for example, two Bt63 copies or two H99 copies, would be undetectable with SNP-based analysis, because recombination would have occurred between identical sequences. Likewise, non-crossover repair such as Synthesis-dependent strand-annealing (SDSA) (42, 43) or very short gene-conversion tracts (44) could fall below our detection thresholds, particularly in regions of low marker density. Thus, our data argue against frequent or widespread genetic exchange between divergent homologs during titan cell depolyploidization, but they remain compatible with recombination on identical templates, a route that could still serve to resolve DNA entanglements or promote chromosome organization and segregation without generating detectable haplotype reshuffling.

Deletion of *DMC1* or *SPO11* in the heterozygous BH background produced a striking reduction in viable progeny from dissected titan cells, and whole-genome sequencing of the survivors revealed widespread LOH and aneuploidy, often at a whole-chromosome scale and linked with smaller colony size. This result suggests that Dmc1 and Spo11 may act to stabilize chromosome segregation during depolyploidization, and that their loss converts the reduction division into an error-prone process that yields uniparental disomy (LOH) and copy number imbalance. Mechanistically, segregation defects can arise from multiple, non-exclusive failures: erroneous kinetochore–microtubule attachments (45, 46), sister-chromatid cohesion defects (47, 48), abnormal centrosome or centriole numbers (49–51), or an impaired spindle assembly checkpoint (SAC) (52–54). Any of these will increase non-disjunction and thereby contribute to the LOH or aneuploidy we observed. The stronger defect in BH *dmc1*Δ/*dmc1*Δ suggests that Dmc1-mediated homology tethering may be especially important to align homologous chromosomes and suppress merotelic attachment. Distinguishing among these possibilities will require further investigation involving live imaging of centromere-tagged chromosomes during titan depolyploidization, quantitative assays of kinetochore occupancy and error correction, cohesion localization, spindle-pole body counting, or dynamics in function of key components of SAC (Mad1, Mad2, or Bub1) in wild type versus *dmc1* and *spo11* deletion mutants.

Based on our observations in this study, when titan cells were induced from a haploid parent (H99), all titan-derived colonies had a haploid DNA content, whereas titan cells induced from diploid parents (CnLC6683 or BH) produced colonies that predominantly have a diploid DNA content. This concordance between the parental ploidy and the major ploidy of progeny implies that the ploidy reduction process proceeds under tight control to restore a balanced genome. Here, we envision a multi-stage process: host-like and environmental cues (nutrient limitation, low pH, CO , serum components, low cell density (13, 14, 16, 18, 19)) trigger the highly regulated process of titanization and endoreplication (15, 18, 20), yielding a uninucleate but highly polyploid cell with altered wall and capsule architecture and stress tolerance (7, 11, 21, 55). During the subsequent reduction phase, titan cells may execute a replication-licensed step analogous to a pre-reduction S-phase, reorganize their spindles, and engage meiotic toolkit components (including Dmc1 and Spo11) to promote homolog engagement and faithful segregation without invoking frequent canonical crossovers. Once the segregation “state” is established, daughter cells emerge by budding. This is consistent with our observation that when we dissected daughter cells budding from titan cells, while in most cases only one daughter cell was produced at a time, there were also occasions where two daughter cells were produced simultaneously at different locations of the titan cell. In addition, the budding process usually occurs after a noticeable lag before the first daughter cell is produced, with subsequent budding events occurring with shorter intervals. We hypothesize that the DNA content for daughter cells might be prepared and packed together inside the titan cell before the budding process starts.

Thus, the nuclei of titan cells could also serve as storage sites for genetic information under stressful and challenging conditions, allowing daughter cells to be quickly produced to repopulate the niche when favorable conditions return, which requires meiosis-specific genes (e.g. *DMC1* and *SPO11*) to ensure faithful return to the original parental ploidy, which likely represents the best adapted fitness state when the stresses are absent.

## Supplemental figure legends

**Supplemental Figure S1. Flow cytometry of titan cell cultures.**

(A) Bright-field image of the titan cell culture analyzed by flow cytometry. Cultures were sonicated immediately prior to analysis. More than 100 titan cells were examined, and no cell clusters were observed, indicating that events measured by flow cytometry represent single cells. Events were also doublet-excluded during analysis. Scale bar, 10 μm. (B) Scatter plots of PI fluorescence (DNA content; x-axis) versus cell size (Area; y-axis) for four samples: haploid control, diploid control, titan cell culture before filtration (pre-filtration), and titan cell culture after passage through a 10 μm cell strainer to enrich for large cells (post-filtration). For each sample, gated populations 2–5 are shown separately (left four columns) and overlaid in the “All events” panel to illustrate the relationship between cell size and ploidy. The rightmost column shows PI intensity histograms for each sample, with the contributions of populations 2–5 indicated in matching colors, revealing the ploidy distribution of haploid and diploid controls and of titan cell cultures before and after filtration.

**Supplemental Figure S2. Pilot assay of carbon source utilization of titan colony cells using Biolog YT microplates.**

Carbon source utilization by progenitors and titan colony cell progeny were assessed using Biolog YT Microplates. Principle component analysis was performed on the normalized absorbance values to determine which carbon sources contributed greatest to variance between tested lines. Loadings (defined as the product of eigenvector and the square root of the eigenvalue, which gives the amount of variance accounted for by each carbon source) for the first two principal components were squared, summed, and square rooted, giving the Euclidean distance to the origin. Carbon sources which accounted for a large amount of the variation between lines are expected to have high values for both progenitors and progeny. D-ribose, L-arabinose, maltose, and sucrose were selected for the following assays in expanded sets of titan colony cells given their relatively high loadings.

**Supplemental Figure S3. Levene’s test of difference in variation among titan colony cells.** Variance in the growth after 6 days for titan colony cells of *dmc1*Δ/*dmc1*Δ, *spo11*Δ/*spo11*Δ mutants, and wild-type BH (A) and CnLC6683 (B) were compared with Levene’s test for equal variances. The mean value of each titan colony cell’s growth in a carbon substrate was normalized by dividing by the mean growth in D-glucose for that titan colony cell. The variance of these normalized means was then compared, with the difference in variance and significance of the test statistic plotted for each genotype and carbon source, from two experimental repeats.

**Supplemental Figure S4. Flow cytometry analysis of titan colony cells.**

Propidium iodide (PI) stained DNA content histograms are shown for control strains and titan colony cells. The top row shows haploid 1C/2C and diploid 2C/4C control strains that served to define the gates. The middle and bottom rows show representative titan colony cells derived from BH *dmc1*Δ/*dmc1*Δ and BH *spo11*Δ/*spo11*Δ strains (sample IDs indicated above each plot). Several titan isolates display predominantly haploid-like DNA content, whereas others exhibit mixed or elevated (triploid/higher) DNA content relative to the haploid and diploid controls.

**Supplemental Figure S5. Illumina Whole genome sequencing analysis of BH and titan colony cells derived from BH strain.**

For each isolate, sequences were aligned to a combined H99 and Bt63 reference genome and copy number variants were called from the resulting read depth. Normalized sequencing depth is shown across all 14 chromosomes of Bt63 (left panel) and H99 (right panel). Putative copy-number variants are annotated, with duplications (green bar on top) and deletions (yellow bar on top), respectively. Simple repeats (blue) and other repetitive elements (brown) are also marked on top. Centromeres (between two dark green dots) and the *MAT* locus (light brown bar on Chromosome 5) are highlighted in the corresponding H99 genome.

**Supplemental Figure S6. Whole genome sequencing analysis of BH *dmc1*Δ/*dmc1*Δ and titan colony cells derived from BH *dmc1*Δ/*dmc1*Δ strain.**

For each isolate, sequences were aligned to a combined H99 and Bt63 reference genome and copy number variants were called from the resulting read depth. Normalized sequencing depth is shown across all 14 chromosomes of Bt63 (left panel) and H99 (right panel). Putative copy-number variants are annotated, with duplications (green bar on top) and deletions (yellow bar on top), respectively. Simple repeats (blue) and other repetitive elements (brown) are also marked on top. Centromeres (between two dark green dots) and the *MAT* locus (light brown bar on Chromosome 5) are highlighted in the corresponding H99 genome.

**Supplemental Figure S7. Whole genome sequencing analysis of BH *spo11*Δ/*spo11*Δ and titan colony cells derived from BH *spo11*Δ/*spo11*Δ strain.**

For each isolate, sequences were aligned to a combined H99 and Bt63 reference genome and copy number variants were called from the resulting read depth. Normalized sequencing depth is shown across all 14 chromosomes of Bt63 (left panel) and H99 (right panel). Putative copy-number variants are annotated, with duplications (green bar on top) and deletions (yellow bar on top), respectively. Simple repeats (blue) and other repetitive elements (brown) are also marked on top. Centromeres (between two dark green dots) and the *MAT* locus (light brown bar on Chromosome 5) are highlighted in the corresponding H99 genome.

**Supplemental Figure S8. Lineage dissection of BH titan cells.**

**(A)** Lineage dissection maps of nine individual BH titan cells (Titan1-Titan9) that produced daughter cells. For each lineage, the titan colony cell is shown in purple, and each gray box represents a progeny colony. Colonies in the same vertical column belong to the same generation, and connecting lines indicate parent-daughter relationships. Colony names (Ba, Ba-a, Ba-a-a, Bb, etc.) denote their position within each lineage. **(B)** Colony morphology of the Titan6 lineage compared with wild-type BH on YPD agar. The BH control forms a large colony, whereas Titan6 and its progeny Ba, Ba-a and Bb all form smaller colonies.

**Supplemental Figure S9. Developed program can identify recombination and partial chromosome LOH event in heterozygous diploid background.**

Reads plots showing breakpoints representing recombination event identified in simulated heterozygous diploid sample using developed program.

## MATERIALS AND METHODS

### Strains, media and growth conditions

*C. neoformans* strains used in this study are listed in Supplemental Table S1. Fresh cultures were revived and maintained on YPD (1% yeast extract, 2% Bacto Peptone, 2% dextrose) agar media, supplemented with appropriate antifungal drugs (nourseothricin (NAT) at 100 μg/mL or neomycin (G418) at 200 μg/mL) when needed, and incubated at 30°C. Titan cell induction was performed using the published methods from Ballou’s lab (7). Briefly, cells were incubated in Yeast Nitrogen Base (YNB) liquid media with 2% glucose at 30°C overnight, followed by adjusting OD to 0.001 and incubated in PBS with 10% Fetal Bovine Serum (FCS) (Thermo Fisher, USA) at 37°C with 5% CO2. Cell cultures were then washed with PBS for 3 times, centrifuged, and resuspended in PBS. Titan cell and titan daughter cell dissection was carried out using a Nikon yeast spore dissection microscope equipped with a glass fiber needle 25 μm in diameter.

### Construction of mutant strains

Deletion mutant strains were generated in the *C. neoformans* H99, Bt63, or CnLC6683 backgrounds. To generate the deletion alleles, the *NAT* or *NEO* gene expression cassette was amplified from plasmids pAI3 and pJAF1, respectively. Approximately 1.5-kb regions (homologous arms) flanking the genes of interest were amplified from wild-type strain genomic DNA and fused with the *NAT* or *NEO* drug resistance marker with overlapping PCR, as previously described (56), to generate the donor DNA cassettes. CRISPR-Cas9-directed mutagenesis was used for mutant generation. The CAS9 cassette was PCR-amplified from plasmid pXL-1 with universal primers M13F and M13R (56). The desired target sequences for the sgRNA constructs were designed using the Eukaryotic Pathogen CRISPR guide RNA/DNA Design Tool (EuPaGDT) with default parameters (57). Two gRNAs were designed and used for each gene of interest. Complete gRNAs were generated by one-step overlap PCR. About 1.5 µg donor DNA cassette, 400 ng CAS9 cassette, and 150 ng of each complete gRNAs fragment were mixed and condensed to a 5 µL volume before being introduced to strains with the transient CRISPR/Cas9 coupled with electroporation (TRACE) transformation approach (56). The obtained transformants were validated by genotyping using diagnostic PCRs for the internal region of the ORF, 5’-junction and 3’-junction regions, as well as spanning PCR of the region encompassing the deletion construct as previously described (58).

### Generation of BH wild type, BH *dmc1*Δ/*dmc1*Δ, and BH *spo11*Δ/*spo11*Δ mutant strains

To generate the heterozygous diploid strain BH, the H99α mutant *ura5*::*NEO* and wild-type Bt63**a** cells were mixed in equal proportions in ddH_2_O and inoculated and incubated on V8 medium for 48 hours at room temperature in the dark to allow cell fusion. The cells were then harvested and plated on YPD + NEO solid medium plates to isolate single colonies, with 150-200 colonies per plate. Colonies from YPD + NEO plate were replica-plated onto SD-uracil agar medium (0.67% nitrogen base w/o amino acid, 0.002% uracil drop out mix, 2% agar, 2% dextrose) and incubated at 30°C. Colonies that could grow on both YPD + NEO and SD-uracil agar plates were recovered on YPD + NEO and maintained as yeast colonies. Colony PCR was performed to confirm the presence of both the **a** and the α mating types. Flow cytometry and whole genome sequencing were used to confirm the ploidy of the resulting diploid strain BH. BH *dmc1*Δ/*dmc1*Δ strain was generated by fusing Bt63 *dmc1*Δ and H99 *dmc1*Δ. BH *spo11*Δ/*spo11*Δ strain was generated by fusing Bt63 *spo11*Δ and H99 *spo11*Δ.

### Generation of *MAT*-homozygous BH and CnLC6683 strains

To generate the *MAT*-homozygous (**a**/**a** and α/α) strains in the BH and CnLC6683 backgrounds, cells were spread onto V8 medium and incubated at room temperature in the dark to induce self-filamentation. Non-selfing cells grow as yeast colonies, indicative of LOH at the *MAT* locus . Cells from these non-selfing colonies were then streaked onto YPD medium plates for colony purification and further verified for their mating type locus by PCR and ploidy by flow cytometry.

### Flow cytometry analysis

Samples for flow cytometry analysis to determine *Cryptococcus* ploidy were prepared as previously described (58, 63). Briefly, samples were fixed in 70% ethanol overnight at 4°C. Fixed samples were then washed once with 1 mL NS buffer (10 mL Tris-HCl (1 M, pH=7.5), 85.6 g sucrose, 2 mL EDTA (0.5 M, pH=8), 0.095 g MgCl_2_, 0.0147 g CaCl_2_, 0.0136 g ZnCl_2_, 0.096 g phenylmethylsulfonyl fluoride, 0.49 mL 2-mercaptoethanol (added freshly), 989 mL H_2_O) and resuspend in 200 µL Master mix (183.5 µL NS buffer, 6 µL PI (1µg/µL), 10.5 µL RNase (20 mg/mL)) and incubated at room temperature overnight. 50 µL stained samples were then added into 500 µL Tris-PI mix (482 µL Tris-HCl (1 M, pH=7.5), 18 µL PI (1µg/µL)) for flow cytometry analysis. The flow cytometry analysis was performed at the Duke Cancer Institute Flow Cytometry Shared Resource Laboratory with a BD FACSCanto II Flow Cytometer (40, 59). Ploidy of titan-derived colonies was assessed by PI flow cytometry with internal standards on the same run: H99 (1C/2C) and CnLC6683 (2C/4C). Events were doublet-excluded (area/width gating). Progeny were called haploid or diploid when their 1C/2C (or 2C/4C) peaks co-aligned with the corresponding control peaks.

### Carbon source utilization assay and statistical analysis

To assess metabolic traits which varied among titan colony cells, Biolog YT Microplates (Biolog, Inc., Hayward, CA, USA) were inoculated in triplicate, using BUY agar, YT inoculating solution, and the suggested optical density, then incubated at 28°C in a humid enclosure. Data were collected at 3 days post inoculation, when absorbance was read at 500 nm for oxidation tests and 590 nm for assimilation tests. Values in each plate were blanked based on the absorbance of the negative control and divided by the growth in D-glucose for normalization. Principle component analysis (PCA) was performed on the normalized values, then the loadings were determined to identify the amount of variance explained by each carbon source in the overall matrix. Carbon sources that explained high amounts of the variance for both were used for testing a smaller number of carbon sources with additional titan colony cells.

Variability of titan colony cell growth was then evaluated via assimilation of selected carbon sources. D-ribose, L-arabinose, maltose, and sucrose were selected as carbon sources, with D-glucose as a control treatment for normalization. All 12 viable titan-colony isolates available for *dmc1*Δ/*dmc1*Δ were used for the phenotype analysis. To match sample size across genotypes, we then randomly selected 12 colonies from BH, BH *spo11*Δ/*spo11*Δ, CnLC6683, CnLC6683 *dmc1*Δ/*dmc1*Δ, and CnLC6683 *spo11*Δ/*spo11*Δ. Selected strains were grown in triplicate in 96 well microtiter plates (Falcon, USA) in 160 μl YNB supplemented with 666 mM per C atom of each carbon source (i.e. 111 mM for D-glucose, 133 mM for D-ribose) for up to 6 days at 28°C, with daily measurements of absorbance at 600 nm. For comparison of variance among progeny, absorbance values were log-transformed, then averaged by strain and normalized to the D-glucose condition. Mean normalized absorbances of each genotype were then used to compare variances of mutant titan colony cells to the variance of wild-type titan colony cells with Levene’s test.

### Whole genome sequencing and nanopore sequencing

Freshly dissected titan colony cells or titan daughter cells were first allowed to grow from micro-dissected titan mothers on YPD at 30 °C for 48 h (72 h for small colony isolates). Each colony was then picked, resuspended in 1 mL of 15% glycerol, and stored at −80 °C. For genomic DNA extraction, isolates were revived by spotting 10 µL of the glycerol suspension onto fresh YPD plates and incubating at 30 °C for 24 h, and cells were then harvested for DNA prep using the previously described protocol (40). No liquid culturing was involved so as to reduce any possible inter-cell competition.

Illumina sequencing was conducted at the Duke Sequencing and Genomic Technologies core facility (https://genome.duke.edu), on a Novaseq X platform with 250 bp paired-end option. High molecular weight genomic DNA for Nanopore long read sequencing was isolated using a modified CTAB method, as previously described (23). Nanopore long-read libraries were prepared and sequenced in house using a MinION device with R10.4.1 flow cells according to the manufacturer’s instructions. All of the sequencing data are available under BioProject ID PRJNA1328773.

### Read depth analysis of short-read sequencing data

Illumina short read sequences for all samples were analyzed using WeavePop (v1.0.0) (60). Briefly, Sequences were aligned to a combined H99 and Bt63 reference genome using Snippy (https://github.com/tseemann/snippy) and copy number variants were called from the resulting read depth. The full parameters and configuration files for the WeavePop pipeline can be found in the following GitHub repository: https://github.com/magwenelab/BH_titan_cell_ploidy_parameters/

### Recombination detection using Nanopore long reads

Long Nanopore reads from progeny samples were aligned to reference genomes using the lra aligner (v1.3.7) (61), with separate alignments performed against the H99 major parental reference, the Bt63 minor parental reference, and a concatenated reference containing both genomes. Alignment files were processed using samtools (v1.18) (62) for sorting, indexing, and extracting unmapped reads. Bedtools (v2.31.0) (63) was used to convert BAM files to BED format for downstream analysis. Reference genomes were preprocessed to ensure consistent chromosome naming and ordering. Whole-genome alignments between parental references were generated with wfmash (v0.10.) (64) and structural variant detection was performed with SyRI (v1.6.3) (65) to identify SNPs, insertions, and deletions distinguishing the parental haplotypes.

Following alignment, known SNPs and indels distinguishing the H99 and Bt63 parental genomes were identified from syri output and filtered to retain only SNPs. The resulting file was analyzed using a pipeline implemented in python (v3.7.12). Using pybedtools (v0.9.0) (66) and pandas (v1.3.5) (67), intersections between aligned reads and SNP positions were computed for each chromosome. For each read overlapping a SNP, the base was compared to both parental references and assigned to H99, Bt63, or marked as unknown. This parent-calling step was parallelized by chromosome using Python’s multiprocess (v0.70.14) (68) and pysam (v0.21.0) libraries.

Consecutive SNPs with identical parent assignments were merged into stretches, and short stretches less than 1000bp and containing fewer than 4 SNPs were merged into the subsequent stretch to reduce the impact of sequencing errors. Reads containing substantial stretches that were assigned to both parents were flagged as chimeric. Breakpoint positions where parent assignment switched were extracted and their frequency across the genome was calculated. Data visualization and further analysis were performed using matplotlib (v3.5.3) (69), seaborn (v0.12.2) (70), and numpy (v1.21.6) (71), with candidate chimeric reads and breakpoint loci summarized in output tables and plots.

### Wild-type progeny comparison with simulated long reads

Simulated long-read datasets were generated with PBSIM3 (v3.0.0) (72), which sampled progeny Nanopore reads to match the error profile and length distribution observed in the data. Recombinant and non-recombinant genomes were constructed by swapping defined regions of random chromosomes between parental references, and concatenated FASTA files were used as templates for simulation. PBSIM3 was run in whole-genome sampling mode with a target depth of 20×, producing simulated FASTQ files for both recombinant and unrecombined genomes. Scripts used for the simulation of long-read datasets have been uploaded to the GitHub repository (https://github.com/magwenelab/BH_titan_cell_ploidy_parameters/).

To account for differences in sequencing depth across samples, the number of chimeric reads at each breakpoint position was normalized by dividing by the average genome-wide read depth for each sample. Average depth was calculated from coverage statistics generated by samtools, and normalization was performed prior to plotting. Breakpoint densities for simulated samples and 10 wild type experimental samples were visualized together with seaborn and matplotlib. For each chromosome, breakpoint positions were binned and plotted as normalized histograms, with each sample assigned a distinct color and legend label. This allowed direct comparison of recombination signatures between simulated and experimental datasets, facilitating identification of regions with elevated chimeric read frequency. Regions with a normalized breakpoint frequency greater than 0.5 were selected and the pileup of reads underlying the loci were inspected for signatures of canonical recombination. A crossover signal was defined as a locus where more than 60% of the reads supported a switching between parental haplotypes at the breakpoint location, and two types of reciprocal chimeric reads were present in approximately equal frequencies. A loss of heterozygosity signal was defined as a locus where more than 20% of the reads supported a switching between parental haplotypes at the breakpoint location, and only one type of chimeric read was present.

## Acknowledgments

We thank Dr. Steven Haase, Dr. Bin Li, and Huarui Zhou for their invaluable efforts and insightful discussions that improved our flow cytometry data. We also thank Dr. Steven Haase, Dr. Thomas Petes, Dr. Xiaorong Lin, and Dr. Kristen Nielsen for their thoughtful comments and suggestions on this manuscript. Their feedback and insights greatly improved the clarity and interpretation of our work. This work was supported by NIH/NIAID awards R01AI050113-20, R01AI039115-28, R01AI133654-08 to JH. JH is a fellow and co-director with Leah Cowen of the CIFAR program Fungal Kingdom: Threats & Opportunities.

