## Supplementary figures and images for "Meiosis-specific genes play roles in ploidy reduction in *Cryptococcus neoformans* titan cells"

### Supplemental Figure 1

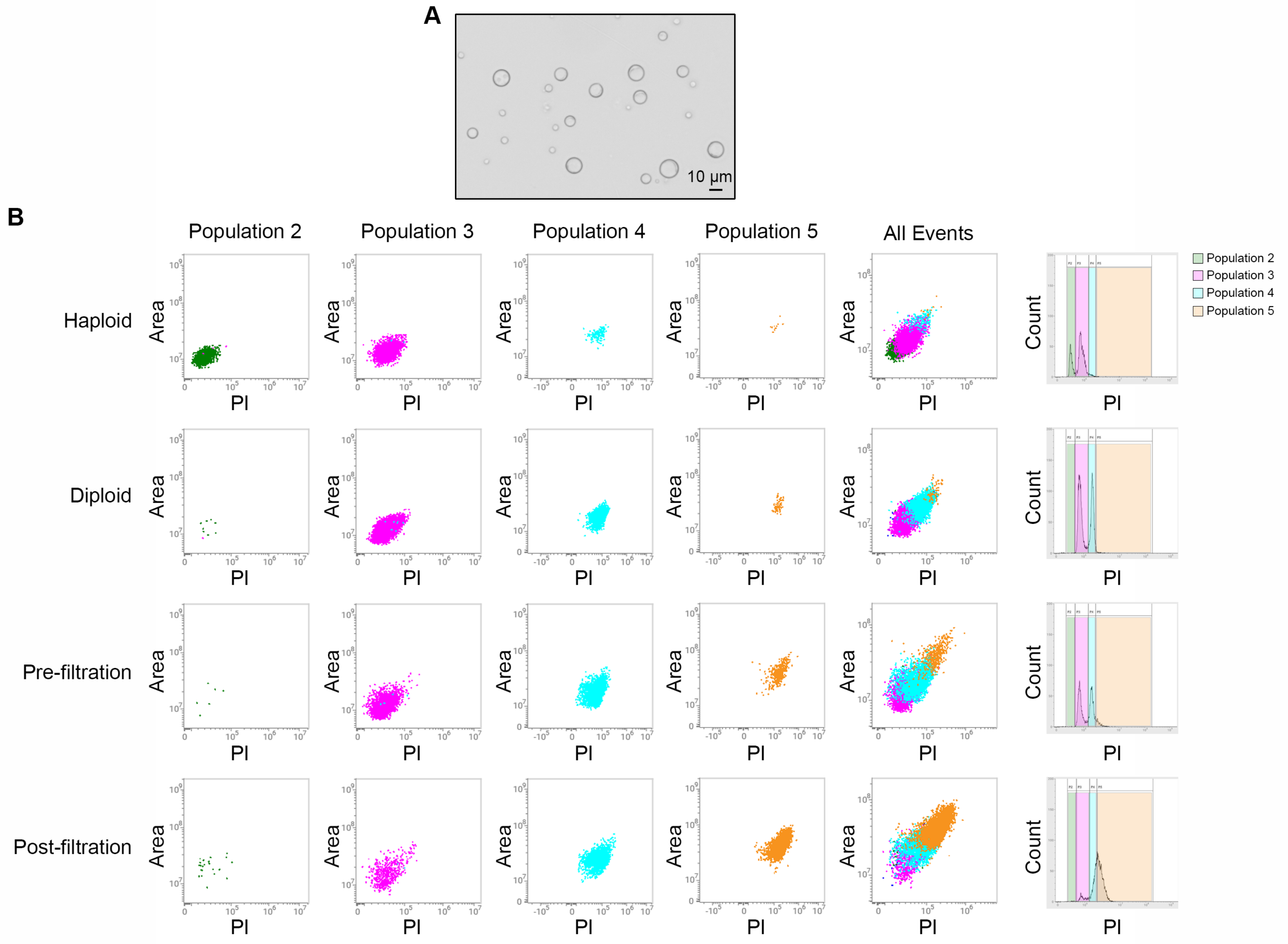

### Supplemental Figure 2

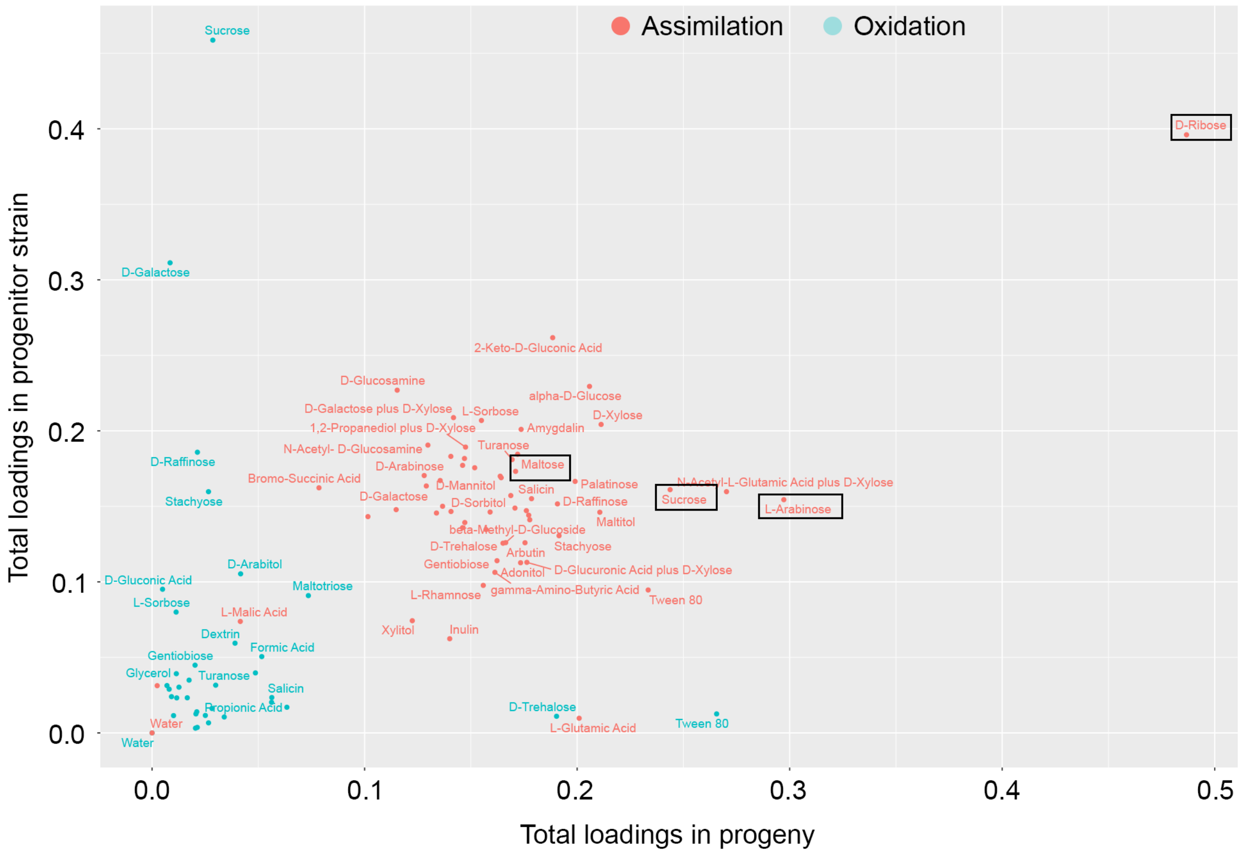

### Supplemental Figure 3

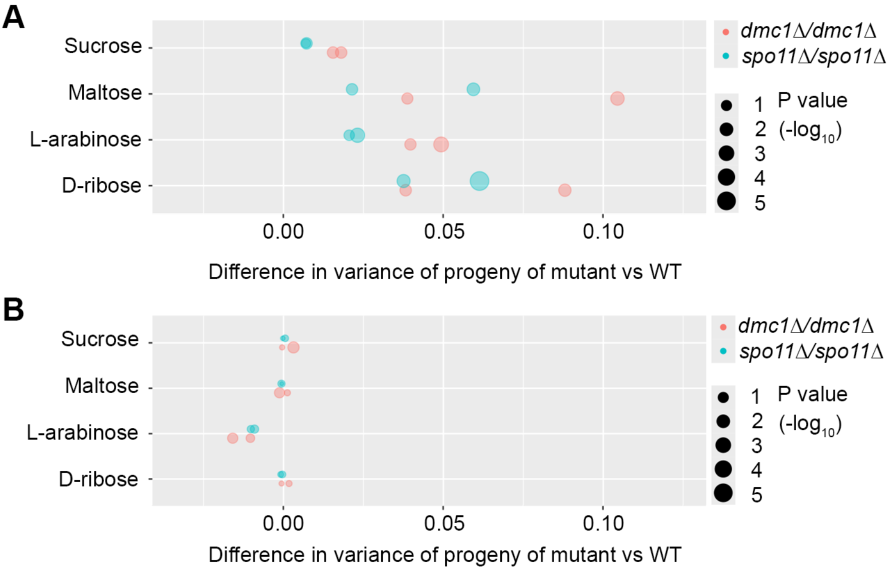

### Supplemental Figure 4

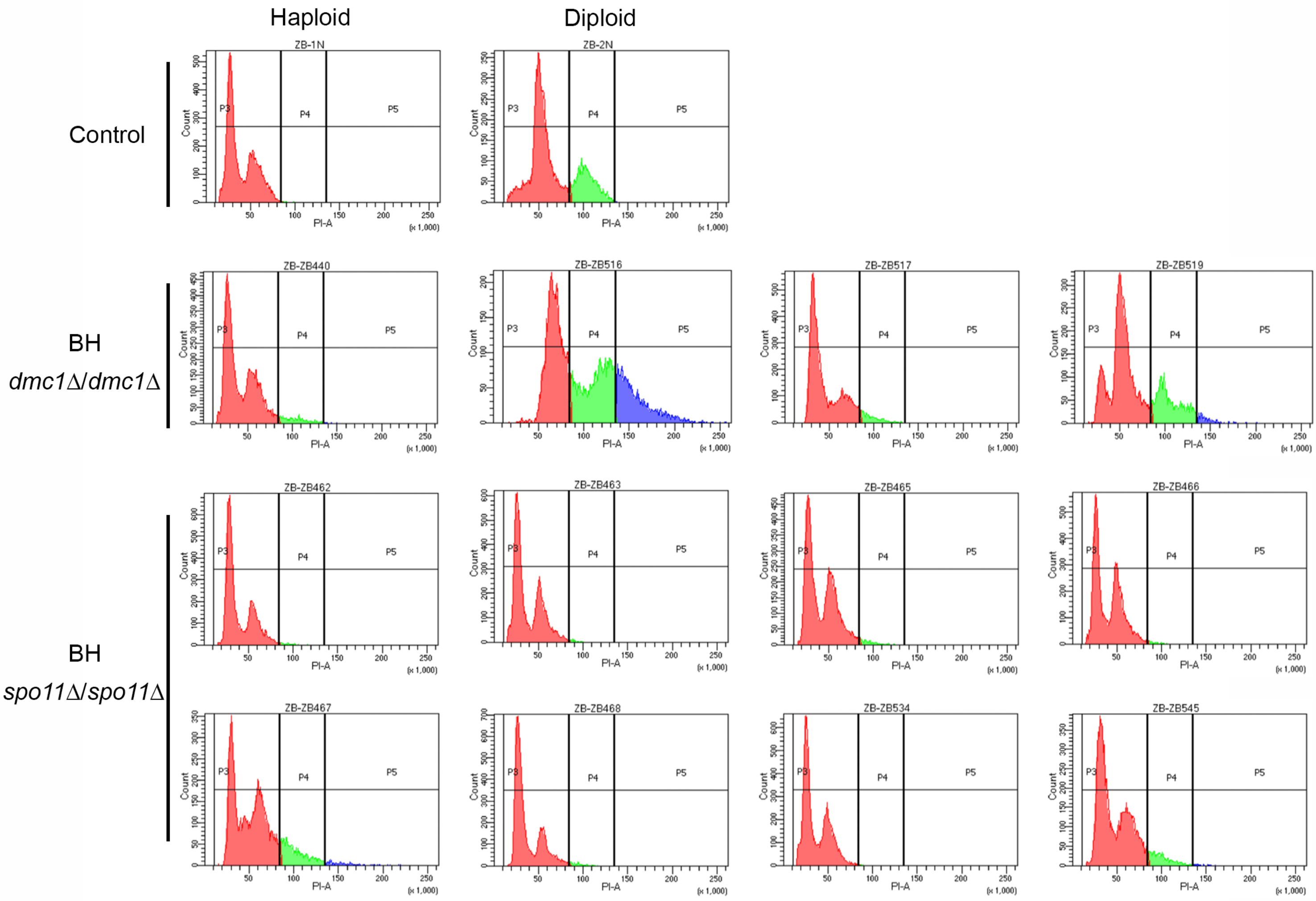

### Supplemental Figure 8

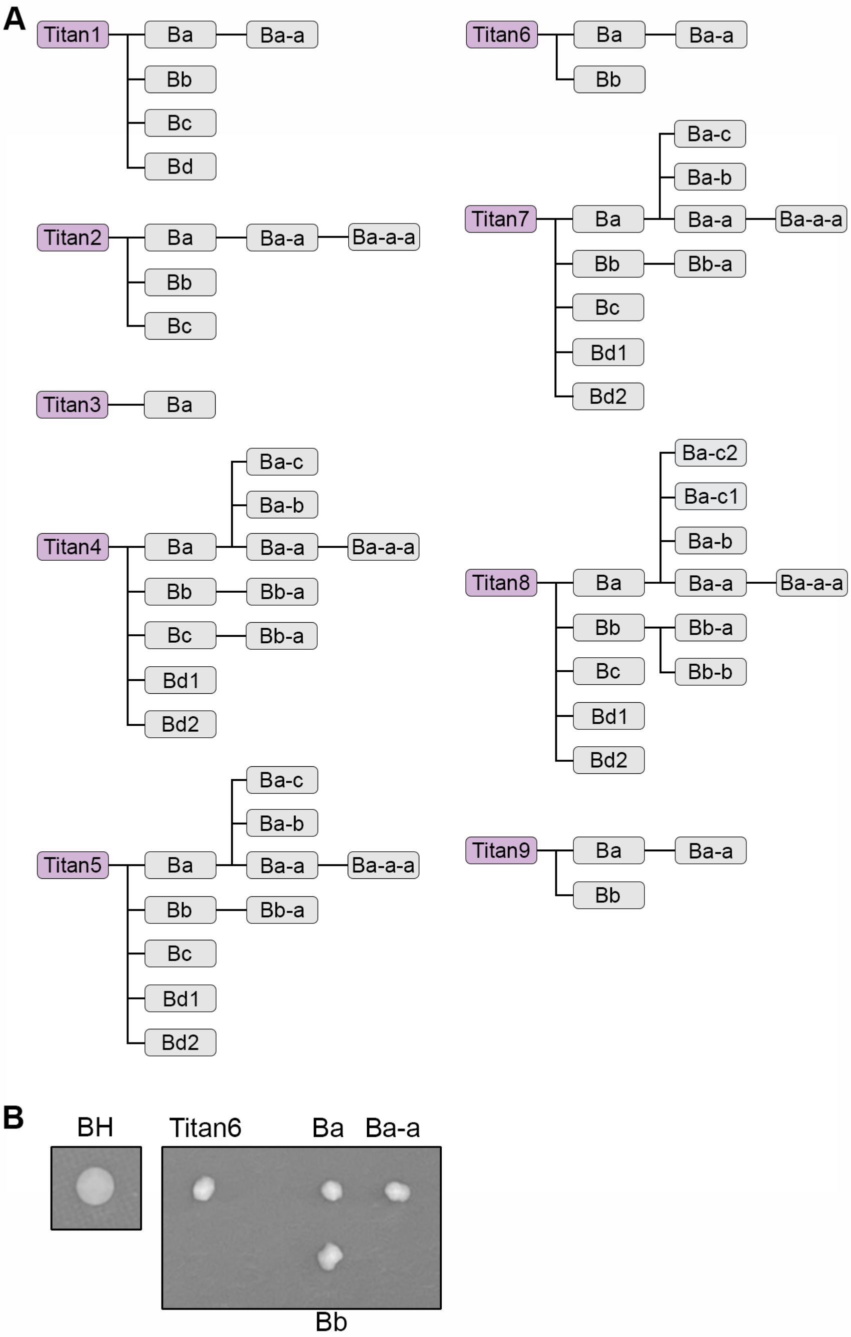

### Supplemental Figure 9

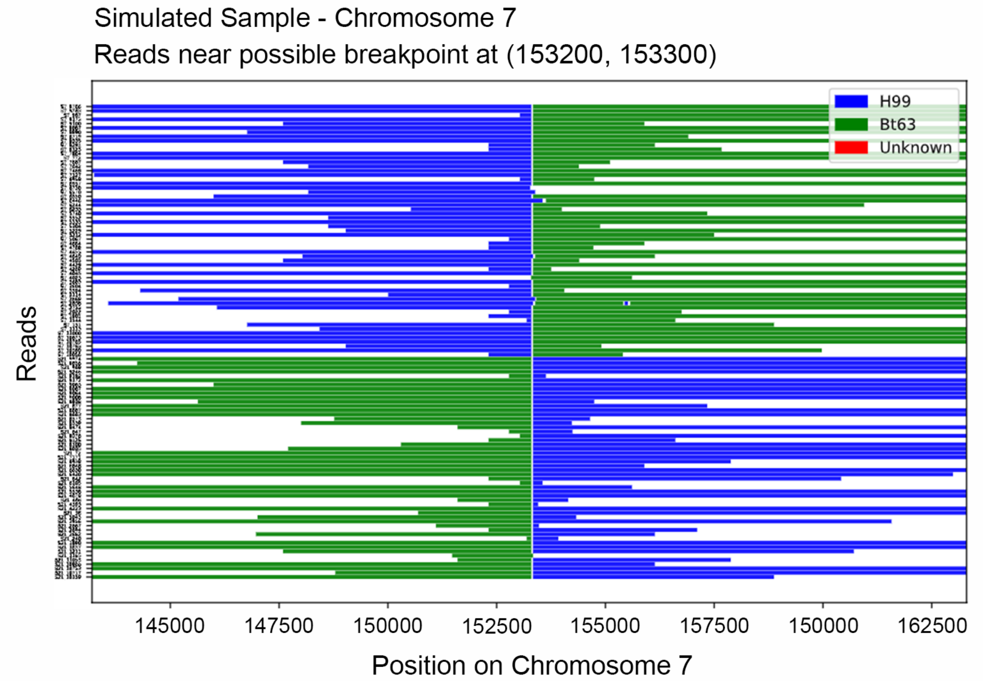
