## Supplemental Figure 5 for "Meiosis-specific genes play roles in ploidy reduction in *Cryptococcus neoformans* titan cells"

### Normalized Depth of Windows Along Chromosomes

Sample: ZB_38_S38_L001 Strain: wildtype_control Lineage: H99_Bt63 Window size: 500

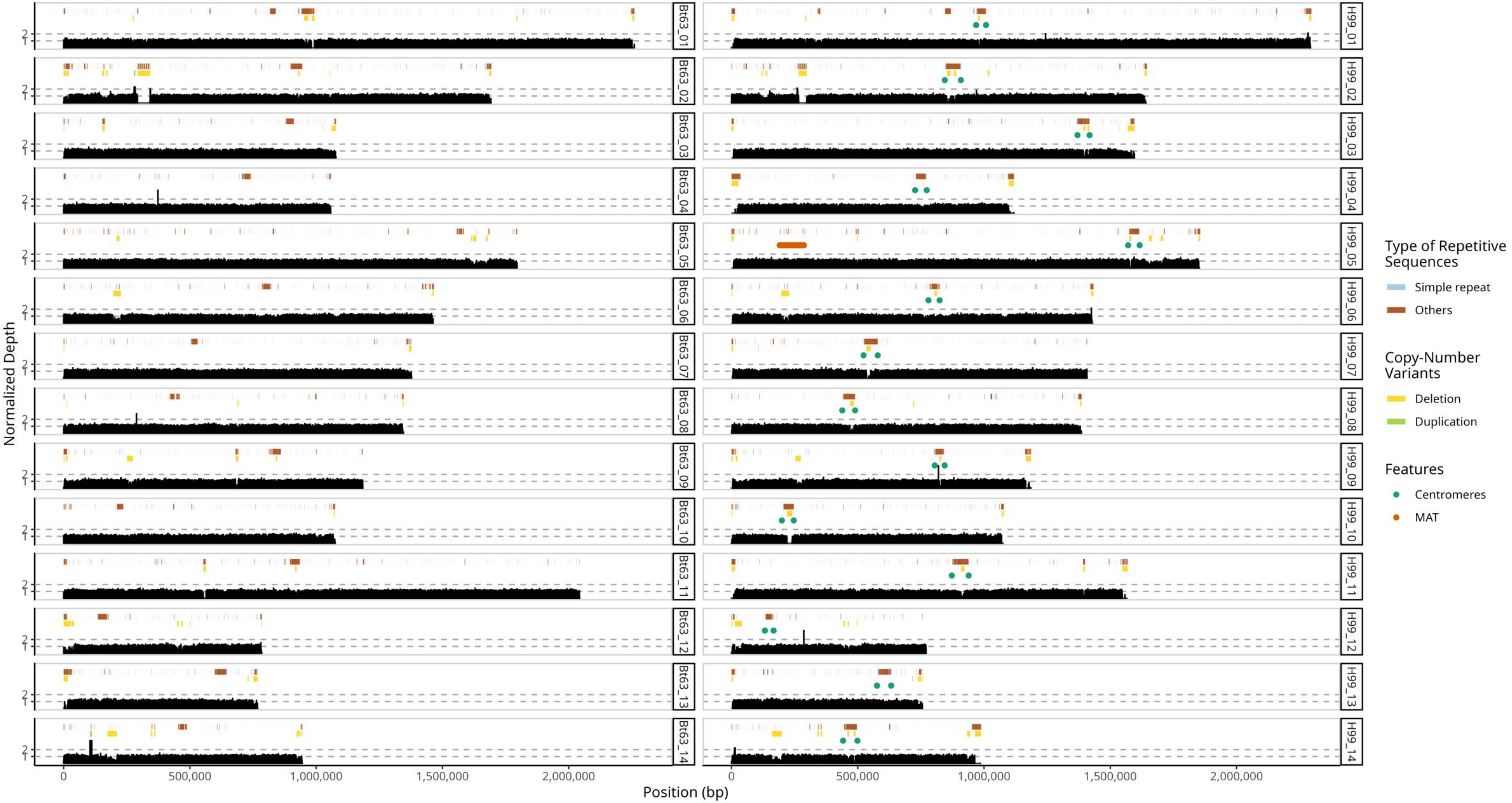

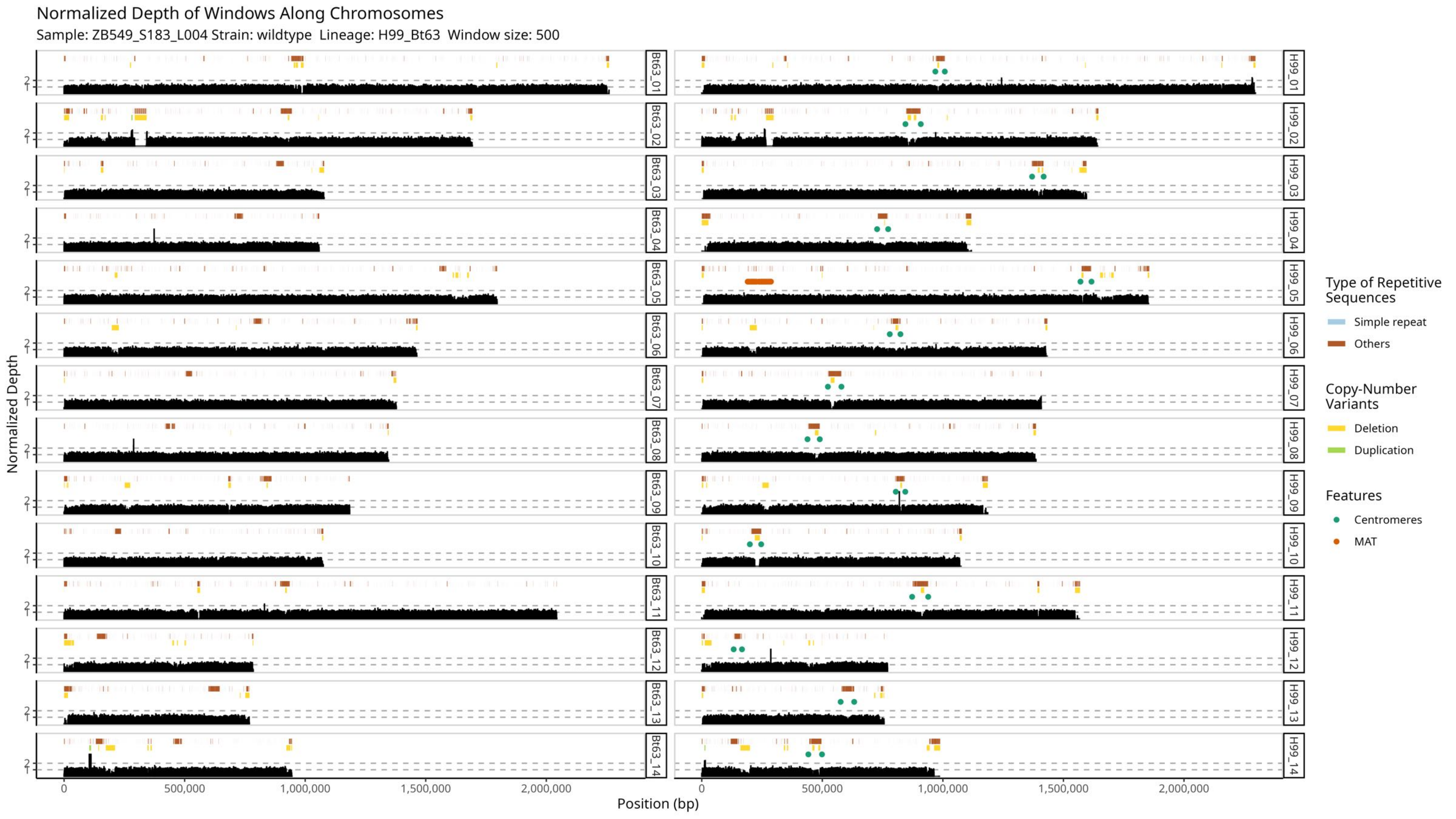
Normalized Depth of Windows Along Chromosomes
Sample: ZB549_S183_L004 Strain: wildtype Lineage: H99_Bt63 Window size: 500
Normalized Depth
Position (bp)
Bt63_01
Bt63_02
Bt63_03
Bt63_04
Bt63_05
Bt63_06
Bt63_07
Bt63_08
Bt63_09
Bt63_10
Bt63_11
Bt63_12
Bt63_13
Bt63_14
H99_01
H99_02
H99_03
H99_04
H99_05
H99_06
H99_07
H99_08
H99_09
H99_10
H99_11
H99_12
H99_13
H99_14
0
500,000
1,000,000
1,500,000
2,000,000
1
2
Type of Repetitive Sequences
Simple repeat
Others
Copy-Number Variants
Deletion
Duplication
Features
Centromeres
MAT

Normalized Depth of Windows Along Chromosomes

Sample: ZB550_S184_L004 Strain: wildtype Lineage: H99_Bt63 Window size: 500

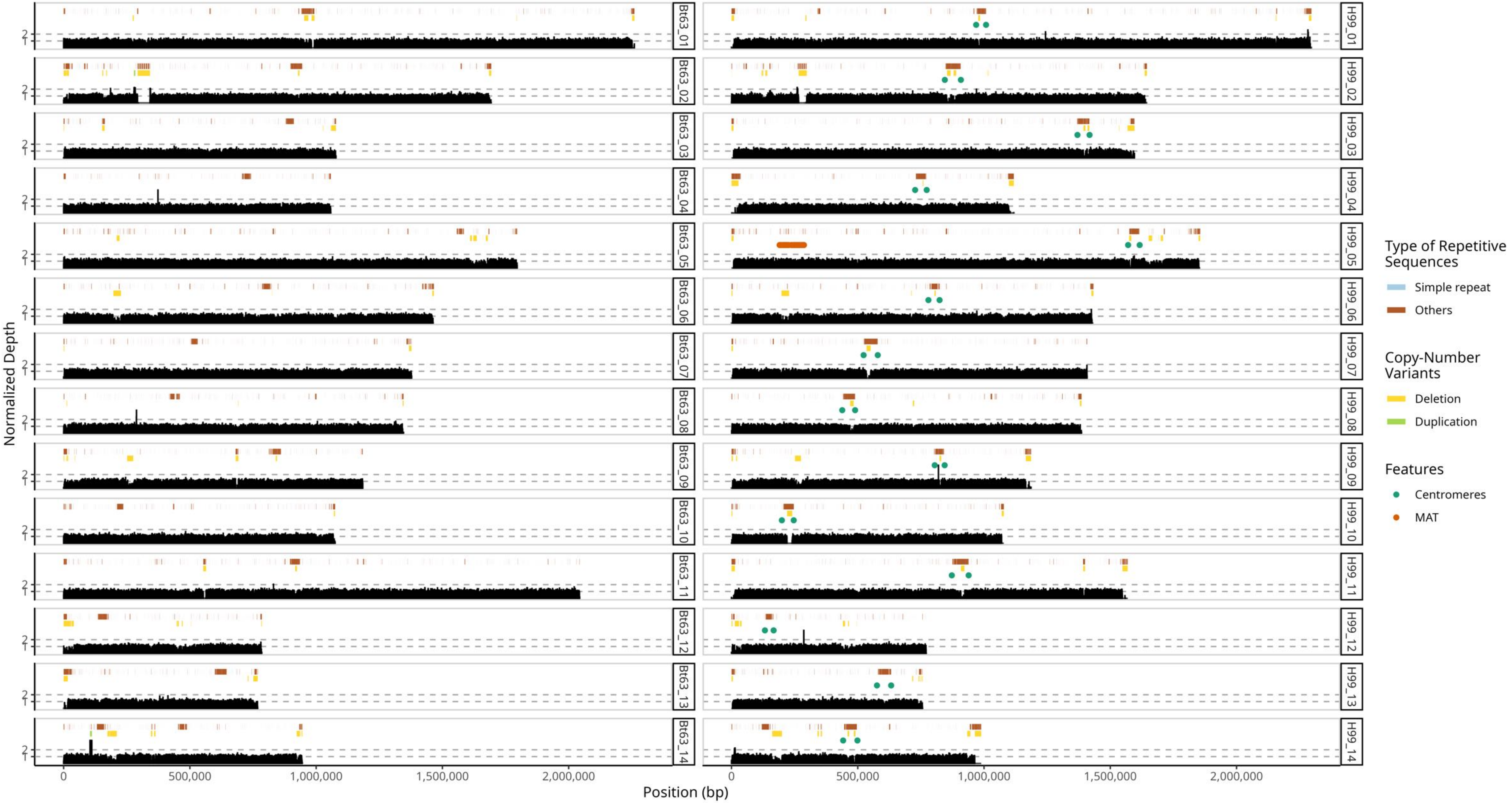

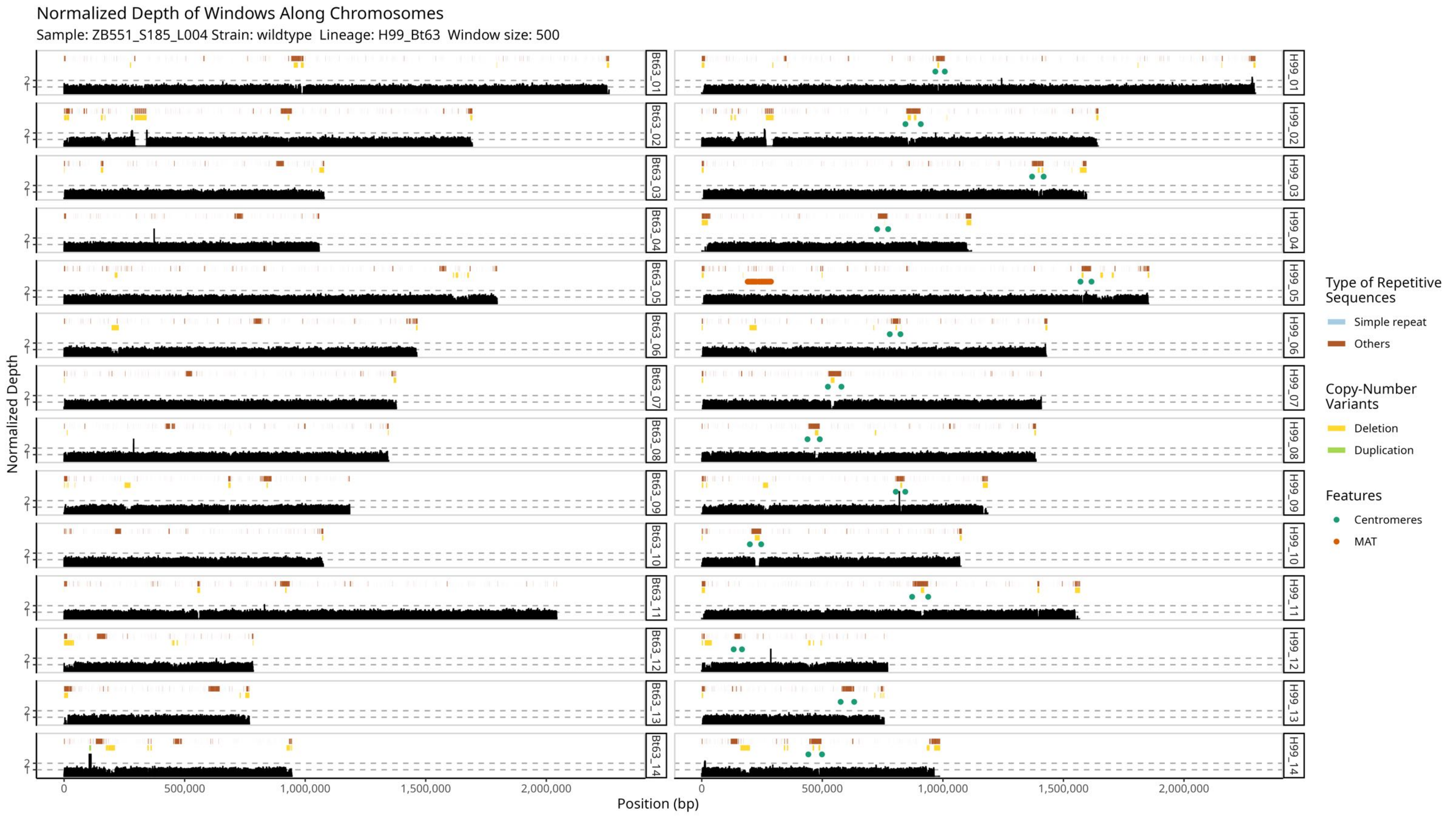

Normalized Depth of Windows Along Chromosomes
Sample: ZB551_S185_L004 Strain: wildtype Lineage: H99_Bt63 Window size: 500
Normalized Depth
Position (bp)
Bt63_01
Bt63_02
Bt63_03
Bt63_04
Bt63_05
Bt63_06
Bt63_07
Bt63_08
Bt63_09
Bt63_10
Bt63_11
Bt63_12
Bt63_13
Bt63_14
H99_01
H99_02
H99_03
H99_04
H99_05
H99_06
H99_07
H99_08
H99_09
H99_10
H99_11
H99_12
H99_13
H99_14
0
500,000
1,000,000
1,500,000
2,000,000
Type of Repetitive Sequences
Simple repeat
Others
Copy-Number Variants
Deletion
Duplication
Features
Centromeres
MAT

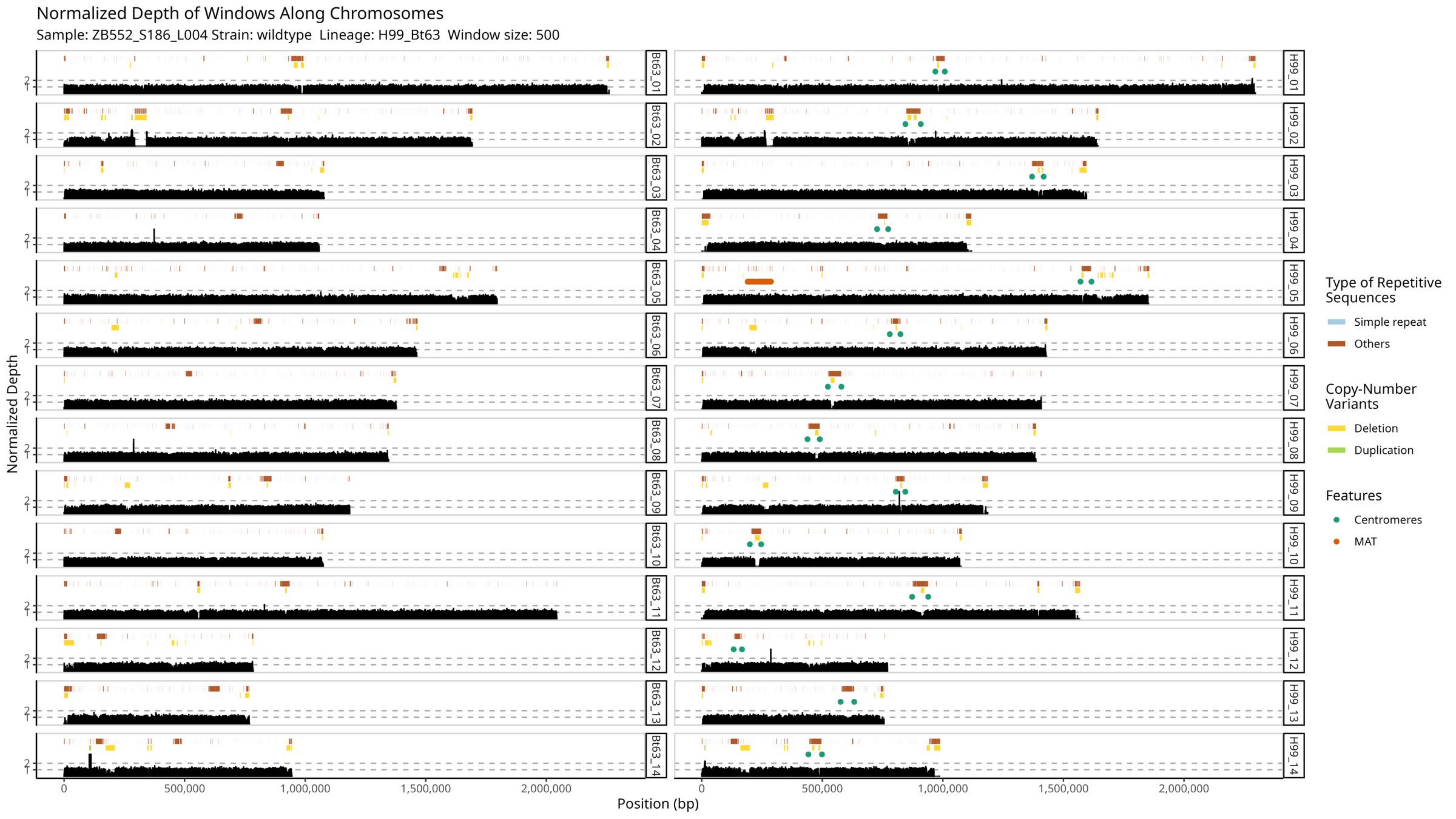

Normalized Depth of Windows Along Chromosomes
Sample: ZB552_S186_L004 Strain: wildtype Lineage: H99_Bt63 Window size: 500
Normalized Depth
Position (bp)
0
500,000
1,000,000
1,500,000
2,000,000
1
2
Bt63_01
Bt63_02
Bt63_03
Bt63_04
Bt63_05
Bt63_06
Bt63_07
Bt63_08
Bt63_09
Bt63_10
Bt63_11
Bt63_12
Bt63_13
Bt63_14
H99_01
H99_02
H99_03
H99_04
H99_05
H99_06
H99_07
H99_08
H99_09
H99_10
H99_11
H99_12
H99_13
H99_14
Type of Repetitive Sequences
Simple repeat
Others
Copy-Number Variants
Deletion
Duplication
Features
Centromeres
MAT

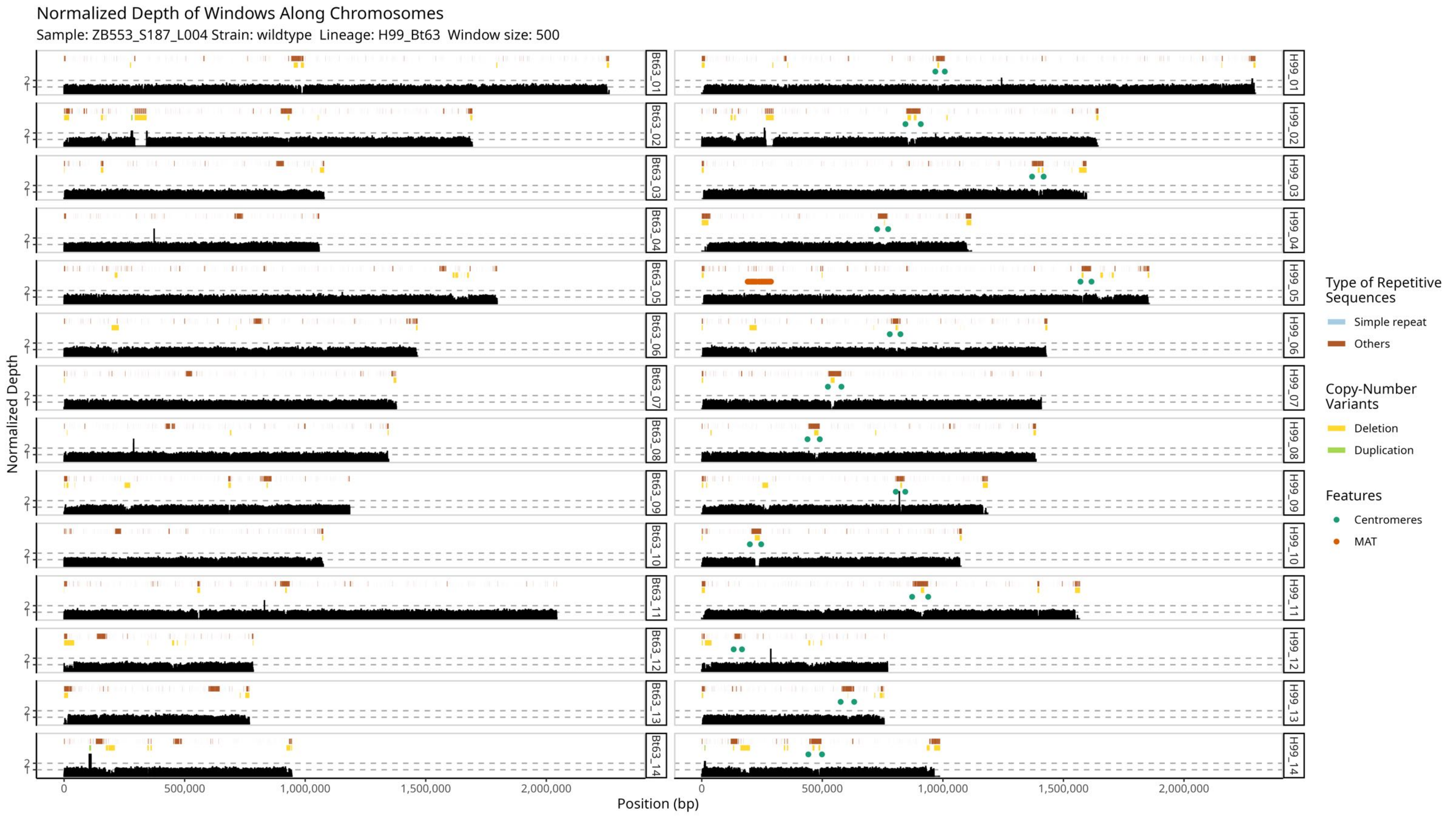
Normalized Depth of Windows Along Chromosomes
Sample: ZB553_S187_L004 Strain: wildtype Lineage: H99_Bt63 Window size: 500
Normalized Depth
Position (bp)
0
500,000
1,000,000
1,500,000
2,000,000
1
2
Bt63_01
Bt63_02
Bt63_03
Bt63_04
Bt63_05
Bt63_06
Bt63_07
Bt63_08
Bt63_09
Bt63_10
Bt63_11
Bt63_12
Bt63_13
Bt63_14
H99_01
H99_02
H99_03
H99_04
H99_05
H99_06
H99_07
H99_08
H99_09
H99_10
H99_11
H99_12
H99_13
H99_14
Type of Repetitive Sequences
Simple repeat
Others
Copy-Number Variants
Deletion
Duplication
Features
Centromeres
MAT

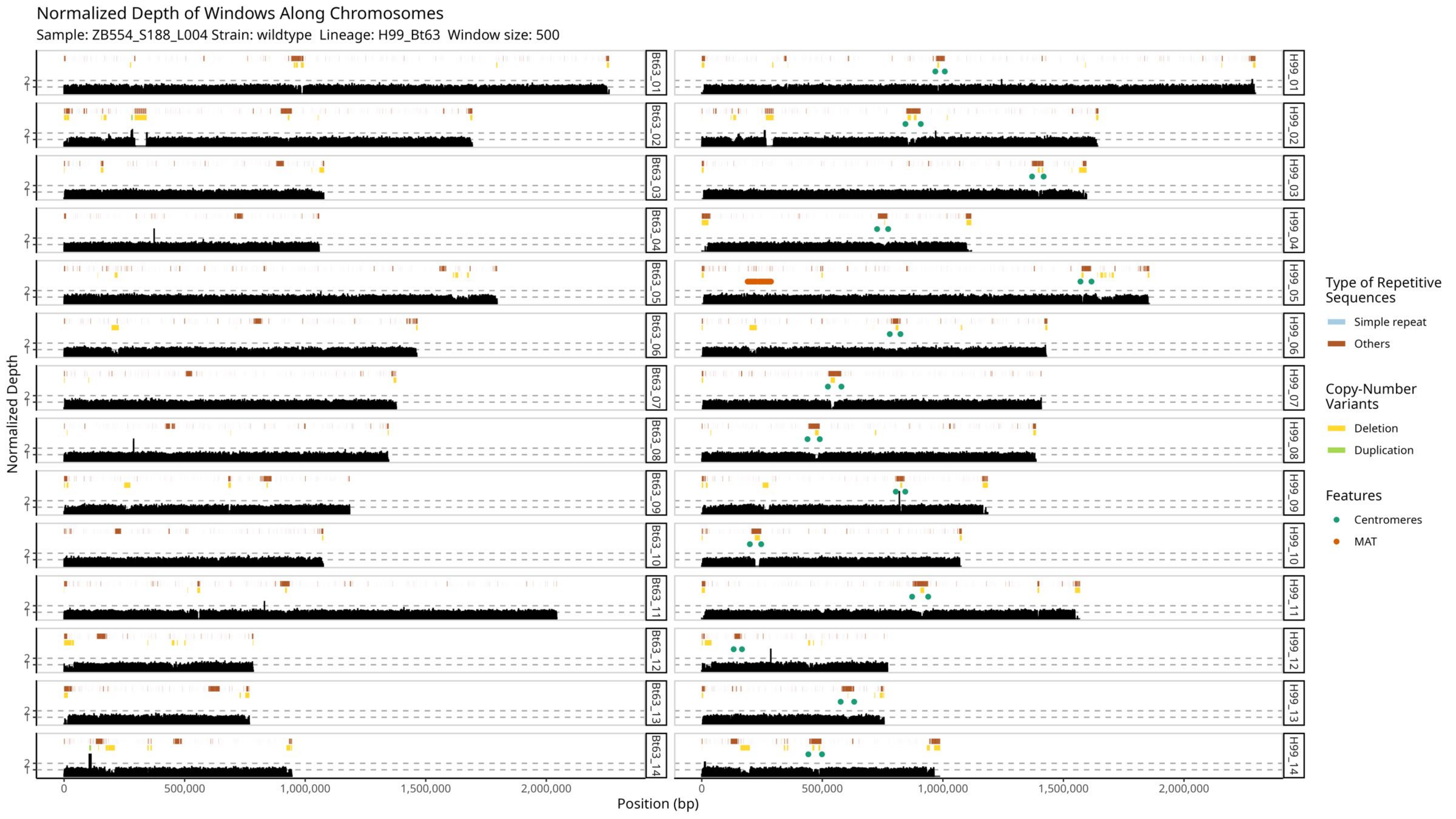

Normalized Depth of Windows Along Chromosomes
Sample: ZB554_S188_L004 Strain: wildtype Lineage: H99_Bt63 Window size: 500
Normalized Depth
Position (bp)
Bt63_01
Bt63_02
Bt63_03
Bt63_04
Bt63_05
Bt63_06
Bt63_07
Bt63_08
Bt63_09
Bt63_10
Bt63_11
Bt63_12
Bt63_13
Bt63_14
H99_01
H99_02
H99_03
H99_04
H99_05
H99_06
H99_07
H99_08
H99_09
H99_10
H99_11
H99_12
H99_13
H99_14
0
500,000
1,000,000
1,500,000
2,000,000
Type of Repetitive Sequences
Simple repeat
Others
Copy-Number Variants
Deletion
Duplication
Features
Centromeres
MAT

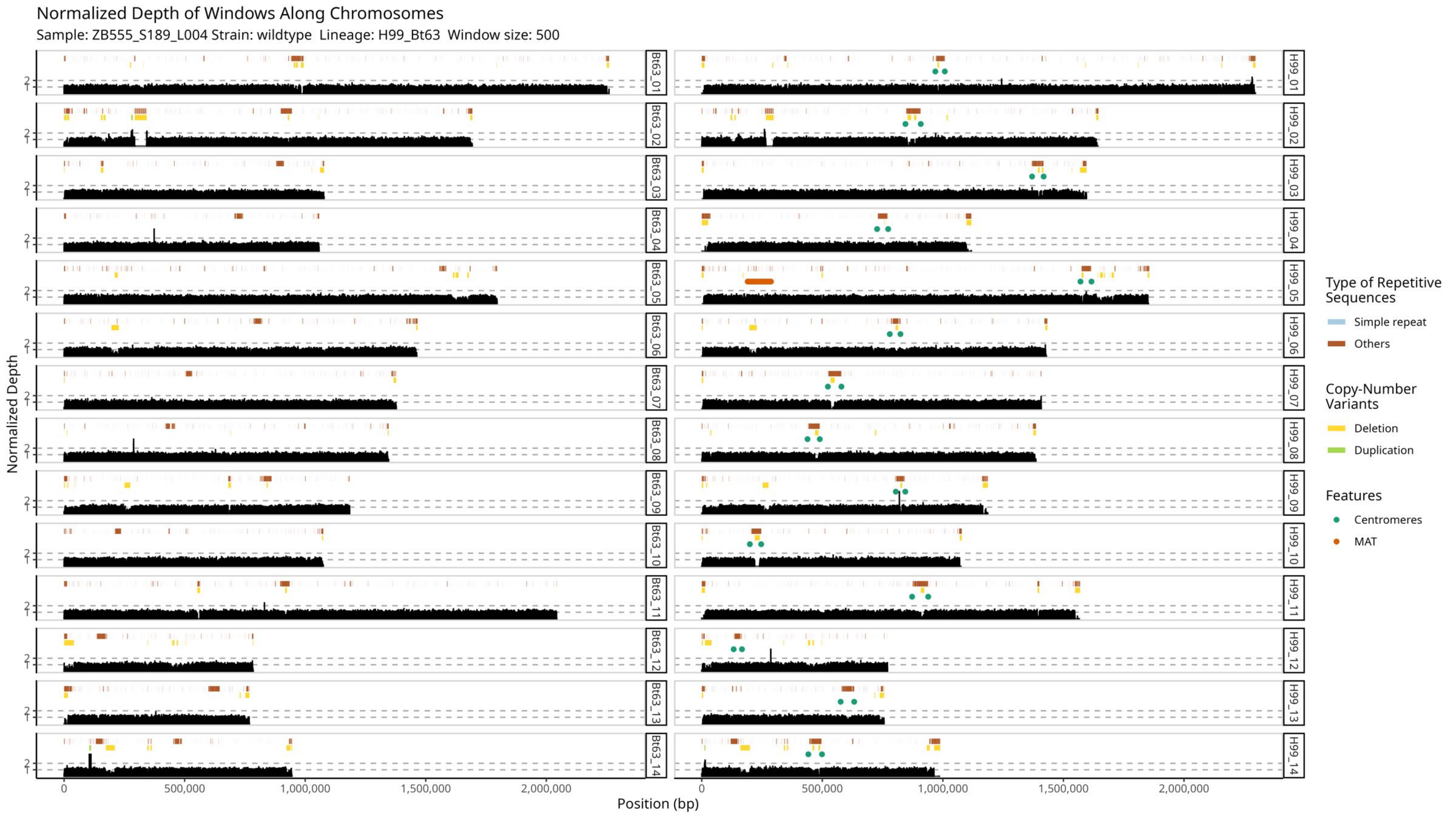

Normalized Depth of Windows Along Chromosomes
Sample: ZB555_S189_L004 Strain: wildtype Lineage: H99_Bt63 Window size: 500
Normalized Depth
Position (bp)
0
500,000
1,000,000
1,500,000
2,000,000
1
2
Bt63_01
Bt63_02
Bt63_03
Bt63_04
Bt63_05
Bt63_06
Bt63_07
Bt63_08
Bt63_09
Bt63_10
Bt63_11
Bt63_12
Bt63_13
Bt63_14
H99_01
H99_02
H99_03
H99_04
H99_05
H99_06
H99_07
H99_08
H99_09
H99_10
H99_11
H99_12
H99_13
H99_14
Type of Repetitive Sequences
Simple repeat
Others
Copy-Number Variants
Deletion
Duplication
Features
Centromeres
MAT

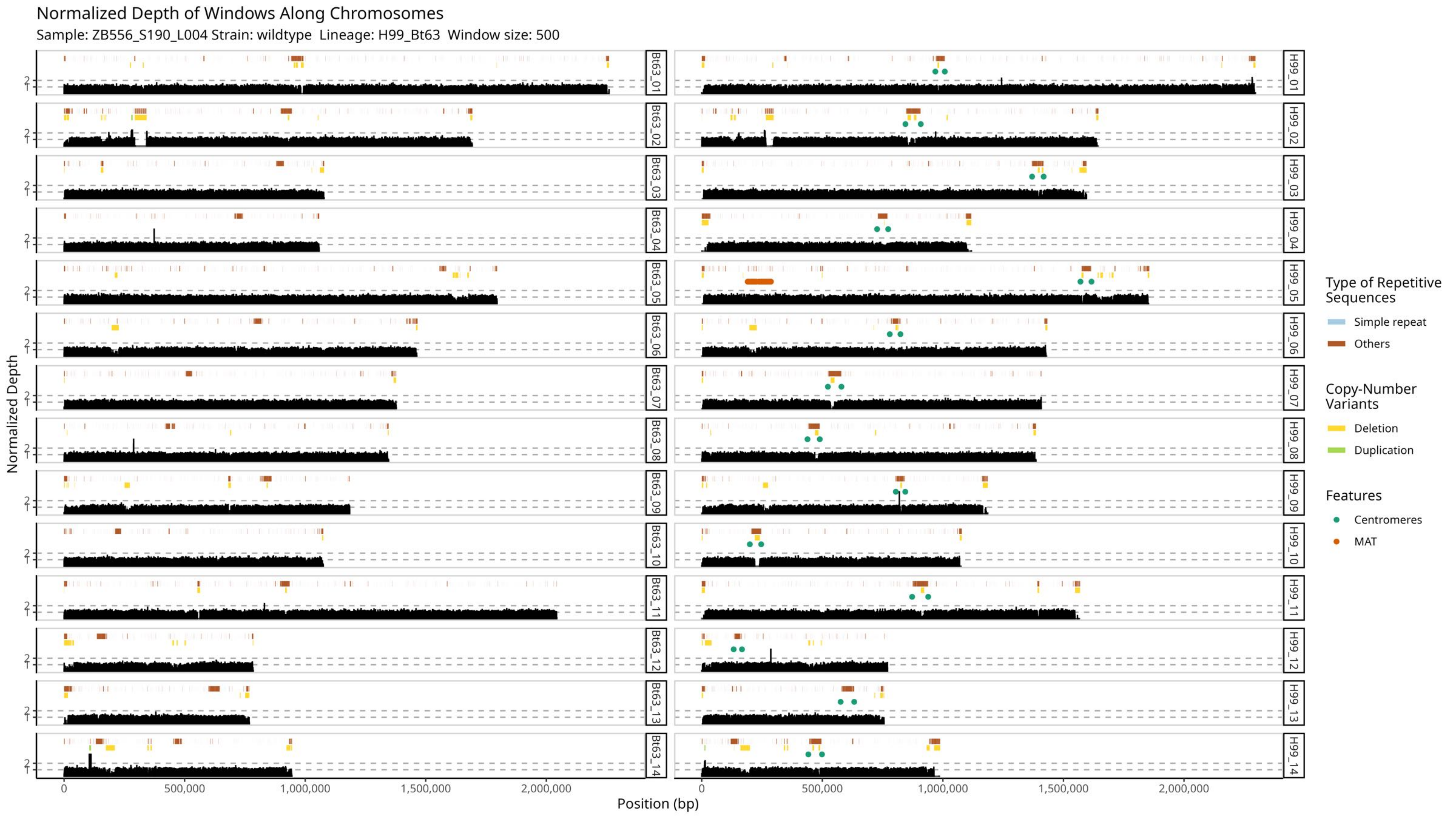
Normalized Depth of Windows Along Chromosomes
Sample: ZB556_S190_L004 Strain: wildtype Lineage: H99_Bt63 Window size: 500
Normalized Depth
Position (bp)
0
500,000
1,000,000
1,500,000
2,000,000
Bt63_01
Bt63_02
Bt63_03
Bt63_04
Bt63_05
Bt63_06
Bt63_07
Bt63_08
Bt63_09
Bt63_10
Bt63_11
Bt63_12
Bt63_13
Bt63_14
H99_01
H99_02
H99_03
H99_04
H99_05
H99_06
H99_07
H99_08
H99_09
H99_10
H99_11
H99_12
H99_13
H99_14
Type of Repetitive Sequences
Simple repeat
Others
Copy-Number Variants
Deletion
Duplication
Features
Centromeres
MAT

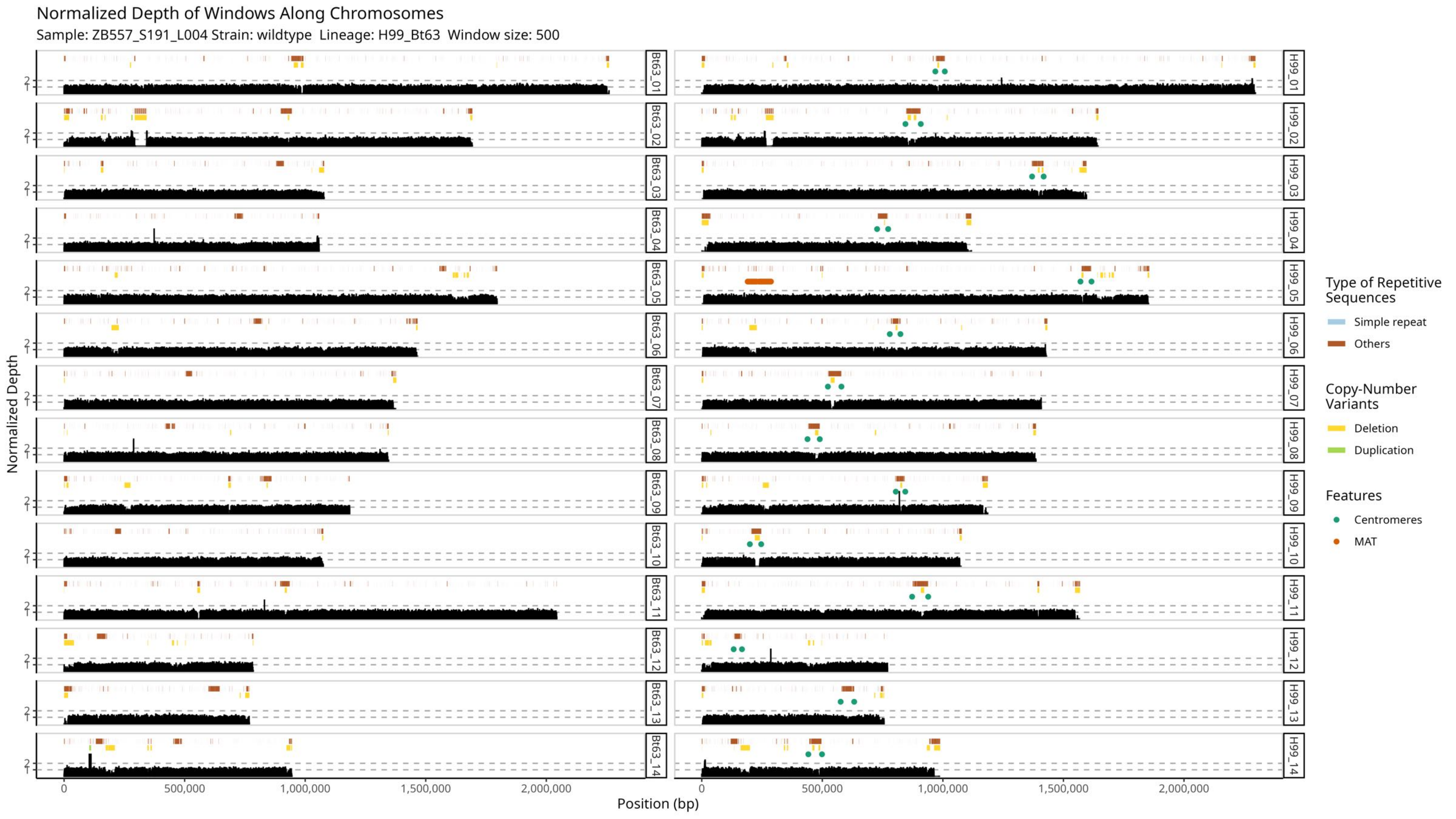

Normalized Depth of Windows Along Chromosomes
Sample: ZB557_S191_L004 Strain: wildtype Lineage: H99_Bt63 Window size: 500
Normalized Depth
Position (bp)
0
500,000
1,000,000
1,500,000
2,000,000
1
2
Bt63_01
Bt63_02
Bt63_03
Bt63_04
Bt63_05
Bt63_06
Bt63_07
Bt63_08
Bt63_09
Bt63_10
Bt63_11
Bt63_12
Bt63_13
Bt63_14
H99_01
H99_02
H99_03
H99_04
H99_05
H99_06
H99_07
H99_08
H99_09
H99_10
H99_11
H99_12
H99_13
H99_14
Type of Repetitive Sequences
Simple repeat
Others
Copy-Number Variants
Deletion
Duplication
Features
Centromeres
MAT

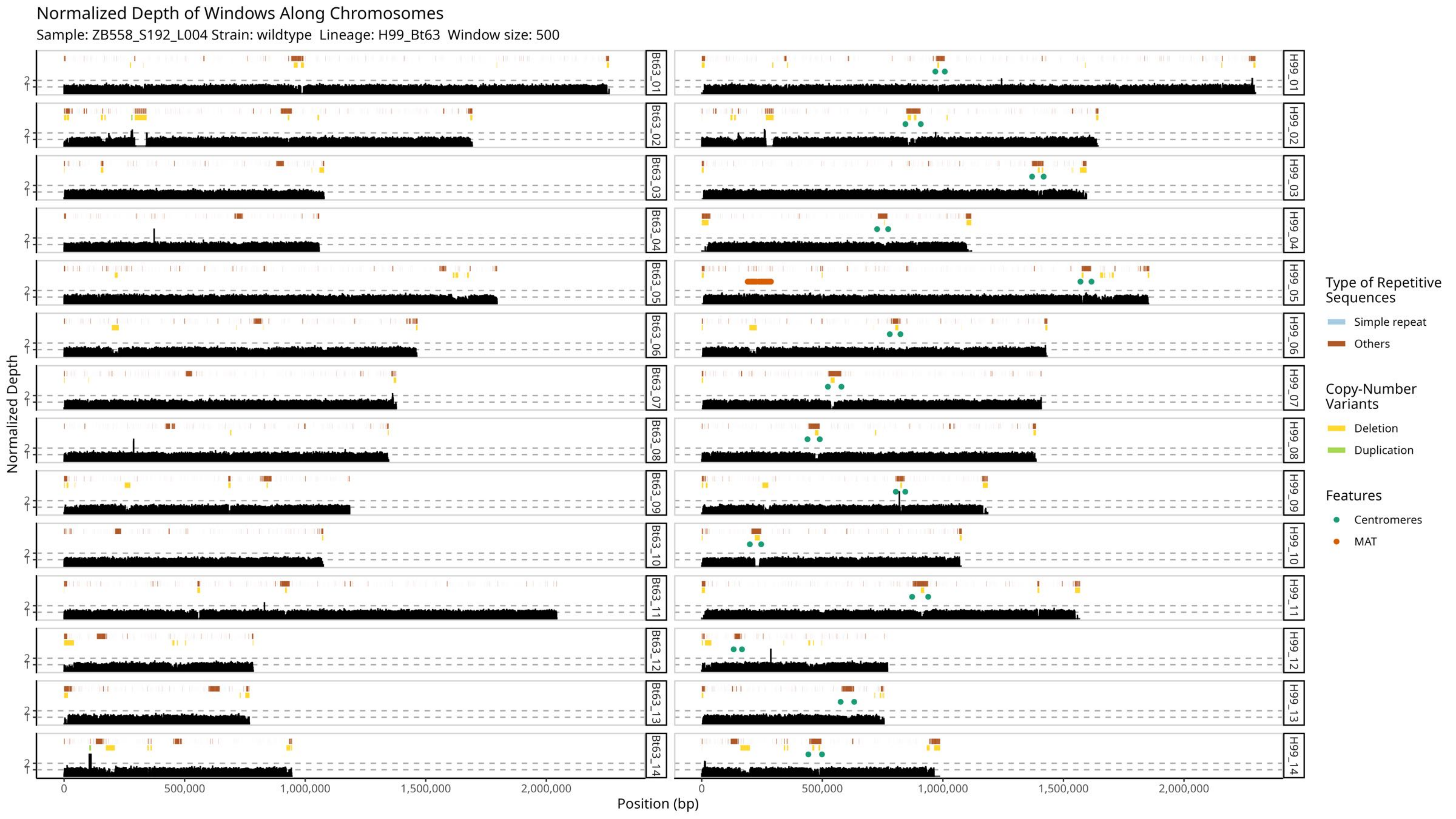

Normalized Depth of Windows Along Chromosomes
Sample: ZB558_S192_L004 Strain: wildtype Lineage: H99_Bt63 Window size: 500
Normalized Depth
Position (bp)
0
500,000
1,000,000
1,500,000
2,000,000
1
2
Bt63_01
Bt63_02
Bt63_03
Bt63_04
Bt63_05
Bt63_06
Bt63_07
Bt63_08
Bt63_09
Bt63_10
Bt63_11
Bt63_12
Bt63_13
Bt63_14
H99_01
H99_02
H99_03
H99_04
H99_05
H99_06
H99_07
H99_08
H99_09
H99_10
H99_11
H99_12
H99_13
H99_14
Type of Repetitive Sequences
Simple repeat
Others
Copy-Number Variants
Deletion
Duplication
Features
Centromeres
MAT

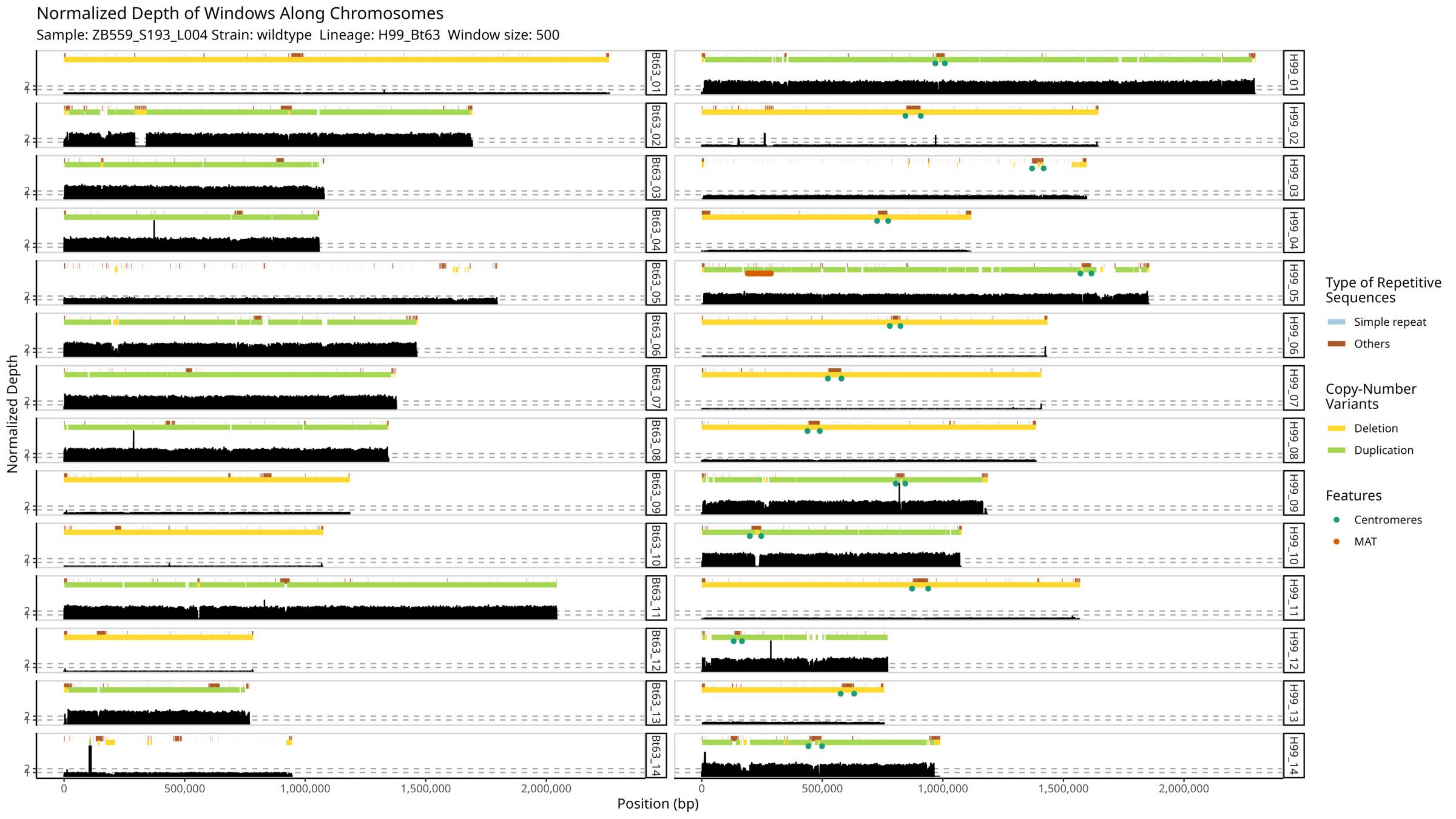

Normalized Depth of Windows Along Chromosomes
Sample: ZB559_S193_L004 Strain: wildtype Lineage: H99_Bt63 Window size: 500
Normalized Depth
Position (bp)
0
500,000
1,000,000
1,500,000
2,000,000
Bt63_01
Bt63_02
Bt63_03
Bt63_04
Bt63_05
Bt63_06
Bt63_07
Bt63_08
Bt63_09
Bt63_10
Bt63_11
Bt63_12
Bt63_13
Bt63_14
H99_01
H99_02
H99_03
H99_04
H99_05
H99_06
H99_07
H99_08
H99_09
H99_10
H99_11
H99_12
H99_13
H99_14
Type of Repetitive Sequences
Simple repeat
Others
Copy-Number Variants
Deletion
Duplication
Features
Centromeres
MAT

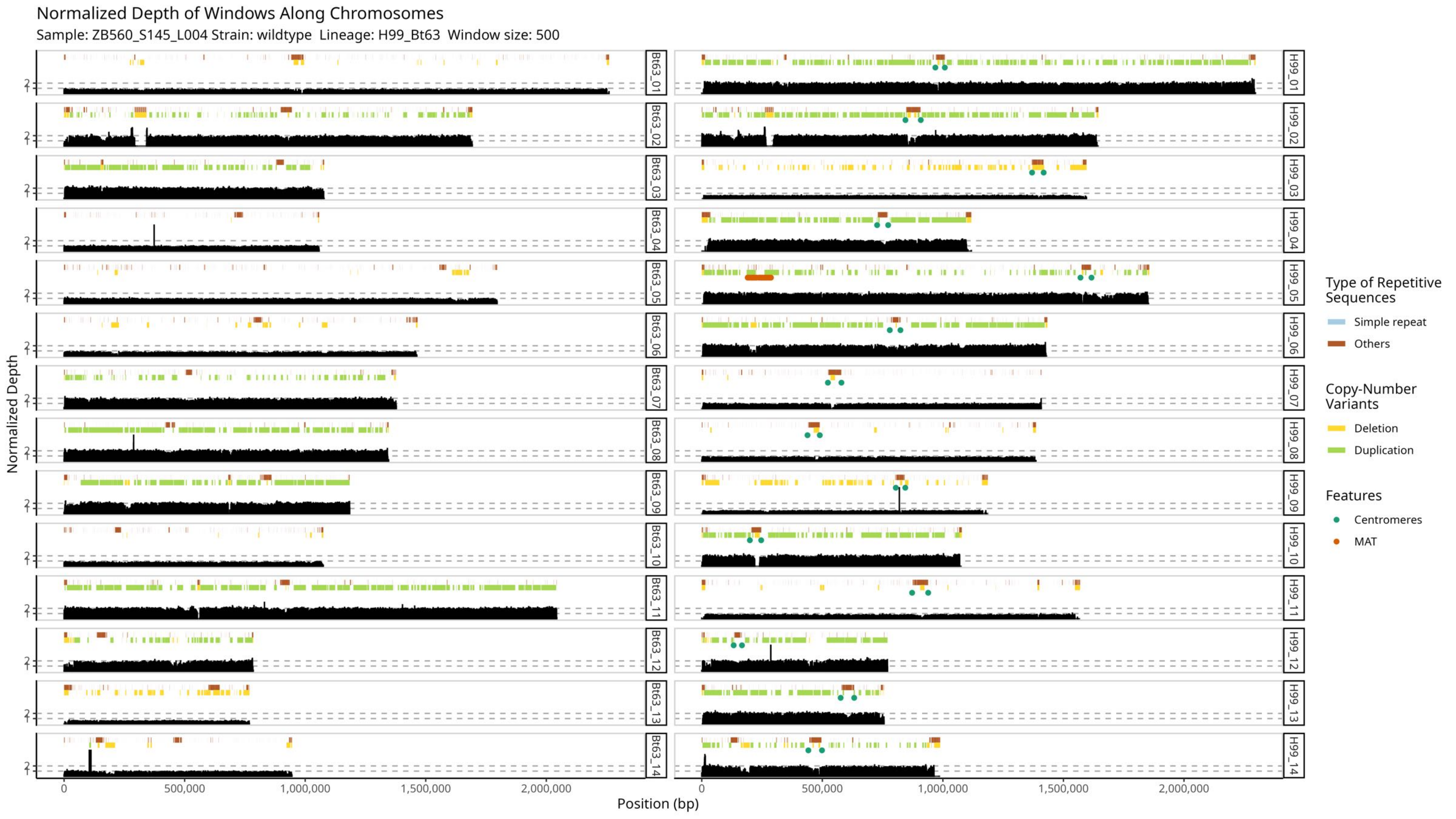

Normalized Depth of Windows Along Chromosomes
Sample: ZB560_S145_L004 Strain: wildtype Lineage: H99_Bt63 Window size: 500
Normalized Depth
Position (bp)
Bt63_01
Bt63_02
Bt63_03
Bt63_04
Bt63_05
Bt63_06
Bt63_07
Bt63_08
Bt63_09
Bt63_10
Bt63_11
Bt63_12
Bt63_13
Bt63_14
H99_01
H99_02
H99_03
H99_04
H99_05
H99_06
H99_07
H99_08
H99_09
H99_10
H99_11
H99_12
H99_13
H99_14
0
500,000
1,000,000
1,500,000
2,000,000
Type of Repetitive Sequences
Simple repeat
Others
Copy-Number Variants
Deletion
Duplication
Features
Centromeres
MAT

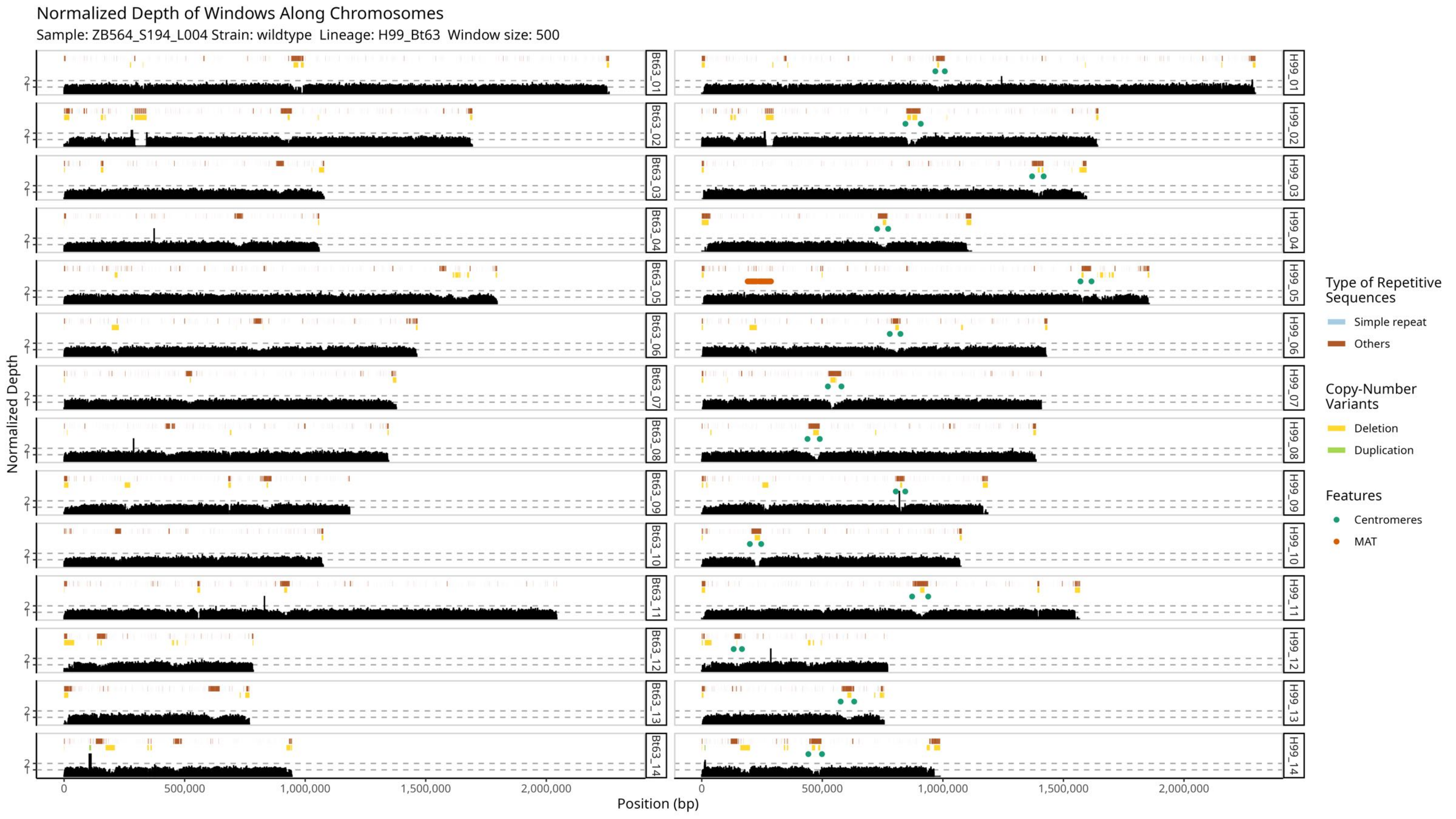

Normalized Depth of Windows Along Chromosomes
Sample: ZB564_S194_L004 Strain: wildtype Lineage: H99_Bt63 Window size: 500
Normalized Depth
Position (bp)
0
500,000
1,000,000
1,500,000
2,000,000
1
2
Bt63_01
Bt63_02
Bt63_03
Bt63_04
Bt63_05
Bt63_06
Bt63_07
Bt63_08
Bt63_09
Bt63_10
Bt63_11
Bt63_12
Bt63_13
Bt63_14
H99_01
H99_02
H99_03
H99_04
H99_05
H99_06
H99_07
H99_08
H99_09
H99_10
H99_11
H99_12
H99_13
H99_14
Type of Repetitive Sequences
Simple repeat
Others
Copy-Number Variants
Deletion
Duplication
Features
Centromeres
MAT

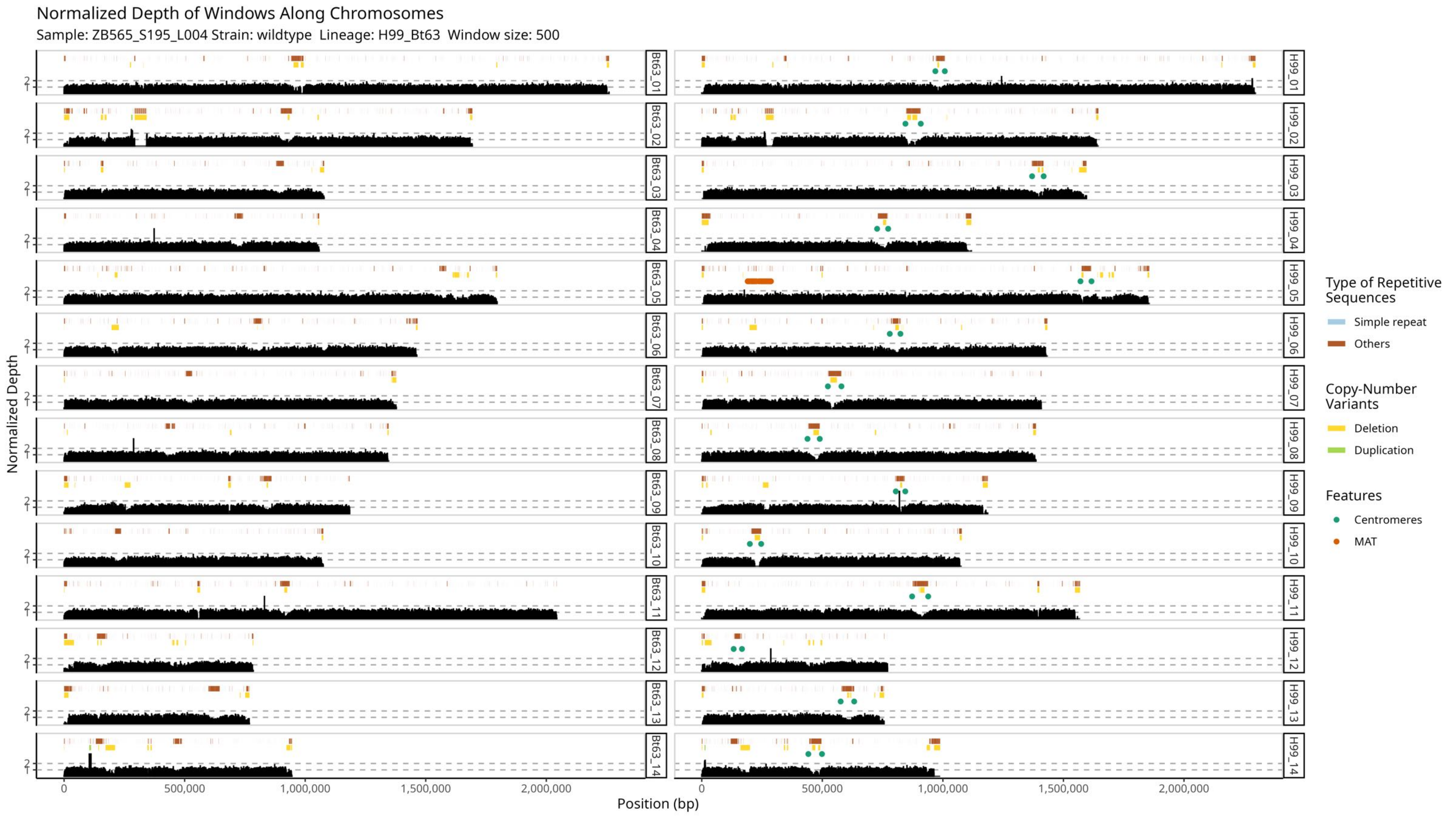
Normalized Depth of Windows Along Chromosomes
Sample: ZB565_S195_L004 Strain: wildtype Lineage: H99_Bt63 Window size: 500
Normalized Depth
Position (bp)
0
500,000
1,000,000
1,500,000
2,000,000
1
2
Bt63_01
Bt63_02
Bt63_03
Bt63_04
Bt63_05
Bt63_06
Bt63_07
Bt63_08
Bt63_09
Bt63_10
Bt63_11
Bt63_12
Bt63_13
Bt63_14
H99_01
H99_02
H99_03
H99_04
H99_05
H99_06
H99_07
H99_08
H99_09
H99_10
H99_11
H99_12
H99_13
H99_14
Type of Repetitive Sequences
Simple repeat
Others
Copy-Number Variants
Deletion
Duplication
Features
Centromeres
MAT

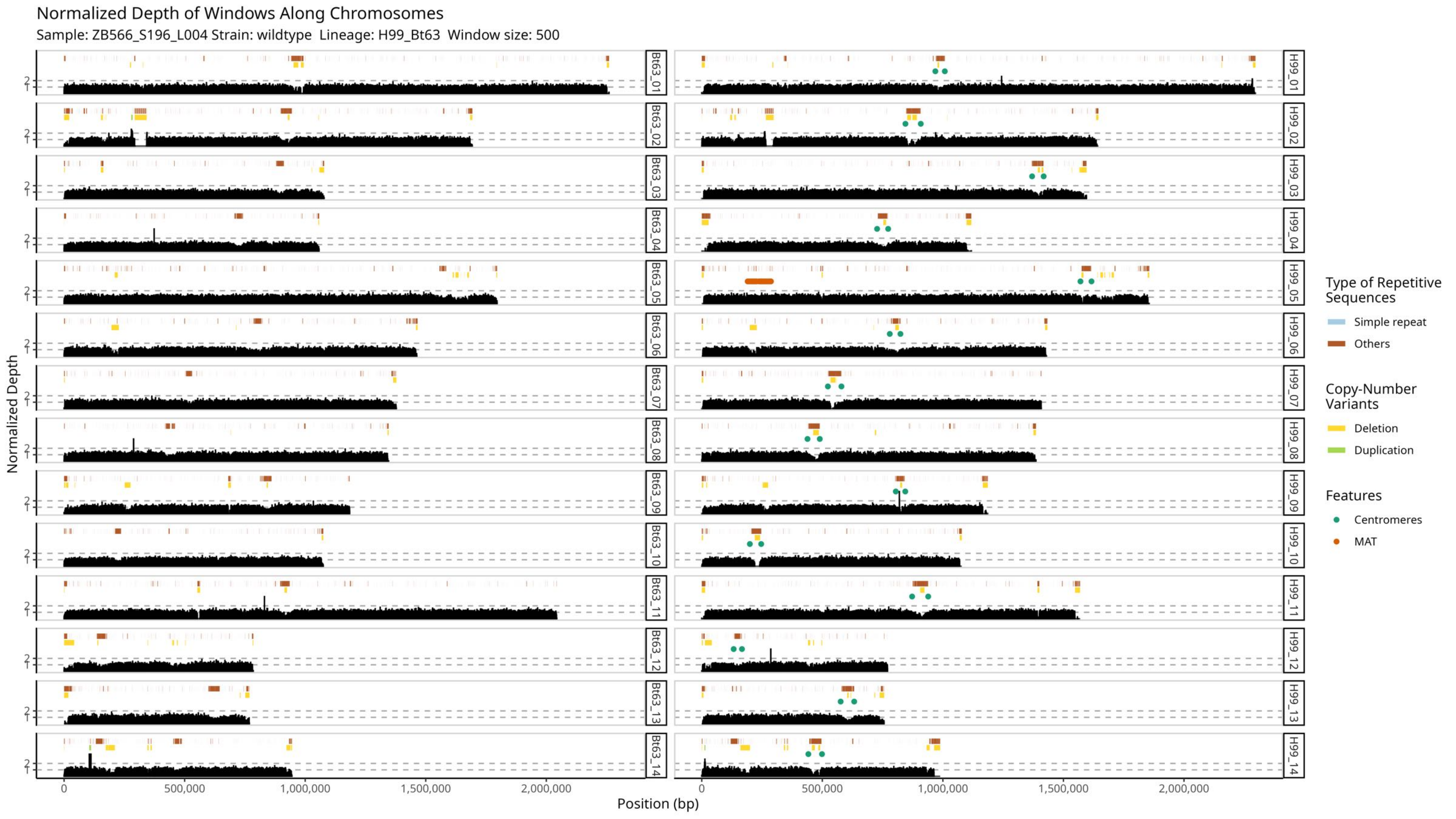

Normalized Depth of Windows Along Chromosomes
Sample: ZB566_S196_L004 Strain: wildtype Lineage: H99_Bt63 Window size: 500
Normalized Depth
Position (bp)
0
500,000
1,000,000
1,500,000
2,000,000
1
2
Bt63_01
Bt63_02
Bt63_03
Bt63_04
Bt63_05
Bt63_06
Bt63_07
Bt63_08
Bt63_09
Bt63_10
Bt63_11
Bt63_12
Bt63_13
Bt63_14
H99_01
H99_02
H99_03
H99_04
H99_05
H99_06
H99_07
H99_08
H99_09
H99_10
H99_11
H99_12
H99_13
H99_14
Type of Repetitive Sequences
Simple repeat
Others
Copy-Number Variants
Deletion
Duplication
Features
Centromeres
MAT

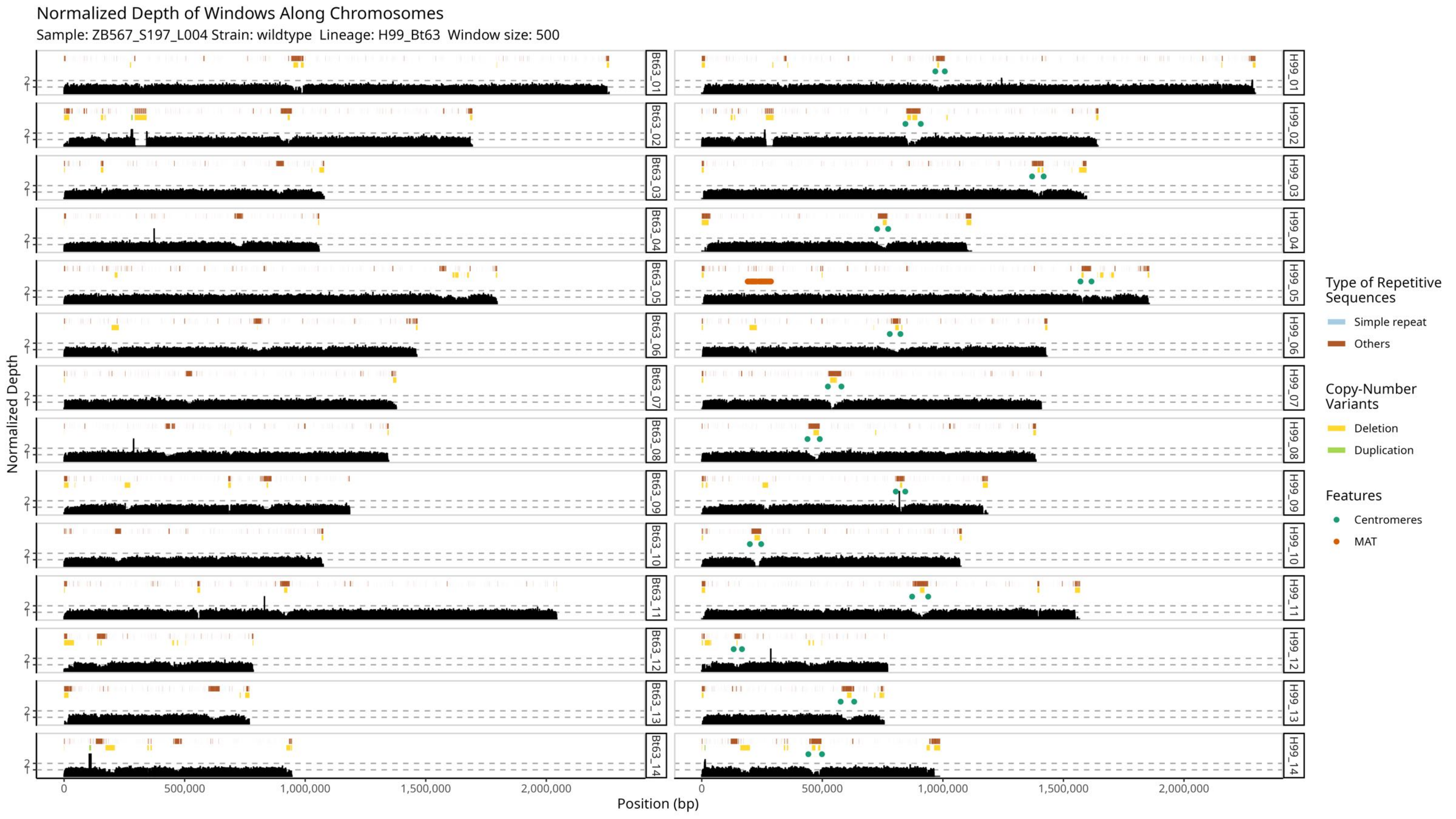

Normalized Depth of Windows Along Chromosomes
Sample: ZB567_S197_L004 Strain: wildtype Lineage: H99_Bt63 Window size: 500
Bt63_01
Bt63_02
Bt63_03
Bt63_04
Bt63_05
Bt63_06
Bt63_07
Bt63_08
Bt63_09
Bt63_10
Bt63_11
Bt63_12
Bt63_13
Bt63_14
H99_01
H99_02
H99_03
H99_04
H99_05
H99_06
H99_07
H99_08
H99_09
H99_10
H99_11
H99_12
H99_13
H99_14
Normalized Depth
Position (bp)
0
500,000
1,000,000
1,500,000
2,000,000
Type of Repetitive Sequences
Simple repeat
Others
Copy-Number Variants
Deletion
Duplication
Features
Centromeres
MAT

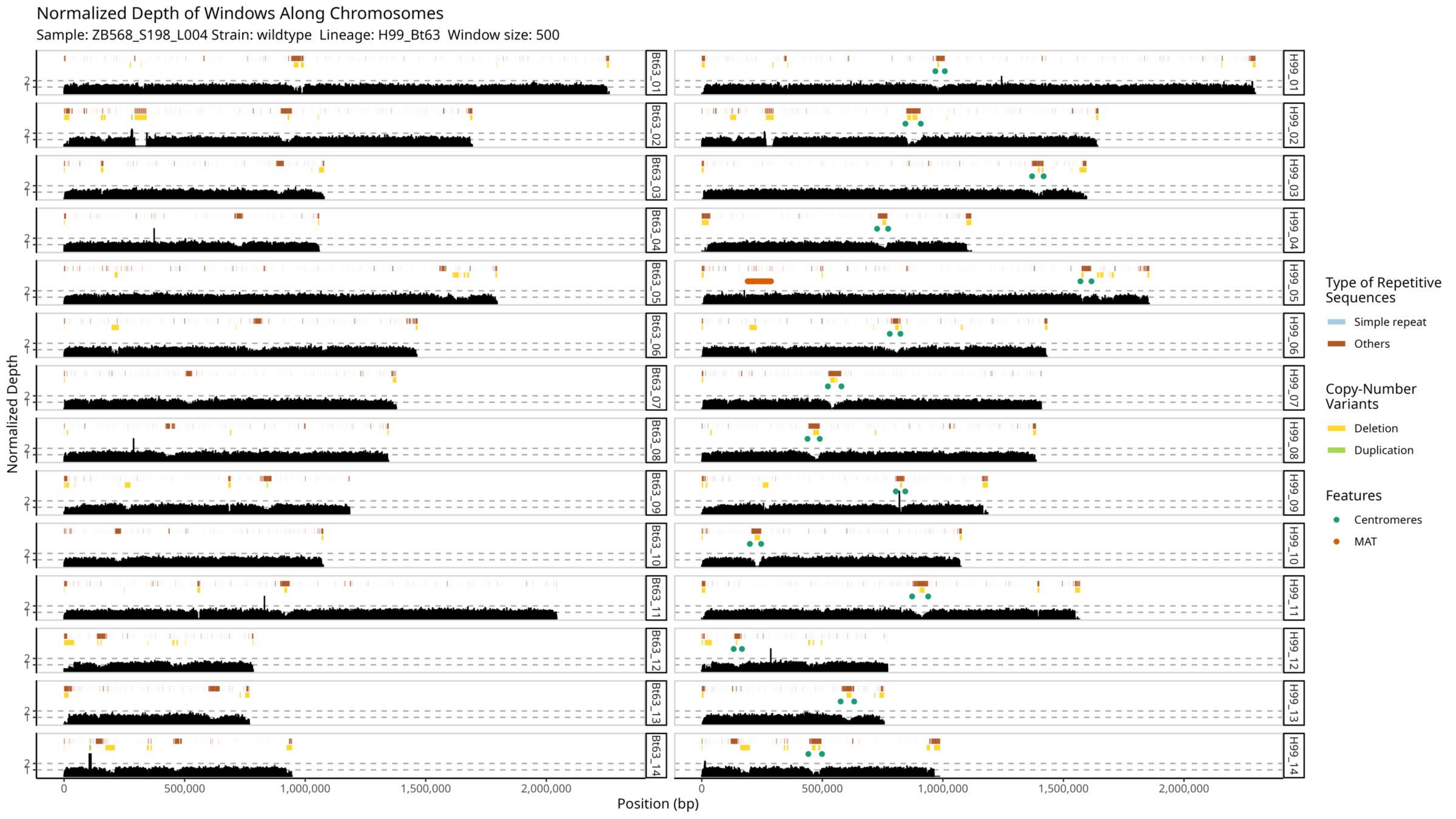
Normalized Depth of Windows Along Chromosomes
Sample: ZB568_S198_L004 Strain: wildtype Lineage: H99_Bt63 Window size: 500
Normalized Depth
Position (bp)
0
500,000
1,000,000
1,500,000
2,000,000
Bt63_01
Bt63_02
Bt63_03
Bt63_04
Bt63_05
Bt63_06
Bt63_07
Bt63_08
Bt63_09
Bt63_10
Bt63_11
Bt63_12
Bt63_13
Bt63_14
H99_01
H99_02
H99_03
H99_04
H99_05
H99_06
H99_07
H99_08
H99_09
H99_10
H99_11
H99_12
H99_13
H99_14
Type of Repetitive Sequences
Simple repeat
Others
Copy-Number Variants
Deletion
Duplication
Features
Centromeres
MAT

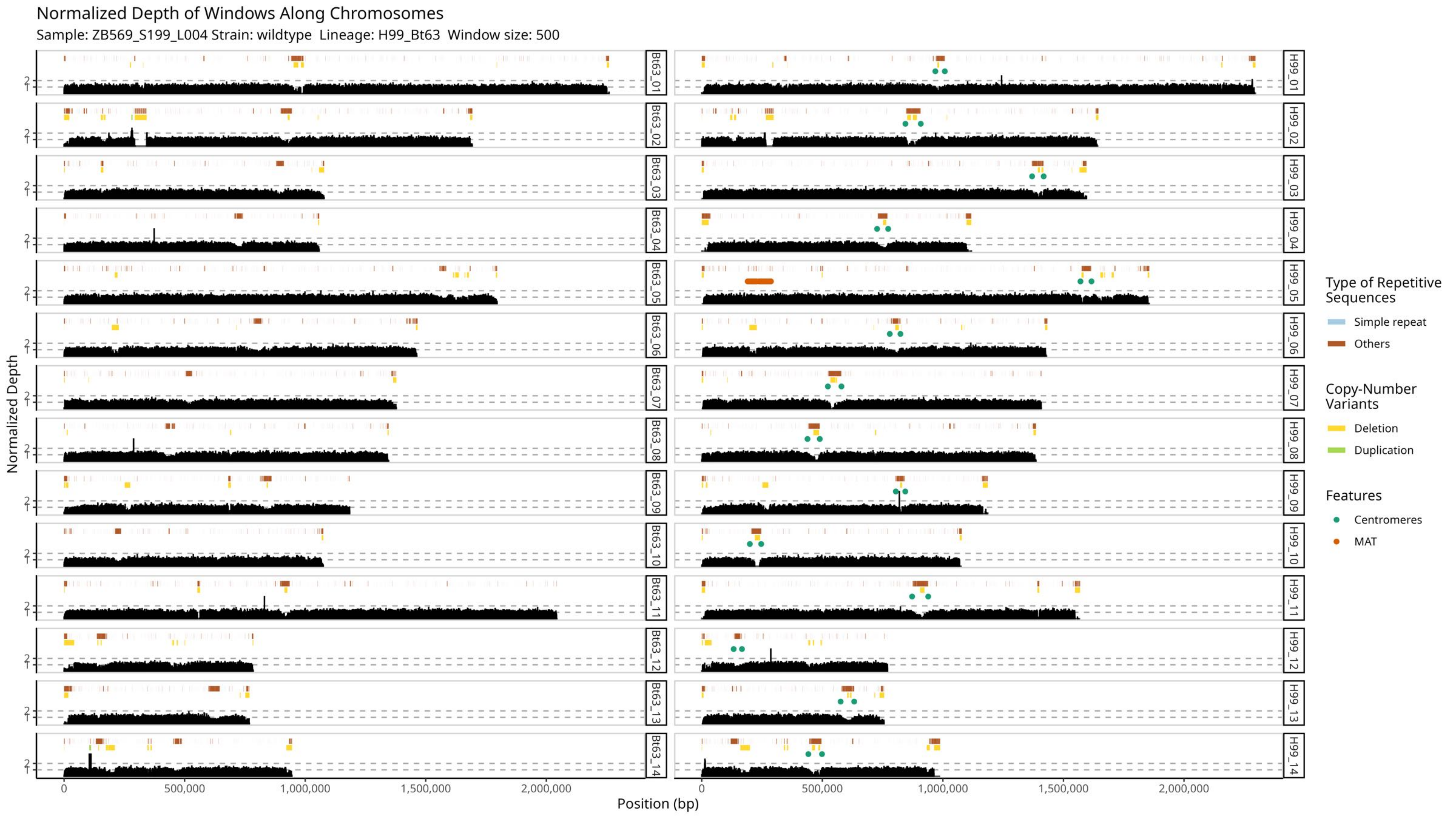

Normalized Depth of Windows Along Chromosomes
Sample: ZB569_S199_L004 Strain: wildtype Lineage: H99_Bt63 Window size: 500
Normalized Depth
Position (bp)
0
500,000
1,000,000
1,500,000
2,000,000
1
2
Bt63_01
Bt63_02
Bt63_03
Bt63_04
Bt63_05
Bt63_06
Bt63_07
Bt63_08
Bt63_09
Bt63_10
Bt63_11
Bt63_12
Bt63_13
Bt63_14
H99_01
H99_02
H99_03
H99_04
H99_05
H99_06
H99_07
H99_08
H99_09
H99_10
H99_11
H99_12
H99_13
H99_14
Type of Repetitive Sequences
Simple repeat
Others
Copy-Number Variants
Deletion
Duplication
Features
Centromeres
MAT

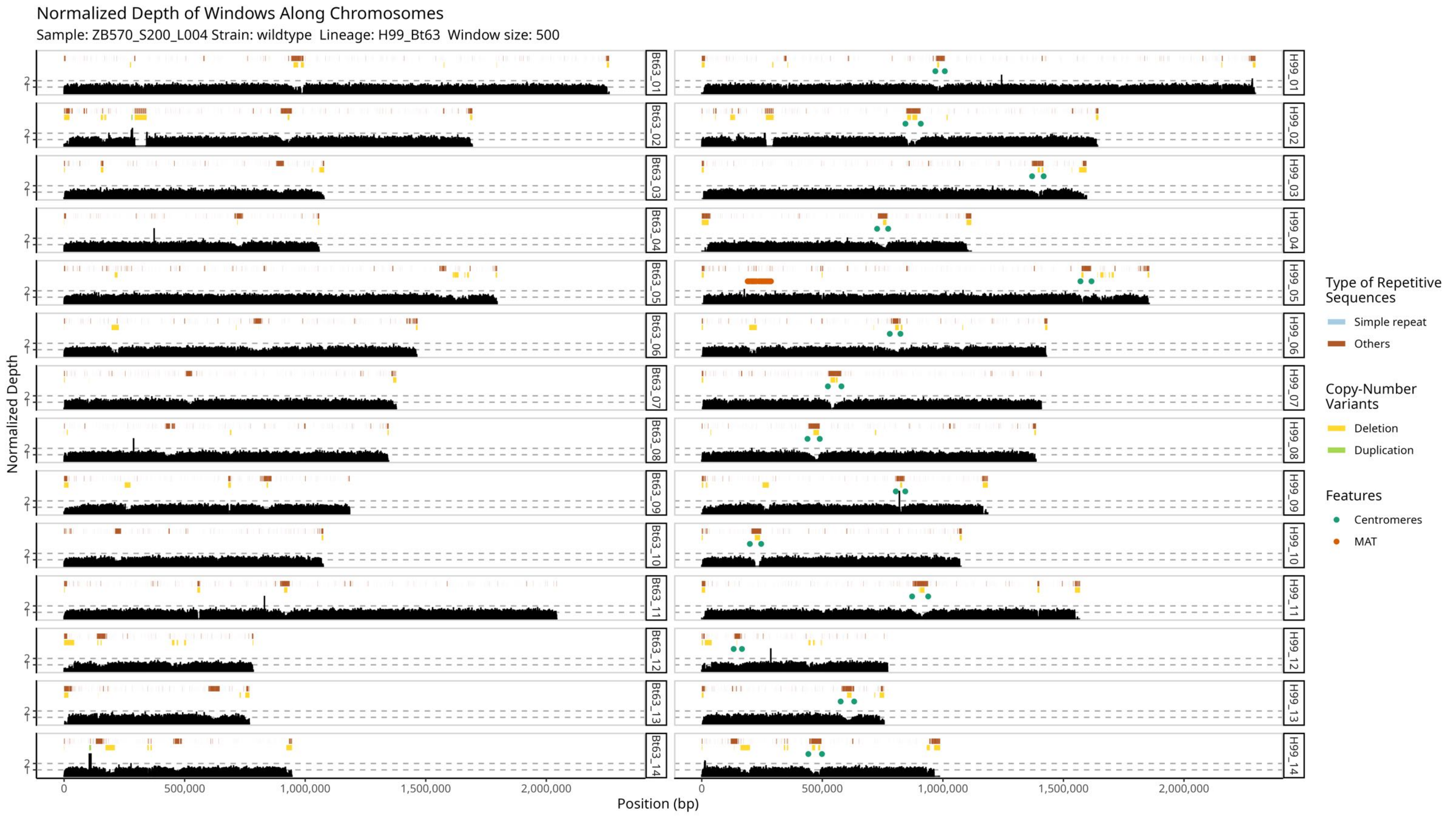

Normalized Depth of Windows Along Chromosomes
Sample: ZB570_S200_L004 Strain: wildtype Lineage: H99_Bt63 Window size: 500
Normalized Depth
Position (bp)
Bt63_01
Bt63_02
Bt63_03
Bt63_04
Bt63_05
Bt63_06
Bt63_07
Bt63_08
Bt63_09
Bt63_10
Bt63_11
Bt63_12
Bt63_13
Bt63_14
H99_01
H99_02
H99_03
H99_04
H99_05
H99_06
H99_07
H99_08
H99_09
H99_10
H99_11
H99_12
H99_13
H99_14
0
500,000
1,000,000
1,500,000
2,000,000
Type of Repetitive Sequences
Simple repeat
Others
Copy-Number Variants
Deletion
Duplication
Features
Centromeres
MAT

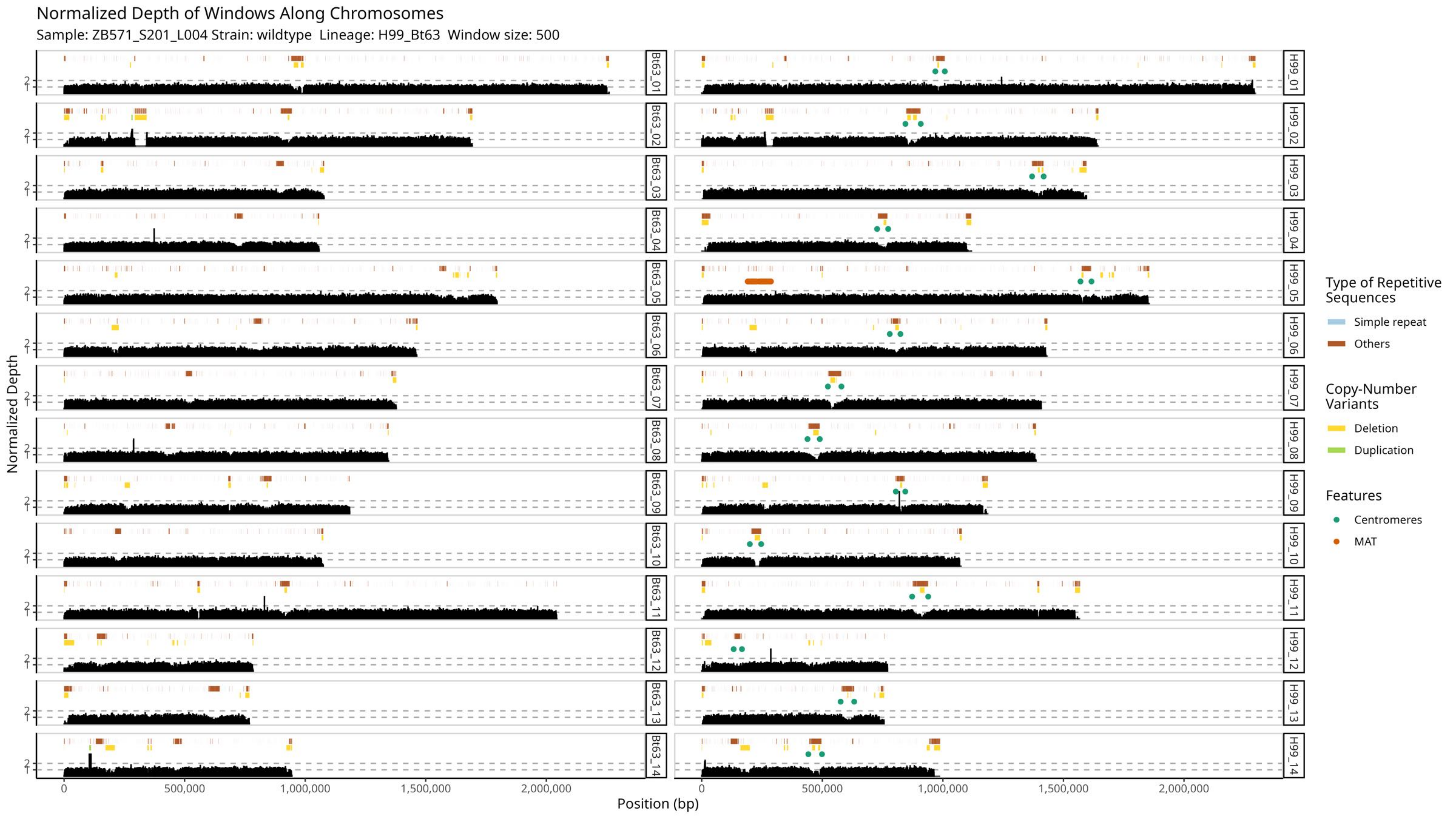

Normalized Depth of Windows Along Chromosomes
Sample: ZB571_S201_L004 Strain: wildtype Lineage: H99_Bt63 Window size: 500
Bt63_01
Bt63_02
Bt63_03
Bt63_04
Bt63_05
Bt63_06
Bt63_07
Bt63_08
Bt63_09
Bt63_10
Bt63_11
Bt63_12
Bt63_13
Bt63_14
H99_01
H99_02
H99_03
H99_04
H99_05
H99_06
H99_07
H99_08
H99_09
H99_10
H99_11
H99_12
H99_13
H99_14
Normalized Depth
Position (bp)
0
500,000
1,000,000
1,500,000
2,000,000
Type of Repetitive Sequences
Simple repeat
Others
Copy-Number Variants
Deletion
Duplication
Features
Centromeres
MAT

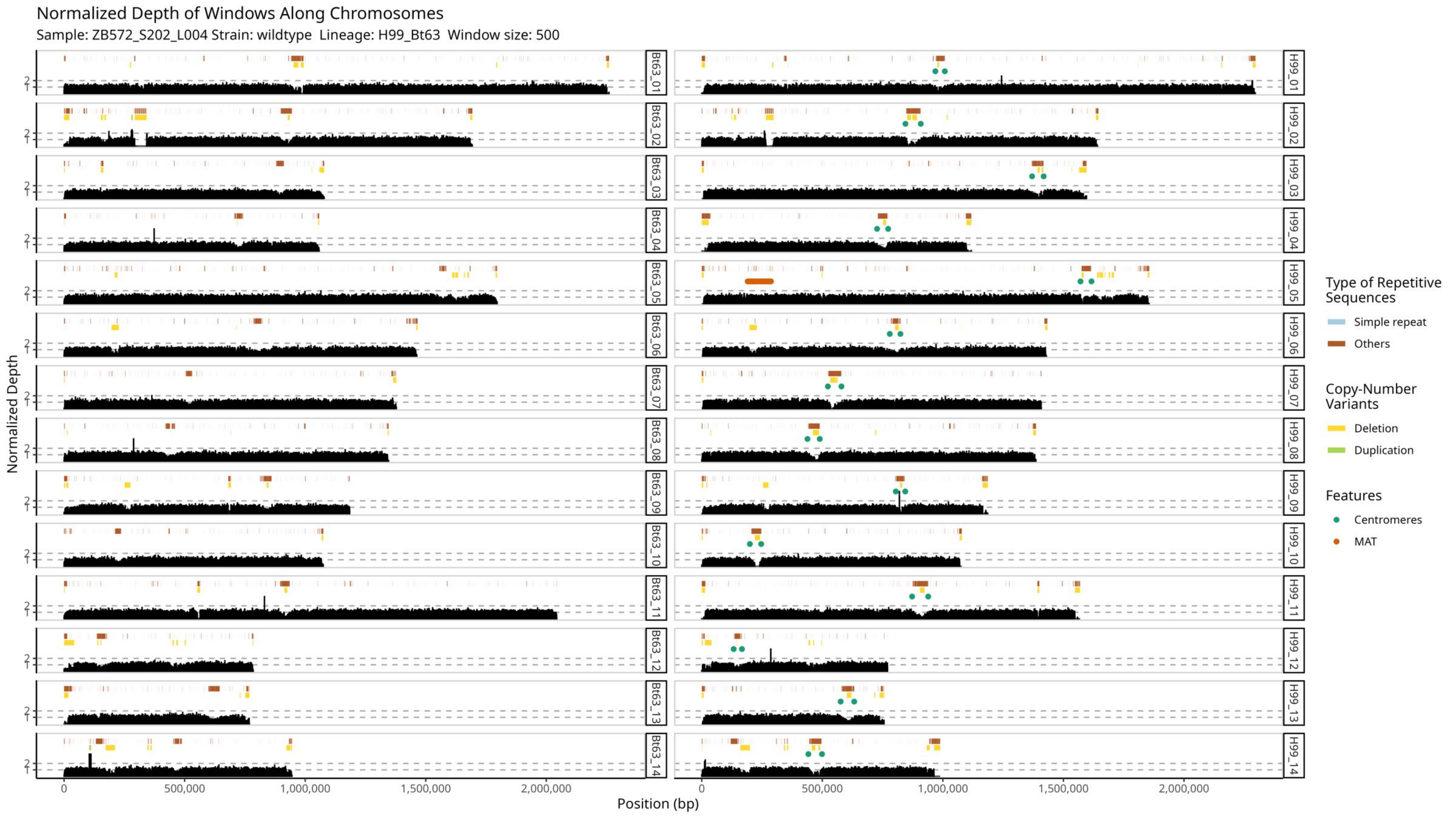

Normalized Depth of Windows Along Chromosomes
Sample: ZB572_S202_L004 Strain: wildtype Lineage: H99_Bt63 Window size: 500
Normalized Depth
Position (bp)
0
500,000
1,000,000
1,500,000
2,000,000
Bt63_01
Bt63_02
Bt63_03
Bt63_04
Bt63_05
Bt63_06
Bt63_07
Bt63_08
Bt63_09
Bt63_10
Bt63_11
Bt63_12
Bt63_13
Bt63_14
H99_01
H99_02
H99_03
H99_04
H99_05
H99_06
H99_07
H99_08
H99_09
H99_10
H99_11
H99_12
H99_13
H99_14
Type of Repetitive Sequences
Simple repeat
Others
Copy-Number Variants
Deletion
Duplication
Features
Centromeres
MAT

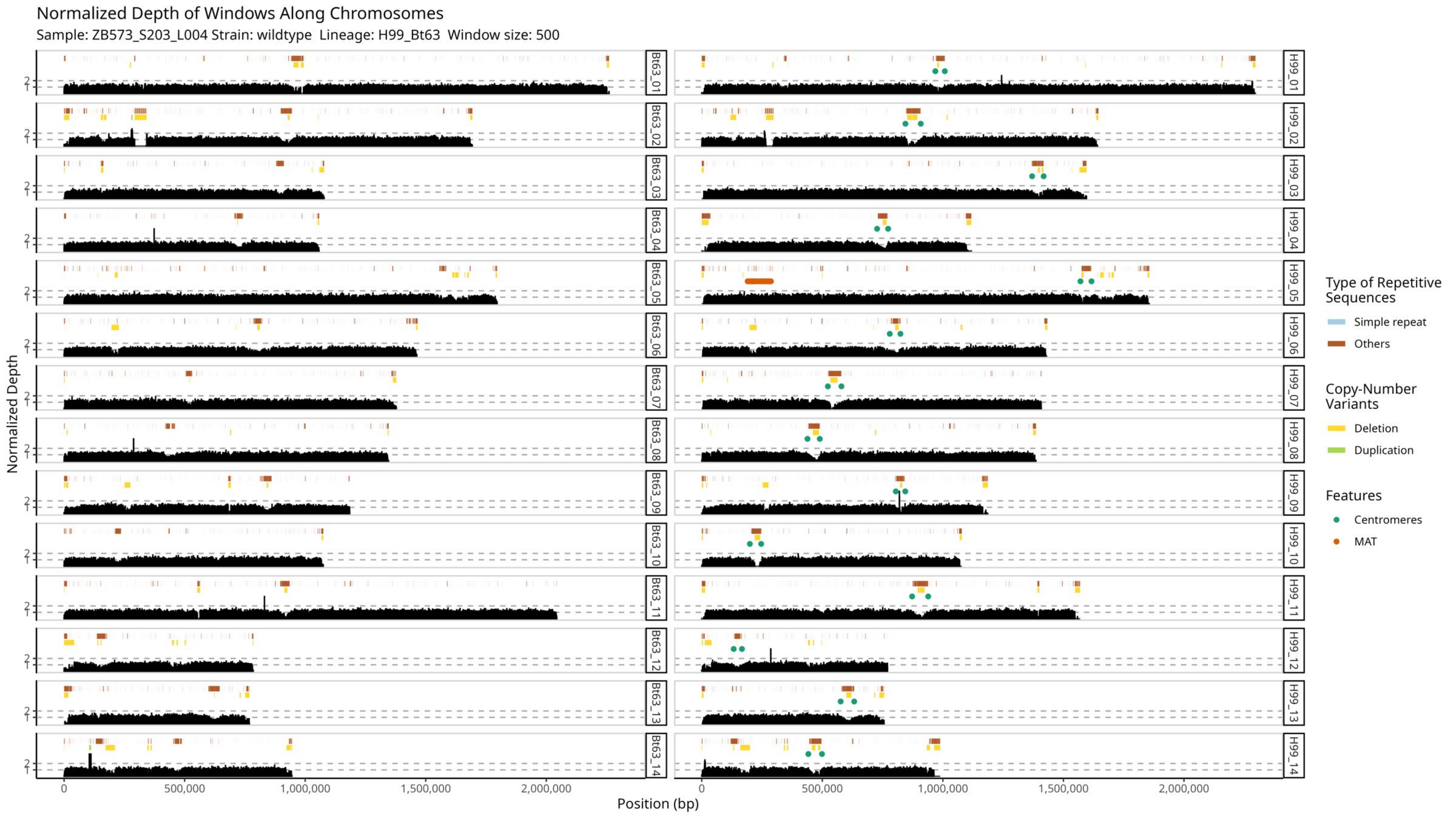

Normalized Depth of Windows Along Chromosomes
Sample: ZB573_S203_L004 Strain: wildtype Lineage: H99_Bt63 Window size: 500
Normalized Depth
Position (bp)
0
500,000
1,000,000
1,500,000
2,000,000
1
2
Bt63_01
Bt63_02
Bt63_03
Bt63_04
Bt63_05
Bt63_06
Bt63_07
Bt63_08
Bt63_09
Bt63_10
Bt63_11
Bt63_12
Bt63_13
Bt63_14
H99_01
H99_02
H99_03
H99_04
H99_05
H99_06
H99_07
H99_08
H99_09
H99_10
H99_11
H99_12
H99_13
H99_14
Type of Repetitive Sequences
Simple repeat
Others
Copy-Number Variants
Deletion
Duplication
Features
Centromeres
MAT

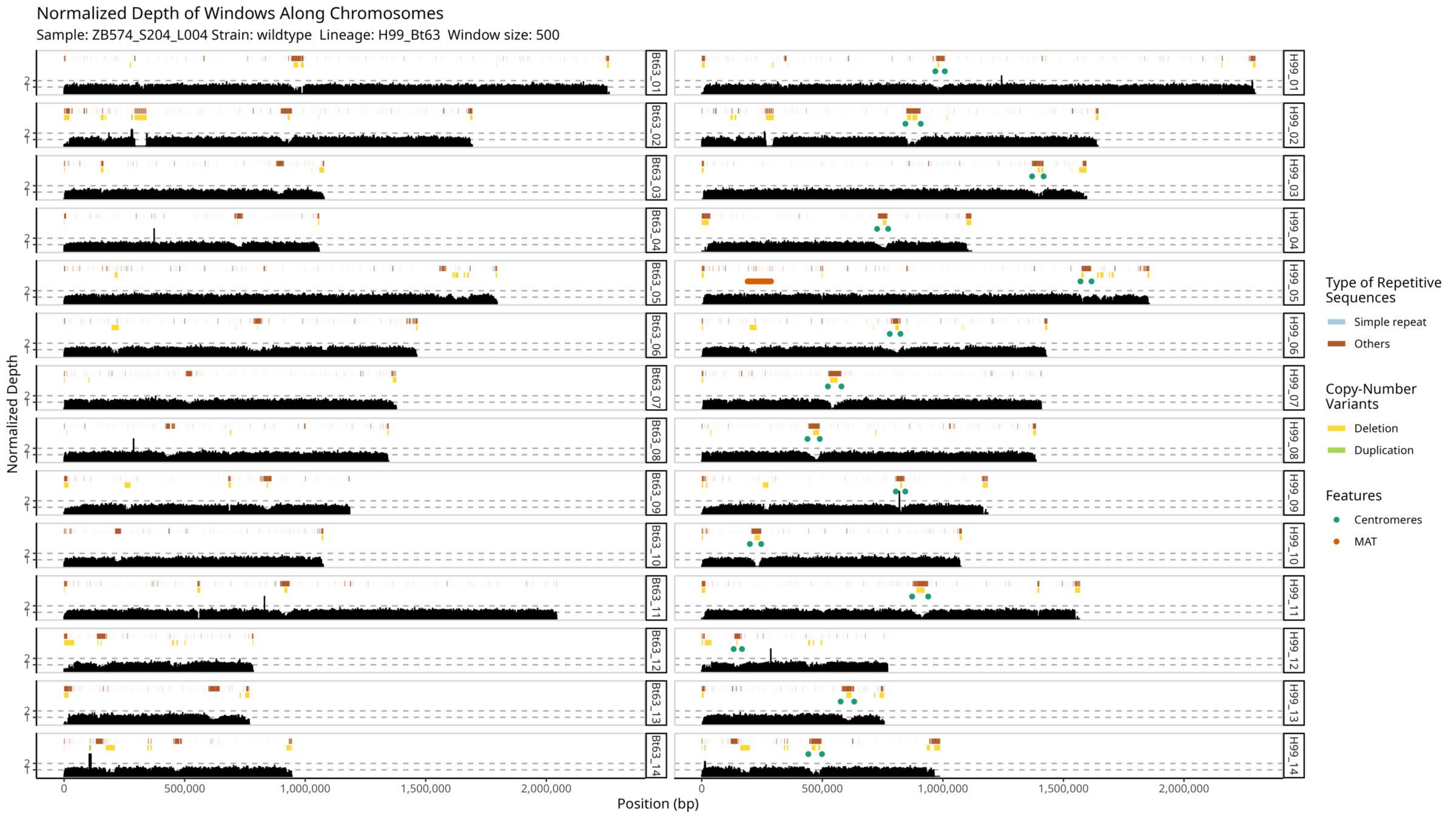

Normalized Depth of Windows Along Chromosomes
Sample: ZB574_S204_L004 Strain: wildtype Lineage: H99_Bt63 Window size: 500
Normalized Depth
Position (bp)
Bt63_01
Bt63_02
Bt63_03
Bt63_04
Bt63_05
Bt63_06
Bt63_07
Bt63_08
Bt63_09
Bt63_10
Bt63_11
Bt63_12
Bt63_13
Bt63_14
H99_01
H99_02
H99_03
H99_04
H99_05
H99_06
H99_07
H99_08
H99_09
H99_10
H99_11
H99_12
H99_13
H99_14
0
500,000
1,000,000
1,500,000
2,000,000
Type of Repetitive Sequences
Simple repeat
Others
Copy-Number Variants
Deletion
Duplication
Features
Centromeres
MAT

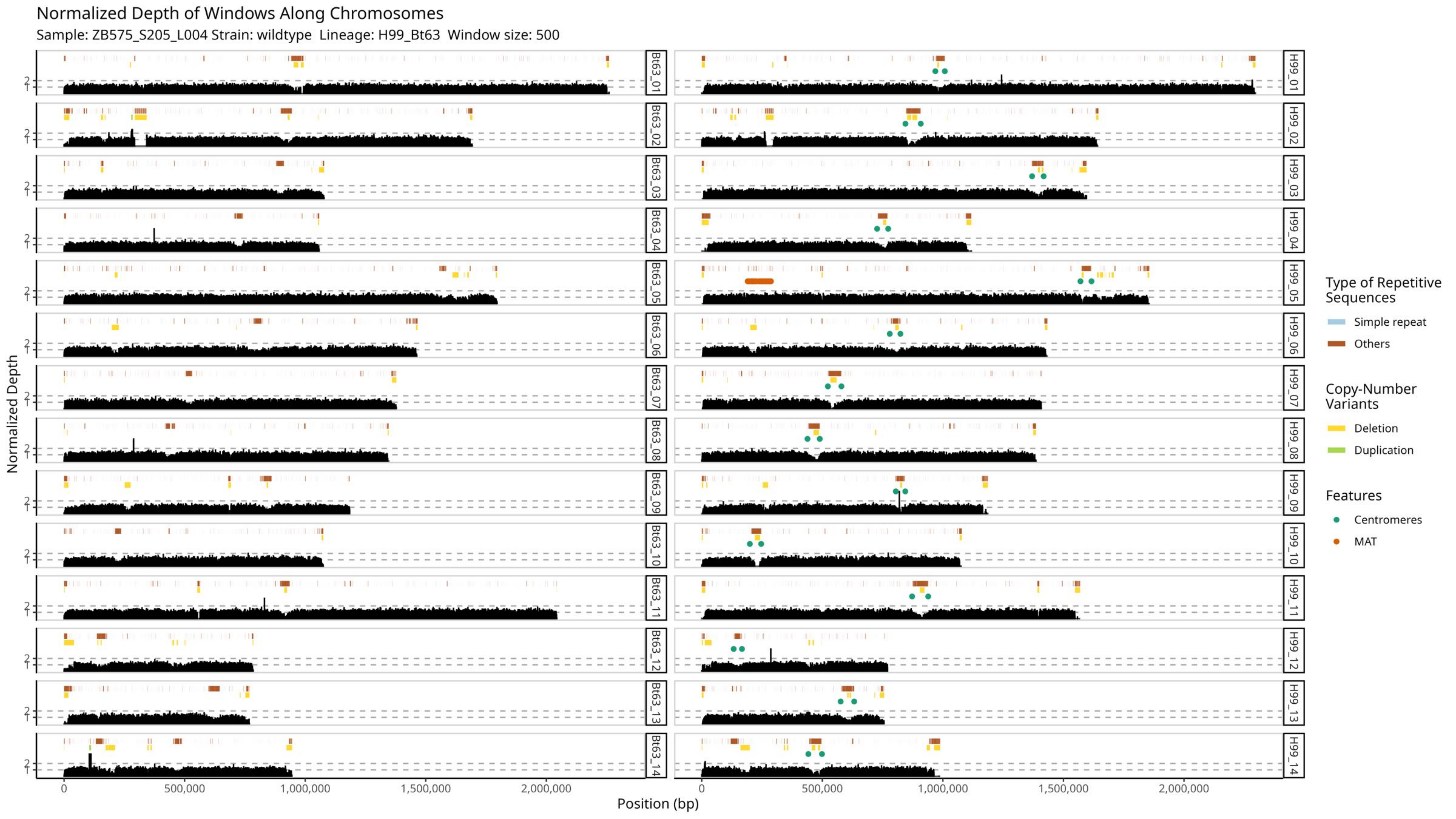
Normalized Depth of Windows Along Chromosomes
Sample: ZB575_S205_L004 Strain: wildtype Lineage: H99_Bt63 Window size: 500
Normalized Depth
Position (bp)
0
500,000
1,000,000
1,500,000
2,000,000
Bt63_01
Bt63_02
Bt63_03
Bt63_04
Bt63_05
Bt63_06
Bt63_07
Bt63_08
Bt63_09
Bt63_10
Bt63_11
Bt63_12
Bt63_13
Bt63_14
H99_01
H99_02
H99_03
H99_04
H99_05
H99_06
H99_07
H99_08
H99_09
H99_10
H99_11
H99_12
H99_13
H99_14
Type of Repetitive Sequences
Simple repeat
Others
Copy-Number Variants
Deletion
Duplication
Features
Centromeres
MAT

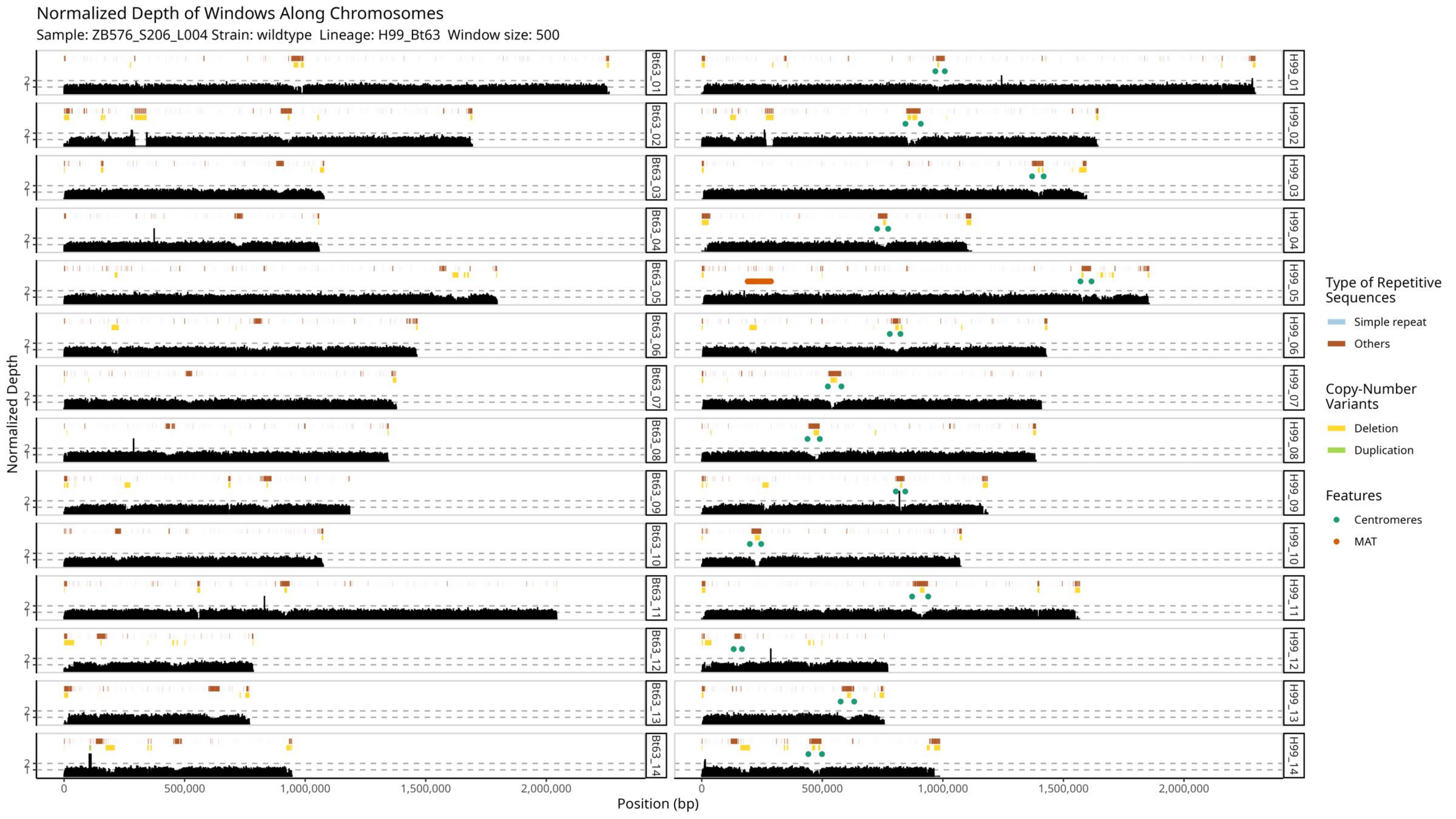

Normalized Depth of Windows Along Chromosomes
Sample: ZB576_S206_L004 Strain: wildtype Lineage: H99_Bt63 Window size: 500
Normalized Depth
Position (bp)
0
500,000
1,000,000
1,500,000
2,000,000
1
2
Bt63_01
Bt63_02
Bt63_03
Bt63_04
Bt63_05
Bt63_06
Bt63_07
Bt63_08
Bt63_09
Bt63_10
Bt63_11
Bt63_12
Bt63_13
Bt63_14
H99_01
H99_02
H99_03
H99_04
H99_05
H99_06
H99_07
H99_08
H99_09
H99_10
H99_11
H99_12
H99_13
H99_14
Type of Repetitive Sequences
Simple repeat
Others
Copy-Number Variants
Deletion
Duplication
Features
Centromeres
MAT

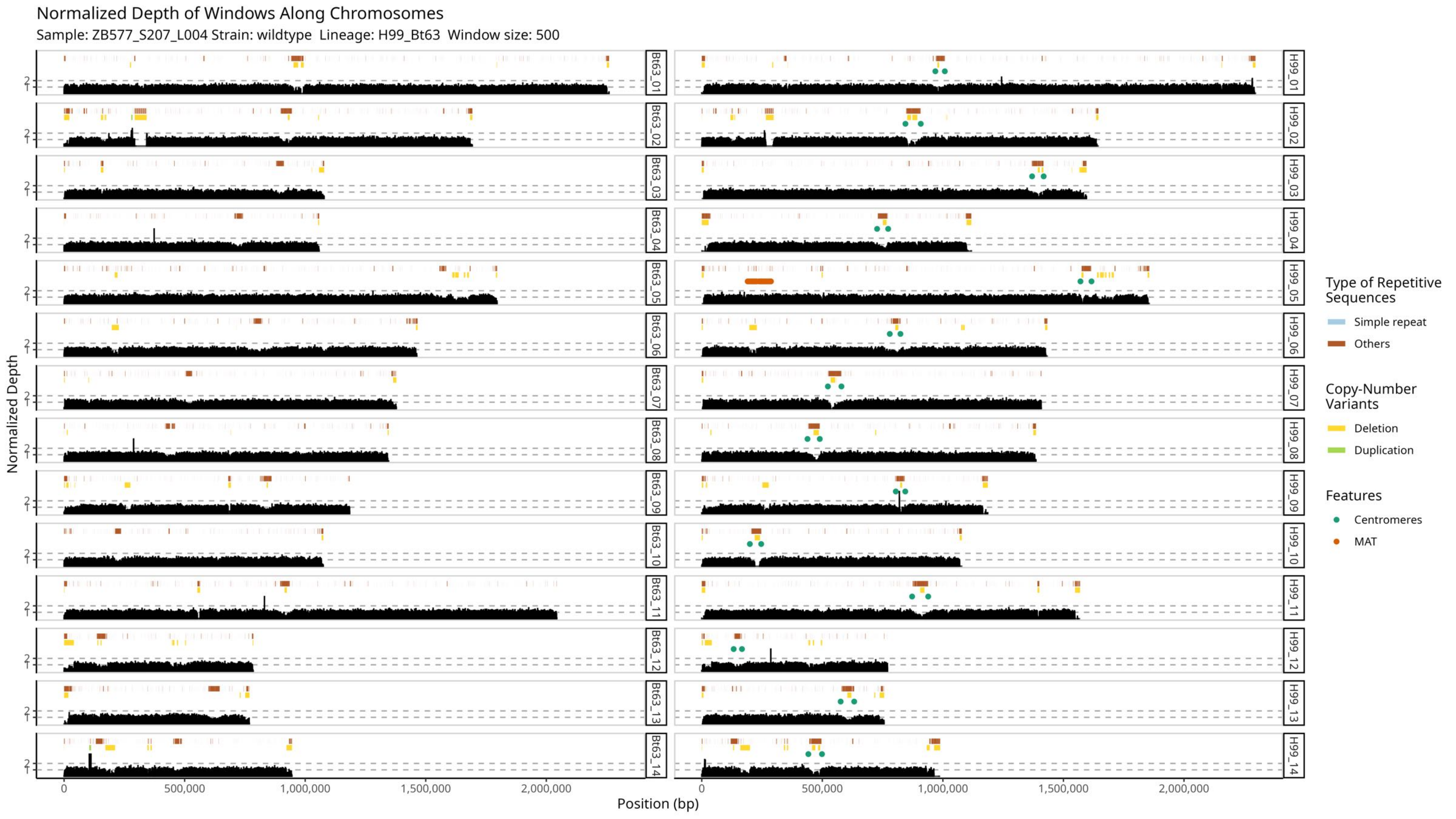

Normalized Depth of Windows Along Chromosomes
Sample: ZB577_S207_L004 Strain: wildtype Lineage: H99_Bt63 Window size: 500
Normalized Depth
Position (bp)
0
500,000
1,000,000
1,500,000
2,000,000
Bt63_01
Bt63_02
Bt63_03
Bt63_04
Bt63_05
Bt63_06
Bt63_07
Bt63_08
Bt63_09
Bt63_10
Bt63_11
Bt63_12
Bt63_13
Bt63_14
H99_01
H99_02
H99_03
H99_04
H99_05
H99_06
H99_07
H99_08
H99_09
H99_10
H99_11
H99_12
H99_13
H99_14
Type of Repetitive Sequences
Simple repeat
Others
Copy-Number Variants
Deletion
Duplication
Features
Centromeres
MAT

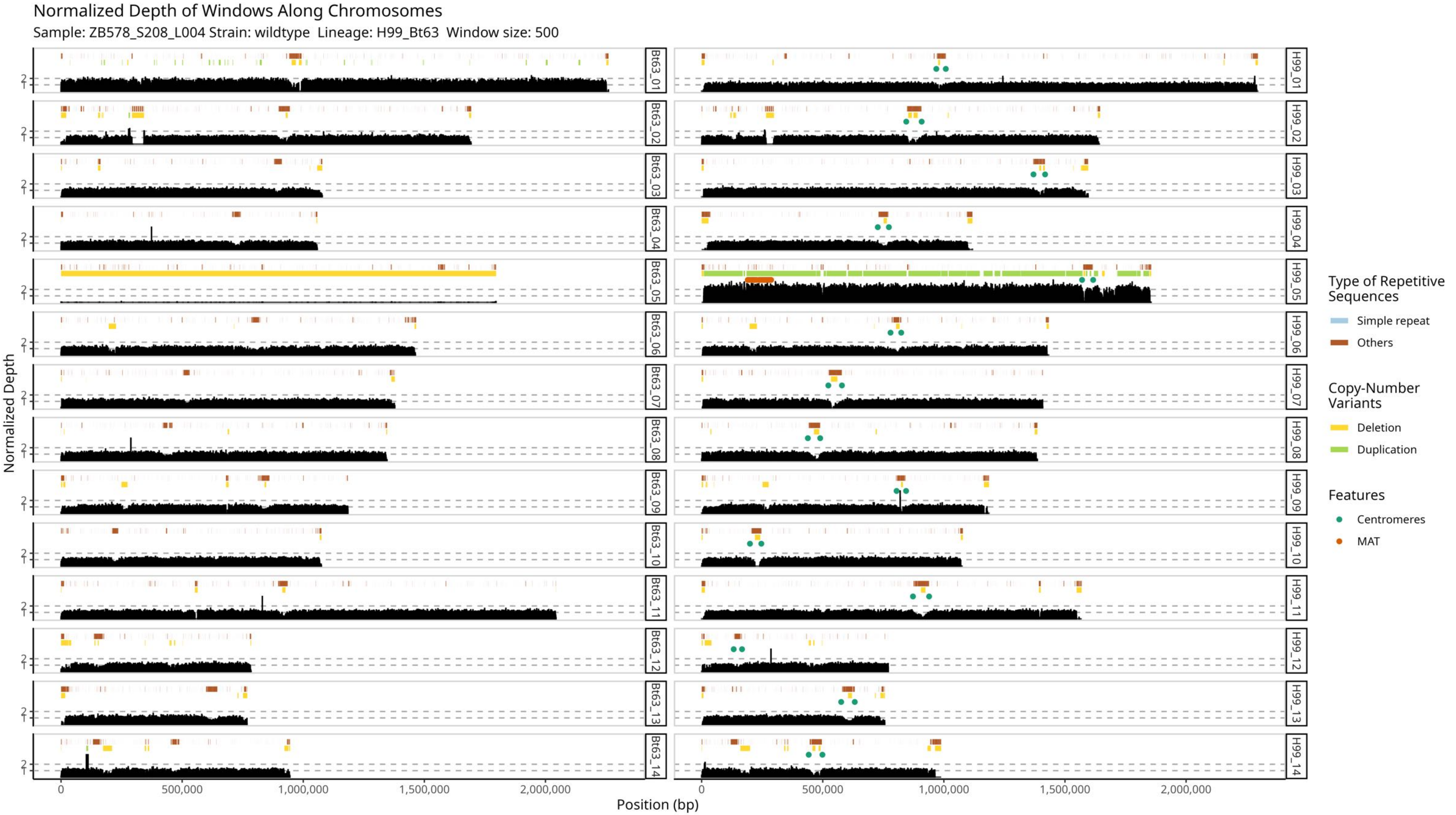

Normalized Depth of Windows Along Chromosomes
Sample: ZB578_S208_L004 Strain: wildtype Lineage: H99_Bt63 Window size: 500
Normalized Depth
Position (bp)
0
500,000
1,000,000
1,500,000
2,000,000
Bt63_01
Bt63_02
Bt63_03
Bt63_04
Bt63_05
Bt63_06
Bt63_07
Bt63_08
Bt63_09
Bt63_10
Bt63_11
Bt63_12
Bt63_13
Bt63_14
H99_01
H99_02
H99_03
H99_04
H99_05
H99_06
H99_07
H99_08
H99_09
H99_10
H99_11
H99_12
H99_13
H99_14
Type of Repetitive Sequences
Simple repeat
Others
Copy-Number Variants
Deletion
Duplication
Features
Centromeres
MAT

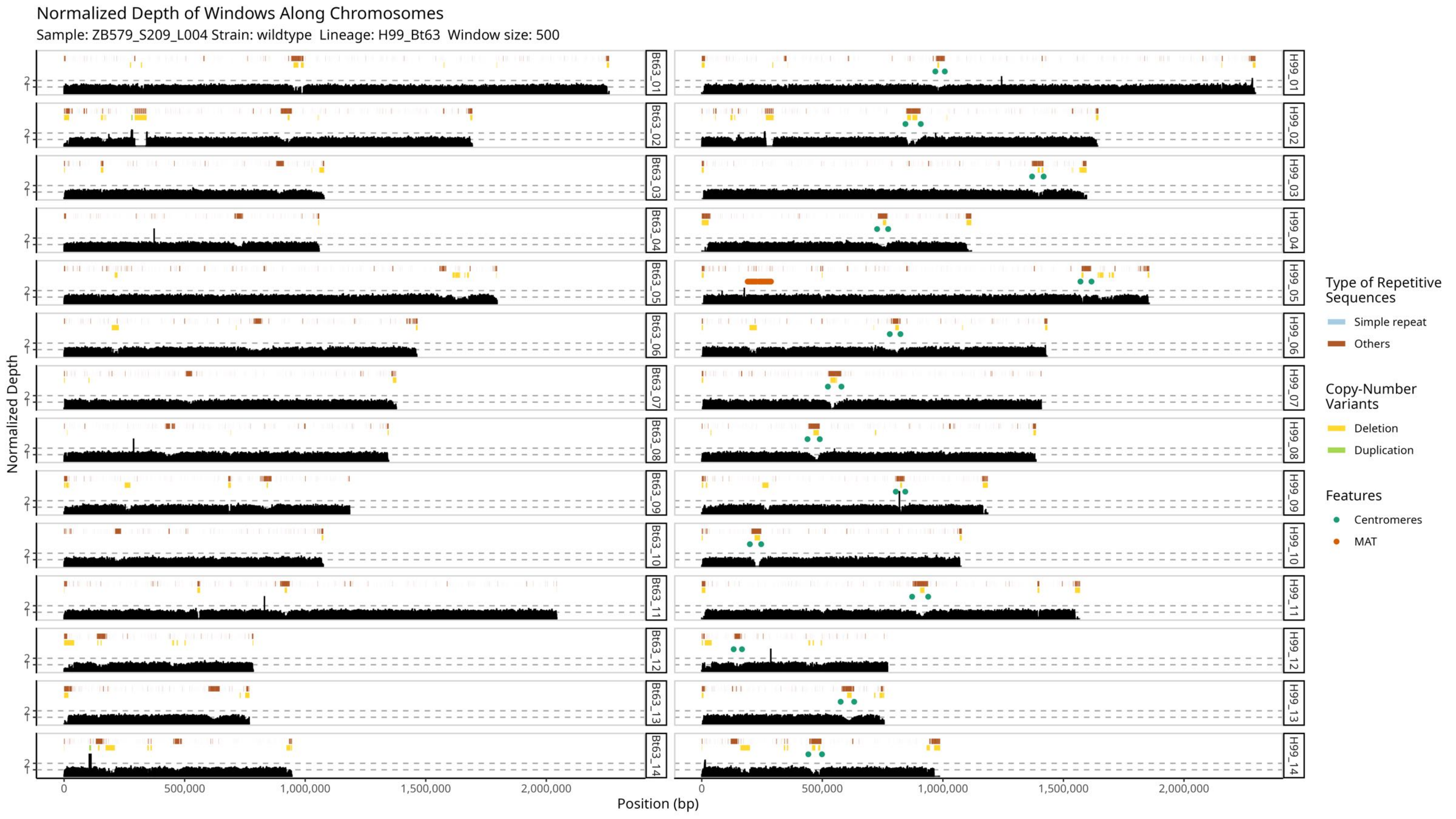

Normalized Depth of Windows Along Chromosomes
Sample: ZB579_S209_L004 Strain: wildtype Lineage: H99_Bt63 Window size: 500
Normalized Depth
Position (bp)
0
500,000
1,000,000
1,500,000
2,000,000
1
2
Bt63_01
Bt63_02
Bt63_03
Bt63_04
Bt63_05
Bt63_06
Bt63_07
Bt63_08
Bt63_09
Bt63_10
Bt63_11
Bt63_12
Bt63_13
Bt63_14
H99_01
H99_02
H99_03
H99_04
H99_05
H99_06
H99_07
H99_08
H99_09
H99_10
H99_11
H99_12
H99_13
H99_14
Type of Repetitive Sequences
Simple repeat
Others
Copy-Number Variants
Deletion
Duplication
Features
Centromeres
MAT

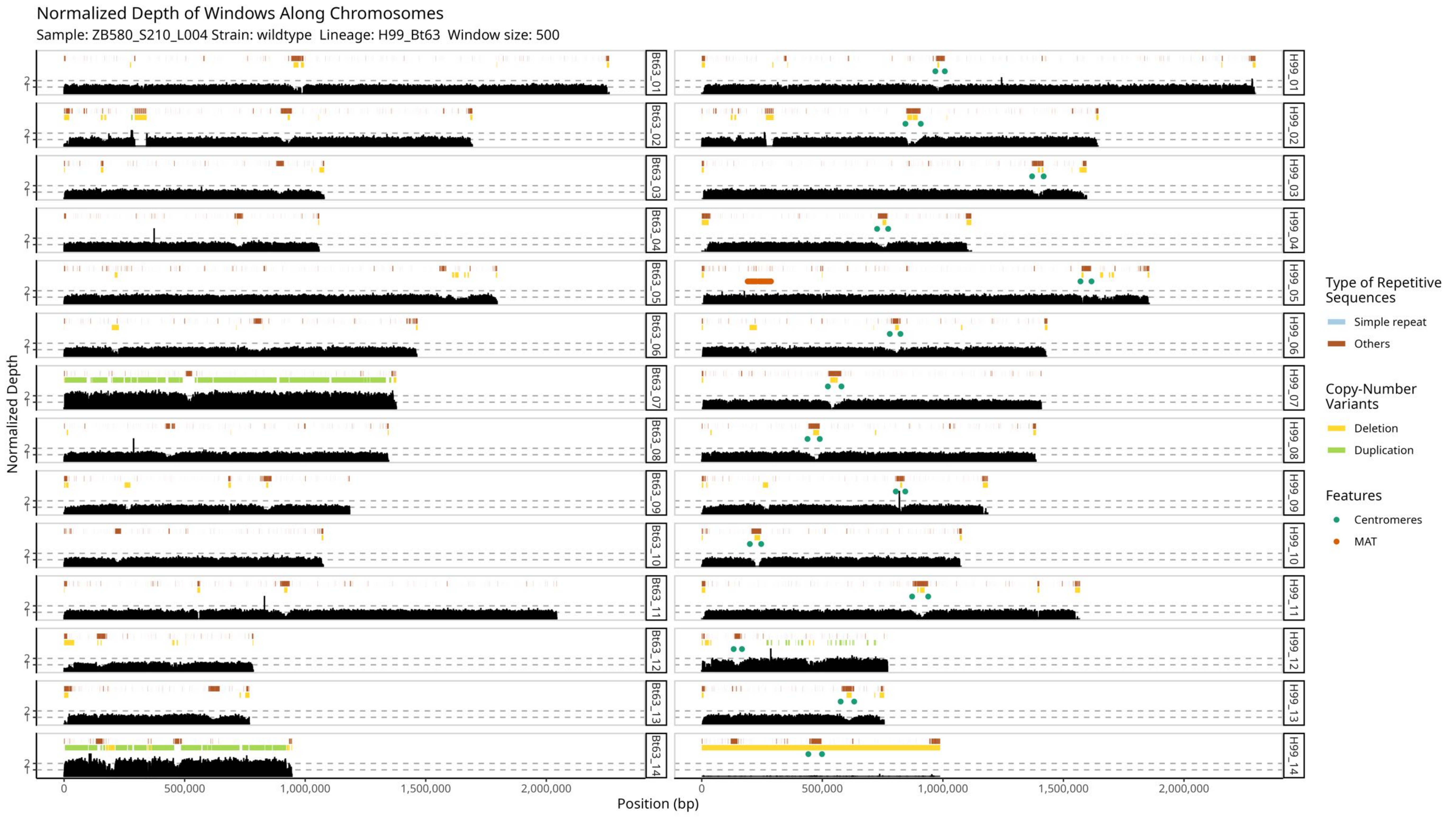

Normalized Depth of Windows Along Chromosomes
Sample: ZB580_S210_L004 Strain: wildtype Lineage: H99_Bt63 Window size: 500
Normalized Depth
Position (bp)
0
500,000
1,000,000
1,500,000
2,000,000
Bt63_01
Bt63_02
Bt63_03
Bt63_04
Bt63_05
Bt63_06
Bt63_07
Bt63_08
Bt63_09
Bt63_10
Bt63_11
Bt63_12
Bt63_13
Bt63_14
H99_01
H99_02
H99_03
H99_04
H99_05
H99_06
H99_07
H99_08
H99_09
H99_10
H99_11
H99_12
H99_13
H99_14
Type of Repetitive Sequences
Simple repeat
Others
Copy-Number Variants
Deletion
Duplication
Features
Centromeres
MAT
