## Supplemental Figure 6 for "Meiosis-specific genes play roles in ploidy reduction in *Cryptococcus neoformans* titan cells"

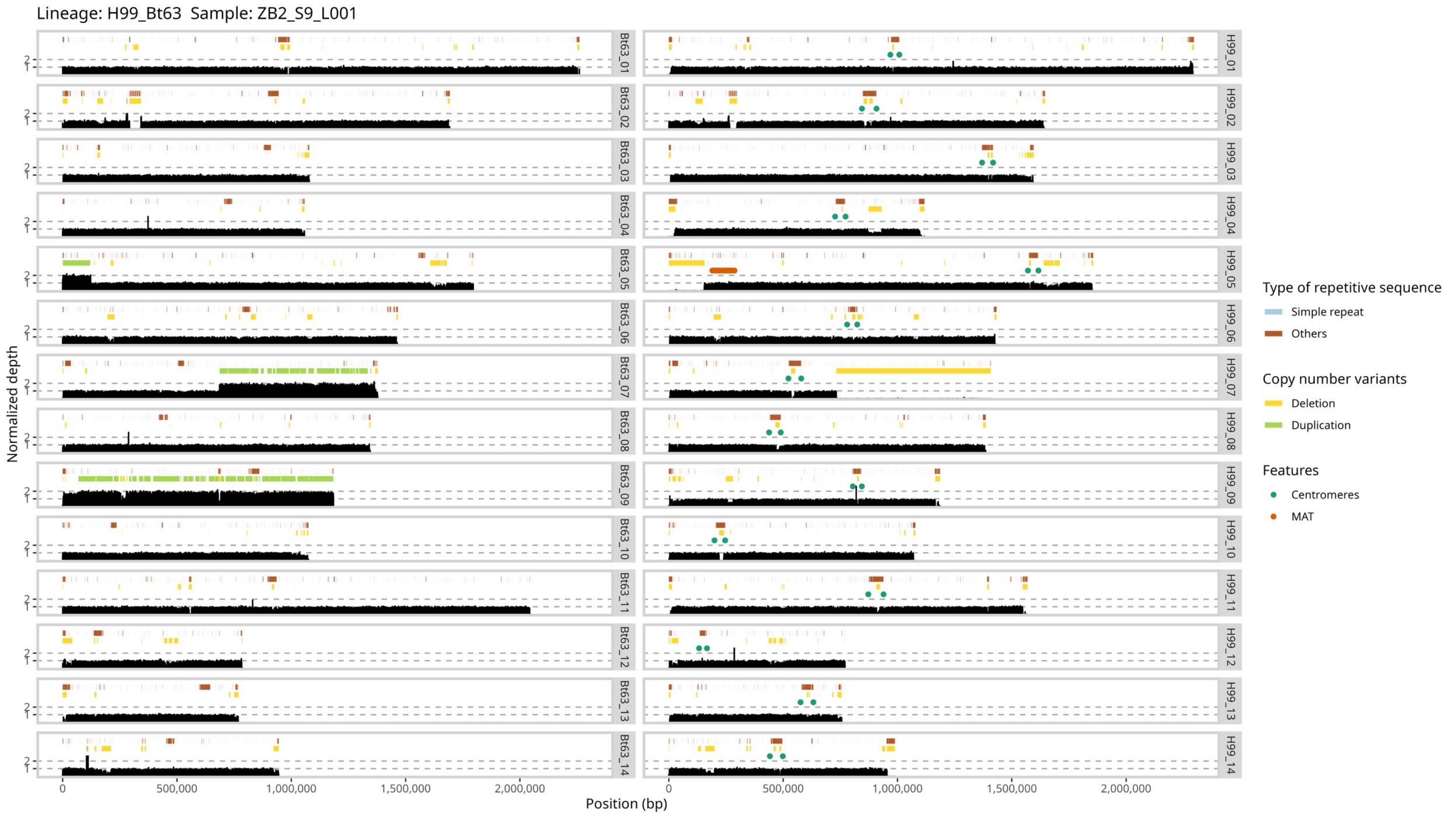
Lineage: H99_Bt63 Sample: ZB2_S9_L001
Normalized depth
Position (bp)
Bt63_01
Bt63_02
Bt63_03
Bt63_04
Bt63_05
Bt63_06
Bt63_07
Bt63_08
Bt63_09
Bt63_10
Bt63_11
Bt63_12
Bt63_13
Bt63_14
H99_01
H99_02
H99_03
H99_04
H99_05
H99_06
H99_07
H99_08
H99_09
H99_10
H99_11
H99_12
H99_13
H99_14
0
500,000
1,000,000
1,500,000
2,000,000
Type of repetitive sequence
Simple repeat
Others
Copy number variants
Deletion
Duplication
Features
Centromeres
MAT

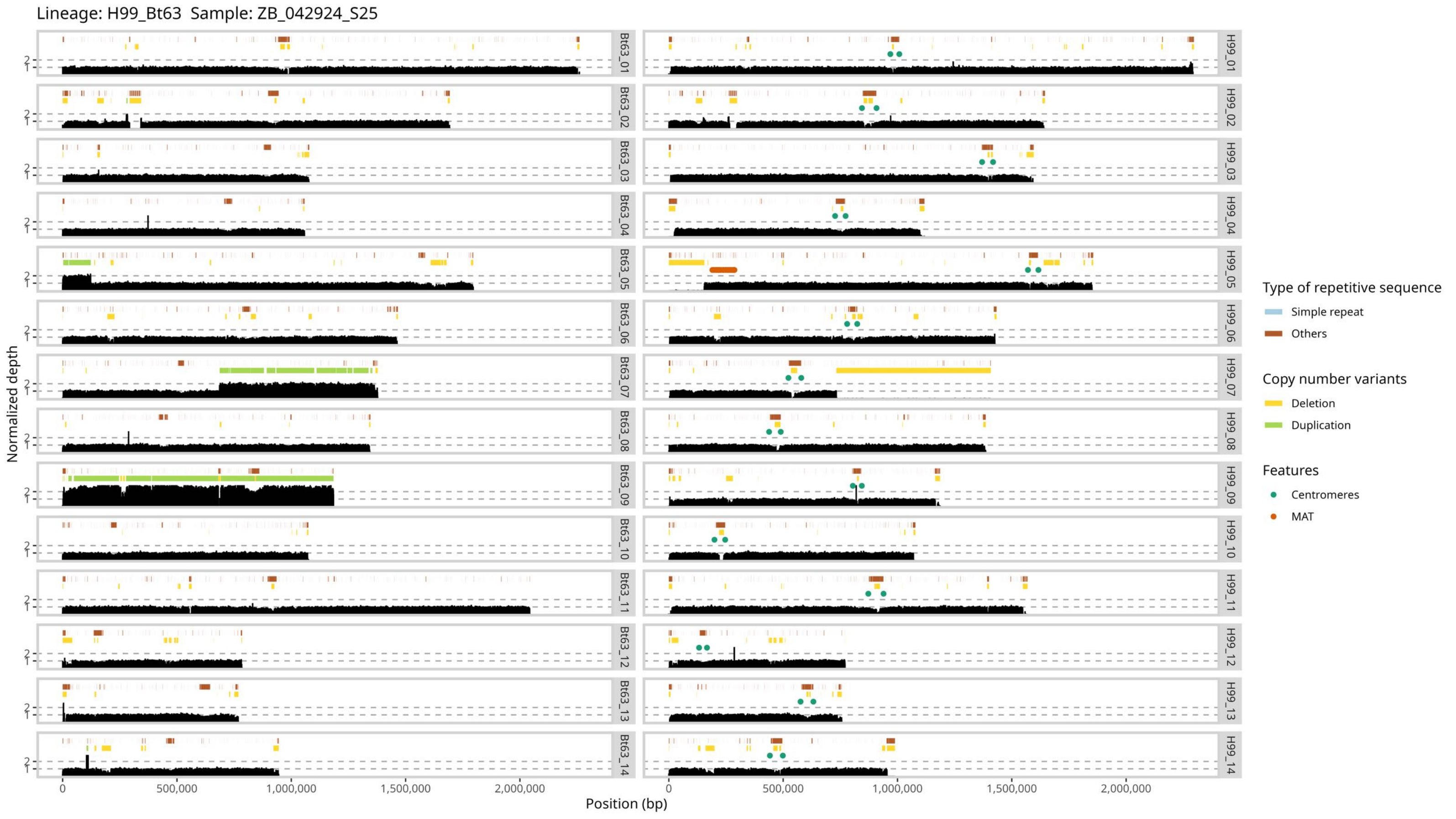
Lineage: H99_Bt63 Sample: ZB_042924_S25
Normalized depth
Position (bp)
0
500,000
1,000,000
1,500,000
2,000,000
Bt63_01
Bt63_02
Bt63_03
Bt63_04
Bt63_05
Bt63_06
Bt63_07
Bt63_08
Bt63_09
Bt63_10
Bt63_11
Bt63_12
Bt63_13
Bt63_14
H99_01
H99_02
H99_03
H99_04
H99_05
H99_06
H99_07
H99_08
H99_09
H99_10
H99_11
H99_12
H99_13
H99_14
Type of repetitive sequence
Simple repeat
Others
Copy number variants
Deletion
Duplication
Features
Centromeres
MAT

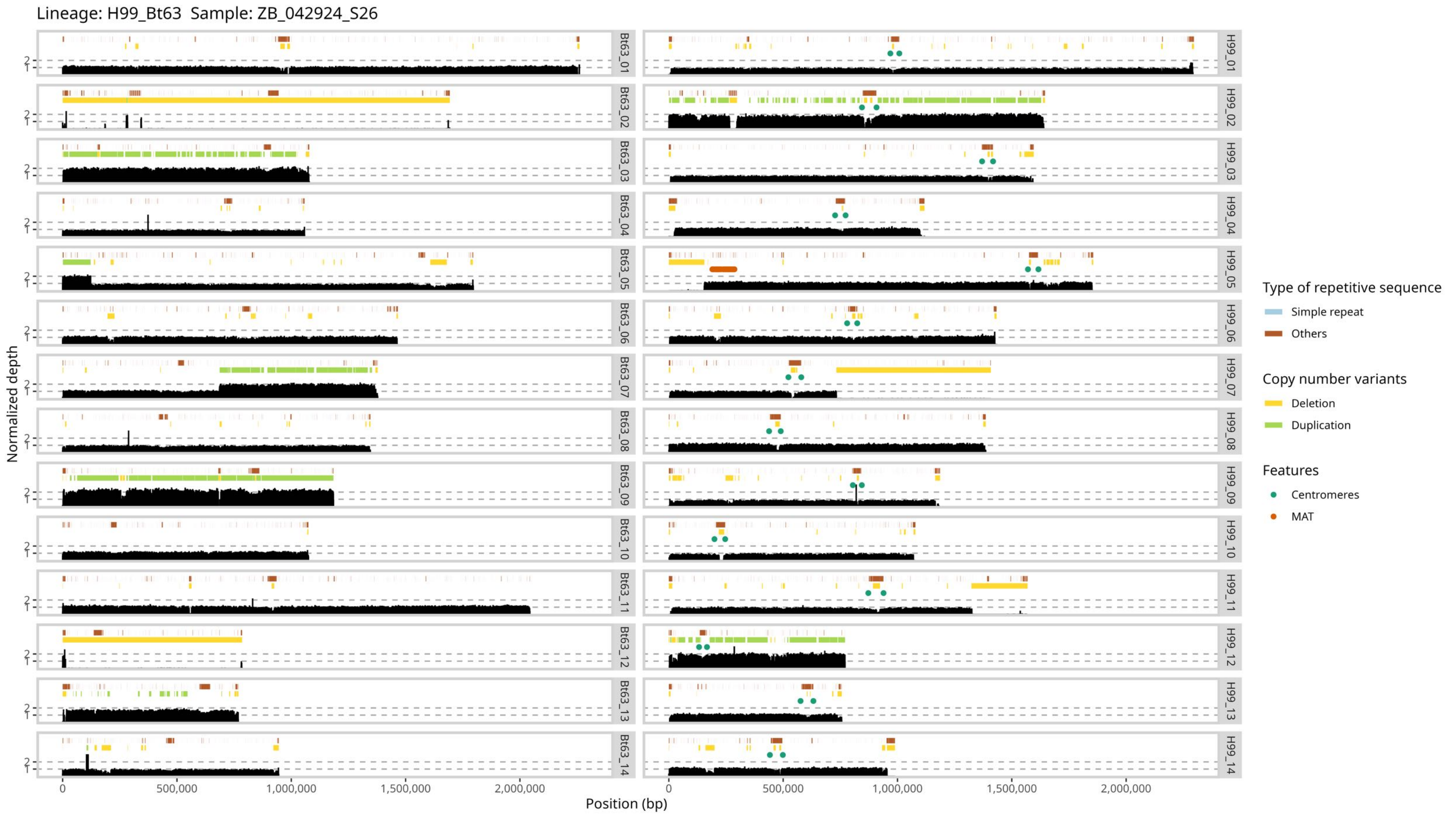

Lineage: H99_Bt63 Sample: ZB_042924_S26
Bt63_01
Bt63_02
Bt63_03
Bt63_04
Bt63_05
Bt63_06
Bt63_07
Bt63_08
Bt63_09
Bt63_10
Bt63_11
Bt63_12
Bt63_13
Bt63_14
H99_01
H99_02
H99_03
H99_04
H99_05
H99_06
H99_07
H99_08
H99_09
H99_10
H99_11
H99_12
H99_13
H99_14
Normalized depth
Position (bp)
0
500,000
1,000,000
1,500,000
2,000,000
Type of repetitive sequence
Simple repeat
Others
Copy number variants
Deletion
Duplication
Features
Centromeres
MAT

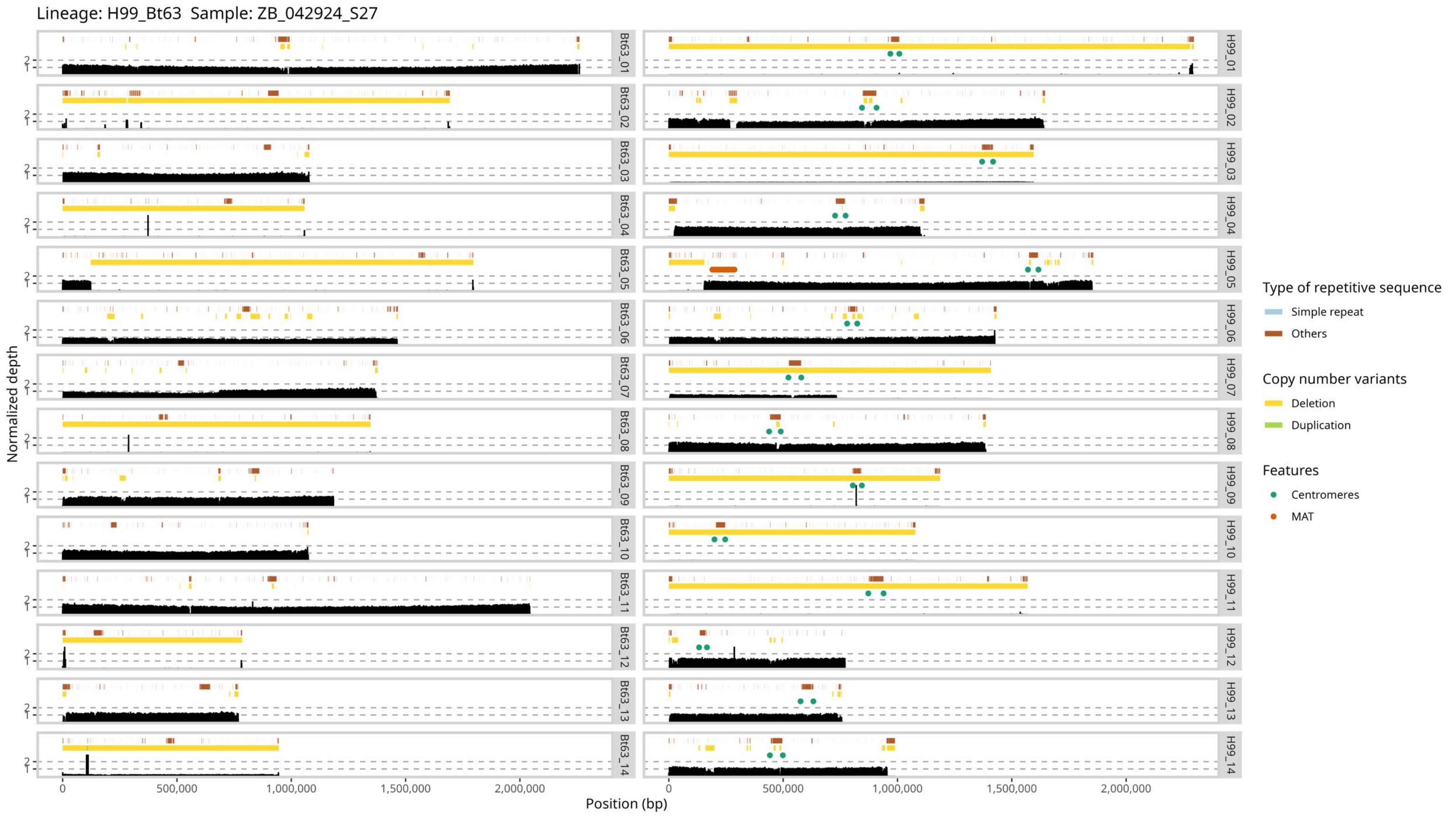

Lineage: H99_Bt63 Sample: ZB_042924_S27
Normalized depth
Position (bp)
0
500,000
1,000,000
1,500,000
2,000,000
Bt63_01
Bt63_02
Bt63_03
Bt63_04
Bt63_05
Bt63_06
Bt63_07
Bt63_08
Bt63_09
Bt63_10
Bt63_11
Bt63_12
Bt63_13
Bt63_14
H99_01
H99_02
H99_03
H99_04
H99_05
H99_06
H99_07
H99_08
H99_09
H99_10
H99_11
H99_12
H99_13
H99_14
Type of repetitive sequence
Simple repeat
Others
Copy number variants
Deletion
Duplication
Features
Centromeres
MAT

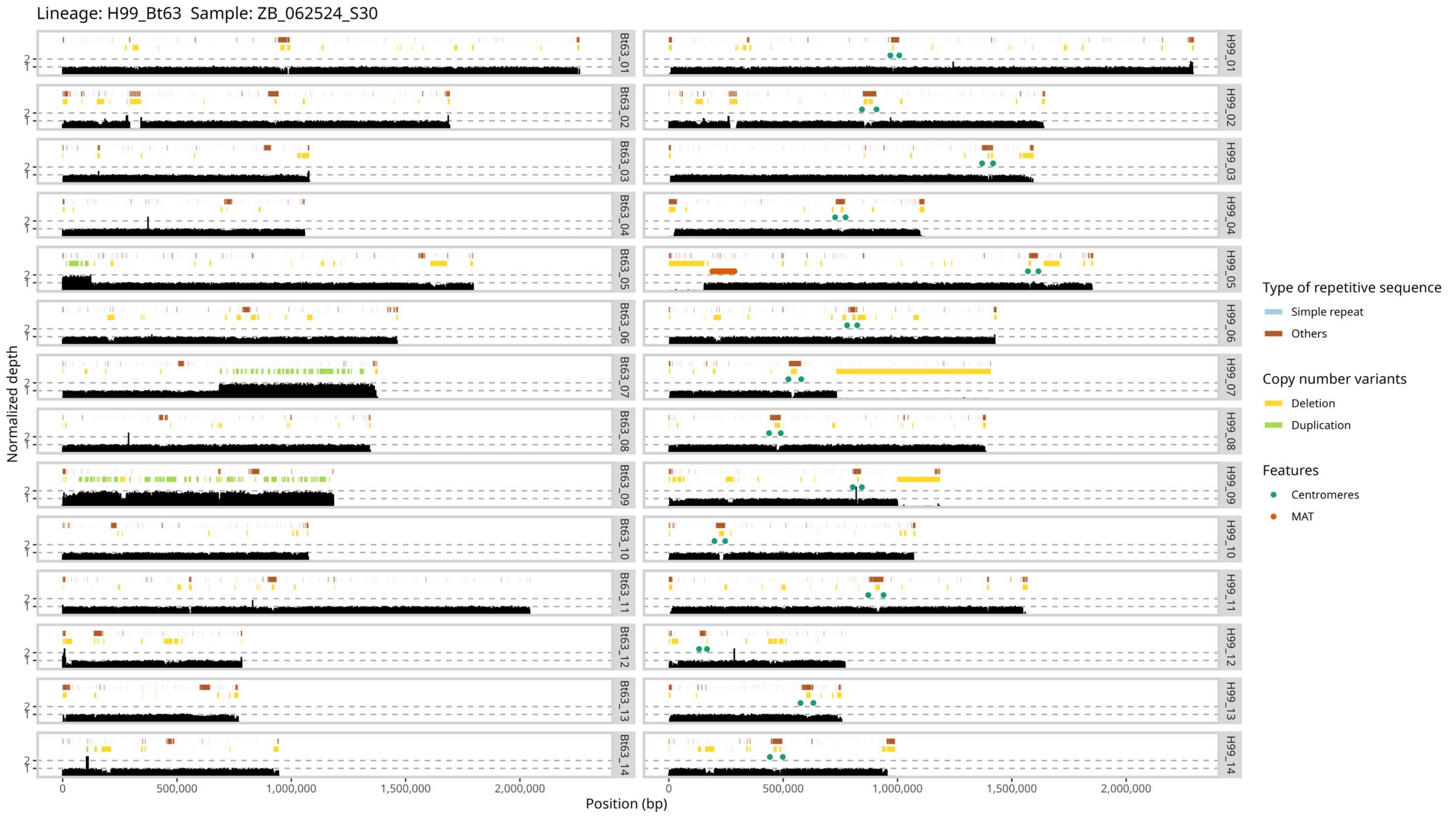
Lineage: H99_Bt63 Sample: ZB_062524_S30
Normalized depth
Position (bp)
0
500,000
1,000,000
1,500,000
2,000,000
Bt63_01
Bt63_02
Bt63_03
Bt63_04
Bt63_05
Bt63_06
Bt63_07
Bt63_08
Bt63_09
Bt63_10
Bt63_11
Bt63_12
Bt63_13
Bt63_14
H99_01
H99_02
H99_03
H99_04
H99_05
H99_06
H99_07
H99_08
H99_09
H99_10
H99_11
H99_12
H99_13
H99_14
Type of repetitive sequence
Simple repeat
Others
Copy number variants
Deletion
Duplication
Features
Centromeres
MAT

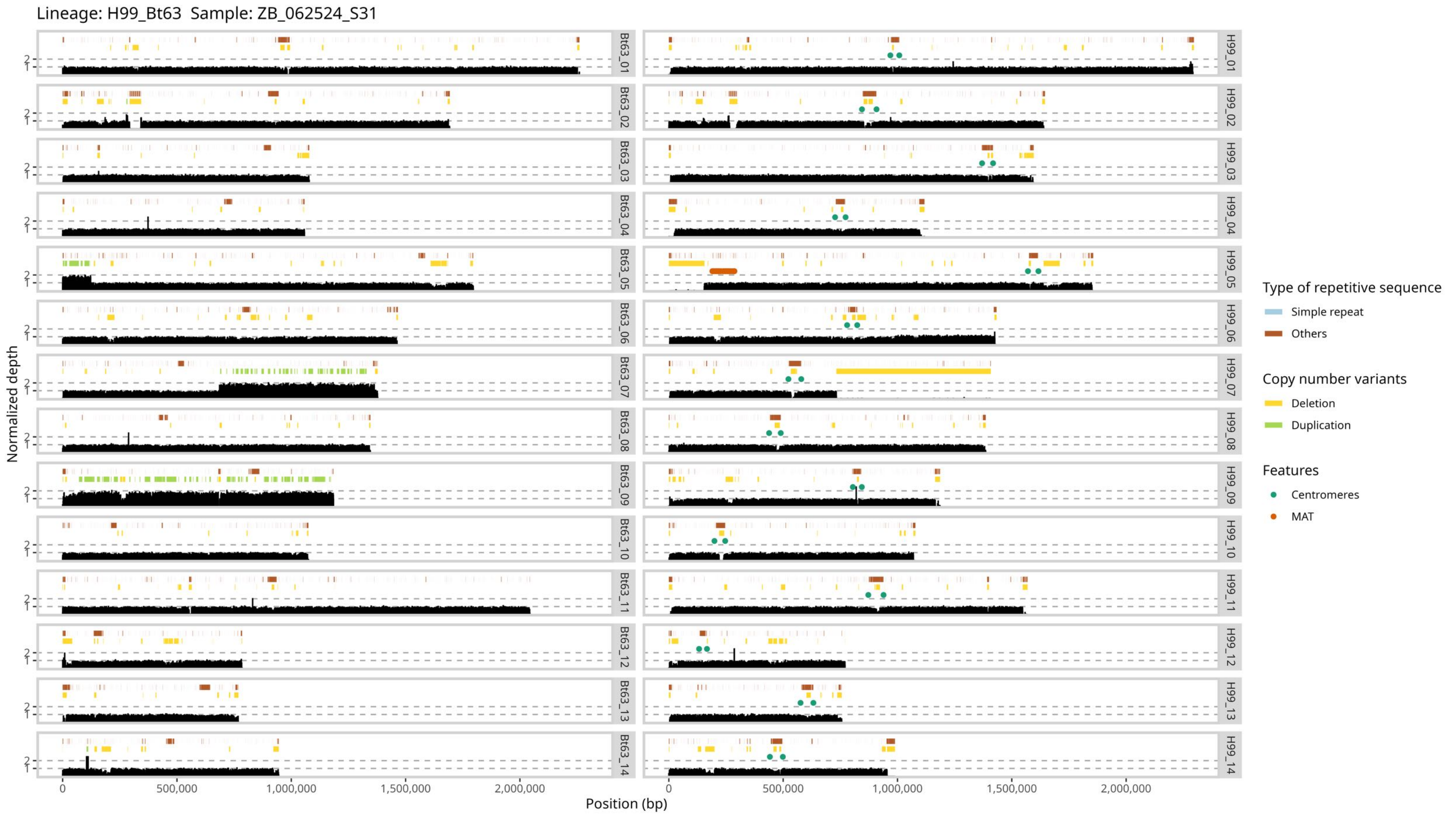

Lineage: H99_Bt63 Sample: ZB_062524_S31
Normalized depth
Position (bp)
0
500,000
1,000,000
1,500,000
2,000,000
Bt63_01
Bt63_02
Bt63_03
Bt63_04
Bt63_05
Bt63_06
Bt63_07
Bt63_08
Bt63_09
Bt63_10
Bt63_11
Bt63_12
Bt63_13
Bt63_14
H99_01
H99_02
H99_03
H99_04
H99_05
H99_06
H99_07
H99_08
H99_09
H99_10
H99_11
H99_12
H99_13
H99_14
Type of repetitive sequence
Simple repeat
Others
Copy number variants
Deletion
Duplication
Features
Centromeres
MAT

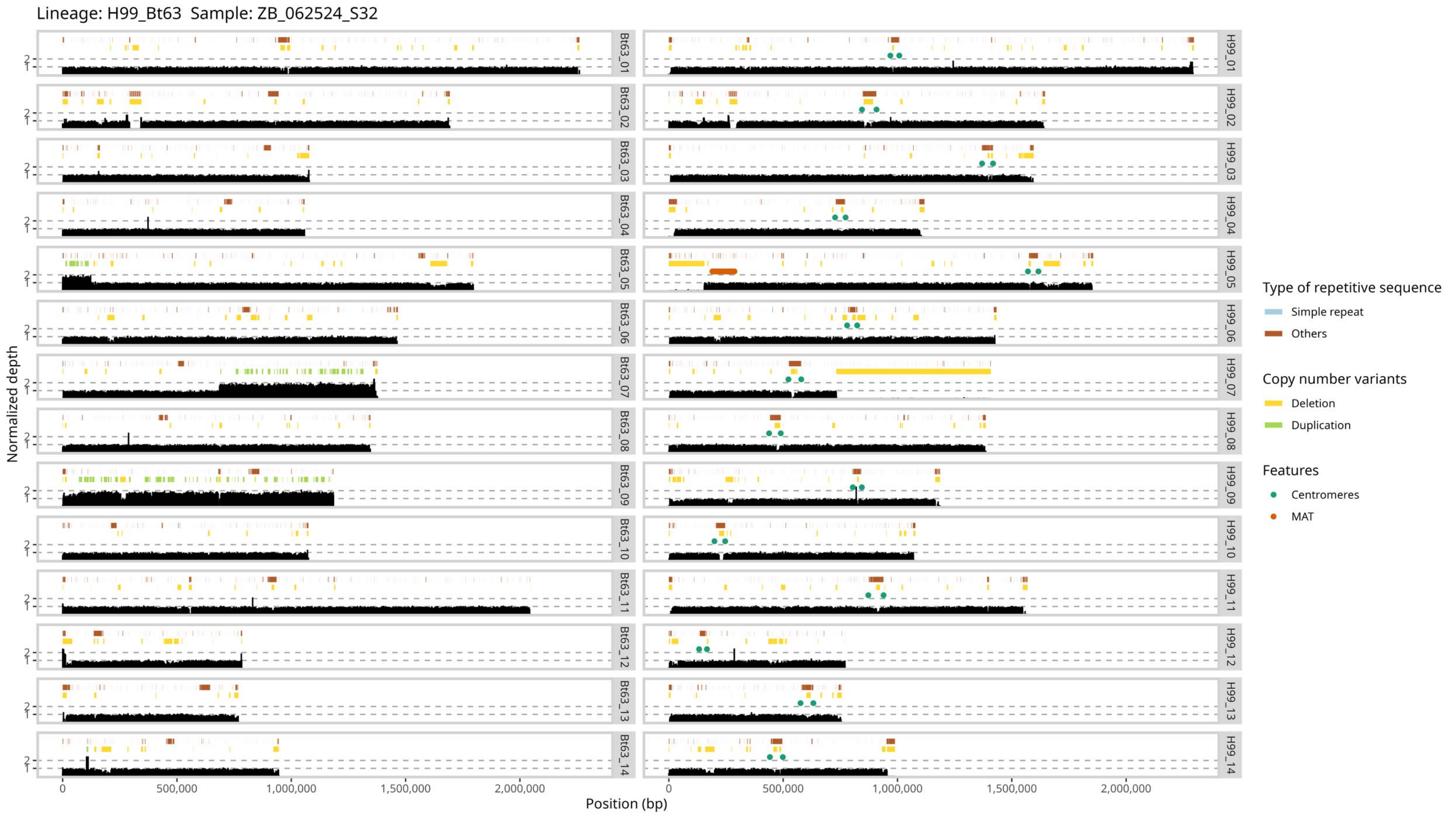

Lineage: H99_Bt63 Sample: ZB_062524_S32
Normalized depth
Position (bp)
Bt63_01
Bt63_02
Bt63_03
Bt63_04
Bt63_05
Bt63_06
Bt63_07
Bt63_08
Bt63_09
Bt63_10
Bt63_11
Bt63_12
Bt63_13
Bt63_14
H99_01
H99_02
H99_03
H99_04
H99_05
H99_06
H99_07
H99_08
H99_09
H99_10
H99_11
H99_12
H99_13
H99_14
0
500,000
1,000,000
1,500,000
2,000,000
Type of repetitive sequence
Simple repeat
Others
Copy number variants
Deletion
Duplication
Features
Centromeres
MAT

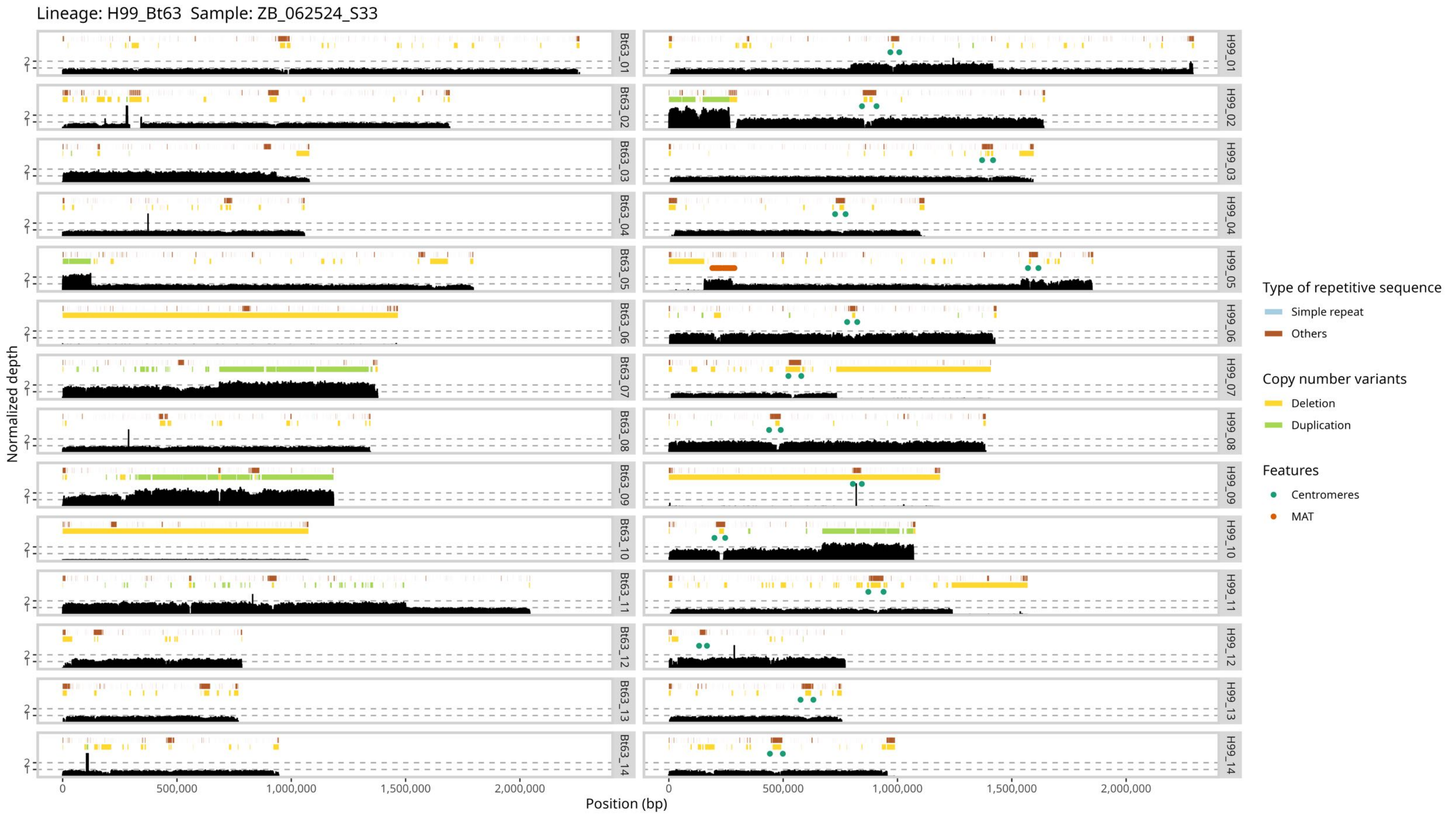

Lineage: H99_Bt63 Sample: ZB_062524_S33
Normalized depth
Position (bp)
0
500,000
1,000,000
1,500,000
2,000,000
1
2
Bt63_01
Bt63_02
Bt63_03
Bt63_04
Bt63_05
Bt63_06
Bt63_07
Bt63_08
Bt63_09
Bt63_10
Bt63_11
Bt63_12
Bt63_13
Bt63_14
H99_01
H99_02
H99_03
H99_04
H99_05
H99_06
H99_07
H99_08
H99_09
H99_10
H99_11
H99_12
H99_13
H99_14
Type of repetitive sequence
Simple repeat
Others
Copy number variants
Deletion
Duplication
Features
Centromeres
MAT

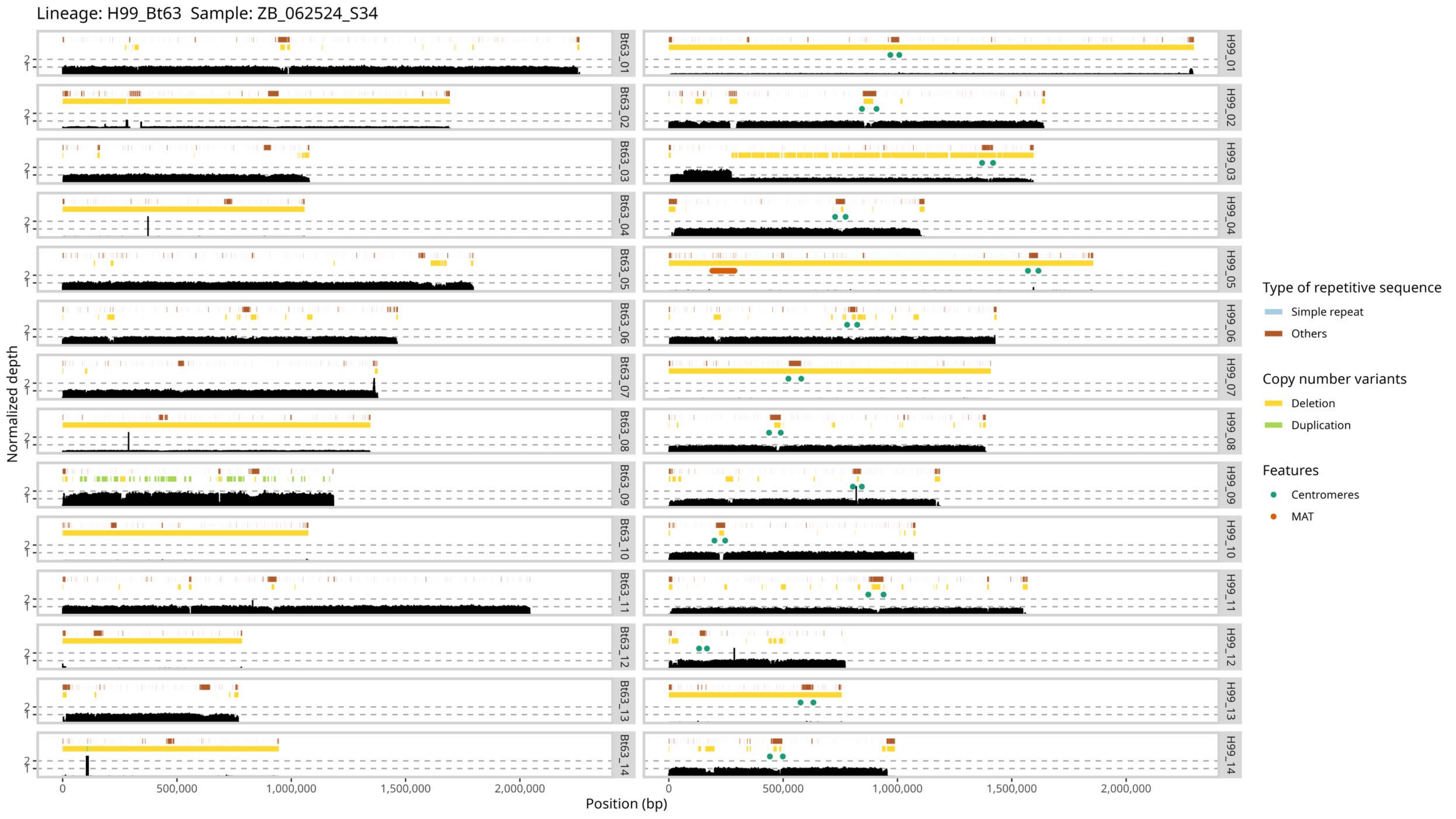

Lineage: H99_Bt63 Sample: ZB_062524_S34
Normalized depth
Position (bp)
0
500,000
1,000,000
1,500,000
2,000,000
Bt63_01
Bt63_02
Bt63_03
Bt63_04
Bt63_05
Bt63_06
Bt63_07
Bt63_08
Bt63_09
Bt63_10
Bt63_11
Bt63_12
Bt63_13
Bt63_14
H99_01
H99_02
H99_03
H99_04
H99_05
H99_06
H99_07
H99_08
H99_09
H99_10
H99_11
H99_12
H99_13
H99_14
Type of repetitive sequence
Simple repeat
Others
Copy number variants
Deletion
Duplication
Features
Centromeres
MAT

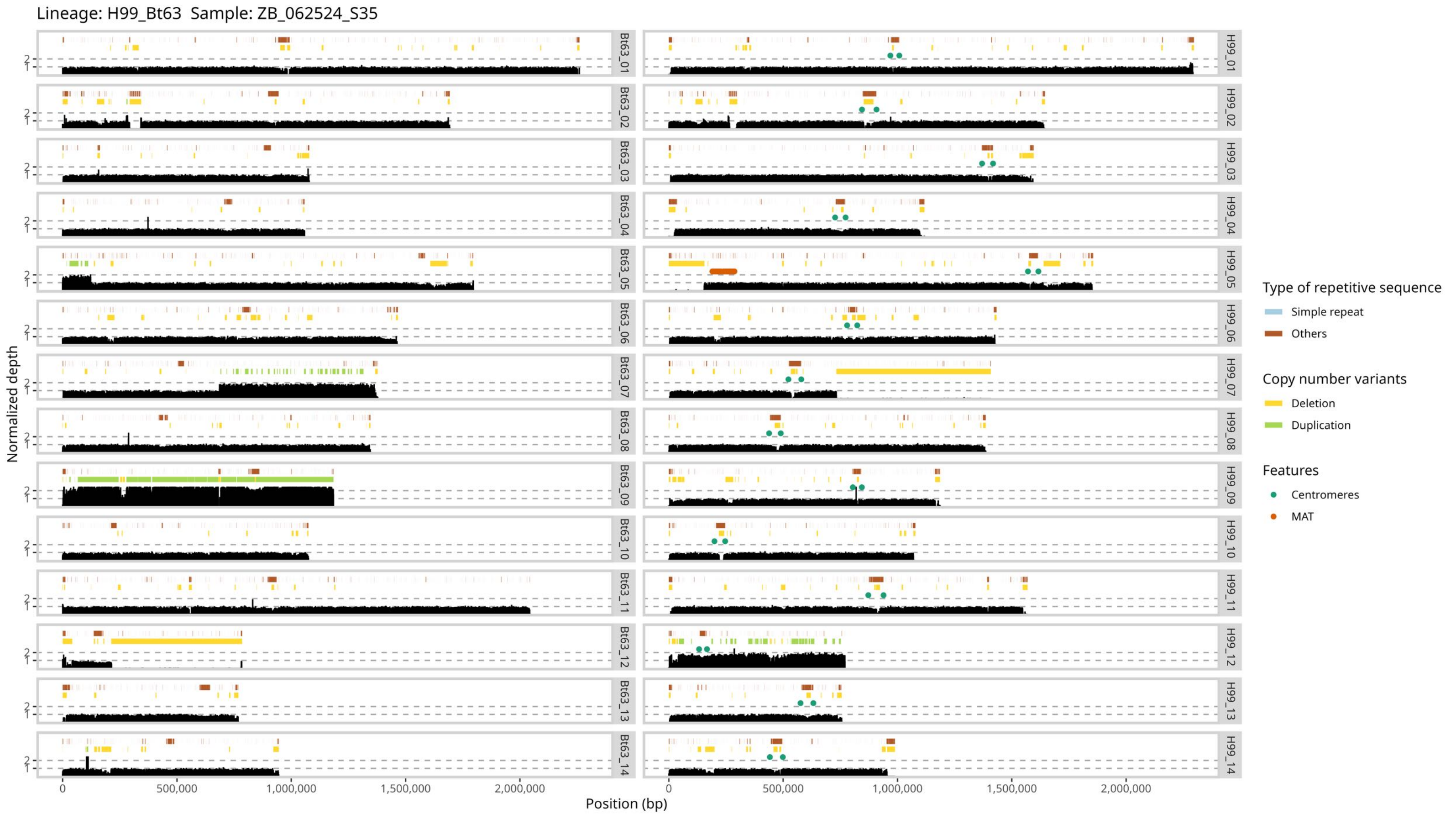
Lineage: H99_Bt63 Sample: ZB_062524_S35
Normalized depth
Position (bp)
0
500,000
1,000,000
1,500,000
2,000,000
Bt63_01
Bt63_02
Bt63_03
Bt63_04
Bt63_05
Bt63_06
Bt63_07
Bt63_08
Bt63_09
Bt63_10
Bt63_11
Bt63_12
Bt63_13
Bt63_14
H99_01
H99_02
H99_03
H99_04
H99_05
H99_06
H99_07
H99_08
H99_09
H99_10
H99_11
H99_12
H99_13
H99_14
Type of repetitive sequence
Simple repeat
Others
Copy number variants
Deletion
Duplication
Features
Centromeres
MAT

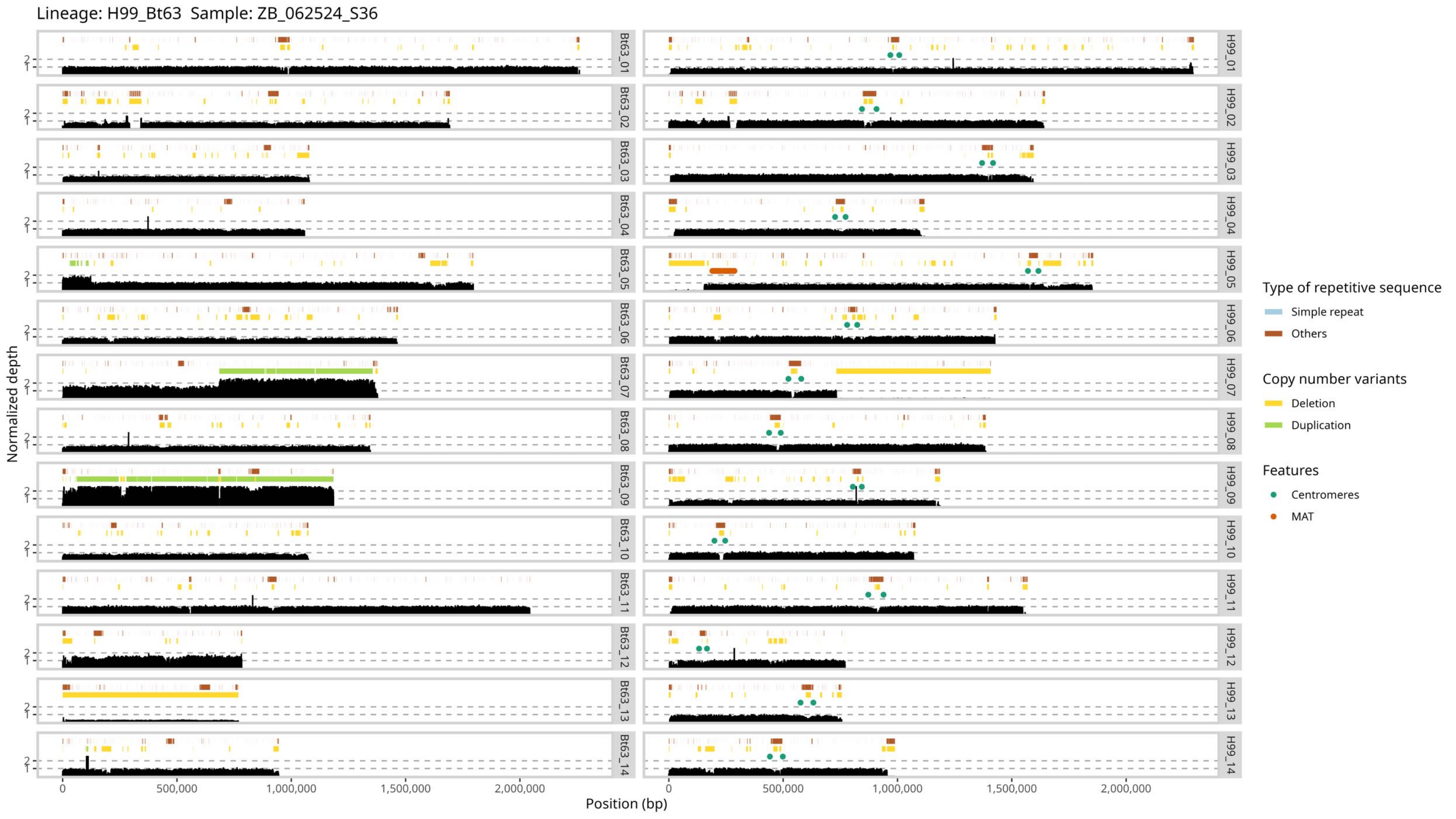

Lineage: H99_Bt63 Sample: ZB_062524_S36
Normalized depth
Position (bp)
0
500,000
1,000,000
1,500,000
2,000,000
Bt63_01
Bt63_02
Bt63_03
Bt63_04
Bt63_05
Bt63_06
Bt63_07
Bt63_08
Bt63_09
Bt63_10
Bt63_11
Bt63_12
Bt63_13
Bt63_14
H99_01
H99_02
H99_03
H99_04
H99_05
H99_06
H99_07
H99_08
H99_09
H99_10
H99_11
H99_12
H99_13
H99_14
Type of repetitive sequence
Simple repeat
Others
Copy number variants
Deletion
Duplication
Features
Centromeres
MAT

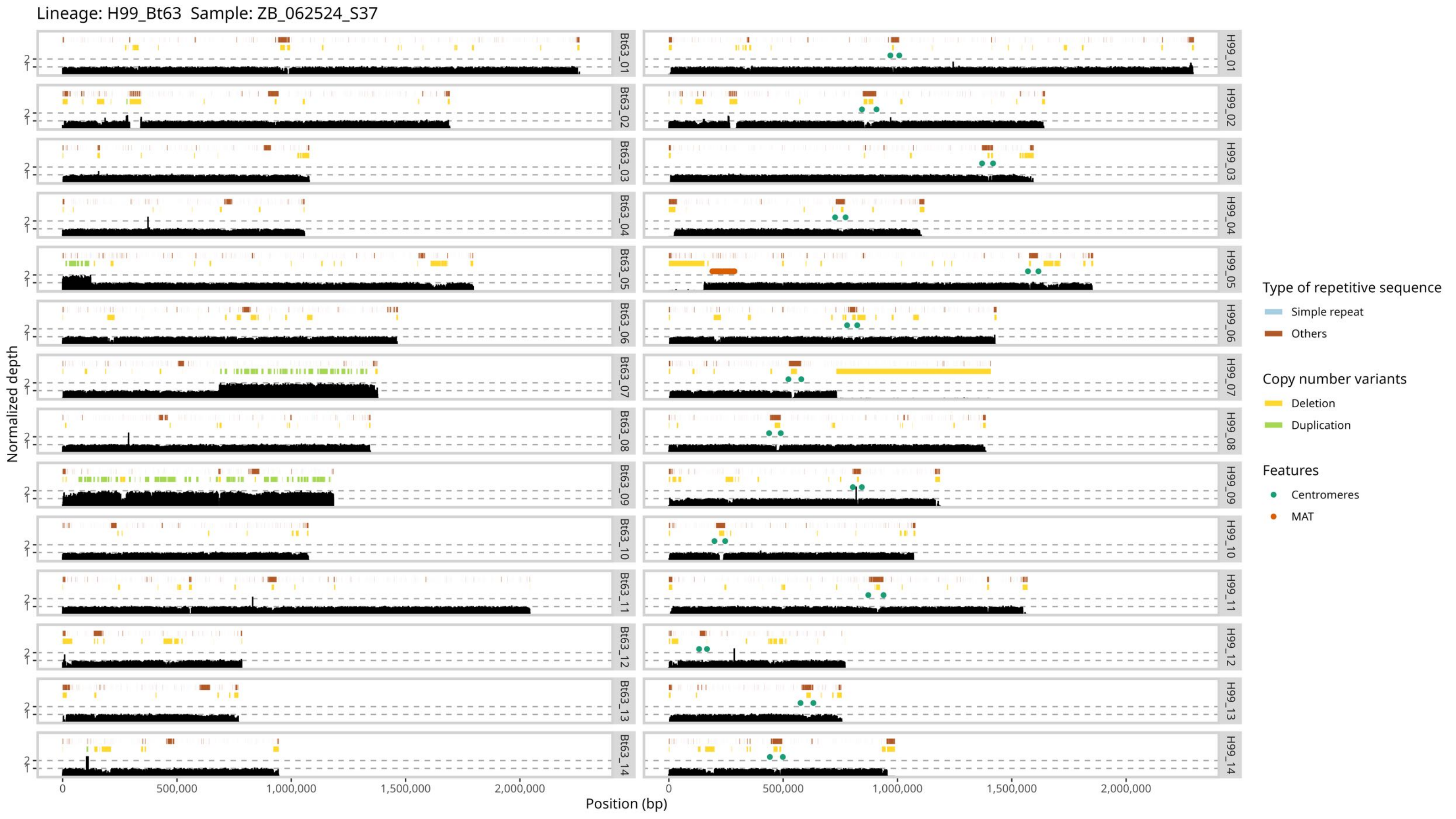

Lineage: H99_Bt63 Sample: ZB_062524_S37
Bt63_01
Bt63_02
Bt63_03
Bt63_04
Bt63_05
Bt63_06
Bt63_07
Bt63_08
Bt63_09
Bt63_10
Bt63_11
Bt63_12
Bt63_13
Bt63_14
H99_01
H99_02
H99_03
H99_04
H99_05
H99_06
H99_07
H99_08
H99_09
H99_10
H99_11
H99_12
H99_13
H99_14
Normalized depth
Position (bp)
0
500,000
1,000,000
1,500,000
2,000,000
Type of repetitive sequence
Simple repeat
Others
Copy number variants
Deletion
Duplication
Features
Centromeres
MAT

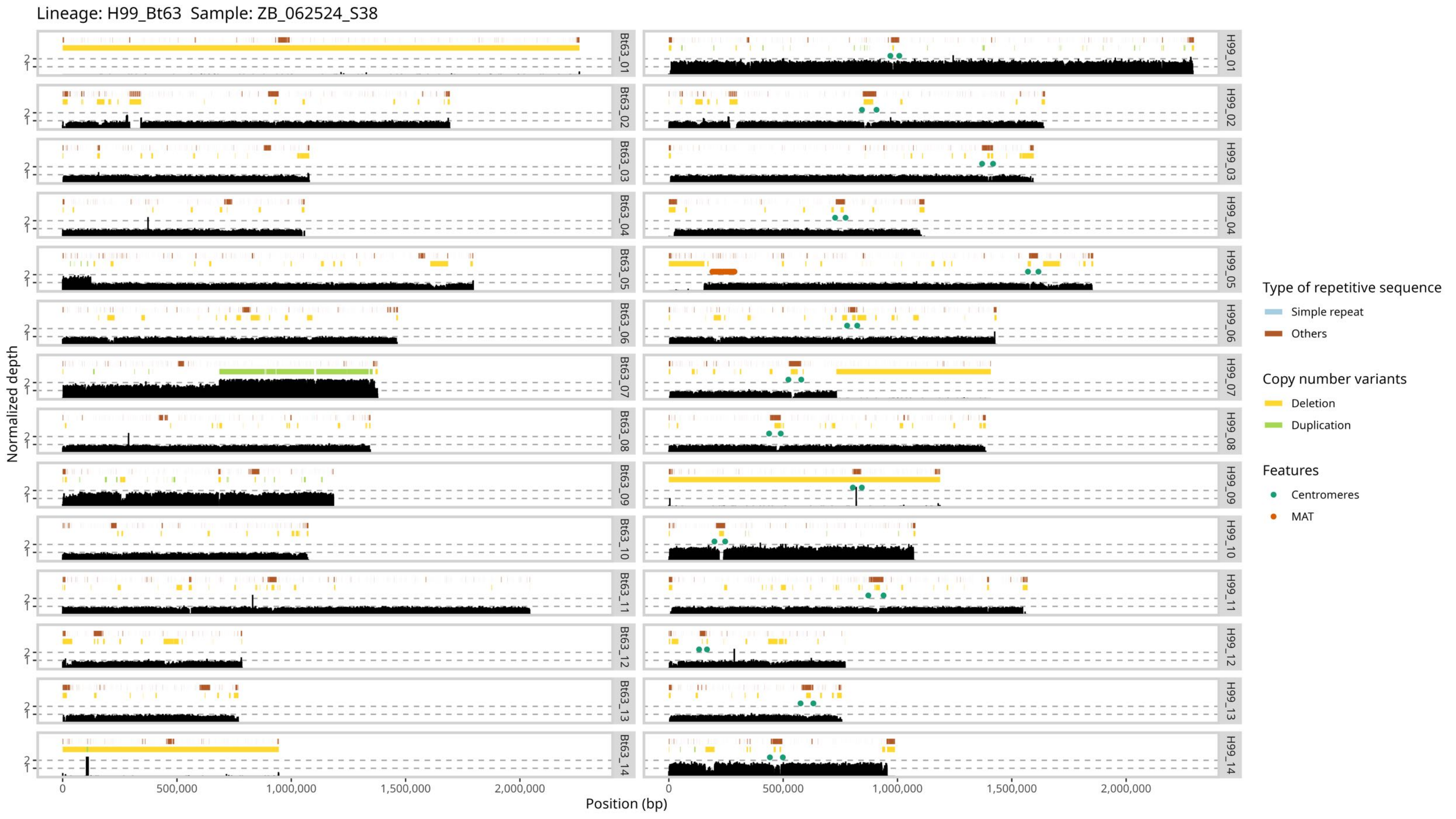

Lineage: H99_Bt63 Sample: ZB_062524_S38
Normalized depth
Position (bp)
Bt63_01
Bt63_02
Bt63_03
Bt63_04
Bt63_05
Bt63_06
Bt63_07
Bt63_08
Bt63_09
Bt63_10
Bt63_11
Bt63_12
Bt63_13
Bt63_14
H99_01
H99_02
H99_03
H99_04
H99_05
H99_06
H99_07
H99_08
H99_09
H99_10
H99_11
H99_12
H99_13
H99_14
0
500,000
1,000,000
1,500,000
2,000,000
Type of repetitive sequence
Simple repeat
Others
Copy number variants
Deletion
Duplication
Features
Centromeres
MAT
