## Supplemental Figure 7 for "Meiosis-specific genes play roles in ploidy reduction in *Cryptococcus neoformans* titan cells"

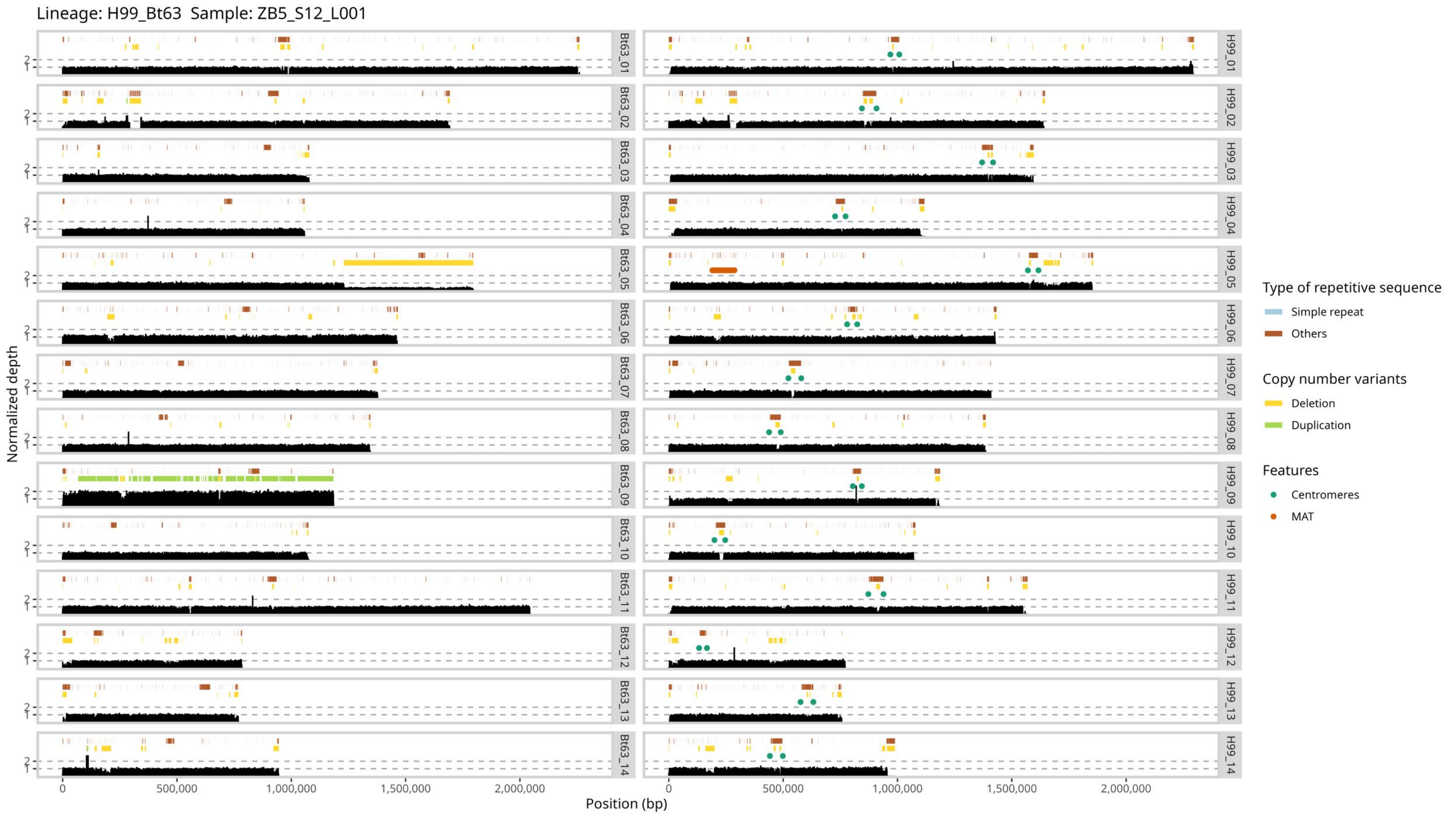

Lineage: H99_Bt63 Sample: ZB5_S12_L001
Bt63_01
Bt63_02
Bt63_03
Bt63_04
Bt63_05
Bt63_06
Bt63_07
Bt63_08
Bt63_09
Bt63_10
Bt63_11
Bt63_12
Bt63_13
Bt63_14
H99_01
H99_02
H99_03
H99_04
H99_05
H99_06
H99_07
H99_08
H99_09
H99_10
H99_11
H99_12
H99_13
H99_14
Normalized depth
Position (bp)
0
500,000
1,000,000
1,500,000
2,000,000
Type of repetitive sequence
Simple repeat
Others
Copy number variants
Deletion
Duplication
Features
Centromeres
MAT

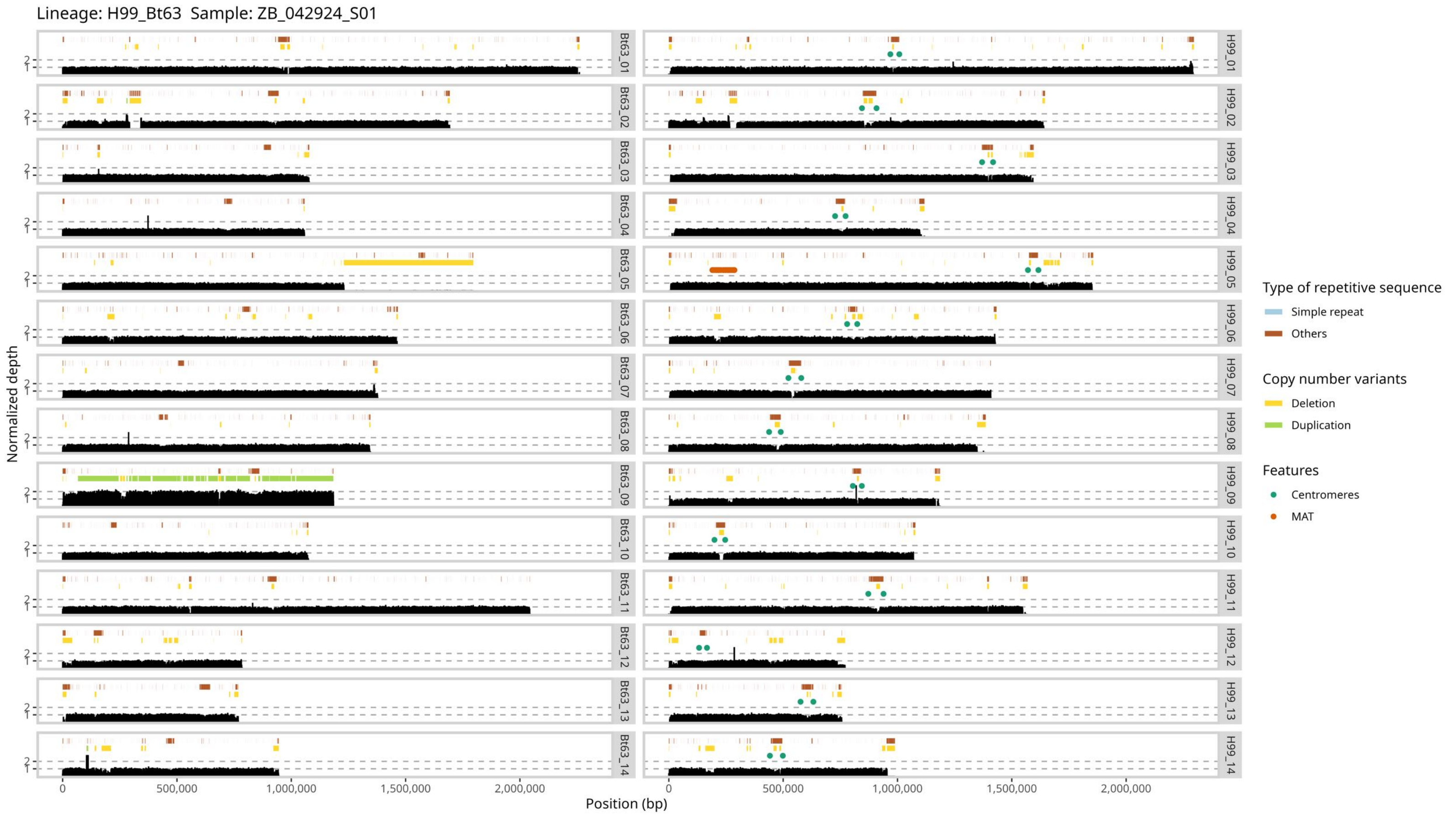

Lineage: H99_Bt63 Sample: ZB_042924_S01
Normalized depth
Position (bp)
0
500,000
1,000,000
1,500,000
2,000,000
Bt63_01
Bt63_02
Bt63_03
Bt63_04
Bt63_05
Bt63_06
Bt63_07
Bt63_08
Bt63_09
Bt63_10
Bt63_11
Bt63_12
Bt63_13
Bt63_14
H99_01
H99_02
H99_03
H99_04
H99_05
H99_06
H99_07
H99_08
H99_09
H99_10
H99_11
H99_12
H99_13
H99_14
Type of repetitive sequence
Simple repeat
Others
Copy number variants
Deletion
Duplication
Features
Centromeres
MAT

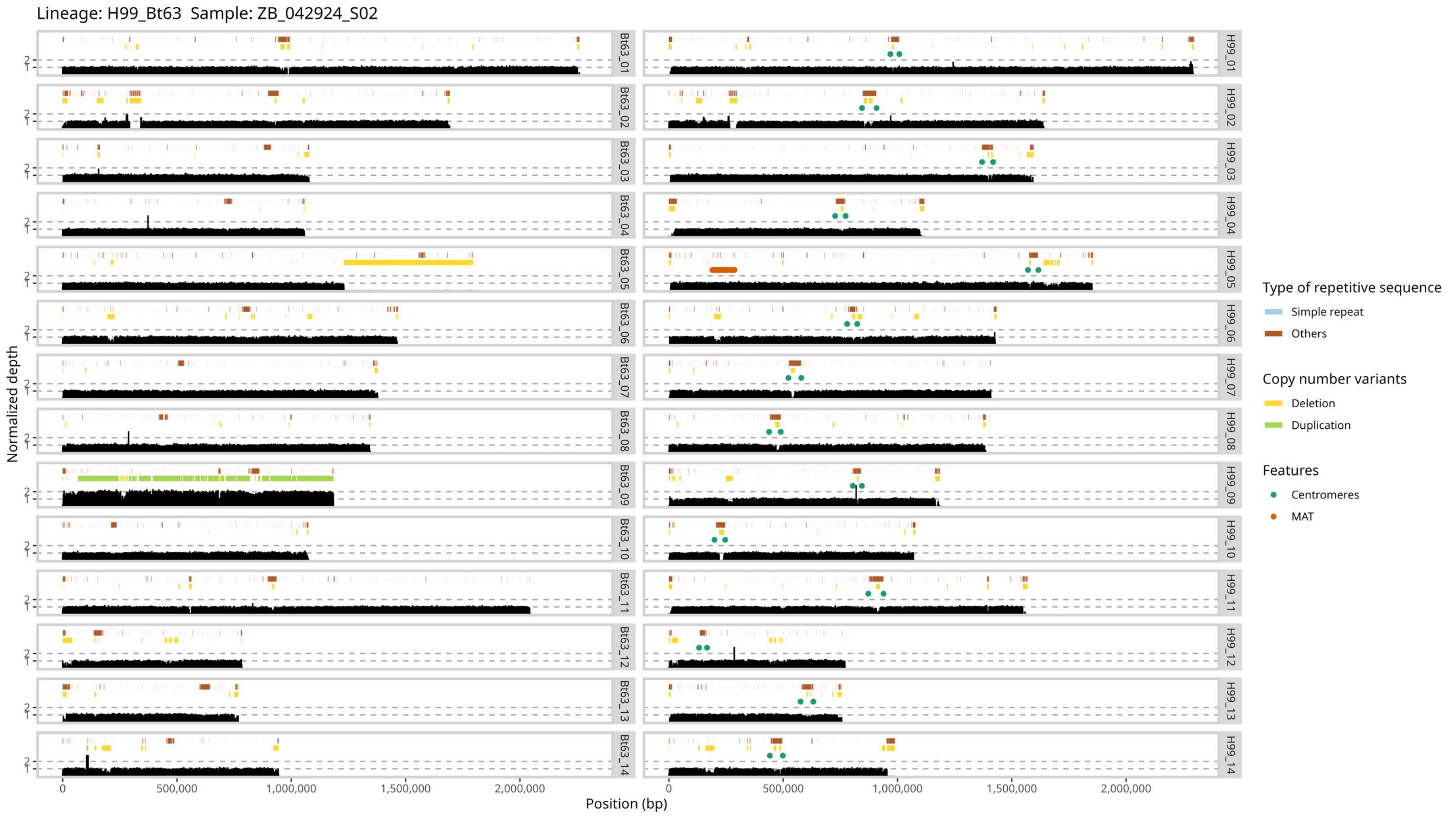

Lineage: H99_Bt63 Sample: ZB_042924_S02
Normalized depth
Position (bp)
0
500,000
1,000,000
1,500,000
2,000,000
Bt63_01
Bt63_02
Bt63_03
Bt63_04
Bt63_05
Bt63_06
Bt63_07
Bt63_08
Bt63_09
Bt63_10
Bt63_11
Bt63_12
Bt63_13
Bt63_14
H99_01
H99_02
H99_03
H99_04
H99_05
H99_06
H99_07
H99_08
H99_09
H99_10
H99_11
H99_12
H99_13
H99_14
Type of repetitive sequence
Simple repeat
Others
Copy number variants
Deletion
Duplication
Features
Centromeres
MAT

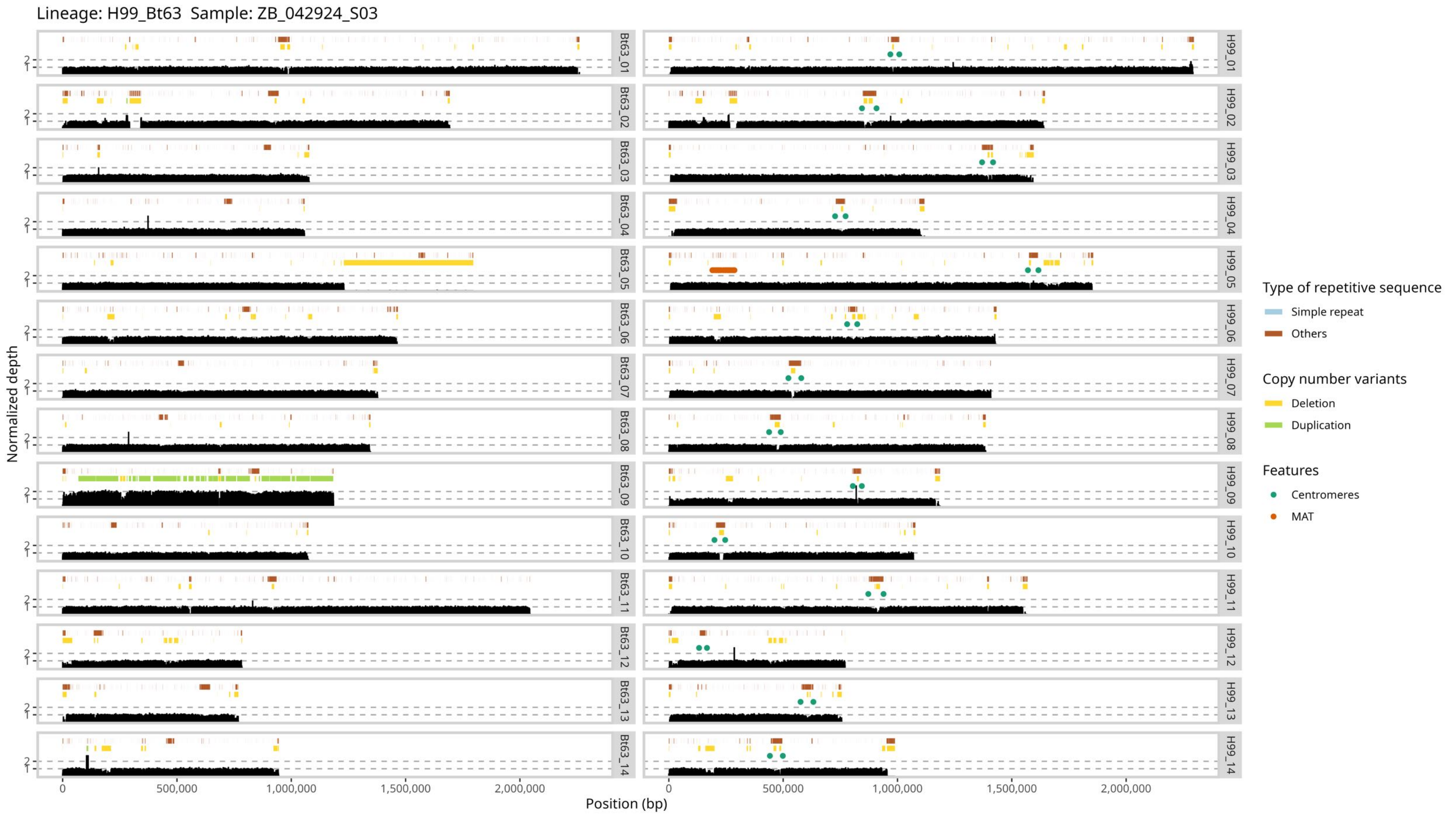

Lineage: H99_Bt63 Sample: ZB_042924_S03
Bt63_01
Bt63_02
Bt63_03
Bt63_04
Bt63_05
Bt63_06
Bt63_07
Bt63_08
Bt63_09
Bt63_10
Bt63_11
Bt63_12
Bt63_13
Bt63_14
H99_01
H99_02
H99_03
H99_04
H99_05
H99_06
H99_07
H99_08
H99_09
H99_10
H99_11
H99_12
H99_13
H99_14
Normalized depth
Position (bp)
0
500,000
1,000,000
1,500,000
2,000,000
Type of repetitive sequence
Simple repeat
Others
Copy number variants
Deletion
Duplication
Features
Centromeres
MAT

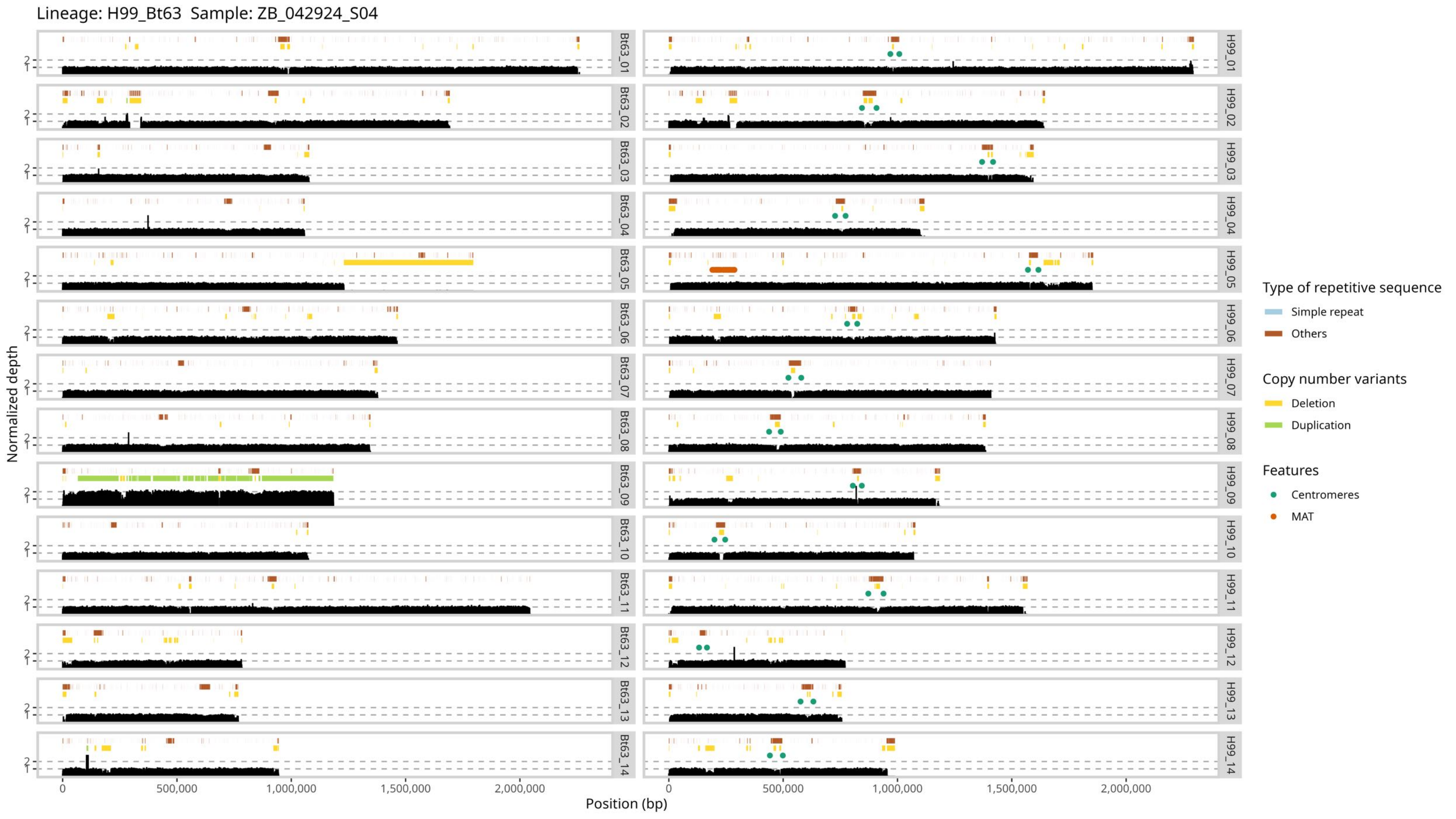

Lineage: H99_Bt63 Sample: ZB_042924_S04
Normalized depth
Position (bp)
Bt63_01
Bt63_02
Bt63_03
Bt63_04
Bt63_05
Bt63_06
Bt63_07
Bt63_08
Bt63_09
Bt63_10
Bt63_11
Bt63_12
Bt63_13
Bt63_14
H99_01
H99_02
H99_03
H99_04
H99_05
H99_06
H99_07
H99_08
H99_09
H99_10
H99_11
H99_12
H99_13
H99_14
0
500,000
1,000,000
1,500,000
2,000,000
Type of repetitive sequence
Simple repeat
Others
Copy number variants
Deletion
Duplication
Features
Centromeres
MAT

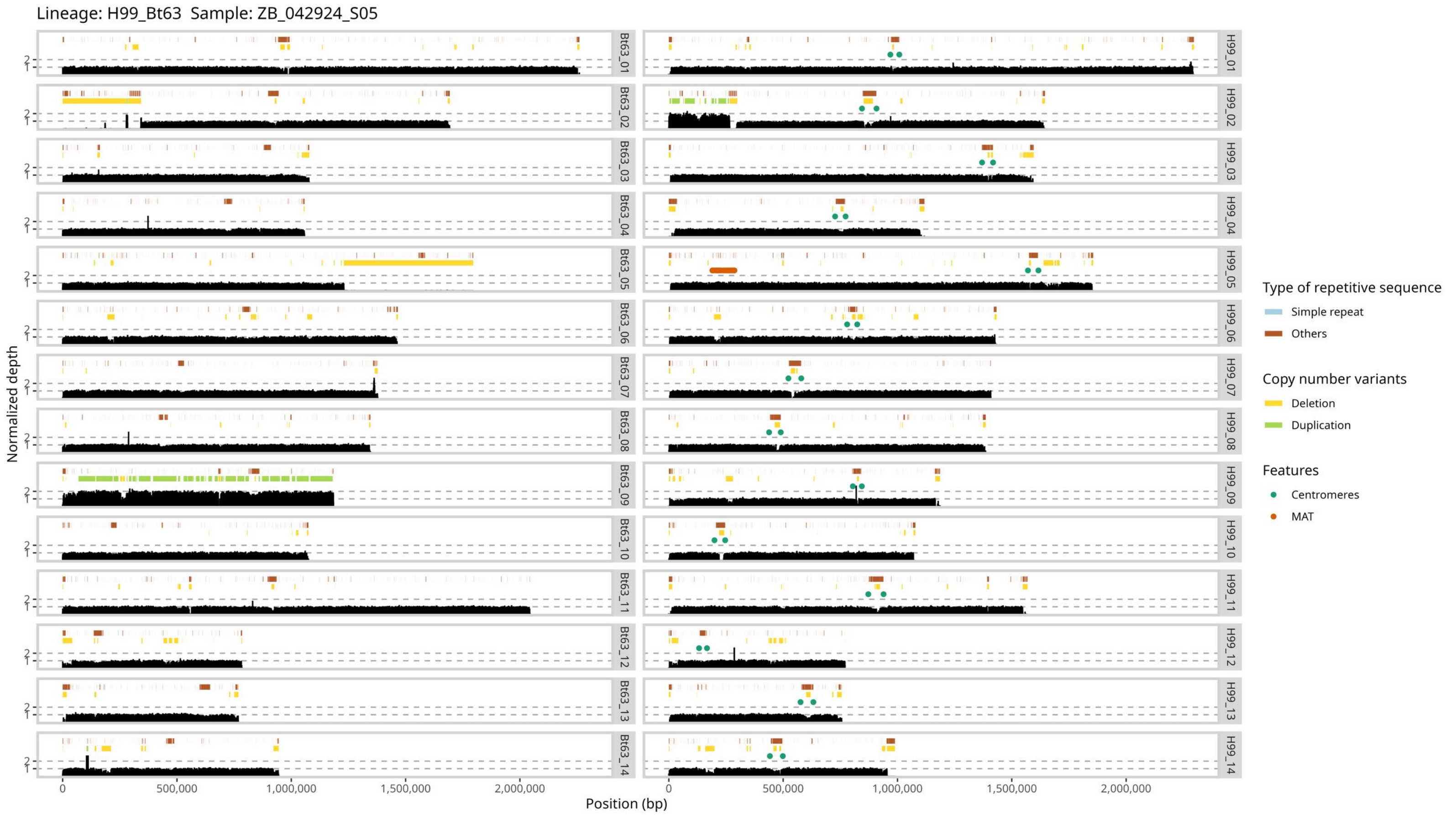

Lineage: H99_Bt63 Sample: ZB_042924_S05
Normalized depth
Position (bp)
Bt63_01
Bt63_02
Bt63_03
Bt63_04
Bt63_05
Bt63_06
Bt63_07
Bt63_08
Bt63_09
Bt63_10
Bt63_11
Bt63_12
Bt63_13
Bt63_14
H99_01
H99_02
H99_03
H99_04
H99_05
H99_06
H99_07
H99_08
H99_09
H99_10
H99_11
H99_12
H99_13
H99_14
0
500,000
1,000,000
1,500,000
2,000,000
Type of repetitive sequence
Simple repeat
Others
Copy number variants
Deletion
Duplication
Features
Centromeres
MAT

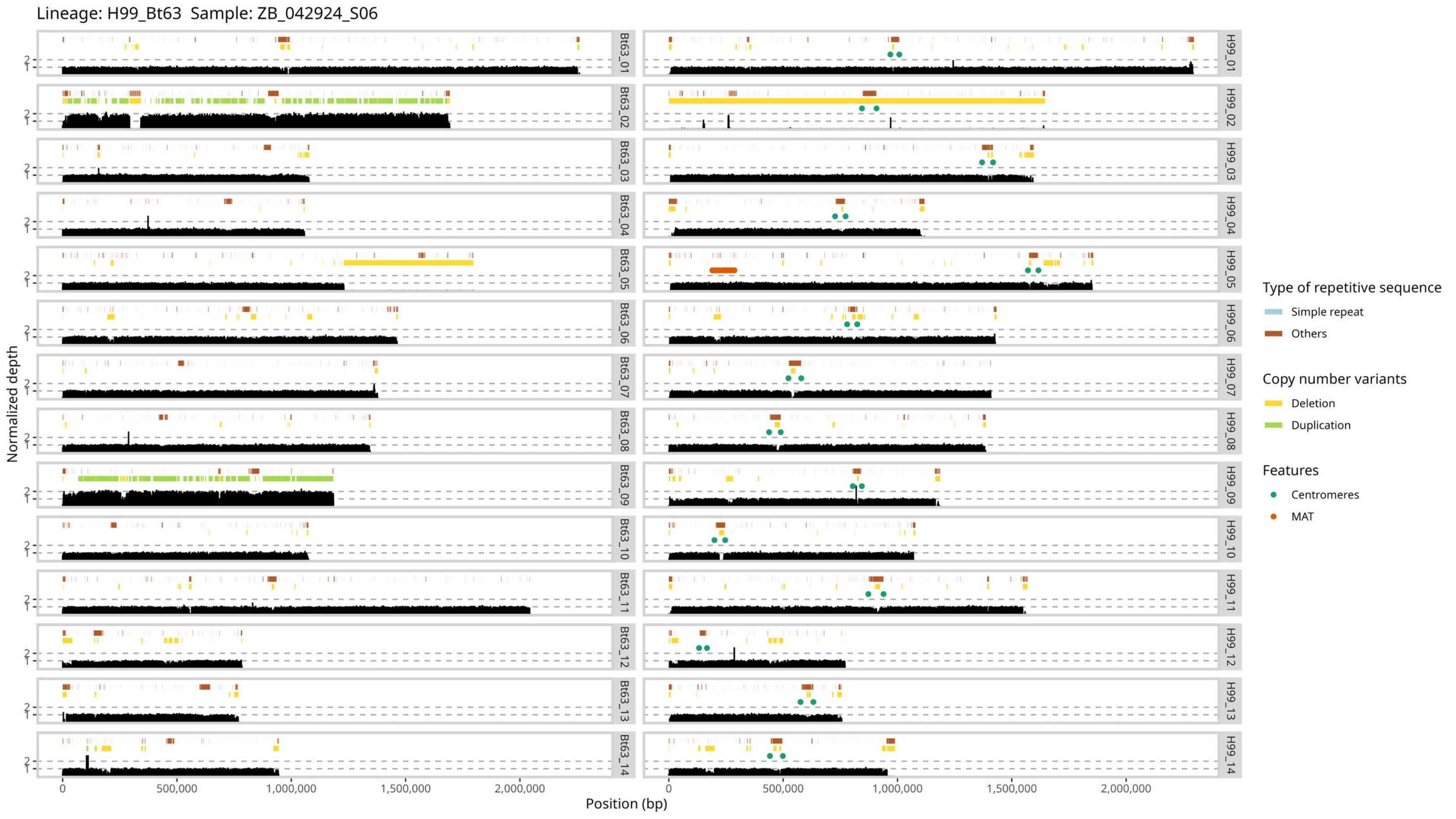

Lineage: H99_Bt63  Sample: ZB_042924_S06
Normalized depth
Position (bp)
Bt63_01
Bt63_02
Bt63_03
Bt63_04
Bt63_05
Bt63_06
Bt63_07
Bt63_08
Bt63_09
Bt63_10
Bt63_11
Bt63_12
Bt63_13
Bt63_14
H99_01
H99_02
H99_03
H99_04
H99_05
H99_06
H99_07
H99_08
H99_09
H99_10
H99_11
H99_12
H99_13
H99_14
0
500,000
1,000,000
1,500,000
2,000,000
Type of repetitive sequence
Simple repeat
Others
Copy number variants
Deletion
Duplication
Features
Centromeres
MAT

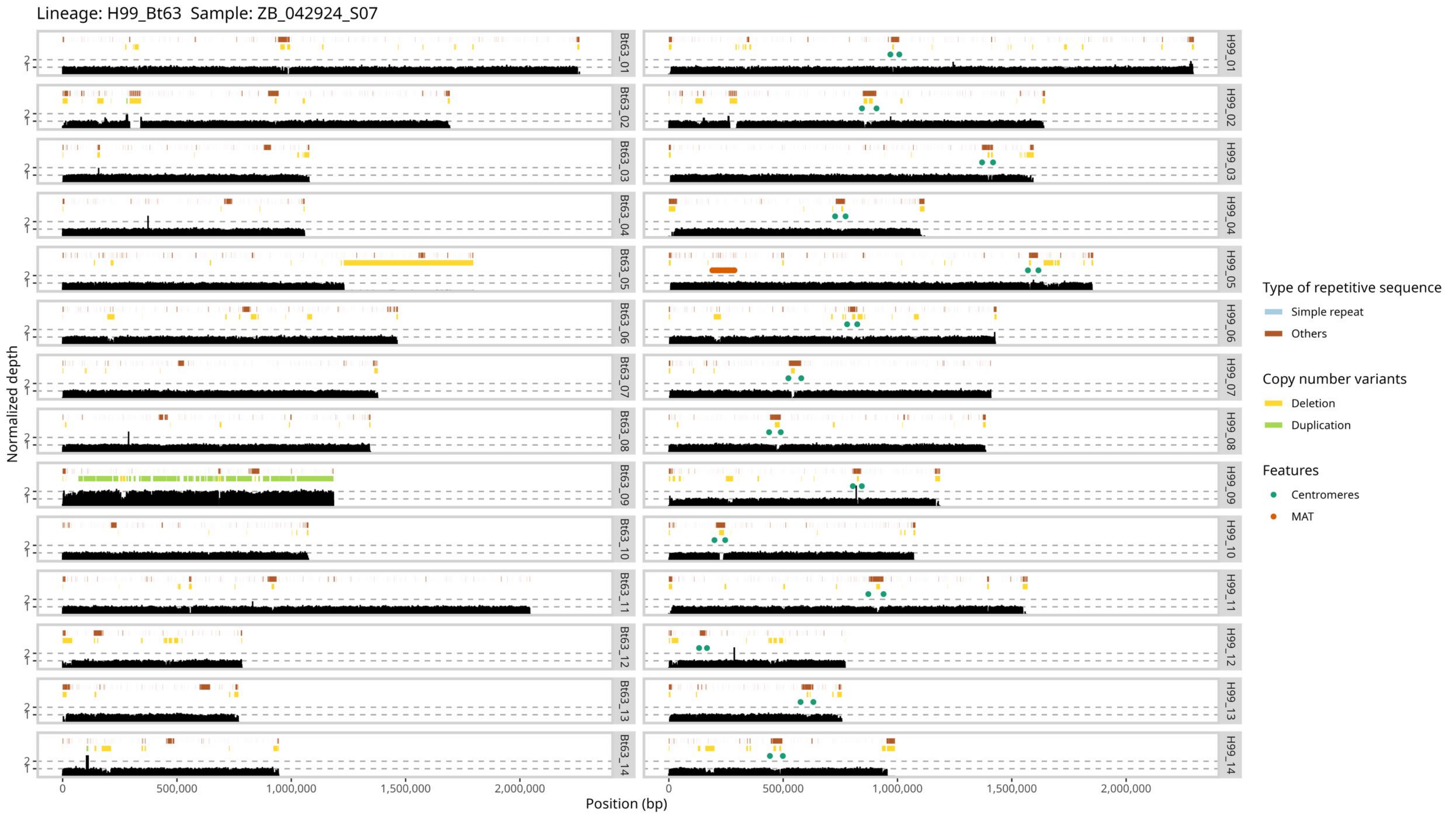
Lineage: H99_Bt63  Sample: ZB_042924_S07
Normalized depth
Position (bp)
0
500,000
1,000,000
1,500,000
2,000,000
1
2
Bt63_01
Bt63_02
Bt63_03
Bt63_04
Bt63_05
Bt63_06
Bt63_07
Bt63_08
Bt63_09
Bt63_10
Bt63_11
Bt63_12
Bt63_13
Bt63_14
H99_01
H99_02
H99_03
H99_04
H99_05
H99_06
H99_07
H99_08
H99_09
H99_10
H99_11
H99_12
H99_13
H99_14
Type of repetitive sequence
Simple repeat
Others
Copy number variants
Deletion
Duplication
Features
Centromeres
MAT

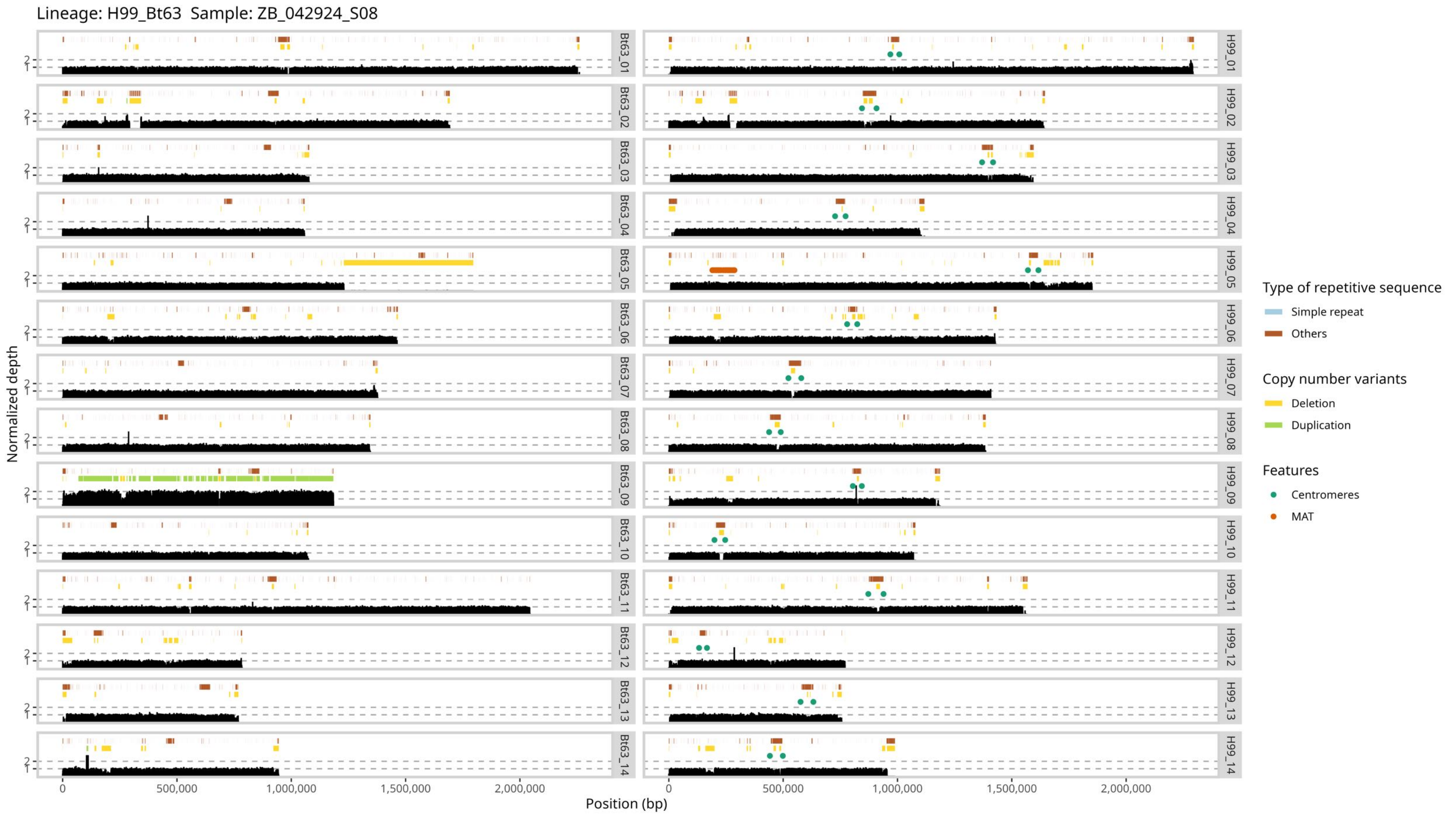

Lineage: H99_Bt63 Sample: ZB_042924_S08
Normalized depth
Position (bp)
0
500,000
1,000,000
1,500,000
2,000,000
1
2
Bt63_01
Bt63_02
Bt63_03
Bt63_04
Bt63_05
Bt63_06
Bt63_07
Bt63_08
Bt63_09
Bt63_10
Bt63_11
Bt63_12
Bt63_13
Bt63_14
H99_01
H99_02
H99_03
H99_04
H99_05
H99_06
H99_07
H99_08
H99_09
H99_10
H99_11
H99_12
H99_13
H99_14
Type of repetitive sequence
Simple repeat
Others
Copy number variants
Deletion
Duplication
Features
Centromeres
MAT

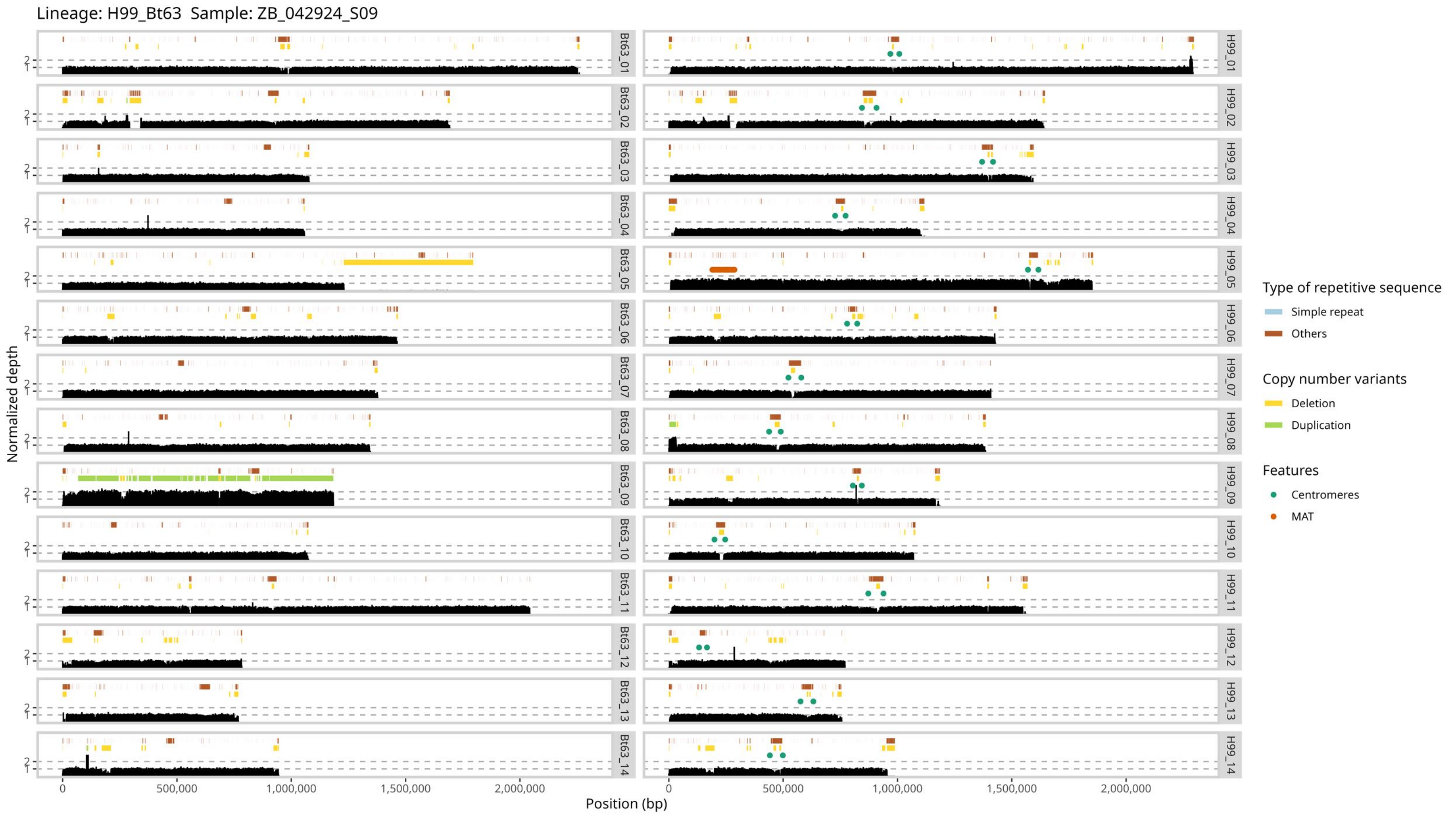

Lineage: H99_Bt63 Sample: ZB_042924_S09
Normalized depth
Position (bp)
0
500,000
1,000,000
1,500,000
2,000,000
Bt63_01
Bt63_02
Bt63_03
Bt63_04
Bt63_05
Bt63_06
Bt63_07
Bt63_08
Bt63_09
Bt63_10
Bt63_11
Bt63_12
Bt63_13
Bt63_14
H99_01
H99_02
H99_03
H99_04
H99_05
H99_06
H99_07
H99_08
H99_09
H99_10
H99_11
H99_12
H99_13
H99_14
Type of repetitive sequence
Simple repeat
Others
Copy number variants
Deletion
Duplication
Features
Centromeres
MAT

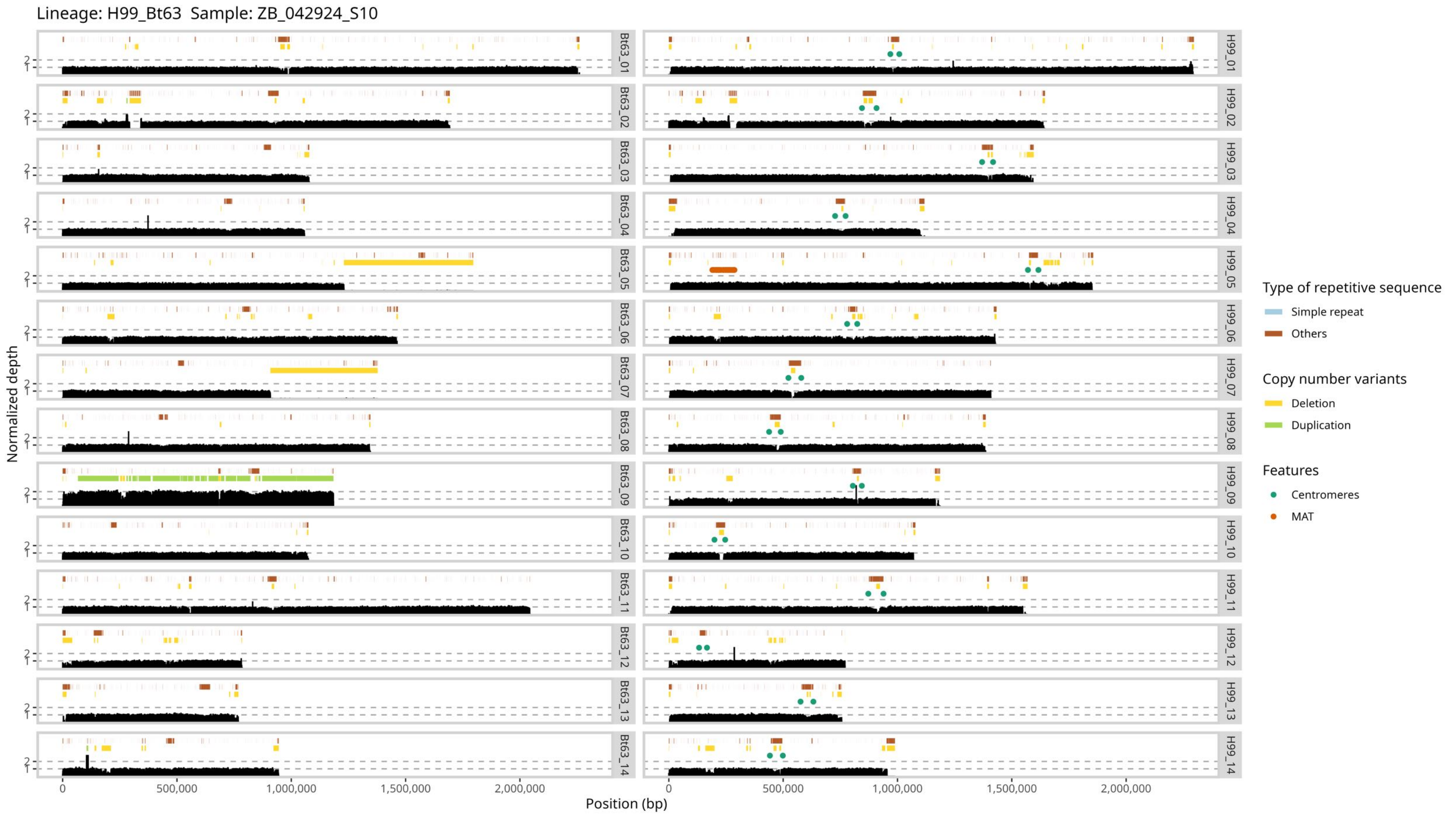

Lineage: H99_Bt63 Sample: ZB_042924_S10
Normalized depth
Position (bp)
Bt63_01
Bt63_02
Bt63_03
Bt63_04
Bt63_05
Bt63_06
Bt63_07
Bt63_08
Bt63_09
Bt63_10
Bt63_11
Bt63_12
Bt63_13
Bt63_14
H99_01
H99_02
H99_03
H99_04
H99_05
H99_06
H99_07
H99_08
H99_09
H99_10
H99_11
H99_12
H99_13
H99_14
0
500,000
1,000,000
1,500,000
2,000,000
Type of repetitive sequence
Simple repeat
Others
Copy number variants
Deletion
Duplication
Features
Centromeres
MAT

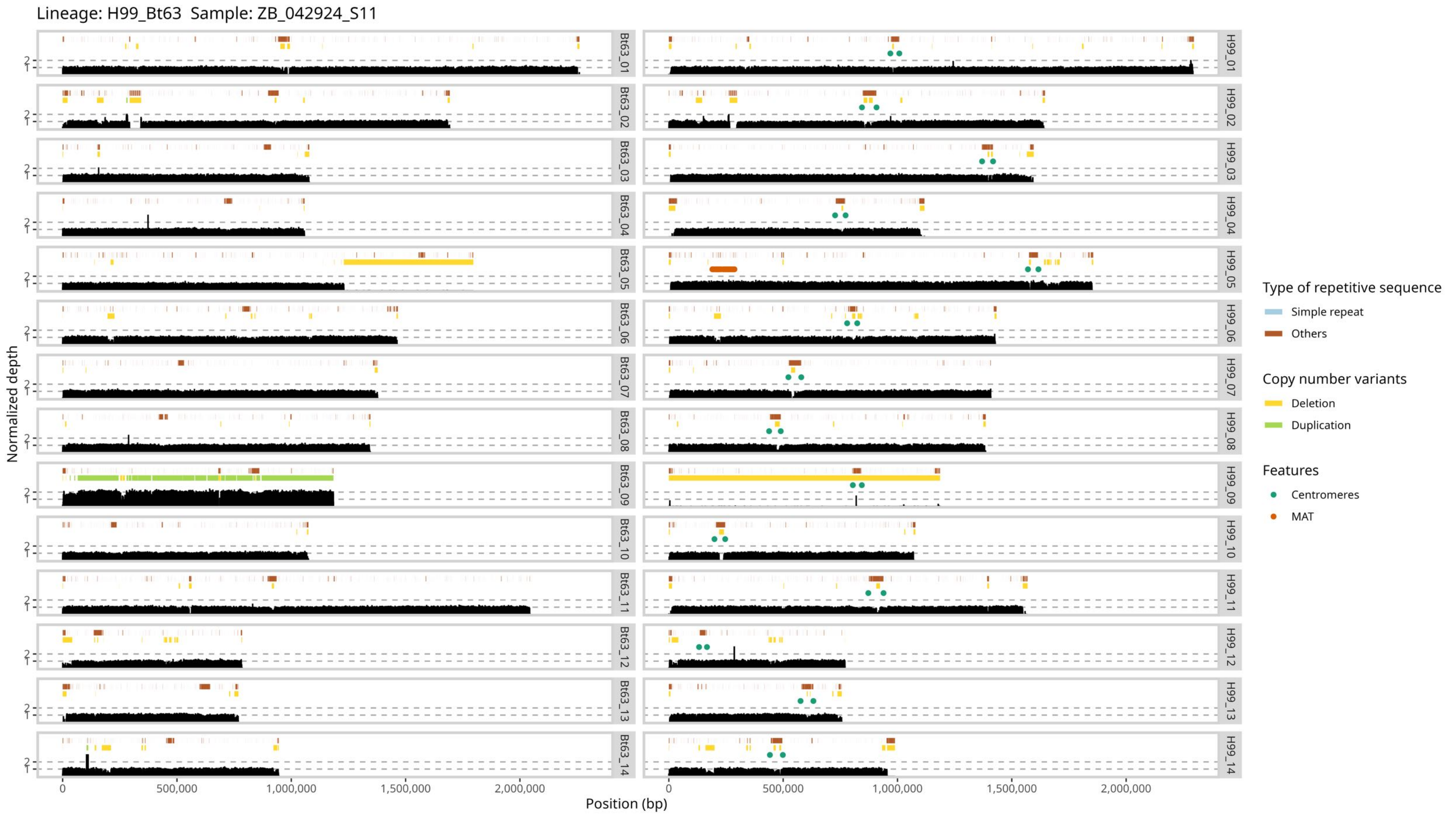

Lineage: H99_Bt63 Sample: ZB_042924_S11
Normalized depth
Position (bp)
Bt63_01
Bt63_02
Bt63_03
Bt63_04
Bt63_05
Bt63_06
Bt63_07
Bt63_08
Bt63_09
Bt63_10
Bt63_11
Bt63_12
Bt63_13
Bt63_14
H99_01
H99_02
H99_03
H99_04
H99_05
H99_06
H99_07
H99_08
H99_09
H99_10
H99_11
H99_12
H99_13
H99_14
0
500,000
1,000,000
1,500,000
2,000,000
Type of repetitive sequence
Simple repeat
Others
Copy number variants
Deletion
Duplication
Features
Centromeres
MAT

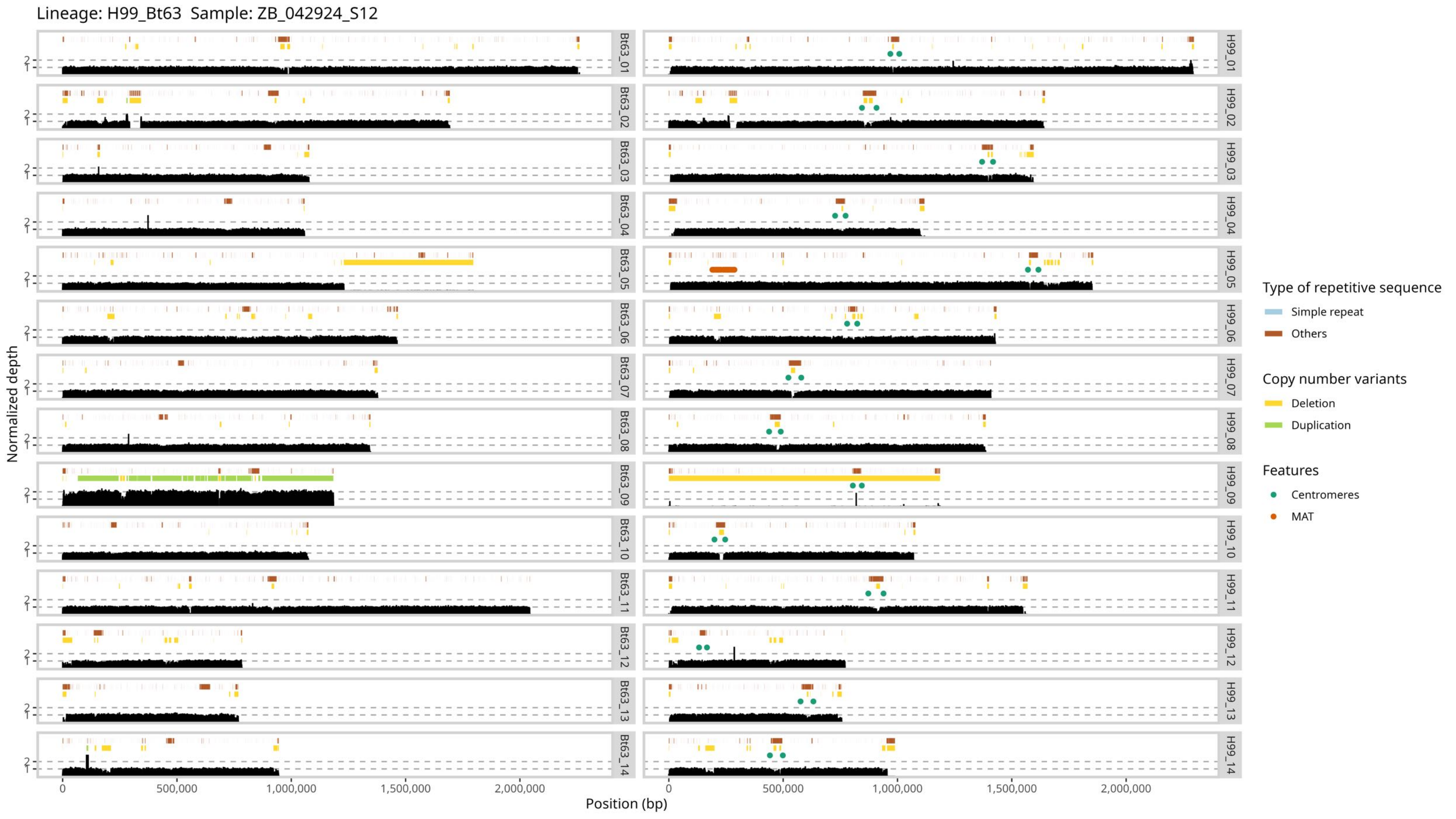
Lineage: H99_Bt63 Sample: ZB_042924_S12
Normalized depth
Position (bp)
0
500,000
1,000,000
1,500,000
2,000,000
Bt63_01
Bt63_02
Bt63_03
Bt63_04
Bt63_05
Bt63_06
Bt63_07
Bt63_08
Bt63_09
Bt63_10
Bt63_11
Bt63_12
Bt63_13
Bt63_14
H99_01
H99_02
H99_03
H99_04
H99_05
H99_06
H99_07
H99_08
H99_09
H99_10
H99_11
H99_12
H99_13
H99_14
Type of repetitive sequence
Simple repeat
Others
Copy number variants
Deletion
Duplication
Features
Centromeres
MAT

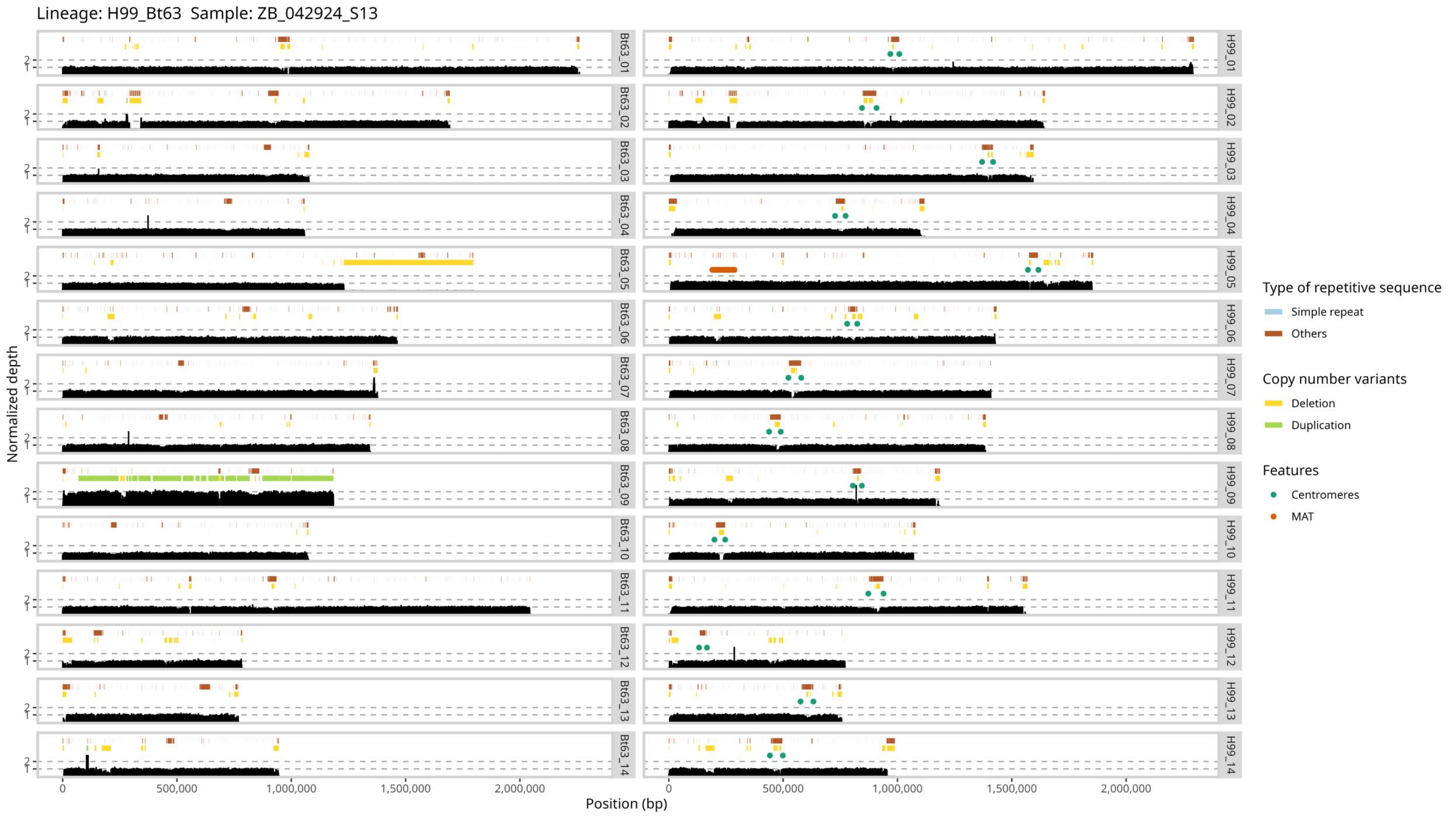

Lineage: H99_Bt63 Sample: ZB_042924_S13
Bt63_01
Bt63_02
Bt63_03
Bt63_04
Bt63_05
Bt63_06
Bt63_07
Bt63_08
Bt63_09
Bt63_10
Bt63_11
Bt63_12
Bt63_13
Bt63_14
H99_01
H99_02
H99_03
H99_04
H99_05
H99_06
H99_07
H99_08
H99_09
H99_10
H99_11
H99_12
H99_13
H99_14
Normalized depth
2
1
0
500,000
1,000,000
1,500,000
2,000,000
Position (bp)
Type of repetitive sequence
Simple repeat
Others
Copy number variants
Deletion
Duplication
Features
Centromeres
MAT

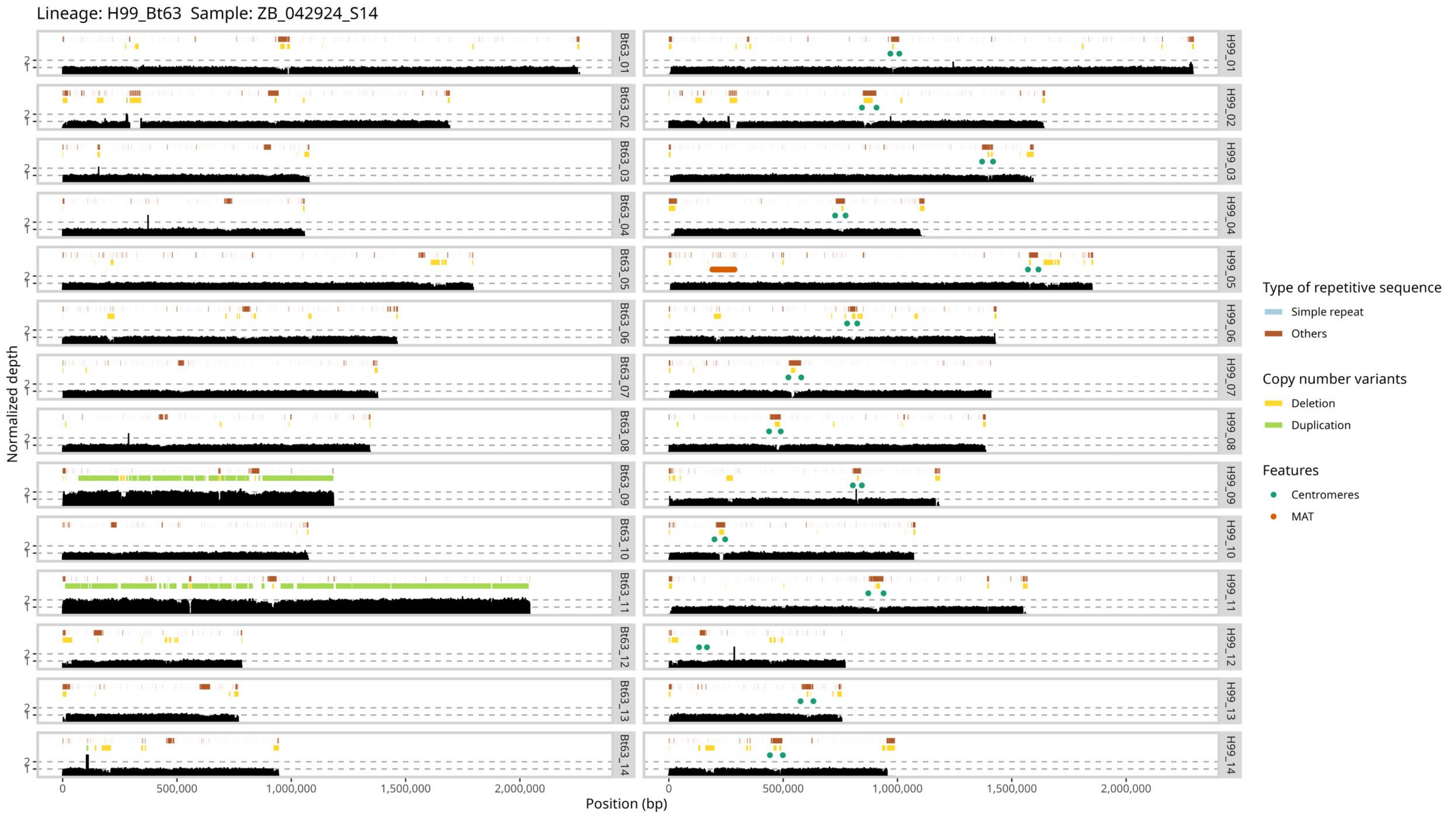

Lineage: H99_Bt63 Sample: ZB_042924_S14
Normalized depth
Position (bp)
0
500,000
1,000,000
1,500,000
2,000,000
Bt63_01
Bt63_02
Bt63_03
Bt63_04
Bt63_05
Bt63_06
Bt63_07
Bt63_08
Bt63_09
Bt63_10
Bt63_11
Bt63_12
Bt63_13
Bt63_14
H99_01
H99_02
H99_03
H99_04
H99_05
H99_06
H99_07
H99_08
H99_09
H99_10
H99_11
H99_12
H99_13
H99_14
Type of repetitive sequence
Simple repeat
Others
Copy number variants
Deletion
Duplication
Features
Centromeres
MAT

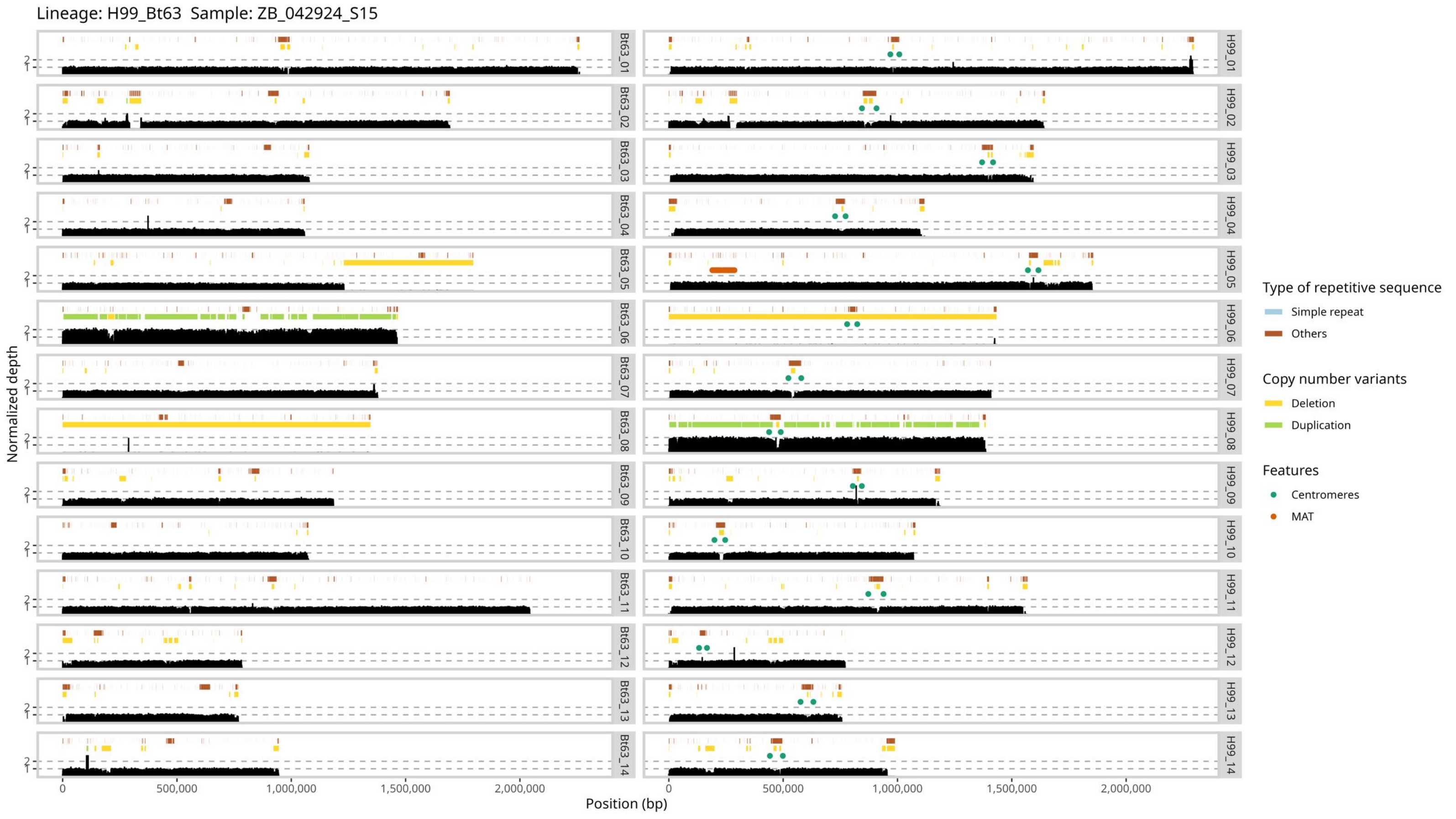

Lineage: H99_Bt63 Sample: ZB_042924_S15
Normalized depth
Position (bp)
0
500,000
1,000,000
1,500,000
2,000,000
Bt63_01
Bt63_02
Bt63_03
Bt63_04
Bt63_05
Bt63_06
Bt63_07
Bt63_08
Bt63_09
Bt63_10
Bt63_11
Bt63_12
Bt63_13
Bt63_14
H99_01
H99_02
H99_03
H99_04
H99_05
H99_06
H99_07
H99_08
H99_09
H99_10
H99_11
H99_12
H99_13
H99_14
Type of repetitive sequence
Simple repeat
Others
Copy number variants
Deletion
Duplication
Features
Centromeres
MAT

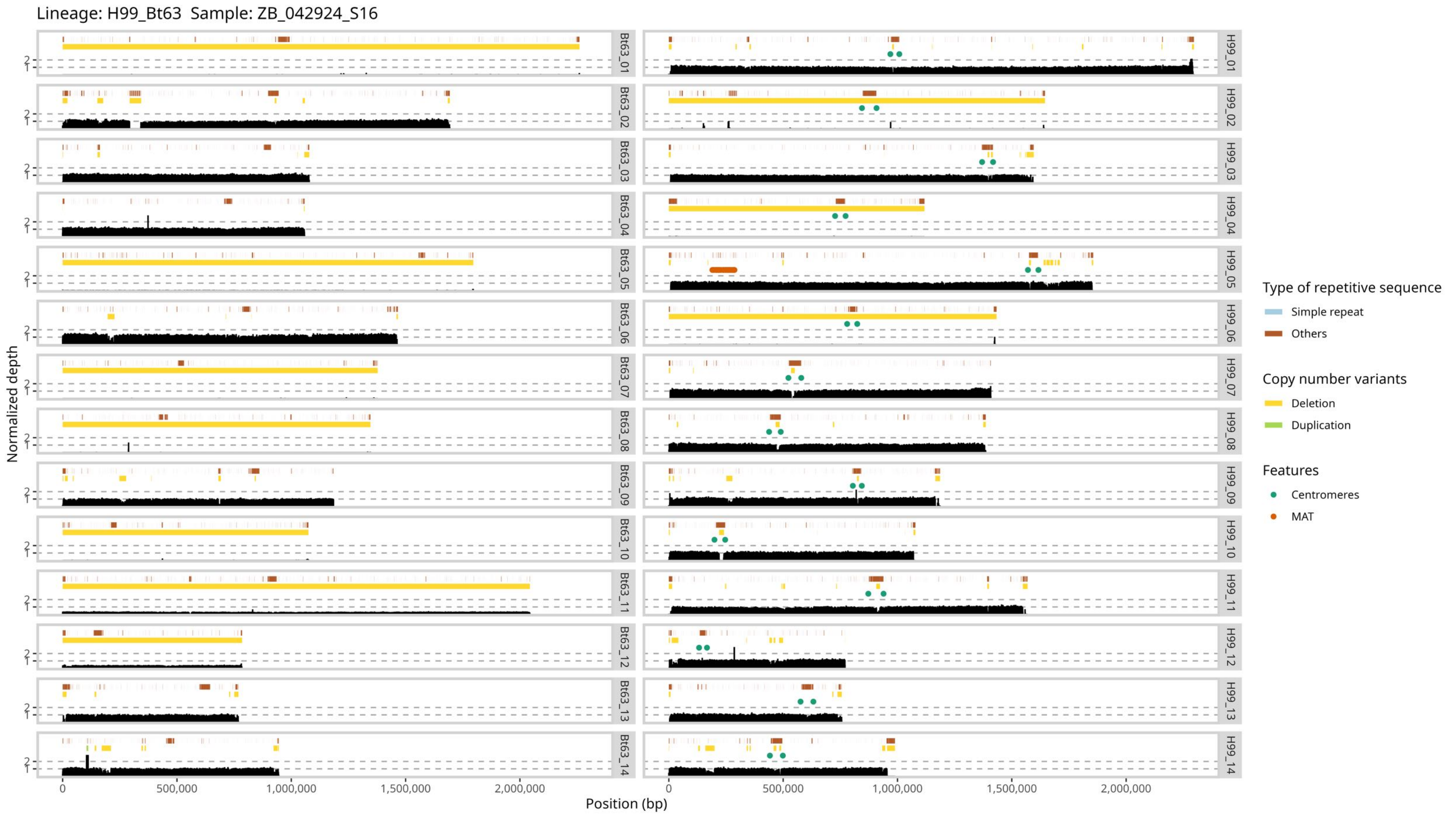

Lineage: H99_Bt63 Sample: ZB_042924_S16
Normalized depth
Position (bp)
0
500,000
1,000,000
1,500,000
2,000,000
Bt63_01
Bt63_02
Bt63_03
Bt63_04
Bt63_05
Bt63_06
Bt63_07
Bt63_08
Bt63_09
Bt63_10
Bt63_11
Bt63_12
Bt63_13
Bt63_14
H99_01
H99_02
H99_03
H99_04
H99_05
H99_06
H99_07
H99_08
H99_09
H99_10
H99_11
H99_12
H99_13
H99_14
Type of repetitive sequence
Simple repeat
Others
Copy number variants
Deletion
Duplication
Features
Centromeres
MAT

Lineage: H99_Bt63  Sample: ZB_042924_S17

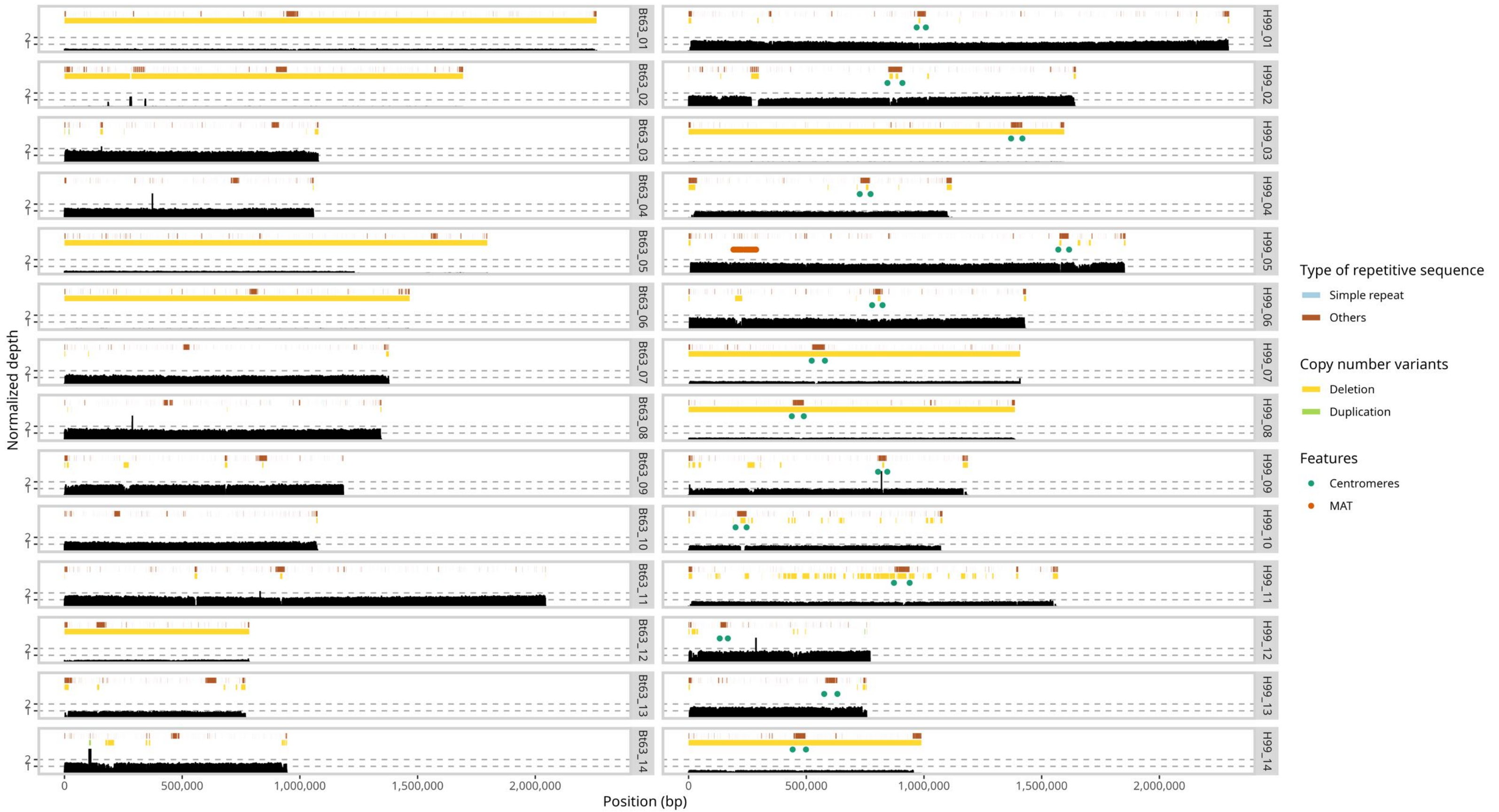

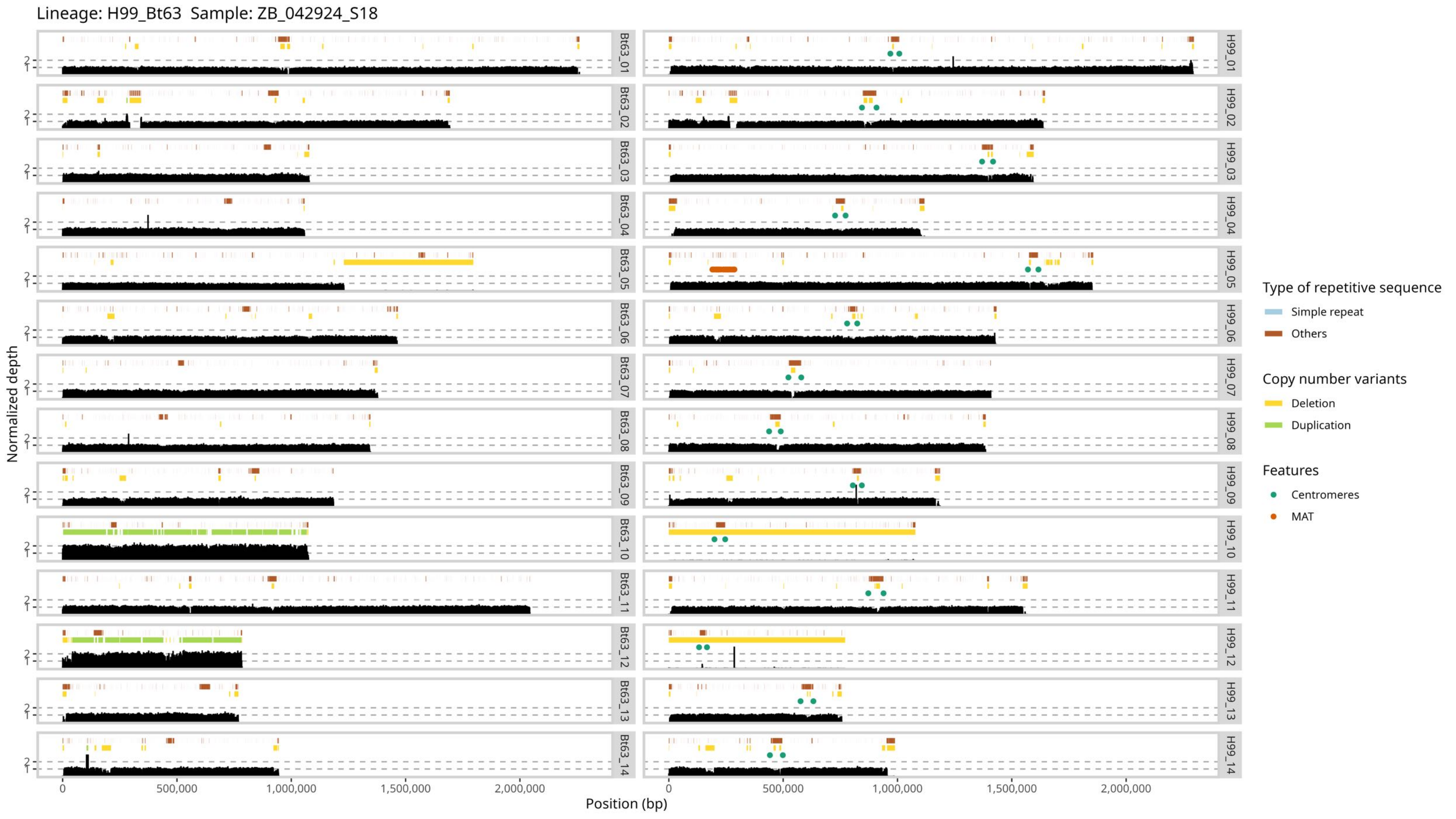

Lineage: H99_Bt63 Sample: ZB_042924_S18
Normalized depth
Position (bp)
0
500,000
1,000,000
1,500,000
2,000,000
1
2
Bt63_01
Bt63_02
Bt63_03
Bt63_04
Bt63_05
Bt63_06
Bt63_07
Bt63_08
Bt63_09
Bt63_10
Bt63_11
Bt63_12
Bt63_13
Bt63_14
H99_01
H99_02
H99_03
H99_04
H99_05
H99_06
H99_07
H99_08
H99_09
H99_10
H99_11
H99_12
H99_13
H99_14
Type of repetitive sequence
Simple repeat
Others
Copy number variants
Deletion
Duplication
Features
Centromeres
MAT

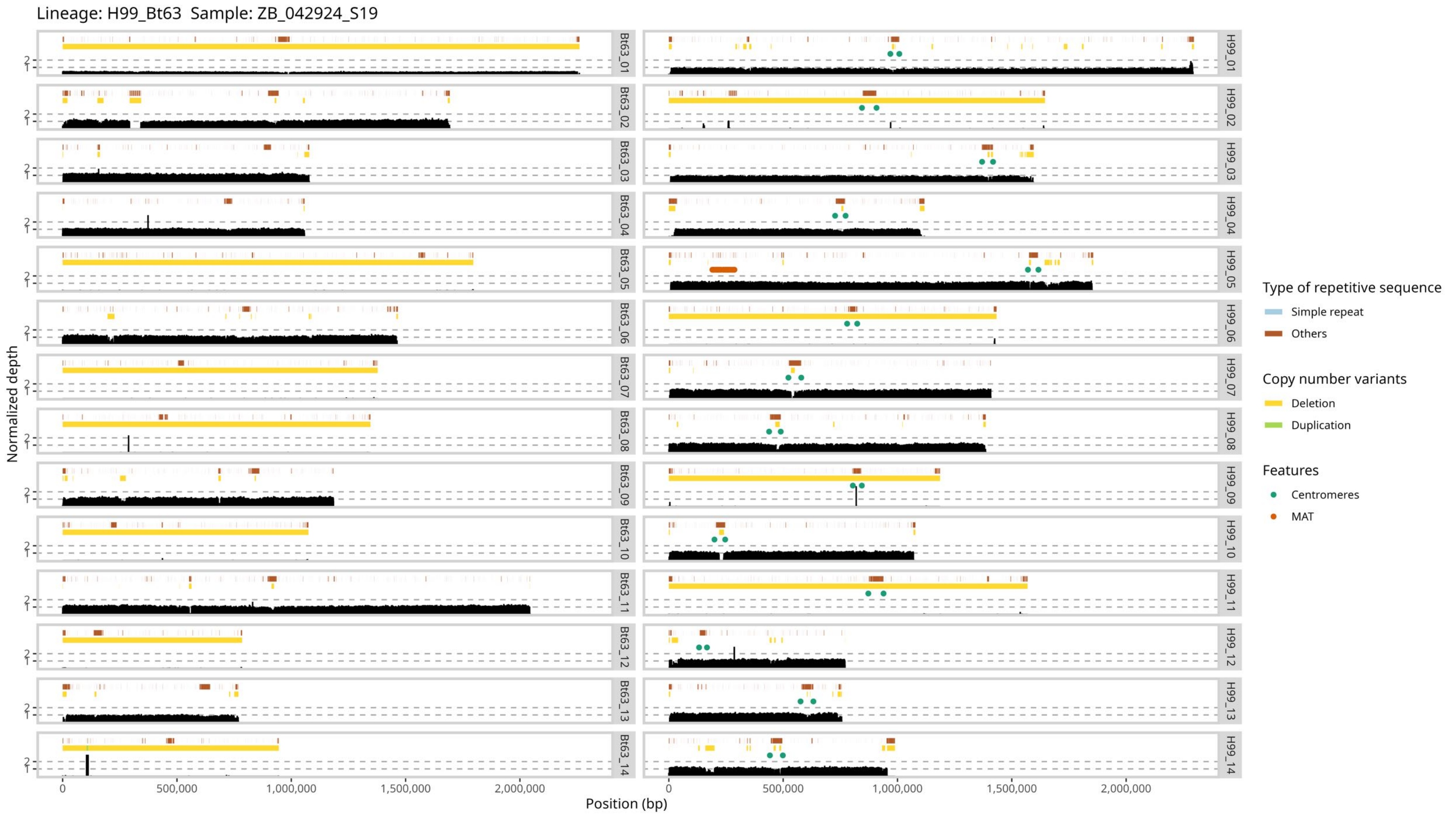

Lineage: H99_Bt63 Sample: ZB_042924_S19
Bt63_01
Bt63_02
Bt63_03
Bt63_04
Bt63_05
Bt63_06
Bt63_07
Bt63_08
Bt63_09
Bt63_10
Bt63_11
Bt63_12
Bt63_13
Bt63_14
H99_01
H99_02
H99_03
H99_04
H99_05
H99_06
H99_07
H99_08
H99_09
H99_10
H99_11
H99_12
H99_13
H99_14
Normalized depth
Position (bp)
0
500,000
1,000,000
1,500,000
2,000,000
Type of repetitive sequence
Simple repeat
Others
Copy number variants
Deletion
Duplication
Features
Centromeres
MAT

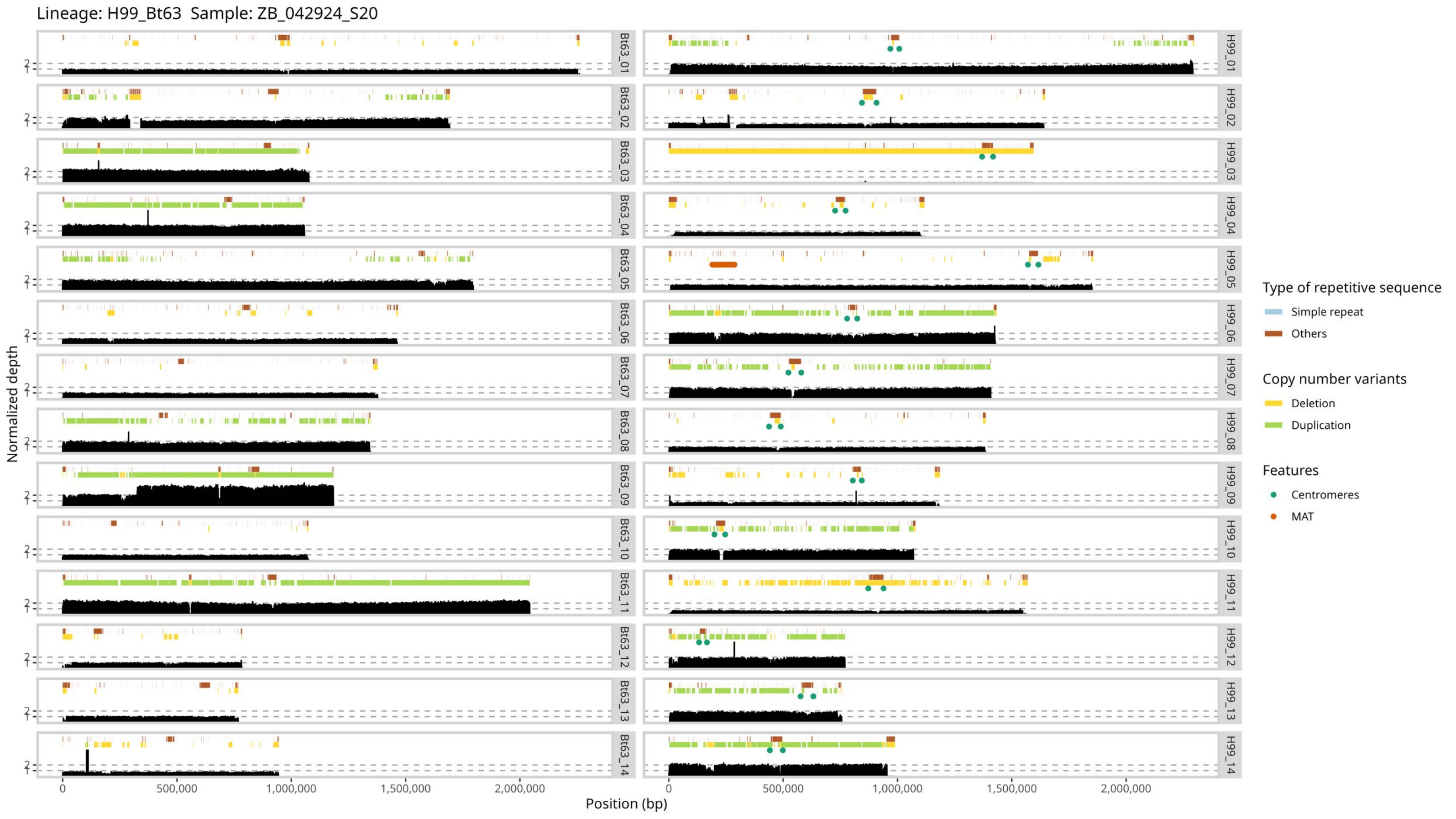

Lineage: H99_Bt63 Sample: ZB_042924_S20
Normalized depth
Position (bp)
0
500,000
1,000,000
1,500,000
2,000,000
Bt63_01
Bt63_02
Bt63_03
Bt63_04
Bt63_05
Bt63_06
Bt63_07
Bt63_08
Bt63_09
Bt63_10
Bt63_11
Bt63_12
Bt63_13
Bt63_14
H99_01
H99_02
H99_03
H99_04
H99_05
H99_06
H99_07
H99_08
H99_09
H99_10
H99_11
H99_12
H99_13
H99_14
Type of repetitive sequence
Simple repeat
Others
Copy number variants
Deletion
Duplication
Features
Centromeres
MAT

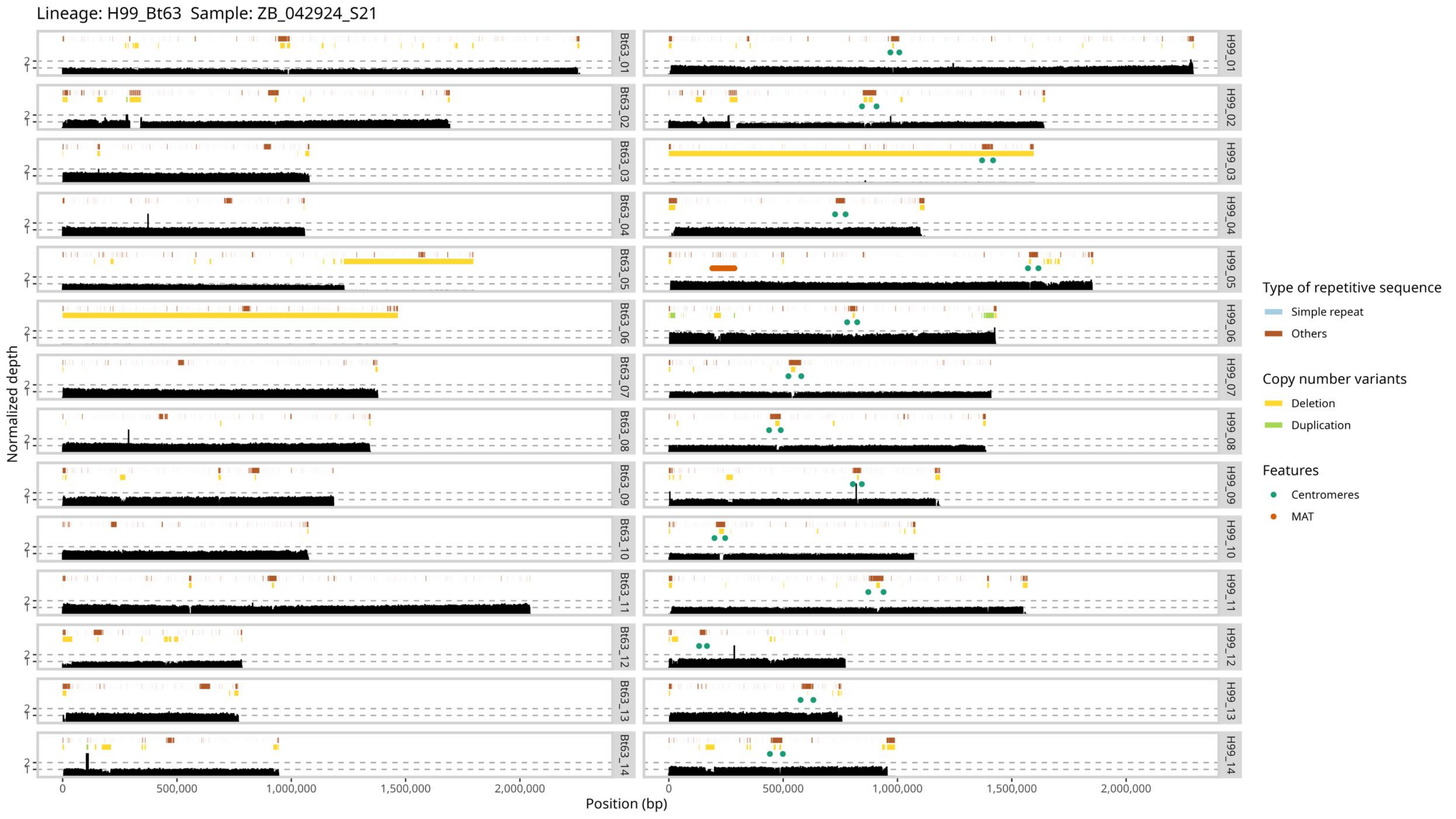

Lineage: H99_Bt63 Sample: ZB_042924_S21
Bt63_01
Bt63_02
Bt63_03
Bt63_04
Bt63_05
Bt63_06
Bt63_07
Bt63_08
Bt63_09
Bt63_10
Bt63_11
Bt63_12
Bt63_13
Bt63_14
H99_01
H99_02
H99_03
H99_04
H99_05
H99_06
H99_07
H99_08
H99_09
H99_10
H99_11
H99_12
H99_13
H99_14
Normalized depth
2
1
0
500,000
1,000,000
1,500,000
2,000,000
Position (bp)
Type of repetitive sequence
Simple repeat
Others
Copy number variants
Deletion
Duplication
Features
Centromeres
MAT

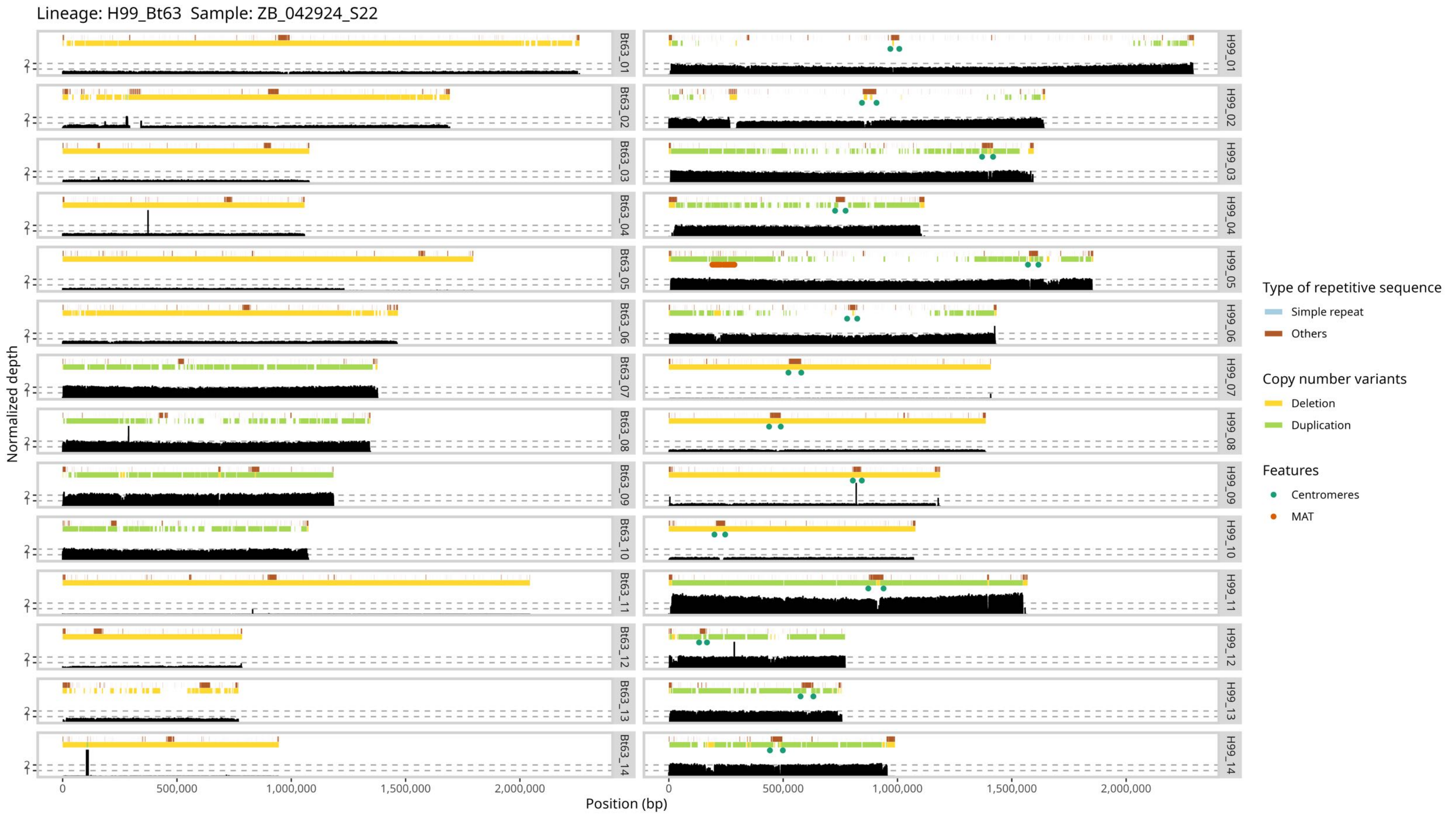

Lineage: H99_Bt63 Sample: ZB_042924_S22
Bt63_01
Bt63_02
Bt63_03
Bt63_04
Bt63_05
Bt63_06
Bt63_07
Bt63_08
Bt63_09
Bt63_10
Bt63_11
Bt63_12
Bt63_13
Bt63_14
H99_01
H99_02
H99_03
H99_04
H99_05
H99_06
H99_07
H99_08
H99_09
H99_10
H99_11
H99_12
H99_13
H99_14
Normalized depth
Position (bp)
0
500,000
1,000,000
1,500,000
2,000,000
Type of repetitive sequence
Simple repeat
Others
Copy number variants
Deletion
Duplication
Features
Centromeres
MAT

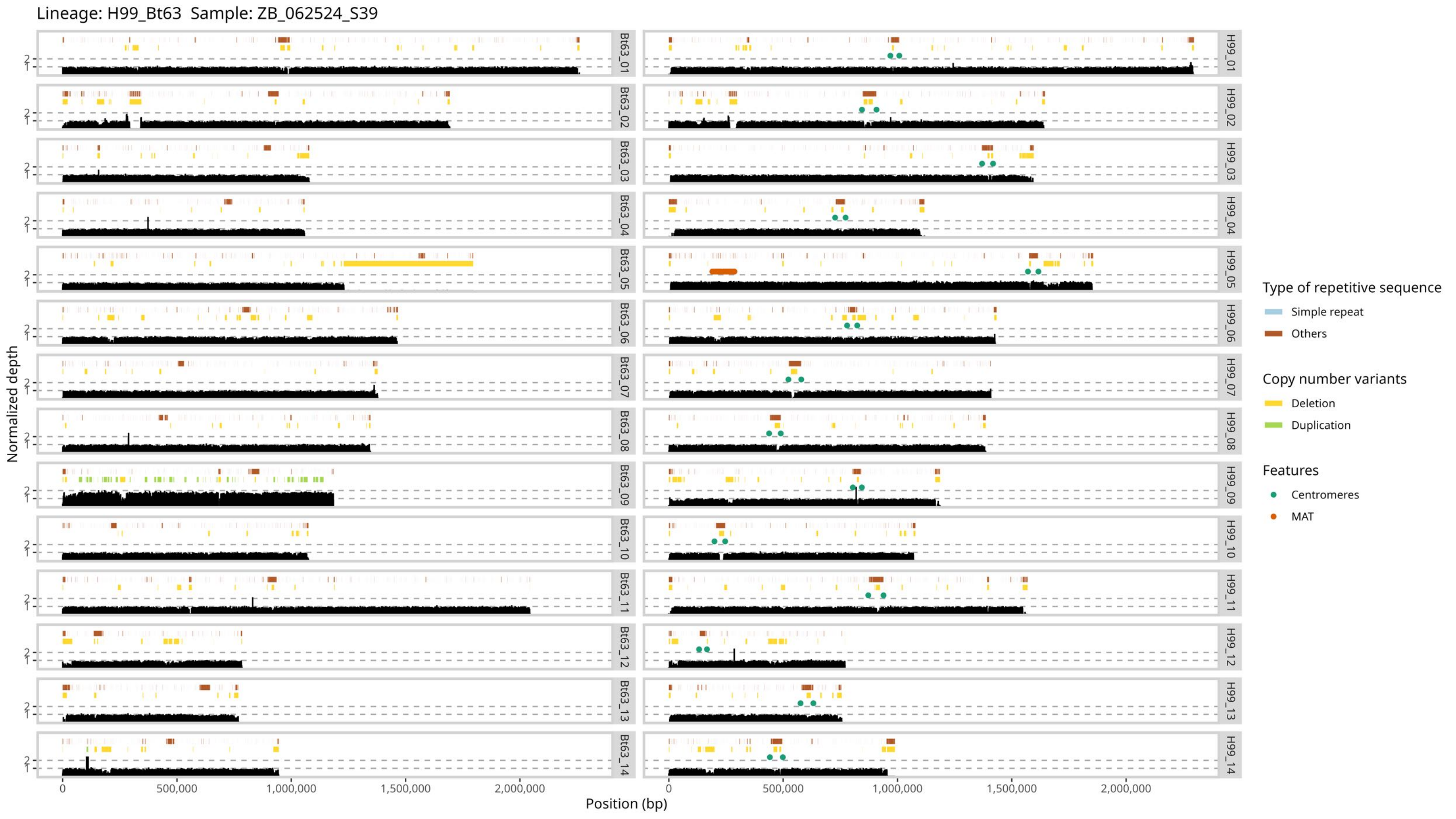

Lineage: H99_Bt63 Sample: ZB_062524_S39
Normalized depth
Position (bp)
0
500,000
1,000,000
1,500,000
2,000,000
Bt63_01
Bt63_02
Bt63_03
Bt63_04
Bt63_05
Bt63_06
Bt63_07
Bt63_08
Bt63_09
Bt63_10
Bt63_11
Bt63_12
Bt63_13
Bt63_14
H99_01
H99_02
H99_03
H99_04
H99_05
H99_06
H99_07
H99_08
H99_09
H99_10
H99_11
H99_12
H99_13
H99_14
Type of repetitive sequence
Simple repeat
Others
Copy number variants
Deletion
Duplication
Features
Centromeres
MAT

Lineage: H99_Bt63  Sample: ZB_062524_S40

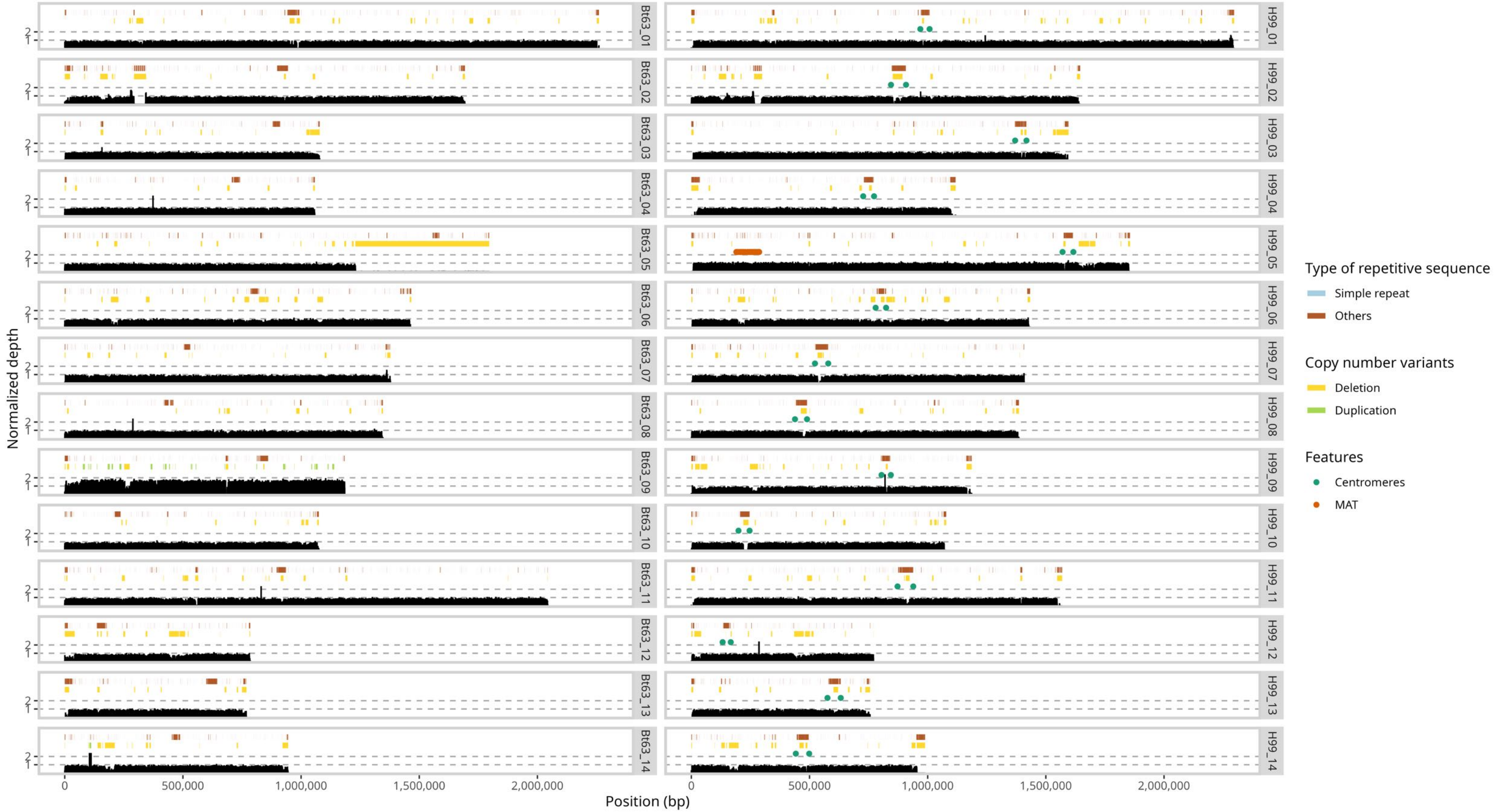

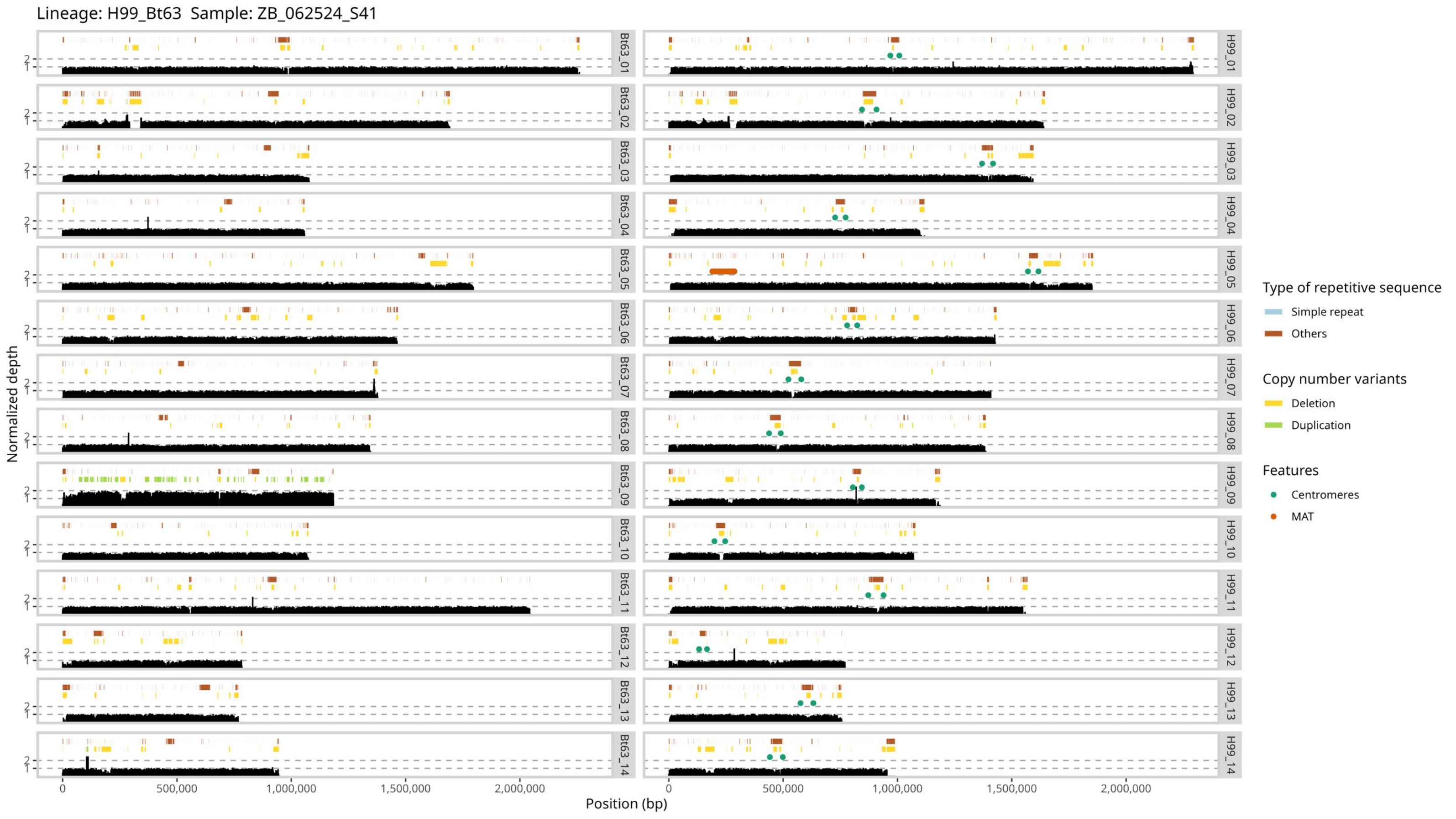
Lineage: H99_Bt63 Sample: ZB_062524_S41
Normalized depth
Position (bp)
Bt63_01
Bt63_02
Bt63_03
Bt63_04
Bt63_05
Bt63_06
Bt63_07
Bt63_08
Bt63_09
Bt63_10
Bt63_11
Bt63_12
Bt63_13
Bt63_14
H99_01
H99_02
H99_03
H99_04
H99_05
H99_06
H99_07
H99_08
H99_09
H99_10
H99_11
H99_12
H99_13
H99_14
0
500,000
1,000,000
1,500,000
2,000,000
Type of repetitive sequence
Simple repeat
Others
Copy number variants
Deletion
Duplication
Features
Centromeres
MAT

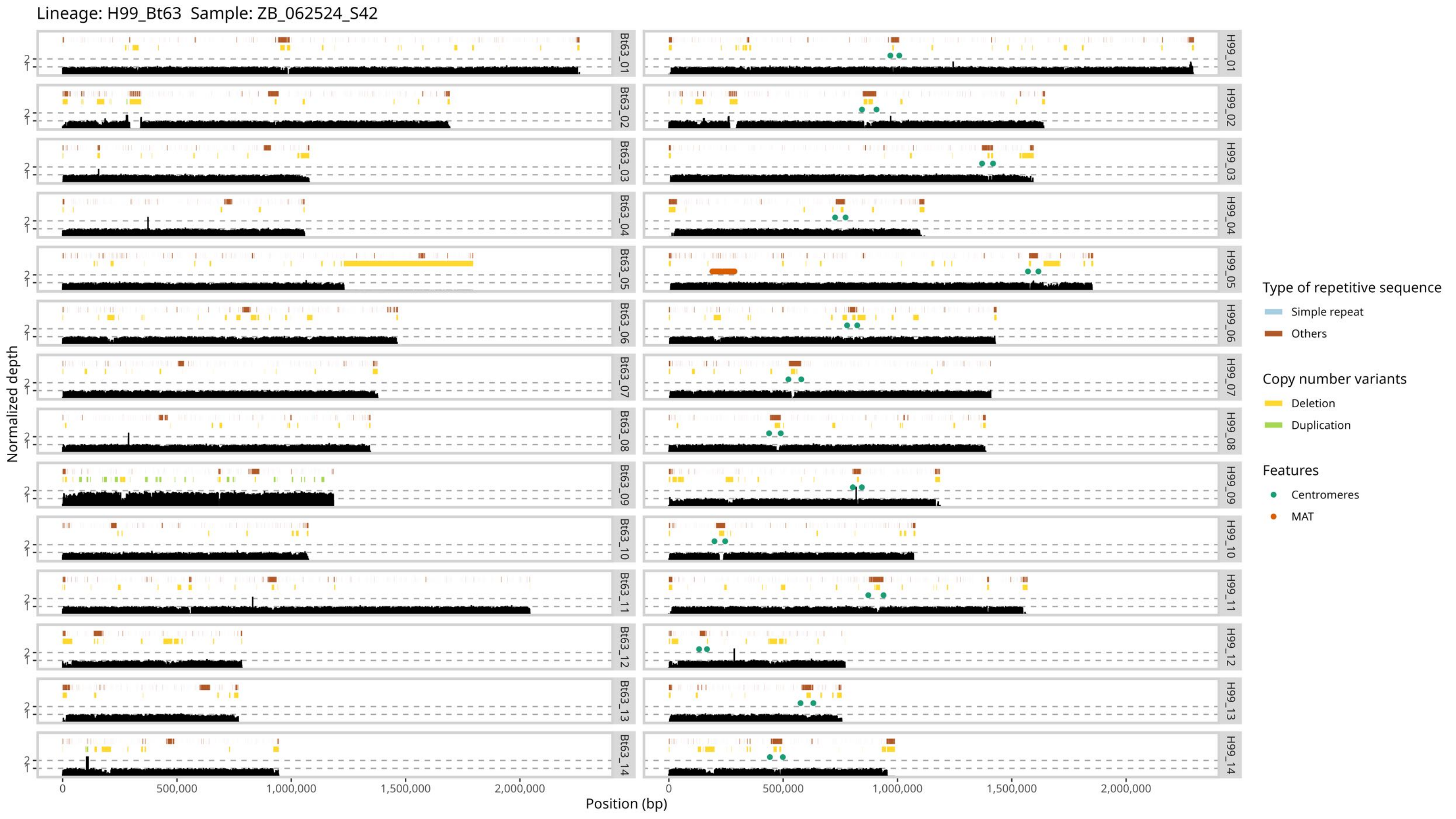

Lineage: H99_Bt63 Sample: ZB_062524_S42
Normalized depth
Position (bp)
0
500,000
1,000,000
1,500,000
2,000,000
Bt63_01
Bt63_02
Bt63_03
Bt63_04
Bt63_05
Bt63_06
Bt63_07
Bt63_08
Bt63_09
Bt63_10
Bt63_11
Bt63_12
Bt63_13
Bt63_14
H99_01
H99_02
H99_03
H99_04
H99_05
H99_06
H99_07
H99_08
H99_09
H99_10
H99_11
H99_12
H99_13
H99_14
Type of repetitive sequence
Simple repeat
Others
Copy number variants
Deletion
Duplication
Features
Centromeres
MAT

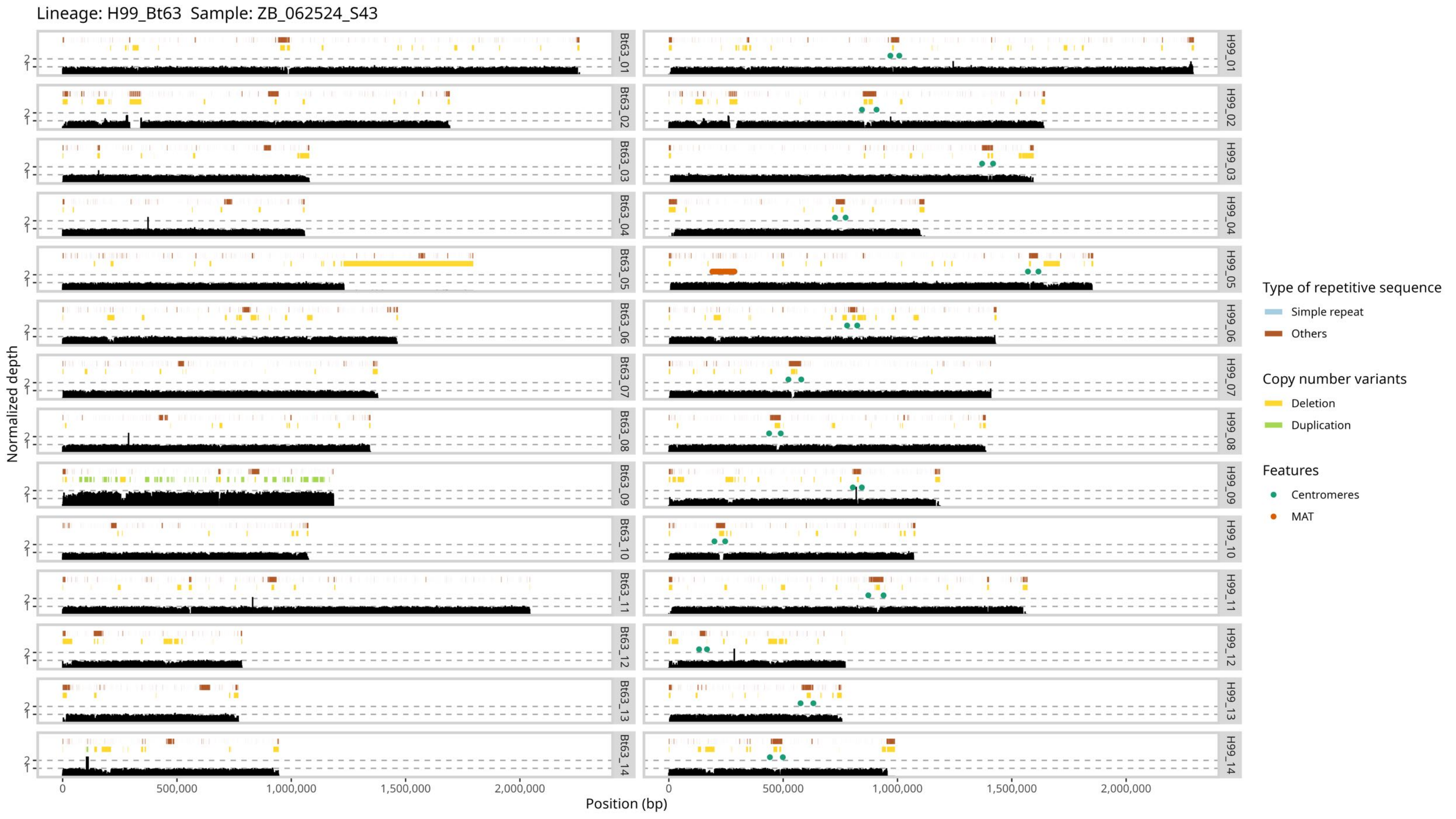
Lineage: H99_Bt63 Sample: ZB_062524_S43
Normalized depth
Position (bp)
0
500,000
1,000,000
1,500,000
2,000,000
1
2
Bt63_01
Bt63_02
Bt63_03
Bt63_04
Bt63_05
Bt63_06
Bt63_07
Bt63_08
Bt63_09
Bt63_10
Bt63_11
Bt63_12
Bt63_13
Bt63_14
H99_01
H99_02
H99_03
H99_04
H99_05
H99_06
H99_07
H99_08
H99_09
H99_10
H99_11
H99_12
H99_13
H99_14
Type of repetitive sequence
Simple repeat
Others
Copy number variants
Deletion
Duplication
Features
Centromeres
MAT

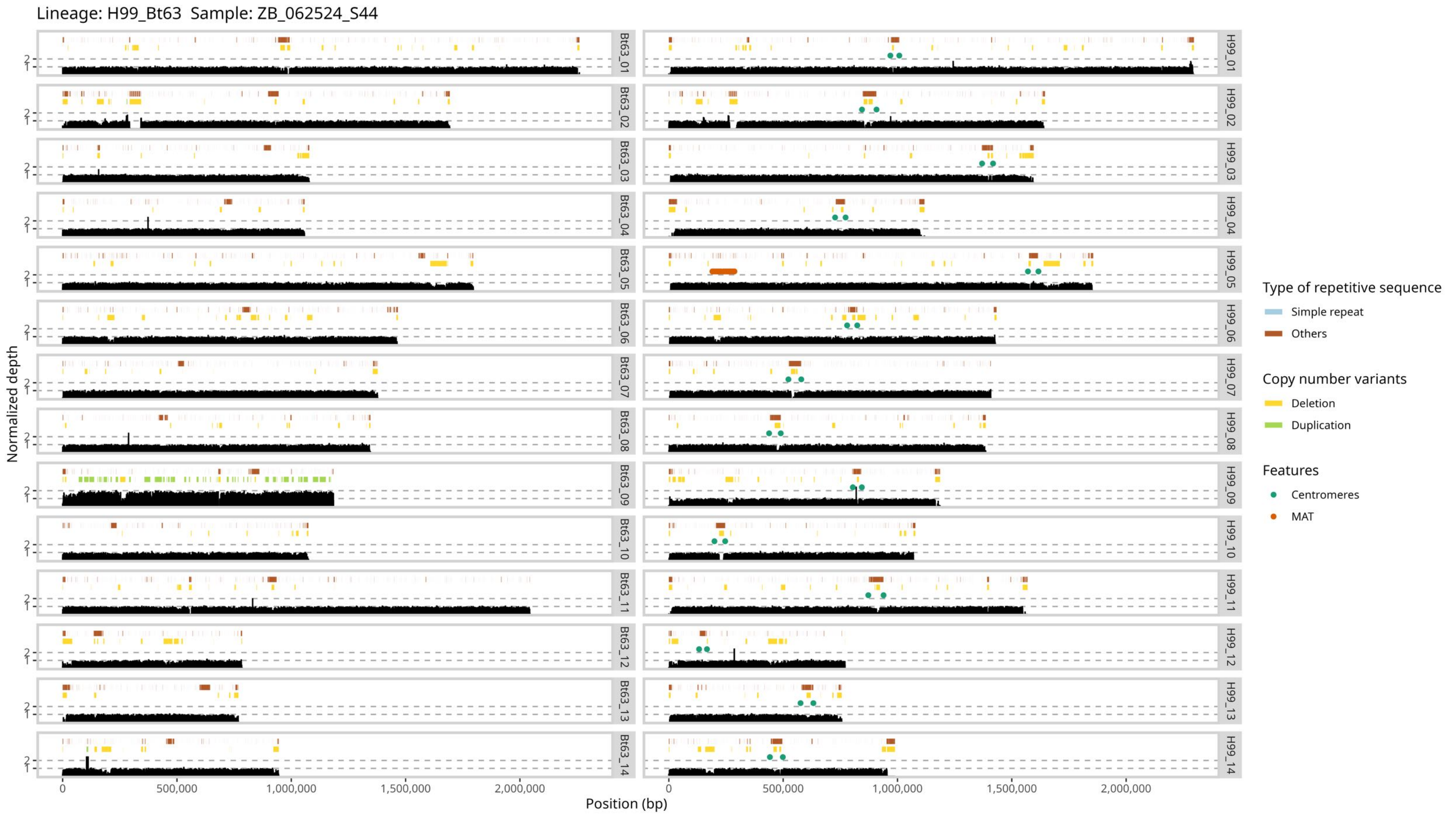

Lineage: H99_Bt63 Sample: ZB_062524_S44
Normalized depth
Position (bp)
0
500,000
1,000,000
1,500,000
2,000,000
1
2
Bt63_01
Bt63_02
Bt63_03
Bt63_04
Bt63_05
Bt63_06
Bt63_07
Bt63_08
Bt63_09
Bt63_10
Bt63_11
Bt63_12
Bt63_13
Bt63_14
H99_01
H99_02
H99_03
H99_04
H99_05
H99_06
H99_07
H99_08
H99_09
H99_10
H99_11
H99_12
H99_13
H99_14
Type of repetitive sequence
Simple repeat
Others
Copy number variants
Deletion
Duplication
Features
Centromeres
MAT

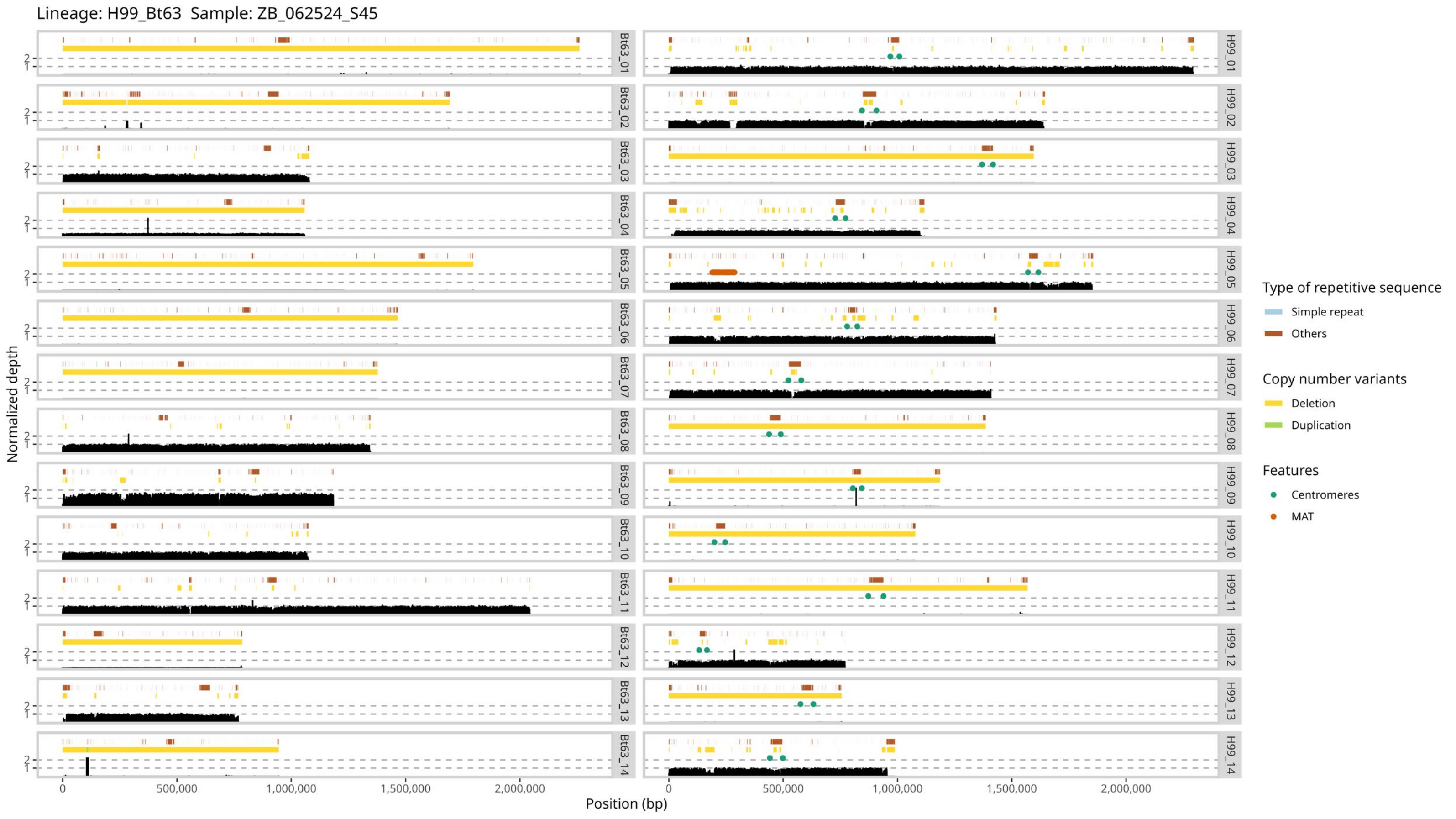

Lineage: H99_Bt63 Sample: ZB_062524_S45
Normalized depth
Position (bp)
0
500,000
1,000,000
1,500,000
2,000,000
Bt63_01
Bt63_02
Bt63_03
Bt63_04
Bt63_05
Bt63_06
Bt63_07
Bt63_08
Bt63_09
Bt63_10
Bt63_11
Bt63_12
Bt63_13
Bt63_14
H99_01
H99_02
H99_03
H99_04
H99_05
H99_06
H99_07
H99_08
H99_09
H99_10
H99_11
H99_12
H99_13
H99_14
Type of repetitive sequence
Simple repeat
Others
Copy number variants
Deletion
Duplication
Features
Centromeres
MAT

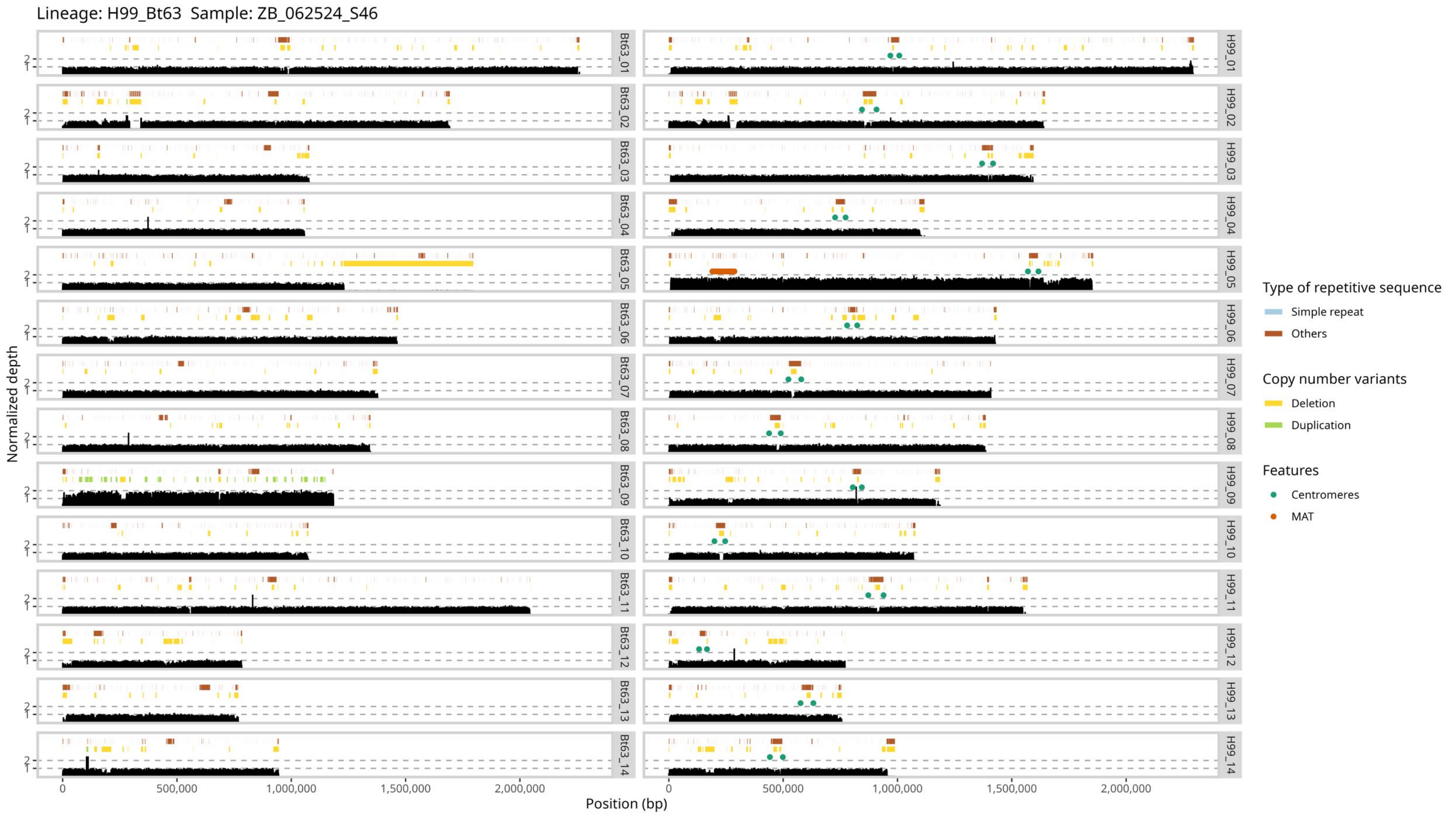

Lineage: H99_Bt63 Sample: ZB_062524_S46
Normalized depth
Position (bp)
0
500,000
1,000,000
1,500,000
2,000,000
1
2
Bt63_01
Bt63_02
Bt63_03
Bt63_04
Bt63_05
Bt63_06
Bt63_07
Bt63_08
Bt63_09
Bt63_10
Bt63_11
Bt63_12
Bt63_13
Bt63_14
H99_01
H99_02
H99_03
H99_04
H99_05
H99_06
H99_07
H99_08
H99_09
H99_10
H99_11
H99_12
H99_13
H99_14
Type of repetitive sequence
Simple repeat
Others
Copy number variants
Deletion
Duplication
Features
Centromeres
MAT

Lineage: H99_Bt63  Sample: ZB_062524_S47

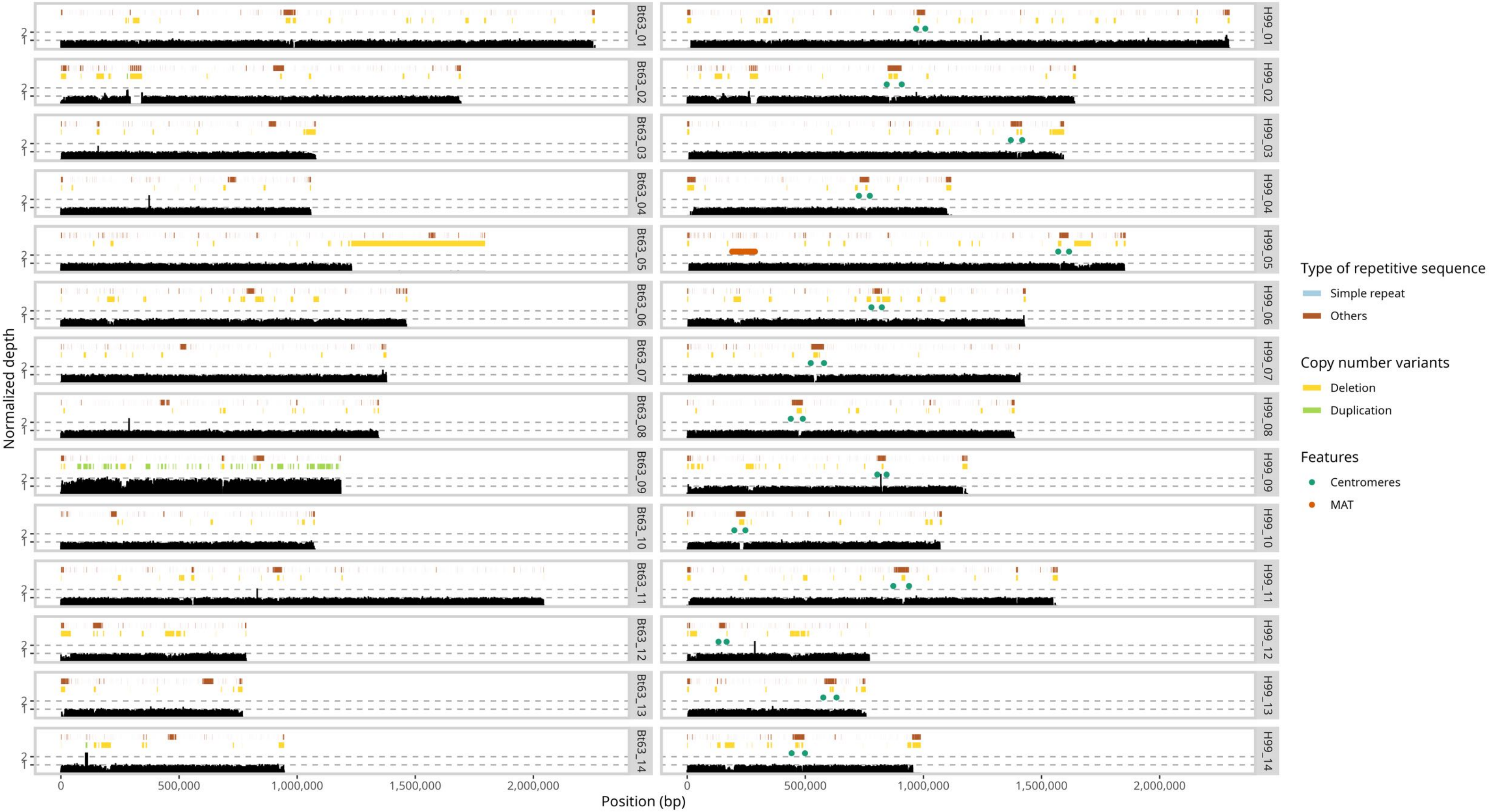

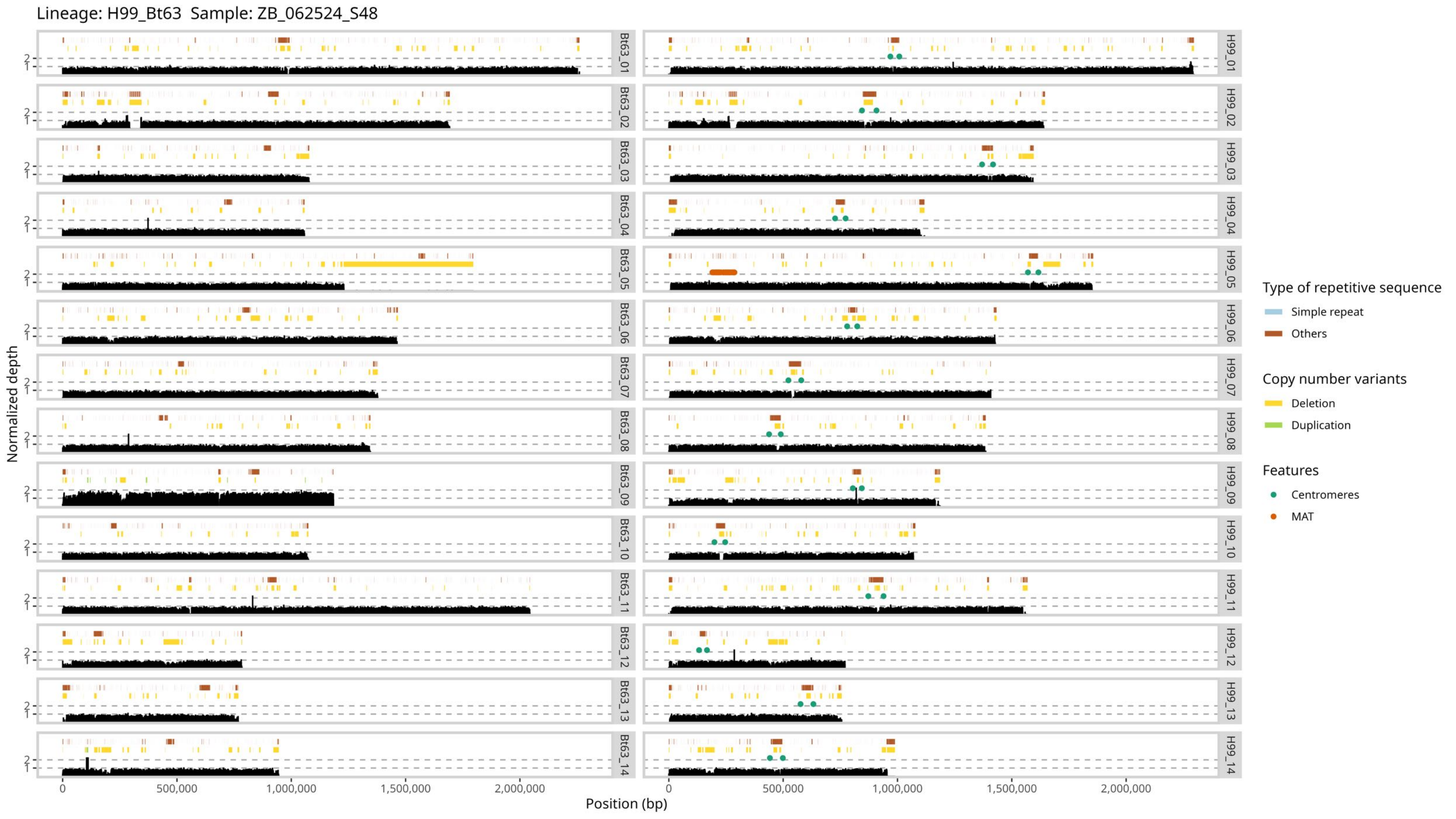

Lineage: H99_Bt63 Sample: ZB_062524_S48
Normalized depth
2
1
Bt63_01
Bt63_02
Bt63_03
Bt63_04
Bt63_05
Bt63_06
Bt63_07
Bt63_08
Bt63_09
Bt63_10
Bt63_11
Bt63_12
Bt63_13
Bt63_14
H99_01
H99_02
H99_03
H99_04
H99_05
H99_06
H99_07
H99_08
H99_09
H99_10
H99_11
H99_12
H99_13
H99_14
0
500,000
1,000,000
1,500,000
2,000,000
Position (bp)
Type of repetitive sequence
Simple repeat
Others
Copy number variants
Deletion
Duplication
Features
Centromeres
MAT

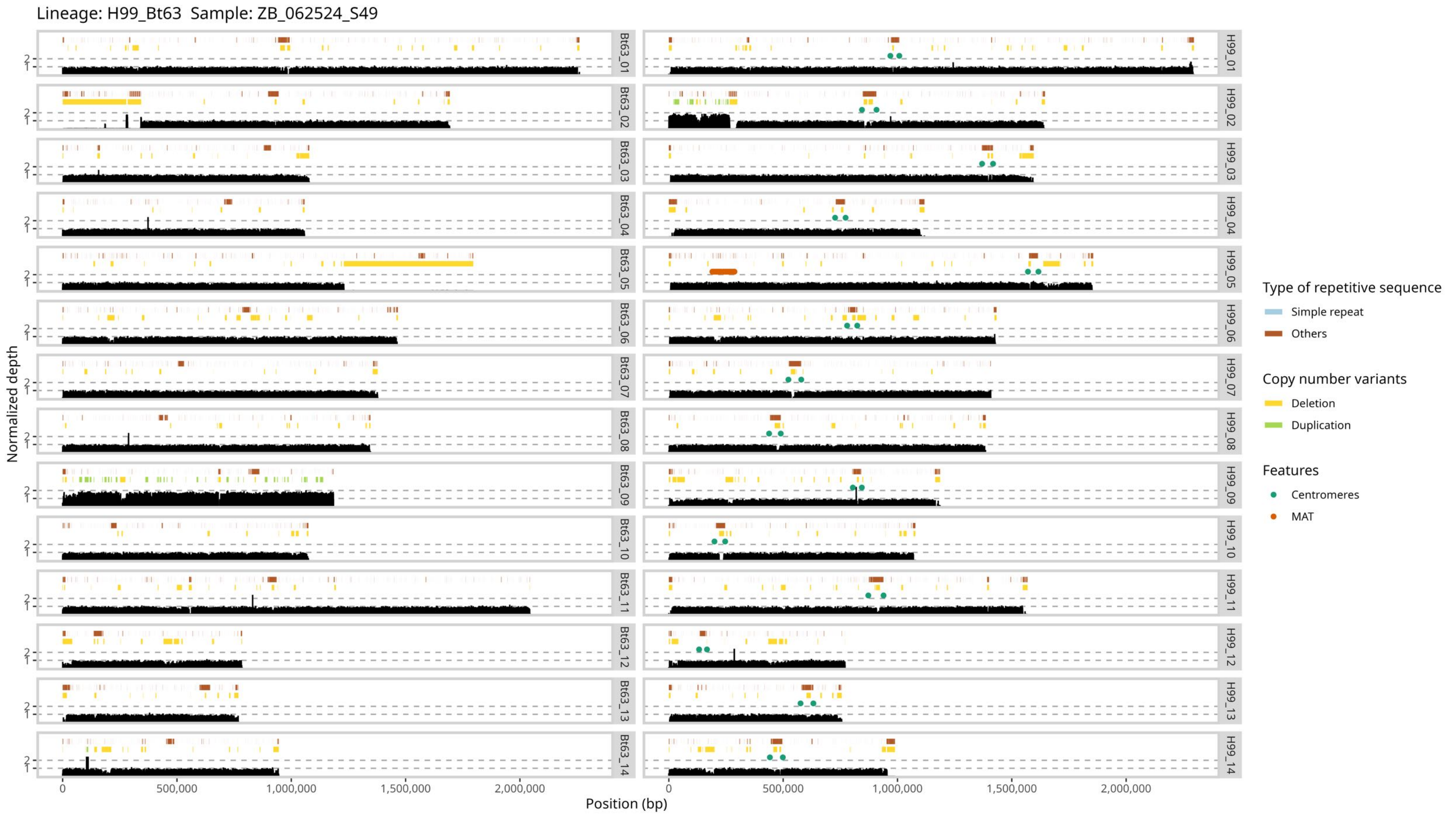
Lineage: H99_Bt63 Sample: ZB_062524_S49
Normalized depth
Position (bp)
0
500,000
1,000,000
1,500,000
2,000,000
Bt63_01
Bt63_02
Bt63_03
Bt63_04
Bt63_05
Bt63_06
Bt63_07
Bt63_08
Bt63_09
Bt63_10
Bt63_11
Bt63_12
Bt63_13
Bt63_14
H99_01
H99_02
H99_03
H99_04
H99_05
H99_06
H99_07
H99_08
H99_09
H99_10
H99_11
H99_12
H99_13
H99_14
Type of repetitive sequence
Simple repeat
Others
Copy number variants
Deletion
Duplication
Features
Centromeres
MAT

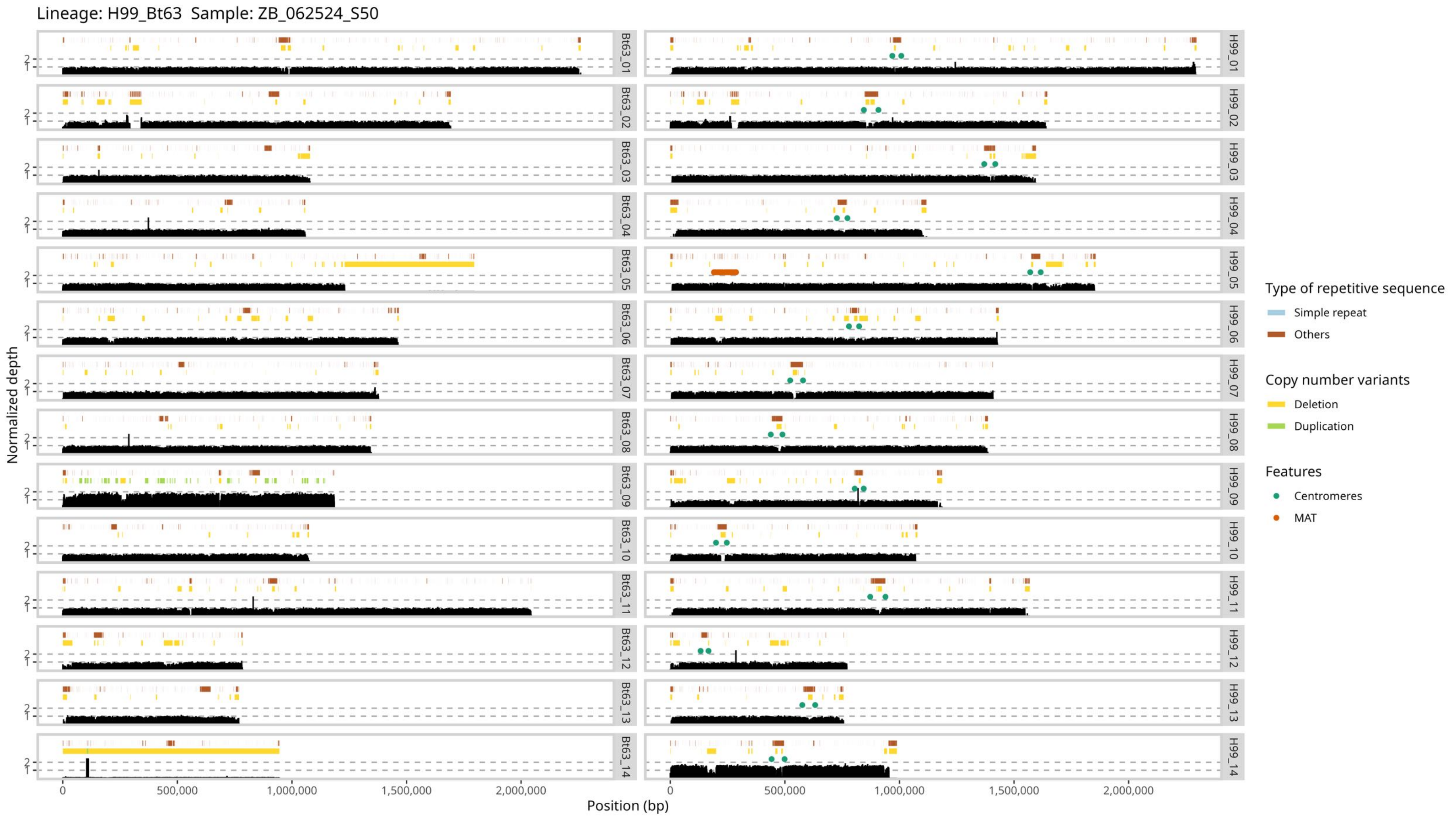

Lineage: H99_Bt63 Sample: ZB_062524_S50
Normalized depth
Position (bp)
0
500,000
1,000,000
1,500,000
2,000,000
Bt63_01
Bt63_02
Bt63_03
Bt63_04
Bt63_05
Bt63_06
Bt63_07
Bt63_08
Bt63_09
Bt63_10
Bt63_11
Bt63_12
Bt63_13
Bt63_14
H99_01
H99_02
H99_03
H99_04
H99_05
H99_06
H99_07
H99_08
H99_09
H99_10
H99_11
H99_12
H99_13
H99_14
Type of repetitive sequence
Simple repeat
Others
Copy number variants
Deletion
Duplication
Features
Centromeres
MAT

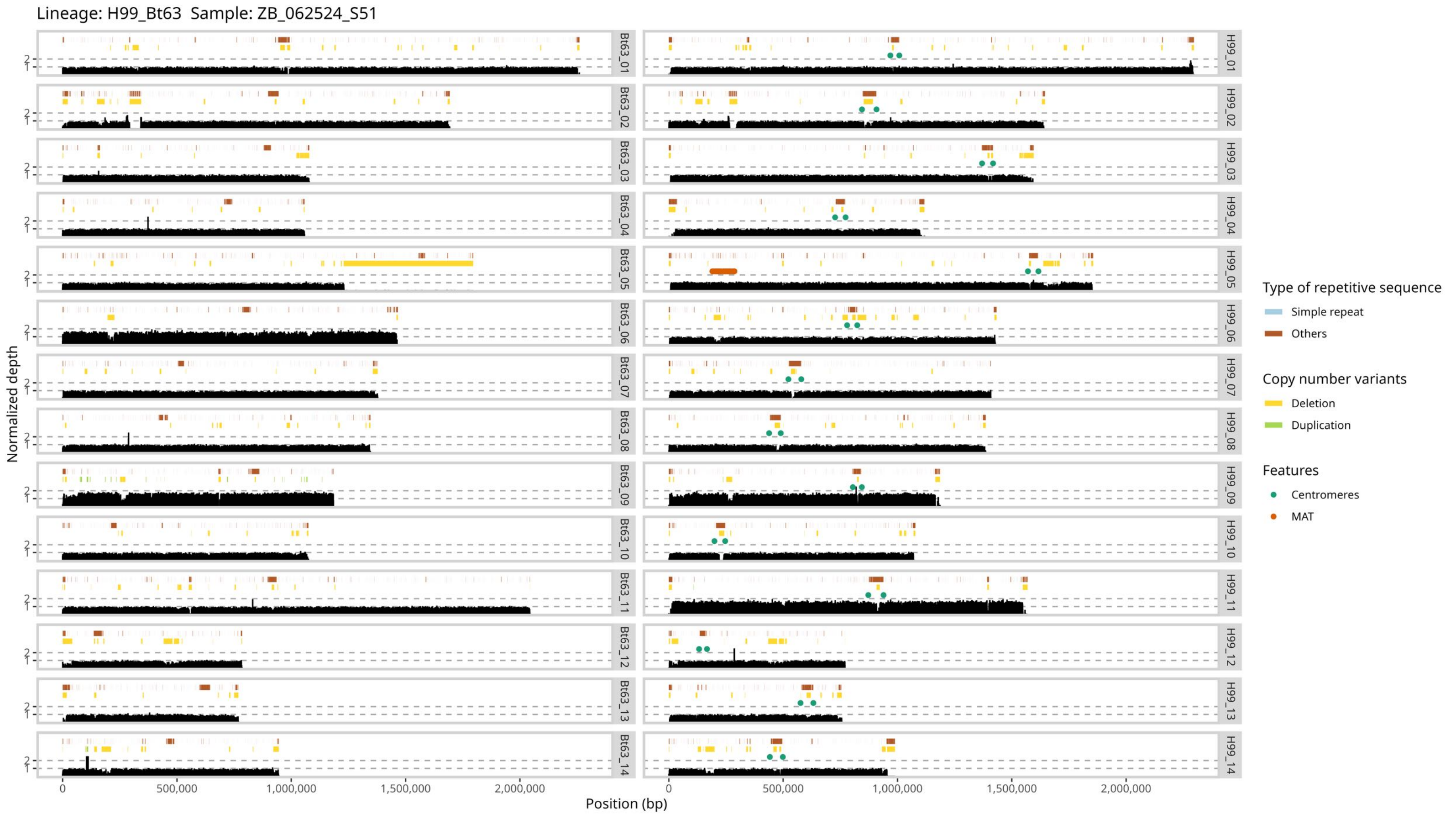
Lineage: H99_Bt63 Sample: ZB_062524_S51
Bt63_01
Bt63_02
Bt63_03
Bt63_04
Bt63_05
Bt63_06
Bt63_07
Bt63_08
Bt63_09
Bt63_10
Bt63_11
Bt63_12
Bt63_13
Bt63_14
H99_01
H99_02
H99_03
H99_04
H99_05
H99_06
H99_07
H99_08
H99_09
H99_10
H99_11
H99_12
H99_13
H99_14
Normalized depth
1
2
Position (bp)
0
500,000
1,000,000
1,500,000
2,000,000
Type of repetitive sequence
Simple repeat
Others
Copy number variants
Deletion
Duplication
Features
Centromeres
MAT

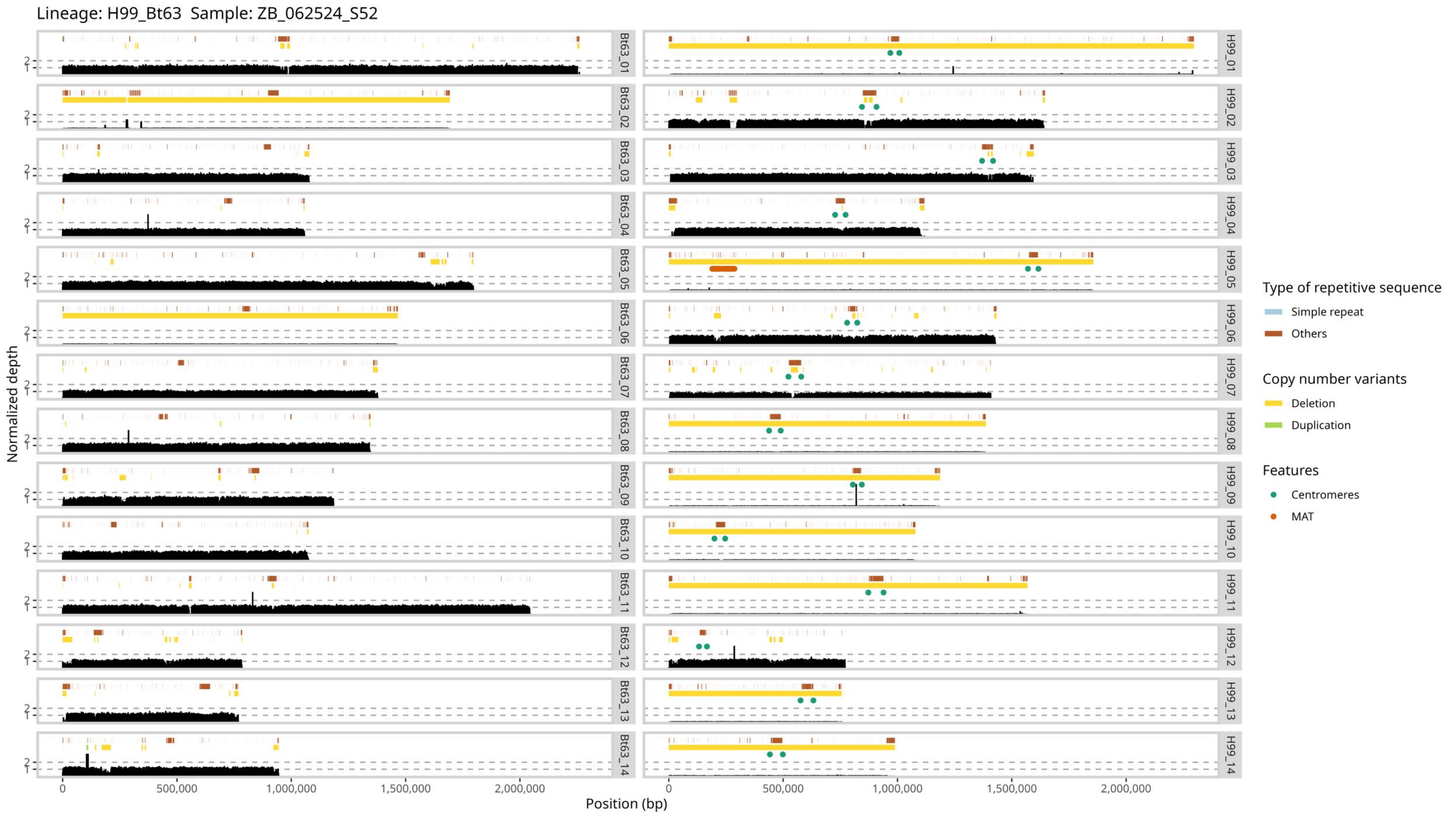

Lineage: H99_Bt63 Sample: ZB_062524_S52
Normalized depth
Position (bp)
Bt63_01
Bt63_02
Bt63_03
Bt63_04
Bt63_05
Bt63_06
Bt63_07
Bt63_08
Bt63_09
Bt63_10
Bt63_11
Bt63_12
Bt63_13
Bt63_14
H99_01
H99_02
H99_03
H99_04
H99_05
H99_06
H99_07
H99_08
H99_09
H99_10
H99_11
H99_12
H99_13
H99_14
0
500,000
1,000,000
1,500,000
2,000,000
Type of repetitive sequence
Simple repeat
Others
Copy number variants
Deletion
Duplication
Features
Centromeres
MAT

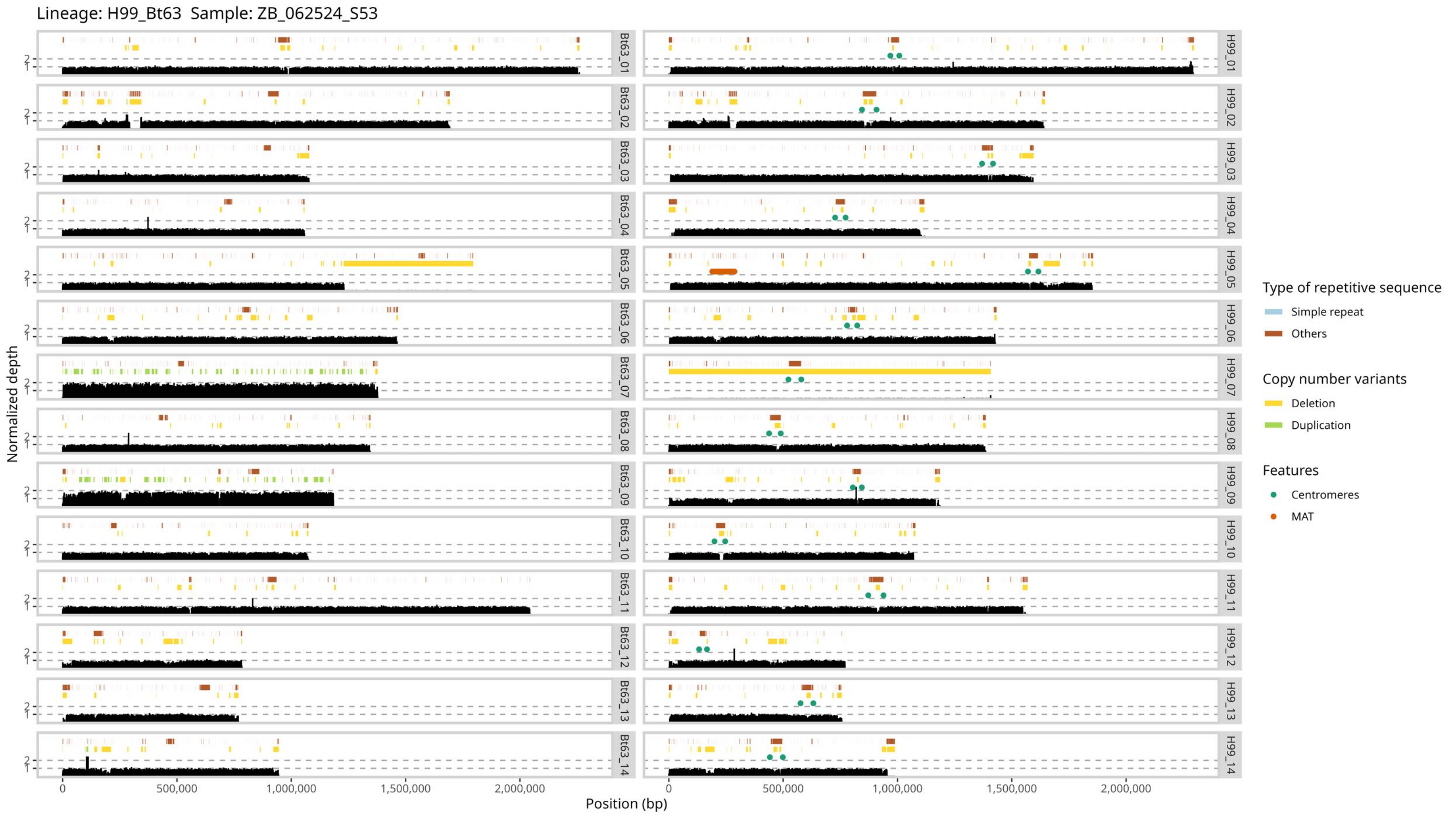
Lineage: H99_Bt63 Sample: ZB_062524_S53
Normalized depth
Position (bp)
0
500,000
1,000,000
1,500,000
2,000,000
1
2
Bt63_01
Bt63_02
Bt63_03
Bt63_04
Bt63_05
Bt63_06
Bt63_07
Bt63_08
Bt63_09
Bt63_10
Bt63_11
Bt63_12
Bt63_13
Bt63_14
H99_01
H99_02
H99_03
H99_04
H99_05
H99_06
H99_07
H99_08
H99_09
H99_10
H99_11
H99_12
H99_13
H99_14
Type of repetitive sequence
Simple repeat
Others
Copy number variants
Deletion
Duplication
Features
Centromeres
MAT
